# Ancient genomics support deep divergence between Eastern and Western Mediterranean Indo-European languages

**DOI:** 10.1101/2024.12.02.626332

**Authors:** Fulya Eylem Yediay, Guus Kroonen, Serena Sabatini, Karin Margarita Frei, Anja B. Frank, Thomaz Pinotti, Andrew Wigman, Rasmus Thorsø, Tharsika Vimala, Hugh McColl, Ioanna Moutafi, Isin Altinkaya, Abigail Ramsøe, Charleen Gaunitz, Gabriel Renaud, Alfredo Mederos Martin, Fabrice Demeter, Gabriele Scorrano, Alessandro Canci, Peter Fischer, Izzet Duyar, Claude Serhal, Alexander Varzari, Murat Türkteki, John O’Shea, Lorenz Rahmstorf, Gürcan Polat, Derya Atamtürk, Lasse Vinner, Sachihiro Omura, Kimiyoshi Matsumura, Jialu Cao, Frederik Valeur Seersholm, Jose Miguel Morillo Leon, Sofia Voutsaki, Raphaël Orgeolet, Brendan Burke, Nicholas P Herrmann, Giulia Recchia, Susi Corazza, Elisabetta Borgna, Mirella Cipolloni Sampò, Flavia Trucco, Ana Pajuelo Pando, Marie Louise Schjellerup Jørkov, Patrice Courtaud, Rebecca Peake, Juan Francisco Gibaja Bao, Györgyi Parditka, Jesper Stenderup, Karl-Göran Sjögren, Jacqueline Staring, Line Olsen, Igor V. Deyneko, György Pálfi, Pedro Manuel López Aldana, Bryan Burns, László Paja, Christian Mühlenbock, Claudio Cavazzuti, Alberto Cazzella, Anna Lagia, Vassilis Lambrinoudakis, Lazaros Kolonas, Jörg Rambach, Eugen Sava, Sergey Agulnikov, Vicente Castañeda Fernández, Mia Broné, Victoria Peña Romo, Fernando Molina González, Juan Antonio Cámara Serrano, Sylvia Jiménez Brobeil, Trinidad Nájera Molino, María Oliva Rodríguez Ariza, Catalina Galán Saulnier, Armando González Martín, Nicolas Cauwe, Claude Mordant, Mafalda Roscio, Luc Staniaszek, Mary Anne Tafuri, Tayfun Yıldırım, Luciano Salzani, Thorfinn Sand Korneliussen, J. Víctor Moreno-Mayar, Morten Erik Allentoft, Martin Sikora, Rasmus Nielsen, Kristian Kristiansen, Eske Willerslev

## Abstract

The Indo-European languages are among the most widely spoken in the world, yet their early diversification remains contentious^1–5^. It is widely accepted that the spread of this language family across Europe from the 5th millennium BP correlates with the expansion and diversification of steppe-related genetic ancestry from the onset of the Bronze Age^6,7^. However, multiple steppe-derived populations co-existed in Europe during this period, and it remains unclear how these populations diverged and which provided the demographic channels for the ancestral forms of the Italic, Celtic, Greek, and Armenian languages^8,9^. To investigate the ancestral histories of Indo-European-speaking groups in Southern Europe, we sequenced genomes from 314 ancient individuals from the Mediterranean and surrounding regions, spanning from 5,200 BP to 2,100 BP, and co-analysed these with published genome data. We additionally conducted strontium isotope analyses on 224 of these individuals. We find a deep east-west divide of steppe ancestry in Southern Europe during the Bronze Age. Specifically, we show that the arrival of steppe ancestry in Spain, France, and Italy was mediated by Bell Beaker (BB) populations of Western Europe, likely contributing to the emergence of the Italic and Celtic languages. In contrast, Armenian and Greek populations acquired steppe ancestry directly from Yamnaya groups of Eastern Europe. These results are consistent with the linguistic Italo-Celtic^10,11^ and Graeco-Armenian^1,12,13^ hypotheses accounting for the origins of most Mediterranean Indo-European languages of Classical Antiquity. Our findings thus align with specific linguistic divergence models for the Indo-European language family while contradicting others. This underlines the power of ancient DNA in uncovering prehistoric diversifications of human populations and language communities.

## Introduction

From 5,000 BP, large-scale human migrations significantly restructured the genetic makeup of human populations across Eurasia^6,7^. Various pulses of steppe-related ancestry, associated with the mobile pastoralists of the Early Bronze Age Yamnaya culture, spread across vast distances, leaving distinct cultural and genetic signatures. These migrations likely also played a key role in the prehistoric dispersal of the Indo-European language family. However, steppe ancestry reached the various regions of Western Eurasia by different mechanisms and at different times. In various historically Indo-European-speaking regions of Europe, the arrival of steppe ancestry was mediated by populations associated with the archaeological complexes of the Corded Ware (CWC) (5,000–4,350 BP)^6,7,14^ and Bell Beakers (4,800–4,300/3,800 BP)^15–17^. However, the extent to which similar dynamics occurred in the Eastern Mediterranean and adjacent Western Asian populations remains less well-understood. Moreover, while steppe ancestry has previously been detected in prehistoric Iberians^18,19^, Italians^20–23^, Greeks^24–27^, and Caucasians^24^, questions remain regarding the interrelatedness of the proximal source populations in the context of the Mediterranean region at large.

The spread of steppe ancestries is closely tied to the diversification of the Indo-European protolanguage into its historically attested subgroups^28^. In the Mediterranean, important Indo-European languages from Classical Antiquity include Gaulish, Latin, Greek, and Armenian, the latter being indigenous to the South Caucasus and Eastern Anatolia. For these, multiple competing phylogenetic linguistic models have been proposed^29^ (Linguistic Supplementary 2-4). The so-called Indo-Greek hypothesis groups Greek as well as the closely related Phrygian with Indo-Iranian^30^, while the competing Graeco-Armenian^12,13^ hypothesis posits that Greek forms a subclade with Armenian, possibly also including Albanian (“Balkan Indo-European”). Similarly, the Italic Indo-European subgroup, which is ancestral to Latin, has been variously grouped with Celtic and Germanic, giving rise to the traditionally popular Italo-Germanic^31^ and contrastive Italo-Celtic^10,32^ hypotheses. While relative linguistic consensus exists on the Graeco-Armenian and Italo-Celtic hypotheses, these are not unchallenged^33,34^. More fundamentally, the lack of a definitive linguistic consensus model for the Indo-European diversification constitutes a key obstacle to the interdisciplinary study of Indo-European language dispersal.

Here, we investigate the various sources of steppe ancestry along the entire northern border of the Mediterranean region, so as to establish the most parsimonious divergence model for the Indo-European languages in this area. We present new whole-genome data from 314 ancient individuals (>0.1X genomic coverage) from Southern and Central-Eastern Europe, as well as from the Eastern Mediterranean comprising Spain, France, Italy, Hungary, Moldova, Greece, Cyprus, Turkey, Syria, and Lebanon. These individuals mostly belong to the Bronze Age but span a time frame from 5,200 BP to 2,100 BP (Fig. 1) (Genetics and Strontium Supplementary S3; Supplementary Table S1). We also provide strontium isotope signatures from 224 individuals and radiocarbon-dating of 144 individuals (Supplementary Tables S8 and S9). Utilizing identity-by-descent (IBD) inferred admixture modelling^17,35^ with specific source populations, we obtained enhanced resolution of genetic ancestries, enabling us to differentiate diverse or common ancestries. Furthermore, we combined our strontium isotope data with our genetic results to deepen the understanding of mobility patterns over time.

**Fig. 1.**
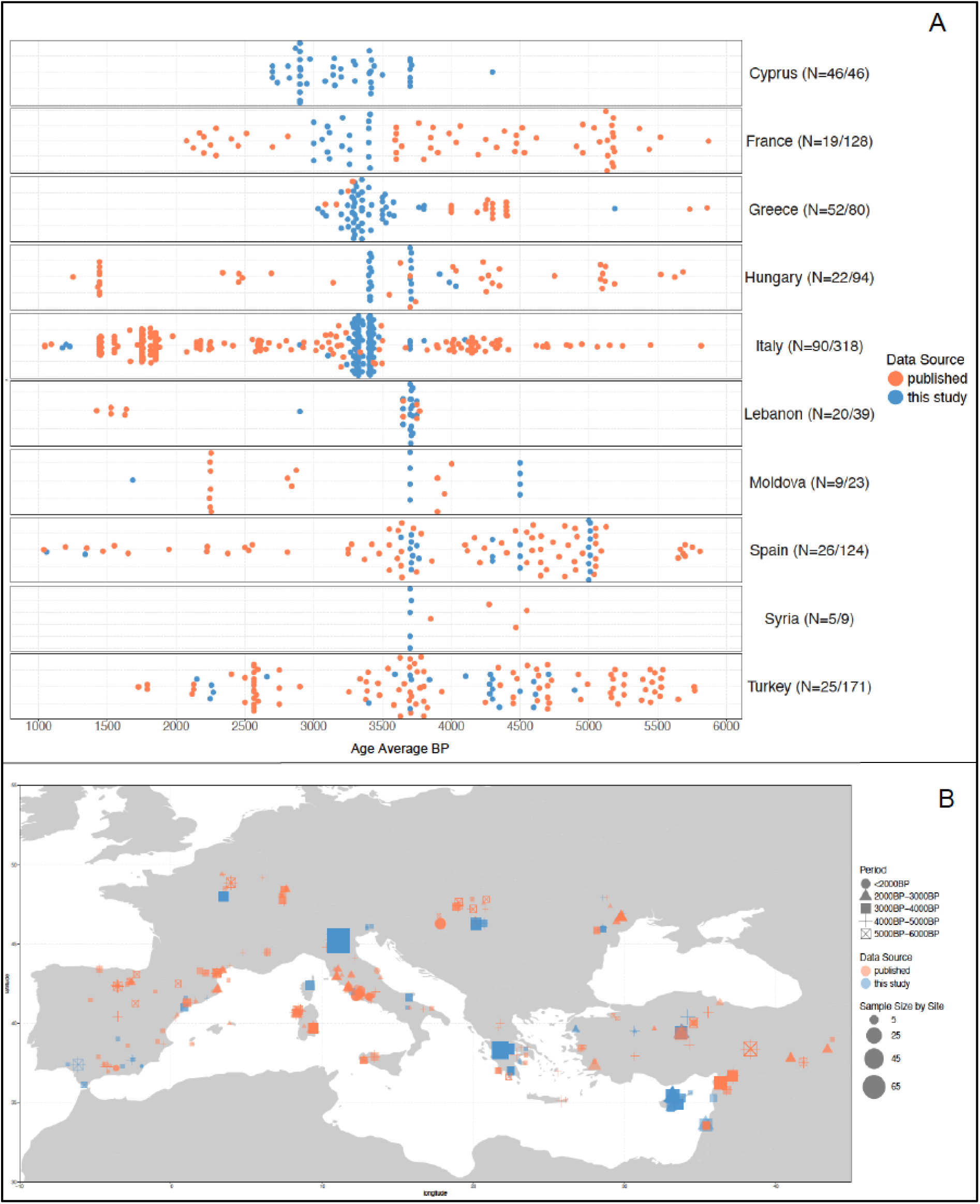
Distribution of ancient individuals distributed by country (A), (N= the number of individuals in this study/total number of individuals in dataset), and locality shown on the map (B). We only demonstrate the individuals limited by time frame (6,000–1,000 cal BP) to avoid overlapping.

### Population structure and overview of IBD clusters

We generated the dataset by shotgun-sequencing genomic DNA extracted from 314 ancient individuals (Genetics and Strontium Supplementary S2; S3). This was combined with 992 published shotgun-sequenced genomes and 1,097 genome-wide SNP-captured ancient individuals (Supplementary Table S2). These were all imputed using established cut-offs^17,35–37^ of 1X capture and 0.1X shotgun data resulting in a total combined dataset of 2,403 imputed genomes across 643,430 SNP (single nucleotide polymorphism) sites (Genetics and Strontium Supplementary S3). We called IBD segments between pairs of individuals using IBDseq^38^, and employed a network-based hierarchical approach^39^ to obtain fine-scale population clusters based on the total length of shared IBD segments between pairs of individuals (Genetics and Strontium Supplementary S5).

Through this clustering approach, we detected six deep clusters corresponding to distinct ancestries previously related to the prehistoric formation of Eurasia: “Farmer-related (Cluster 0_1)”, “European-Hunter-Gatherers (Cluster 0_2)”, “Caucasus – Iran (Cluster 0_3)”, “Steppe-related (Cluster 0_4)”, “Central Asia – Siberia (Cluster 0_5)”, “Moroccan-Hunter-Gatherers (Cluster 0_6)” (Extended Fig. 1 and Fig. 2; Supplementary Table S3; Genetics and Strontium Supplementary Table S5.1). We deeply investigated “Farmer-related (0_1)” and “Steppe-related (0_4)” clusters, since the ancient individuals reported here fell within a number of subclusters of these main clusters. The “Farmer-related (0_1)” cluster is further divided into four subclusters, distinguishing between East (0_1_2 and 0_1_3) and West Mediterranean (0_1_1 and 0_1_4). The “Steppe-related (0_4)” cluster comprises three subclusters with individuals from Central Asia and Europe, “Russia-Central Asia (0_4_1)”, “Bell Beaker-related (0_4_2)”, and “CWC-Yamnaya-related (0_4_3)” (Fig. 2; Genetics and Strontium Supplementary Table S5.2; Extended Data Fig. 3). The occurrence of these subclusters refers to the spread of Yamnaya-related ancestry both eastward and westward, as well as the formation of new genetic profiles throughout Europe and Central Asia^6,7,40–42^. When focusing specifically on the time frame of this study (between 5,000 BP and 3,000 BP), we observe a distinct pattern emerging between the 5th and 4th millennia in terms of the distribution of these Steppe-related clusters (Fig. 2). The distribution of the “CWC-Yamnaya-related (0_4_3)” cluster originating from the Pontic Steppe, extends into Central Eastern Europe and Northern Greece before the 4th millennium BP. This cluster persisted west of the Black Sea and in Armenia after the 4th millennium BP, whereas the “Bell Beaker-related (0_4_2)” cluster became prevalent in Europe. Further south to Greece, during this time we see a shift of the border between the “Steppe-related (0_4)” cluster and the “Farmer-related (0_1)” cluster. The first instance we detect of cross-border interactions occurs during an advanced phase of the 4th millennium BP. Here we find that two published Greece Bronze Age individuals^43,44^ (Log04 and G23) fall within the “CWC-Yamnaya-related (0_4_3)” cluster, specifically within subclusters corresponding to the “Yamnaya-related (0_4_3_1)” culture and the “Armenian-Caucasus (0_4_3_3)”, respectively (Genetics and Strontium Supplementary Fig. S5.9). At the east end of this border zone, individuals from Moldova fall within the subclusters of the “Steppe-related (0_4)” cluster, whereas all Eastern Mediterranean individuals including those from Greece fall within the subclusters of “Farmer-related (0_1)” cluster. Moreover, we detect the newly sequenced Early Bronze age individuals from Moldova within the subcluster of “Yamnaya-related (0_4_3_1)”, while the Middle Bronze Age individuals within the subcluster corresponding to “Corded Ware (0_4_3_2)” individuals like previously published Moldovan individuals^24^ (Supplementary Table S3). Additionally, we found all individuals from Anatolia, Cyprus, and Levant within the “Mediterranean (0_1_2)” cluster during the 3rd and 4th millennia, except one Early Bronze age individual and two European outliers from Cyprus are found within “Early farmers (0_1_3)” and “Bell Beaker-related (0_4_2)” throughout the 4th and 3rd millennia BP.

**Fig. 2.**
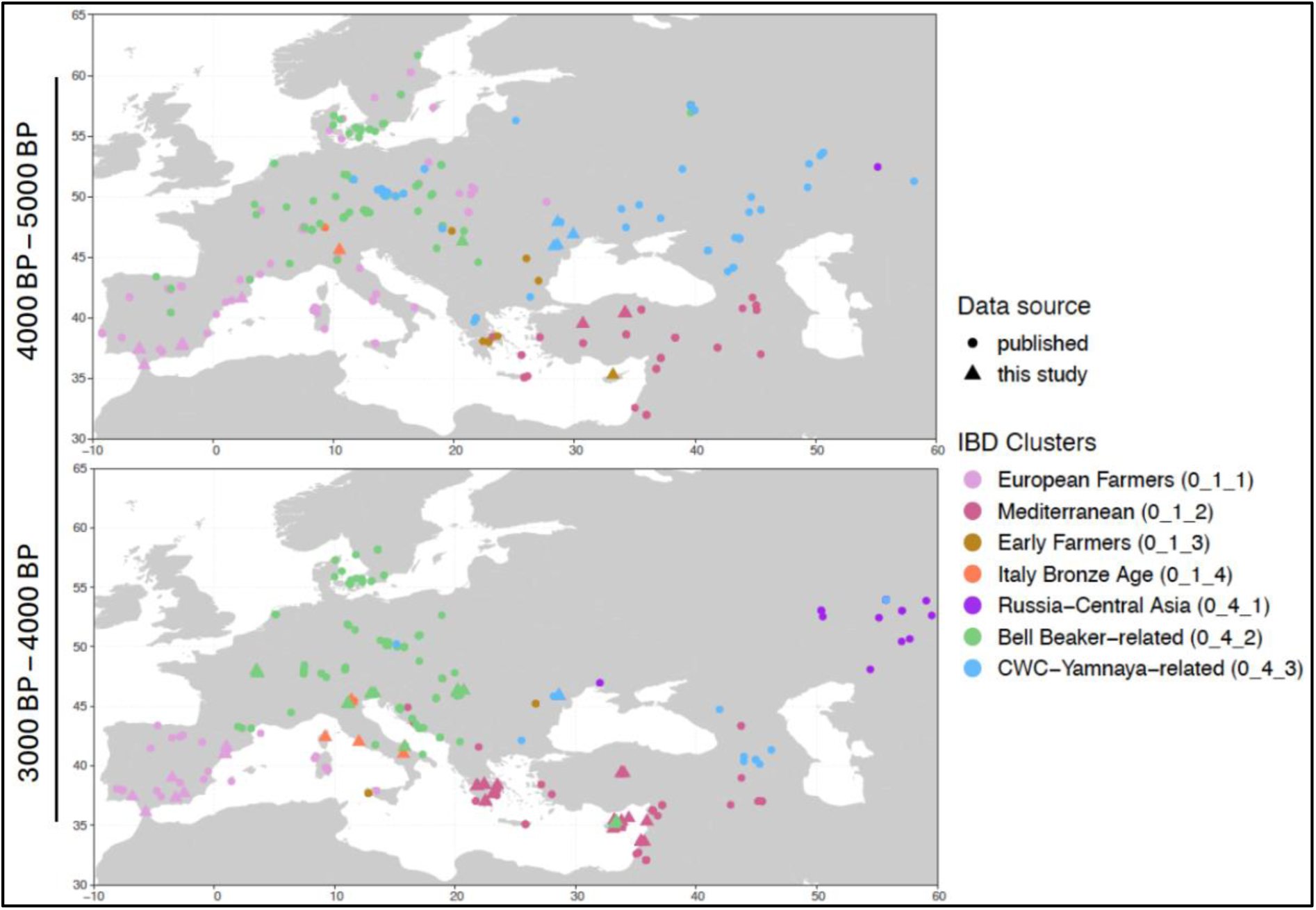
Geographical distribution of the IBD clusters of individuals in the 5th and 4th millennia BP.

### Southern and Central Eastern Europe

The spread of steppe-related ancestry in Europe has been traced to the Yamnaya populations and their subsequent admixture with local European populations after admixing with Globular Amphora culture-related (GAC-related) populations in the east^6,7^. To differentiate between closely-related steppe ancestries in Southern European populations, we first generated an ancestry “palette” for every ancient individual in the database, representing the individual sharing patterns with all clusters in the dataset (Genetics and Strontium Supplementary S6.2). By designating specific clusters as sources, we are able to employ IBD Mixture Modelling^17,35^ to model the palettes of target individuals as best-fitting combinations of the source palettes. Of relevance here were various steppe-related populations from the Pontic Steppe and Europe (Genetics and Strontium Supplementary S6.2). We built on from a basic source set^35^ that includes the Hunter-Gatherer and early Farming-related populations involved in the formation of European population structure, including a series of outgroups from Eurasia and Africa (Supplementary Table S4). We then progressively added a series of more recent populations as sources. By including two steppe sources, early Corded Ware culture (CWC) and Yamnaya (Genetics and Strontium Supplementary S6.2; Supplementary Table S4), we revealed distinctly dissimilar patterns, contrasting Greece with France, Spain, and Italy, reflecting the expansion of two separate steppe ancestries—one mediated through Corded Ware groups and the other derived from Yamnaya populations (Extended Data Fig. 4; Genetics and Strontium Supplementary S6.2). The distinction between these two expansions is also distinguishable by the Farming-related ancestry they carry. The arrival of steppe ancestry in Europe was accompanied by GAC-related Farming ancestry: this farming ancestry is present in the Corded Ware carrying GAC-related ancestry, but absent in Greece (Genetics and Strontium Supplementary S6.2; Fig. S6.18; Supplementary Table S5). We thus detect a clear division in steppe ancestry between the Eastern and Western Mediterranean.

By introducing a third steppe-related source from the “Bell Beaker-related (0_4_2)” cluster, Southwestern Europe shows ancestry that is BB-related, while that of Greece is Yamnaya-related, particularly in the Peloponnese (Figs. 3 and 4). The Balkans, however, exhibit mixed ancestries of CWC/BB and Yamnaya, suggestive of interactions between migrants from the Pontic Steppe and with local populations associated with CWC/BB-related ancestry, or an unsampled steppe source emerging west of the Black Sea (Fig. 4). During the Mycenaean period in Greece (or Late Helladic period, c. 3,700–3,200 BP), Yamnaya ancestry became widespread, even extending into the Eastern Mediterranean (Fig. 3). We interpret this as evidence of a direct Yamnaya migration into the Peloponnese, supported also by the detection of the typical Yamnaya Y-chromosome haplogroup lineage R1b-Z2103 (Z2108)^24,45^ in two individuals from Ayios Vasileios, one from the Mycenaean cemetery Voudeni, and one from Kirrha (Fig. 5, Supplementary Table S7). However, the dominant Y-haplogroup lineage in Greece, J2a-Y7011, is also detected in individuals from Lapithos, Iron Age Cyprus, who are additionally characterized by non-local strontium isotope signatures. This haplogroup is also found in Szőreg, Hungary, suggesting a connection between the Balkans and the Eastern Mediterranean, as this lineage is present in early Balkan populations. To identify a potential connection with the Balkans, we present new data from Middle and Late Bronze Age Hungary revealing a complex population structure. We observe a mixed pattern of BB and Yamnaya components in Hungary during the Bronze Age (Fig. 4; Genetics and Strontium Supplementary S6.3). This suggests either another source population for those individuals or an admixture between two groups.

**Fig. 3.**
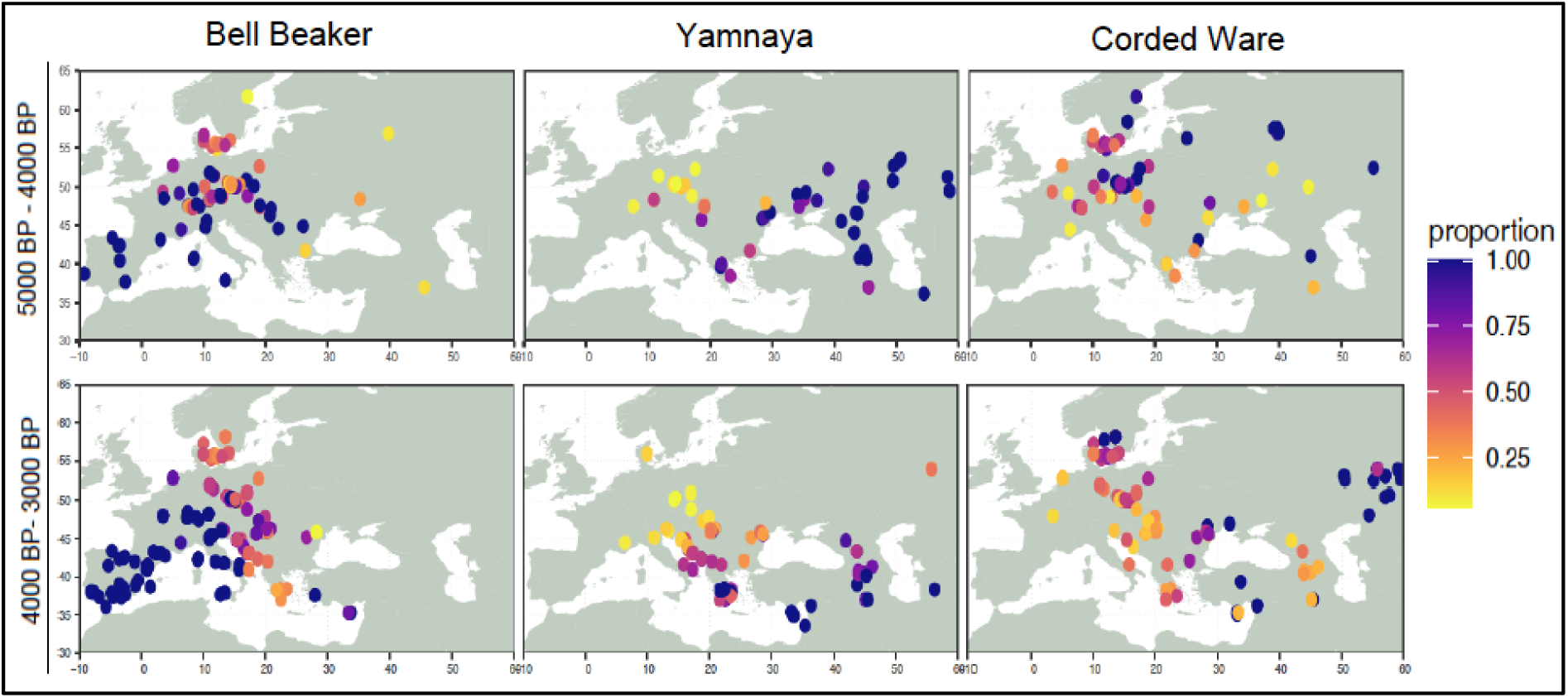
Distribution of Bell Beaker-derived and Yamnaya-derived ancestry proportions obtained from the IBD admixture model. The proportion of each steppe source is standardized by the total steppe contributions, i.e. the sum of Corded Ware, Bell Beaker and Yamnaya_Samara contributions.

**Fig. 4.**
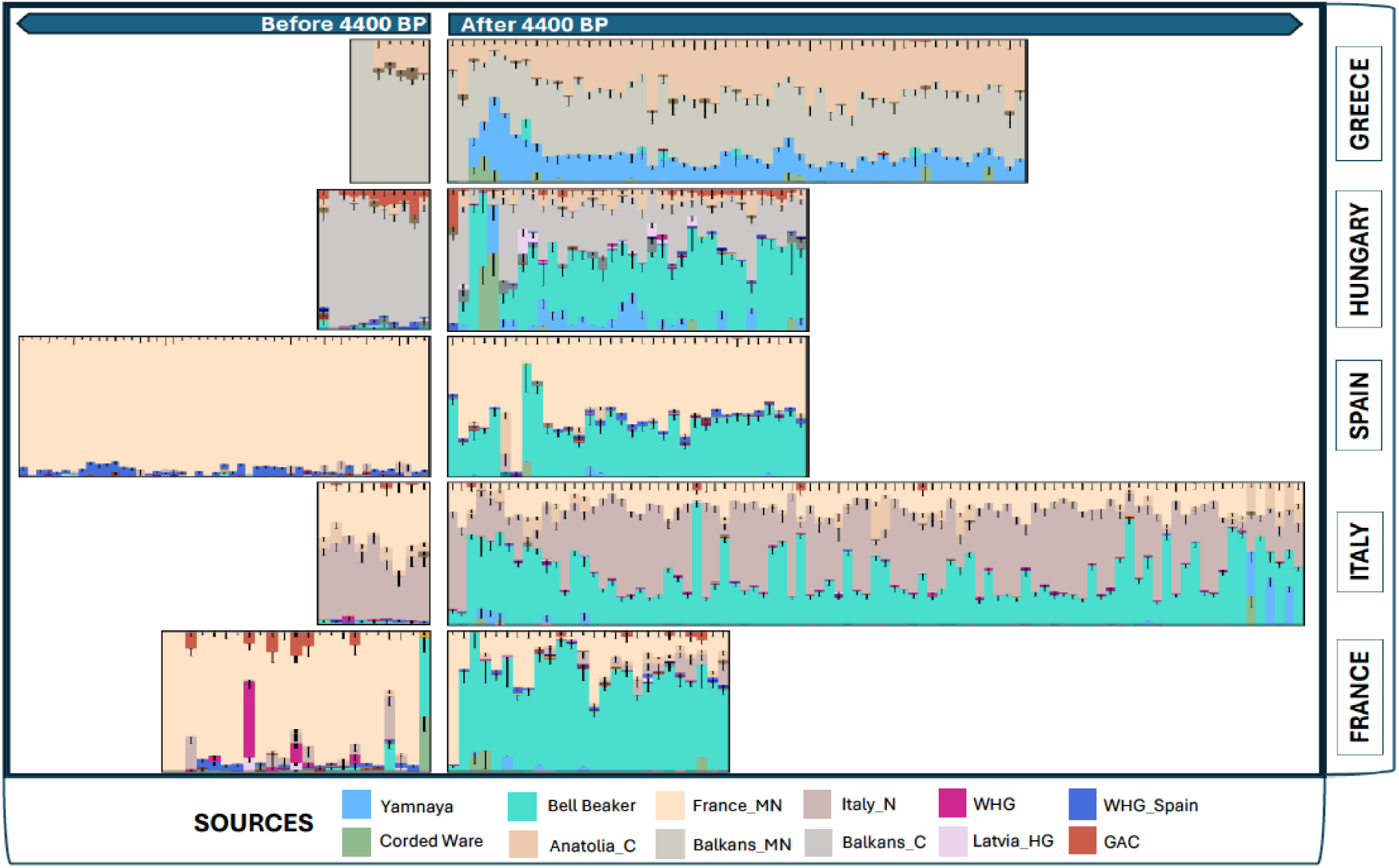
Ancestry bar plots generated for each individual using source population proportions of IBD admixture modelling sorted by time BP and divided into two time series, before and after 4,400 BP, illustrating a Southern Europe split (Italy, France, and Spain vs Greece).

**Fig. 5.**
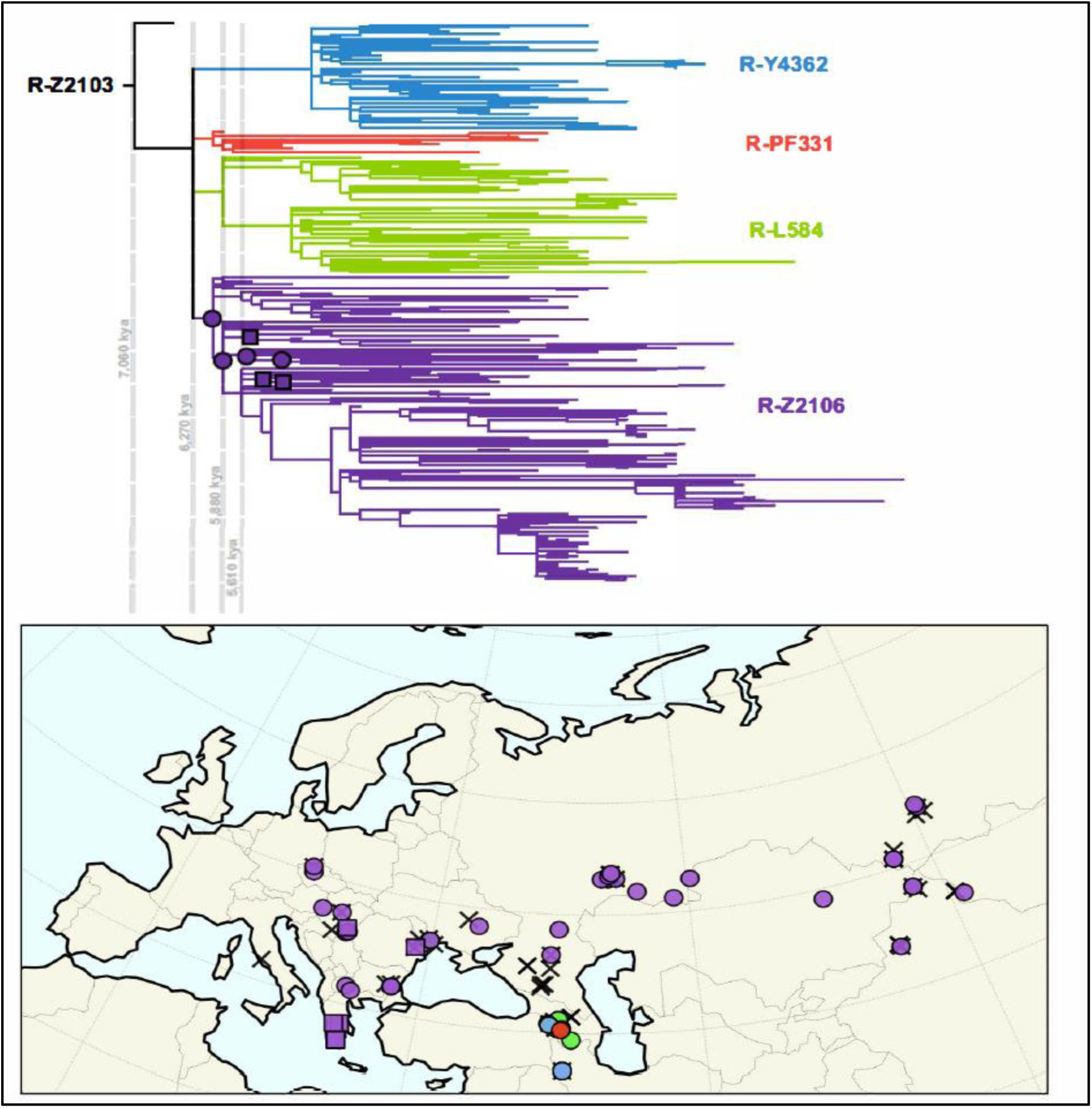
Phylogeography of Y-chromosome haplogroup R-Z2103. Phylogram of haplogroup R-Z2103, with branch lengths proportional to SNP number, built from a dataset of unique variants in private datasets. The haplogroup resolves in a four-way polytomy. We plotted all occurrences older than 2,000 years ago in the ancient DNA record in Western Eurasia on an Albers Equal Area map, colour-coded by clades downstream of R-Z2103. Only haplogroup R-Z2106 extends beyond the Caucasus and Northern Iran, and we indicate phylogenetic position of individuals in the tree. Crosses mark R-Z2103 individuals with uncertain clade assignment due to low coverage, and squares indicate individuals from Greece, Moldova and Hungary generated in this study. Dotted lines denote split time estimates of key haplogroups, calculated using rho statistics.

Italian Bronze Age individuals show at least three distinct admixture patterns. The first is similar to the Bronze Age Bell Beaker-related groups of France and Spain. Interestingly, that group includes all individuals from Central Italy and Corsica, a few individuals from Olmo, and one published Bell Beaker-context individual from Northern Italy^15^ (I2478, 4,006 BP). A second group, primarily from Olmo, shows increased Neolithic farmer ancestry with lower amounts of steppe ancestry, suggesting an ongoing local admixture process. A third limited group with increased Yamnaya ancestry, similar to Balkan and Greek Bronze Age populations, is observed in individuals from the Adriatic coast of Italy (Fig. 4; Genetics and Strontium Supplementary S6.2, CGG_2_100646, NEO806, R1)^6,20^. Interestingly, three Early Bronze Age individuals from Northeastern Italy carry an additional CWC or Yamnaya component, and such a profile is also observed in the individuals from Hungary (Fig. 4; Genetics and Strontium Supplementary S6.3). These individuals, found in monumental tumulus graves and a coffin burial (CGG_2_022653), likely held a special social or cultural status, as suggested by the complexity of the tumuli and the ritual activities linked to those monuments^46^ (Archaeological Supplementary 2.7.4; 2.7.9–11).

To compare the BB source with local contribution, we added two Early Bronze Age individuals from Northeastern Italy as an ancestral source to model Late Bronze Age Italian individuals (Genetics and Strontium Supplementary S6.2, Fig. S6.13; Supplementary Table S5). We found this new, local source to better model the ancestry component previously modelled by the BB source, except in a few individuals who show a mixed pattern of Bell Beaker and local ancestry, rejecting the model. This local source also replaced the steppe proportion in Corsica and Sicily, suggesting a maritime spread of steppe ancestry through the Mediterranean (Genetics and Strontium Supplementary Fig. S6.15). This ancestry also increased in Late Bronze Age individuals from Migennes, France (Genetics and Strontium Supplementary S6.2; Fig. S6.15), suggesting later formation in steppe ancestry in Western Europe could also connect the Balkans and Italy.

### Eastern Mediterranean

The Caucasus region has experienced a complex history of human migrations, interactions, and cultural exchanges, marked by admixture among diverse population groups. With the spread of farming to Iran and the Caucasus, populations in the region were shaped by admixture between early Neolithic farmers from the Fertile Crescent and two genetically similar population groups, Iran Neolithic and Caucasus Hunter-Gatherers (CHG)^24,41^. Additionally, the expansion of the Kura-Araxes culture during the 5th millennium BP connected the Caucasus with the Levant and Mesopotamia through extensive trade networks^47^. Interaction between Anatolia and the Caucasus increased during the Chalcolithic and the Bronze Ages, leading to the spread of CHG ancestry. It also diffused into the Mediterranean, an early indication of which is found in Anatolian farmer groups from Tepecik-Çiftlik^21,25,43,44^.

Here, we report new genomic data from 25 individuals from Anatolia, including individuals from Resuloğlu, an Early Bronze Age cemetery associated with Hattian culture, from Western Anatolia (Küllüoba, Keçiçayırı and Antandros), and from Central Anatolia (Kaman Kalehöyük), representing both the Bronze and Iron Ages. To assess the population structure of Anatolian individuals, we applied IBD mixture modelling, removing the CWC and BB sources, and instead progressively added a series of eastern ancestry sources from Iran and Caucasus Chalcolithic individuals, also including a Central Anatolian farmer source from Tepecik (Genetics and Strontium Supplementary S6.4; Supplementary Table S4). In our base modelling, all Chalcolithic and Bronze Age genomes from Anatolia appeared as mixtures of three components: local Neolithic farmers (from Tepecik), CHG (Caucasus Hunter-Gatherer) and a small proportion of Iranian Neolithic ancestry, with the exception of a minor proportion of EHG (Eastern Hunter-Gatherer) found in some individuals from Kalehöyük, Arslantepe, and Western Anatolia (Genetics and Strontium Supplementary Fig. S6.25; Supplementary Table S5). Together with CHG, this proportion was slightly elevated in the Iron Age. To understand the origin of the CHG and EHG contributions, we added Chalcolithic individuals from Iran and the Southern Caucasus as geographically proximal sources (Supplementary Table S4). The model revealed that Anatolian Chalcolithic and Bronze Age individuals received various proportions of ancestry from both Iran and the Southern Caucasus, with a higher amount of Iranian ancestry in Eastern than Western and Central Anatolia (Fig. 6). Moreover, the Chalcolithic Caucasus proportion increased in Central Anatolia, such as in Kaman Kalehöyük, in the Middle Bronze Age, and Iranian ancestry is minimal or absent in Western Anatolia. Admixture patterns in Eastern Anatolian individuals resemble those of Bronze Age individuals from Lebanon. Notably, Chalcolithic and Bronze Age individuals from Arslantepe, who received both Caucasus and Iranian ancestries, played a pivotal role within the Kura-Araxes region, spanning from Southern Caucasus to Levant.

**Fig. 6.**
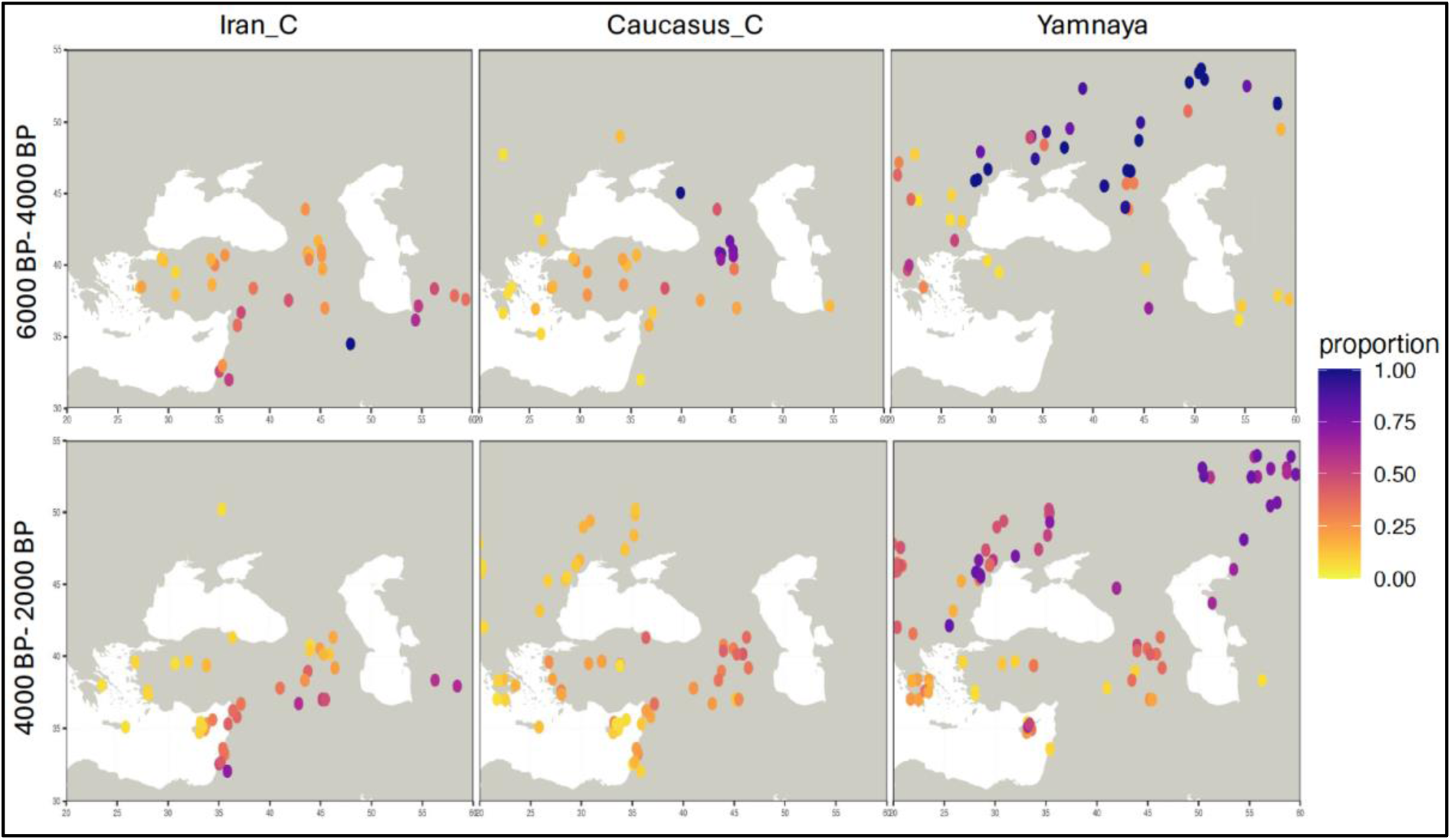
Distribution of Iran Chalcolithic (Iran_C), Caucasus Chalcolithic (Caucasus_C) and Yamnaya-related ancestry proportions obtained from the IBD admixture model.

To distinguish increased proportions of steppe ancestry in the Iron Age, we included multiple steppe sources (Yamnaya, CWC, BB) that revealed different signatures depending on the geographical location. In the newly sequenced Iron Age samples from Central and Northwestern Anatolia (Kalehöyük, Antandros and Keçiçayırı), we observed minor proportions of steppe ancestry with the pattern found in Balkans/Greek Late Bronze Age and probably reflects migrations from the Balkans (Genetics and Strontium Supplementary Fig. S6.37; S6.38; S6.39; Supplementary Table S5). Given that the individual from Keçiçayırı (CGG_2_022162) was unearthed from the Phrygian valley, the appearance of this ancestry may be associated with the emergence of the Phrygian state during the late 4th millennium BP^48^ (Archaeology Supplementary 2.12.5; Linguistic Supplementary 3.3).

To trace the interaction between Eastern Mediterranean and Southern Europe, we sequenced Bronze Age and Iron Age individuals from Cyprus and Lebanon. Cypriot individuals make up a highly diverse group connecting all of the Eastern Mediterranean (Fig. 6). Our results suggest that Cyprus, and in particular its coastal towns, were a genetic and cultural “melting pot” during the Bronze Age. The Middle and Late Bronze Age individuals from Cyprus show a genetic pattern similar to that of individuals from Lebanon and Eastern Anatolia Bronze Age, while one (the earliest) individual from Karavas (CGG_2_022531), dated to 5,000– 4,500 BP (Archaeology Supplementary 2.2.3), show extra early Anatolian farmer ancestry. A long-distance genetic connection is observed at Hala Sultan Tekke, one genetic outlier carrying Balkan/Aegean ancestry similar to that of Late Bronze Age Greece individuals. Thus, our dataset suggests close contacts of Cyprus with the Levant and the Aegean during the Bronze Age and even earlier periods. During the subsequent Iron Age, data from both Lapithos and Amathus suggest an increasingly uniform population across the island. Moreover, Iron Age genomes show a formation similar to populations of the Aegean and Western Anatolia Iron Age, carrying a small proportion of Yamnaya ancestry which reflects Greece ancestry (Fig. 6). Additionally, there is genetic evidence of long-distance interaction with Northern Europe, as seen in a Scandinavian genetic outlier (CGG_2_022535) from a rock-cut tomb at Vounous Bellapais, excavated by the Swedish-Cyprus expedition and dated to c. 4,000–3,800 BP. (Genetics and Strontium Supplementary Fig. S6.45, Archaeology Supplementary 2.2.7). This outlier clusters with Scandinavian Bronze Age individuals and, intriguingly, this origin is also supported by the Y-haplogroup I1 and by a non-local highly radiogenic strontium isotope signature compatible with some parts of Scandinavia (Genetics and Strontium Supplementary S10; Supplementary Table S8). The implications of this observation are not conclusive, since we do not have radiocarbon dating from this individual. Due to its rich copper sources, Cyprus maintained extensive trading networks with most of the Mediterranean; this situation is well mirrored by the presence of Anatolian, Levantine, Greek, and European ancestries in the Cypriot genomes.

### Increasing mobility towards the end of the Bronze Age

Considering the highly dynamic Bronze Age period in the Mediterranean, we analysed strontium isotope ratios from 224 ancient individuals across Cyprus, Greece, Italy, and Spain to trace the mobility of individuals (Genetics and Strontium Supplementary S10; Supplementary Table S8). We identified 56 individuals as potential non-locals and compared them to their genetic profiles.

In Greece, nine individuals from Kirrha, Voudeni, Eleon, and Apollo Maleatas returned non-local strontium isotope signatures. These might have travelled from regions within Greece where strontium isotope baselines are slightly more radiogenic. Among these nine individuals, we only have genetic data for four individuals, as several closely-related individuals were removed to avoid skewing the IBD admixture analysis. Genetically, these four individuals are similar to others from the same area, except for one individual from Eleon, who carries a small proportion of Bell Beaker ancestry reflecting a connection with Central Eastern Europe. Interestingly, two individuals (from Kirrha and Apollo Maleatas, Genetics and Strontium Supplementary S10; Supplementary Table S6) with non-local signatures have second-degree relatives who fall within the local baseline, indicating different mobility patterns within the same family. However, the relatives falling within the local baseline could also have originated from a different region given the large overlap in strontium signatures across Greece (Genetics and Strontium Supplementary S10; Supplementary Table S8). Finally, the father of a non-local individual from Apollo Maleatas (CGG_2_023933) differs from other Late Bronze Age Greece individuals by having a higher Yamnaya proportion compare to Late Bronze Age individuals, similar to the Middle Bronze Age individuals (>∼3,800 BP).

In Cyprus, despite the high levels of genetic variation, only four Cypriot samples are characterized by non-local strontium isotope signatures (Supplementary Table S8). The genetic outlier from Vounous-Bellapais (CGG_2_022535) with unusual Scandinavian ancestry is also confirmed by the strontium isotope results. Another non-local Iron Age individual (CGG_2_022526) exhibits an admixture signature consistent with Greece Late Bronze Age (Genetics and Strontium Supplementary Fig. S6.45; S6.46). In contrast, the remainder of the outliers resemble populations from the Anatolia/Levant Bronze Age (Genetics and Strontium Supplementary S10). For many genetic outliers in which strontium data is also presented, long-term mobility was not detected (Genetics and Strontium Supplementary S10).

We additionally identified several individuals with non-local strontium isotope signatures within Italy, consistent with a highly dynamic society. These individuals come from Olmo, Pian Sultano, Scalvinetto, and from Coppa Nevigata. One individual (CGG_2_101264) from Pian Sultano had a bone and a tooth sample investigated for strontium isotopes, returning one signature (tooth) within their local strontium isotope baseline and another (petrous) outside of it. These results suggest that she did not originate from the area of Pian Sultano, but moved there in her early childhood/adolescence (Genetics and Strontium Supplementary S10). Yet another interesting non-local (CGG_2_022591), from Olmo, carries the highest proportion of steppe-related ancestry found among Italian Bronze Age individuals, displaying a different composition farmer and steppe proportion than other Italian Bronze Age individuals, carrying no Italian Neolithic ancestry. Another non-local from Coppa Nevigata (CGG_2_100646) genetically can be distinguished from the other Italian individuals by carrying Yamnaya-related steppe ancestry and Mediterranean farmer proportion similar to the Aegean/Balkans. Overall, non-local individuals of Bronze Age Italy cannot be distinguished from each other in terms of the steppe source obtained from IBD admixture modelling except for two cases from Coppa Nevigata and Olmo (CGG_2_100646 and CGG_2_022591; Genetics and Strontium Supplementary Fig. S6.13). Moreover, most of the non-local individuals in Olmo come from a group among Bronze Age Italians that exhibit a lower proportion of steppe-related ancestry, thus resembling individuals from Mediterranean islands (Sicily, Corsica)(Genetics and Strontium Supplementary Fig. S6.13; S6.15).

In Spain, we identified individuals with non-local strontium isotope signatures genetically belonging to two different groups: one with local farmer-related ancestry, and the other group dated to the Early Bronze Age (∼4,300 BP) showing Bell Beaker-related ancestry. Since we observe only small-scale variation among farmer populations in Spain, the outliers with farmer ancestry may have originated from other farmer sites within the region.

## Discussion

To elucidate the genetic formation and infer the divergence of the Indo-European language family, we investigated the contribution of potential ancestral source populations to the wider Mediterranean region, including from areas in which the Italic, Celtic, Greek, and Armenian languages are historically spoken. The genetic results obtained by IBD admixture modelling with putative steppe-related source populations support a deep divergence between Eastern and Western Mediterranean Indo-European-speaking groups through the detection of Yamnaya-related and Bell Beaker-related steppe ancestry respectively (Fig. 4). This divergence supports the linguistic hypotheses on the existence of the so-called Graeco-Armenian and Italo-Celtic subclades. In contrast, it disqualifies the rival cladistic hypotheses known as Indo-Greek and Italo-Germanic, the steppe ancestry among the populations of historically Germanic- and Indo-Iranian-speaking areas previously having been characterized as primarily Corded Ware-related^35^.

### South-central Europe: Italic

Prior to its Romanization, Italy was characterized by marked linguistic complexity, harbouring multiple Indo-European and non-Indo-European language groups^49^. The Italic languages, including Latin, Oscan, Umbrian, and possibly Venetic, are Indo-European in origin, just like the more distantly related Cisalpine Gaulish, Messapic, and Greek. Etruscan and Rhaetic, on the other hand, belong to the so-called Tyrrhenian family (Linguistic Supplementary 2). Due to this complexity, tracing the spread of the Italic group, which eventually came to dominate the Italian Peninsula, is notoriously difficult^50^. Archaeological interpretations have variously linked the spread to the Remedello culture, the Rinaldone culture, the Terramare culture, and the Proto-Villanovan as well as the Villanovan cultures^51^. However, the oldest direct linguistic evidence for the presence of Italic is first found in Old Latin and Umbrian inscriptions from around 2,650 BP.

Previous genetic studies have dated the arrival of steppe ancestry in Northern Italy to around 4,000 BP and in Central Italy by 3,600 BP^21^. In our data, steppe ancestry appears in Central Italy a century earlier, in two newly reported individuals from Latium (Pian Sultano, CGG_2_101264, CGG_2_101266), the eventual epicentre of the Latin language. According to our IBD admixture modelling, the steppe ancestry of these individuals, along with that of published Late Bronze Age individuals from Grotta Regina Margherita and Toppo Daguzzo, is characterized as genetically Bell Beaker rather than Yamnaya-related, similar to historically Celtic-speaking populations in Western Europe. This, as well as the prevalence of this ancestry throughout the Bronze and Iron Age, is consistent with the linguistic Italo-Celtic hypothesis.

### The Eastern Mediterranean: Greek

The timing and trajectory of the entry of the Greek language into Greece are traditionally much-debated topics^2,25^ (Linguistic Supplementary 4.1). In Late Bronze Age Greece, writing emerges in the form of the Linear A Minoan (c. 3,800–3,400 BP) and Linear B Mycenaean (c. 3,350–3,150 BP) syllabary scripts^52^. In Cyprus, it appears with the Cypro-Minoan syllabary (c. 3,500–3,300 BP), a local variant of Linear A, and the derived Cypriot syllabary (c. 3,000–2,300 BP)^53^. While the Linear A text corpus remains largely undeciphered, Linear B as well as the Cypriot syllabary have been shown to represent the earliest written evidence of Greek^54^.

The arrival of steppe ancestry has previously been documented in individuals from Northern Greece as early as ∼4,200 BP, suggesting a connection to the Pontic Steppe^44,55,25,43^. Using IBD admixture modelling, we establish that the Steppe source in Greece originates directly from Yamnaya-related populations, and differs from CWC populations who are formed with Yamnaya and GAC and widespread across most of Europe. Notably, the proportion of this Yamnaya-related ancestry is higher in the individuals before ∼3,800 BP, including a previously unpublished male from Argolis (Apollo Maleatas, CGG_2_022928-23929), dated to around 3,800 BP. This provides the hitherto clearest evidence for the intrusion of steppe-derived, potentially Greek-speaking groups into the Peloponnese. The appearance of steppe ancestry here thus predates the oldest direct evidence of Greek in the form of the Linear B script, by which time it had stabilized at lower levels^43^. At the same time, the preliterary interactions with local populations find a possible analogue in the absorption of so-called “Pre-Greek” vocabulary^56^, reflecting contact of Greek with non-Indo-European language varieties.

In Cyprus, the Arcadocypriot Greek dialect was spoken from at least the Early Iron Age, next to Phoenician and one or more unknown languages attested through the Cypro-Minoan and Cypriot syllabaries^57^. A diverse population is also suggested by genetic links between LBA and EIA Cyprus with the Levant and the Aegean. Steppe ancestry is identified in a number of unpublished individuals from Hala Sultan Tekke (CGG_2_022123, CGG_2_022924) and from Lapithos (CGG_2_022517), reflecting an affiliation with Late Helladic (i.e. Mycenaean Age) populations of the Peloponnese. This aligns with the appearance of Mycenaean pottery imports in the Late Bronze Age^58^ and with the linguistic classification of Arcadocypriot as a descendant of the same South Greek dialect as Mycenaean, in contrast to other Greek dialects such as Doric and Ionic^54,59^. However, one other individual from Lapithos (CGG_2_22488), although buried 4,100–4,000 BP, already clusters with LBA Greeks, suggesting Pre-Mycenaean connectivity with steppe-impacted populations of the Aegean.

### The Caucasus and Eastern Anatolia: Armenian

The Armenian language, attested from c. 1,550 BP, is historically spoken in the South Caucasus and Eastern Anatolia (Linguistic Supplementary 4.3). Its earliest presence there has previously been estimated to date to around 3,100 BP^51,60^. During the Late Iron Age, most of the region was part of the Urartian Kingdom. This state was culturally diverse^61^ and likely also contained an Armenian linguistic element, as evidenced by the exchange of vocabulary between Urartian and an early form of Armenian^62^. This is potentially supported by the presence of steppe ancestry among the published individuals from Urartian and pre-Urartian contexts^26^.

Steppe ancestry has previously been detected in the South Caucasus from the Middle Bronze Age^24^, coinciding with the transition from the Kura-Araxes culture to the Trialeti culture by the end of the 5th millennium BP. We can now demonstrate that these individuals, as well as those from Urartian contexts, received steppe ancestry from the same, western Yamnaya population as 4th millennium BP individuals from the Aegean (Extended Data Fig. 6, Genetics and Strontium Supplementary Fig. S6.19; S6.21). These findings support the linguistic Graeco-Armenian hypothesis and suggest that the linguistic precursor of Armenian was introduced to the Caucasus by the end of the 5th millennium BP^24,63^.

### Archaeological implications

Bell Beaker populations exerted a pronounced genetic and cultural impact in Iberia, whereas the situation in Italy is more complex. In Northeastern Italy, Bell Beaker groups appear to have arrived in relatively small numbers, with some individuals receiving prominent tumulus burials. However, during the Early Bronze Age and at the transition to the Middle Bronze Age, evidence emerges of a connection between Central Europe and Italy in the distribution of triangular daggers, often found in hoards^64,65,66^. These daggers are distributed across Italy, with their appearance coinciding with the widespread diffusion of Bell Beaker-related ancestry across the Italian Peninsula between 3,800 and 3,500 BP.

In the Terramare region, population growth significantly exceeded local development, suggesting an influx from surrounding areas^67,68^. Archaeological evidence suggests connections with Hungary and regions north of the Alps^69–72^, although these links only become evident in genetic ancestries later, during the Iron Age. This suggests that Northern Italy functioned as a cultural “melting pot”, which would later impact other Italian regions. After 3,200 BP, a noticeable exodus is archaeologically documented, particularly from the southern part of the Terramare region in the Po Valley^67,73^.

Further south along the Adriatic coast, three genomes from the Central and Southern Italian litoral (CGG_2_100646; NEO806; R1) reveal affinities with Balkan populations, reflecting sustained contacts between both sides of the Adriatic Sea during the 5th and 4th millennia BP^74–76^. These contacts extended beyond the Adriatic, reaching as far as the Aegean, as indicated by the presence of Mycenaean pottery and other goods in settlements along the Italian coast^77^. Genetic ancestries further suggest small-scale movements of people, possibly involving specialized craftspeople and traders, or potentially driven by exogamy practices, contributing to the observed genetic admixture in the region.

Greece reveals a different picture. A distinct steppe proportion was introduced into both the mainland and the Peloponnese around 3,800 BP, subsequently becoming prevalent in burial sites over the ensuing centuries. The arrival of Mycenaean culture, along with the Greek language, has been subject to divergent interpretations, ranging from gradual, peaceful infiltration to more rapid acquisition of political control^78–80^. Neither of these models are contradicted by the new genetic evidence: following the collapse of Early Helladic society, the brief Middle Helladic period saw new migrants from the north. These eventually consolidated political control with the advent of the Late Helladic period and the formation of Early Mycenaean Culture around 3,700 BP^81^. The process also entailed a cultural and linguistic encounter between newcomers and the residing population. Over time the original steppe signal diminished due to admixture with local populations with farmer ancestry. Archaeological evidence documents sustained contacts with the genetic source region in Moldova and the surrounding Carpathian region, as reflected in the introduction of steppe horses and chariots^82^, as well as the trade in silver from Carpathian mines^83^.

Thus, a long-standing debate regarding the origins of Mycenaean culture can be resolved, at least in part. Genetic links with Anatolia and Crete persisted, mirroring the cultural influences that shaped the formation of Mycenaean society. Moreover, Mycenaean culture exhibits striking similarities with the slightly earlier Trialeti culture of the Southern Caucasus^68^, which likely contributed to processes behind the subsequent Armenian ethnogenesis. In the South Caucasus, we see the rise of new Bronze Age elites in the mineral-rich regions of present-day Georgia and Armenia. These elites had links to both steppe chariot traditions and the Hittite civilization in Anatolia. Trialeti burial inventories, characterized by monumental tumulus chamber burials featuring precious imports, exhibit parallels with both Hittite city-states and early Mycenaean shaft graves. This suggests the formation of new commercial and military networks linking steppe societies and Near Eastern civilizations^68^, as now also corroborated by the genetics.

The genetic evidence from Cyprus underscores its significant role as a trade hub owing to its abundant copper resources from Chalcolithic until the Iron Age. Moreover, this evidence highlights the island’s strong ties to Western Anatolia and the Aegean. The Late Bronze Age marked the peak of Cypriot copper mining and trade across the Mediterranean, fostering a flourishing international culture^84,85^. Also, these connections are mirrored in the genetic evidence, which reveals links between Cyprus and both the Eastern and Western Mediterranean regions. By the early 4th millennium BP, an archaeological connection additionally existed between the Únětice culture and the Eastern Mediterranean, including Cyprus, as exemplified by the presence of Únětice ring ingots and dress pins^81^. This connection is potentially supported by genetic evidence from a single individual, although its historical significance remains enigmatic.

## Conclusion

For the first time, we are able to coherently combine evidence from ancient DNA, strontium isotopes, linguistics and archaeology to support a dual model for the divergence and dispersal of the Italic, Greek and Armenian languages during the Bronze Age. Specifically, we show that the arrival of steppe ancestry in Spain, France and Italy was mediated by Bell Beaker populations of Western Europe, likely contributing to the emergence of the Italic and Celtic languages. In contrast, Armenian and Greek populations acquired steppe ancestry directly from Yamnaya groups of Eastern Europe. These results are consistent with the linguistic Italo-Celtic and Graeco-Armenian hypotheses accounting for the origins of most Mediterranean Indo-European languages of Classical Antiquity. In contrast, however, our results fail to support the competing Italo-Germanic and Indo-Greek hypotheses, as the steppe ancestry among the populations of historically Germanic- and Indo-Iranian-speaking areas has been characterized as primarily Corded Ware-related. Our findings thus align with specific linguistic divergence models for the Indo-European language family while contradicting others. This underlines the power of ancient DNA in uncovering prehistoric diversifications of human populations and language communities.

## Supporting information

Genetics and Strontium Supplementary

Linguistic Supplementary

Archaeological Supplementary

## Methods

### Data generation

All steps of generating the ancient genomes were conducted in dedicated ancient DNA lab facilities at Lundbeck Centre for GeoGenetics, University of Copenhagen, using well established protocols within ancient DNA^17,35^ (Genetics and Strontium Supplementary S1). Drilling was performed manually, if possible, on both petrous bone and tooth. DNA extractions and library builds were carried out in the ancient DNA clean lab both manually and with automatisation. Double stranded “USER” and “non-USER” libraries^86^ were built and sequenced on the Illumina Hiseq 4000 and the Novaseq 6000. non-USER were evaluated for the purpose of authenticating ancient reads, to allow for the detection of post-mortem damage^87^. Where possible, USER treated libraries were built from the authenticated extracts to minimise the effects of post-mortem damage on downstream analysis^88^ (Supplementary Table S1). We merged the USER and non-USER libraries as described in Genetics and Strontium Supplementary S2. All reads were mapped to the human reference genomes (build hs37d5) using bwa aln (0.7.17)^89^, and removed the adapters using AdapterRemoval (2.3.2)^90^, duplicates by using picard MarkDuplicates (2.25.0). to ensure the authenticity of the sequenced data, we performed mapDamage2.0 (v2.2.1)^87^ (Supplementary Table S1).

Contaminated libraries were identified using contamix^91^, schmutzi^92^ and for the X chromosome contamination level, we ran ANGSD^93^. The total 380 samples yielded DNA ranging from 0.004X to 15.5X in autosomal coverage, 314 samples out of total samples provided DNA coverage >0.1X which were used for downstream analysis (Supplementary Table S1).

### Imputation

We imputed^36^ a total 2,403 ancient samples (merged whole genome shotgun and capture), 314 newly sequenced whole genome shotgun samples by following McColl et al. 2024^35^ and Allentoft et al 2024^17^ (Genetics and Strontium Supplementary S3).

### Principal component analysis (PCA)

To get an overview of the basic structure of our data, we carried out a principal component analysis (PCA) on an imputed dataset, comprising a whole set (2,228 ancient individuals) and a subset (1,837 ancient individuals) from both shotgun and capture (1,240K) data using Plink (v1.90b6.21) (Genetics and Strontium Supplementary S4). We first restricted the SNPs to capture sites, and applied MAF (0.05) filtering, resulting in a dataset with 580,130 SNPs. We visualized the main IBD clusters by colouring for all ancient genomes used for clustering (Extended Data Fig. 2) and focused on the subclusters of “Farmer-related (0_1)” and “Steppe-related (0_4)” in the PCA plot of a subset individuals (Extended Data Fig. 3).

### IBD clustering and modelling

filtered the merged VCF files INFO>0.5, MAF (minor allele frequency) 0.01 and restricted to 1,240K capture SNP sites. To run IBDseq, we included the samples with >0.1x coverage for shotgun, >1x coverage for capture, and >0.90 average genotype probability across 643,430 SNP (single nucleotide polymorphism) sites^38^. We ran IBDseq following the implementation in McColl et al. 2024 on ancient samples within a broad geographical area and period (300 BP–45,000 BP)^17,35,37^. After removing close relatives, we ran IBD admixture models with a total of 2,228 ancient individuals, of which 274 were newly sequenced Bronze Age Mediterranean individuals Prior to running Leiden network-based hierarchical clustering, we applied filters by removing the IBD segments of less than 1 cM, LOD score of less than 3, and hotspot regions. We also removed one pair of the first and second-degree relatives to minimize small clusters formed only with close relatives. We then ran the clustering with 2,228 ancient individuals by setting a minimum total shared IBD of 5 cM and a permutation of 200.

IBD admixture modelling is a method combining allele-matching profiling^94^ and chromopainting^95^ using the shared IBD length instead of allele frequency to get a better resolution for distinguishing especially genetically similar populations. Before aggregating the IBD sharing, we filtered out IBD segments of less than 1 cM, with no limit to the upper bound of total shared segments^17,35^. The models and the source individuals for IBD Admixture modelling were given in the Supplementary Table S4.

### Genetic sex determination

We estimated the depth of coverage for the individuals with newly generated data using pysam, by counting and measuring the length of the reads (MQ > 30) and dividing the sum by the reference contig length of chromosomes 1–22, X and Y. Because the Y-chromosome presents large regions of repetitive sequence not mappable using short-read sequencing technologies^96–98^, we restricted all analyses to the 10 Mb single-copy region defined in Poznik et al 2013^97^. We called chromosomal sex for all individuals in the dataset by calculating the ratio of the depth of coverage of X to the autosomes, Y to the autosomes, and Y and X chromosomes.

### Relatedness

To identify relatedness, we ran NGSRelate (v2)^99^ on the imputed dataset, calculating allele frequency by using only our samples since we have a dense population structure from Eurasia (Genetics and Strontium Supplementary S7).

### Y-chromosome analysis

We used bcftools^100^ mpileup and call functions to call genotypes within the 10 Mb accessible region of the Y-chromosome^97^. We excluded indels, triallelic positions, and genotypes that were not called in more than 95% of the population of non-clonal reads. To determine haplogroups, we matched ancestral and derived calls to the ISOGG 2019–2020 database using an in-house script that generates haplogroup paths in a root-to-tip manner. Those paths are ranked by the number of supporting variants – while also distinguishing C to T in forward and G to A in reverse strands – and then manually verified.

### Mitochondrial DNA analyses

We re-aligned the newly sequenced ancient DNA reads to the revised Cambridge Reference Sequence (rCRS) for the human mitochondrial DNA sequence using *bwa aln v. 0.7.17*^89^ *(options: -l10000)* and filtered for reads with a mapping quality of minimum 30 using SAMtools v. 1.17^100^. We then reconstructed consensus sequences of the mitogenomes with *bcftools*^101^ *v. 1.18* using *mpileup* (*options: --no-BAQ*) to obtain read pileups along the reference sequence, which were then inputted to *bcftools call (options: --multiallelic-caller -- ploidy 1)* for haploid genotype calling. We kept variants covered by at least five reads and a genotype quality above 25. We generated the final consensus sequences with *bcftools consensus.* The reconstructed mitogenomes were aligned with *mafft*^102,103^ *v.7.490,* while we restricted the phylogenetic analysis to the coding region located at 577–16,023 base pairs (rCRS coordinates). We carried out a Maximum Likelihood (ML)-based phylogenetic tree analysis with *RAxML-NG*^104^ *v. 1.2.2* under the substitution model GTR+I+G4 *(options: --all - -bs-trees 100)*.

### Sr isotope analysis

We performed Sr isotope analysis on 232 skeletal samples (139 teeth and 93 petrous bones) from 224 ancient individuals from Cyprus, Greece, Italy, and Spain (Supplementary Table S8). A diamond-tipped dental drill was used to cut a clean enamel sample (1-2 mg) from the tooth samples and to drill 1-2 mg of sample from the densest part of the otic capsule of the petrous bones. The tooth and bone samples were dissolved using a 1:1 solution of 0.5 ml 6M HCl and 0.5 ml 30% H2O2. Selected samples were spiked with a ^84^Sr-enriched tracer to determine Sr concentration via isotope dilution (ID).

The Sr column separation was done according to the methods of Frei et al^105^, using disposable 1 ml pipette tips fitted with pre-cleaned filters and charged using 200 µl pre-cleaned SrSpec™ resin (50–100 mesh; Eichrome Inc./Tristchem) as disposable extraction columns. The prepared samples were dissolved, loaded onto the columns, and washed using 3M HNO3, before the Sr was collected using mq. All Sr concentrations and isotope measurements were performed at the University of Copenhagen using A VG Sector 54 IT mass spectrometer equipped with eight Faraday detectors.

## Data availability

Sequence data were deposited in the ENA under accession: xxxxxxxx

## Methods References

References 17, 35–36, 86–105

## Acknowledgements

The Lundbeck Foundation GeoGenetics Centre is supported by grants from the Lundbeck Foundation (R302-2018-2155, R155-2013-16338), the Novo Nordisk Foundation (NNF18SA0035006), the Wellcome Trust (WT214300), Carlsberg Foundation (CF18-0024), the Danish National Research Foundation (DNRF94, DNRF174), the University of Copenhagen (KU2016 programme) and Ferring Pharmaceuticals A/S, to E.W. This work was further supported by the Swedish Foundation for Humanities and Social Sciences grant (Riksbankens Jubileumsfond M16-0455:1) and funded by the European Research Council (ERC) under the European Union’s Horizon 2020 research and innovation programme (Grant agreement no. 95138) to K.K., G.K, K.G.S, K.M.F. and F.E.Y. A.M.W., F.E.Y., G.J.K. and R.T. received support from the European Research Council (Starting Grant no. 716732). We are grateful for the support of the Carlsberg Foundation Semper Ardens research grant (CF18-0005) “Tales of Bronze Age People” and the research grant “Tales of Bronze Age Women” (CF-15 0878) both to K.M.F. and F.E.Y. S.S. and G.S. were supported by Riksbankens Jubileumsfond, grant P22-0641 Italy before Rome. J.V.M. is supported by the European Research Council (101078151) and VILLUM FONDEN (VIL53099). R.N. received support from NIH R35 GM153400-01. We would like to thank to Sapienza Università di Roma - progetti di Ricerca e Grandi Scavi, Regione Friuli Venezia Giulia; Fondazione Friuli; Università di Udine (projects “Dai tumuli ai castellieri: 1500 anni di storia in Friuli”, “Protostoria della media pianura friulana”), The M.H. Wiener Laboratory for Archaeological Science, American School of Classical Studies at Athens, Sapienza Università di Roma; Soprintendenza Archeologia Belle Arti Paesaggio per le province di Foggia, Andria – Barletta, Soprintendenza Archeologia Belle Arti Paesaggio Friuli Venezia Giulia, Japanese Institute of Anatolian Archaeology, Kaman Kalehoyük Archaeology Museum, Balikesir Kuva-yi Milliye Archaeology Museum, Eskisehir Eti Archaeology Museum, Çorum Archaeology Museum, Ministry of Culture and Tourism in Republic of Turkey for providing access to archeological samples. We thank Henrik Bjerre Hansen, Maria Madrona, Lærke Daniela Kjærsgaard Hansen, Tina-Louise Marie Mortensen, Marcus Andreas Behnke Hjorth, Andreas Bak Pørksen and Nils August Thomasen for their assistance.

## Author contributions

K.K., E.W., F.E.Y, K.M.F. initiated the study. F.E.Y., K.M.F. (Strontium part), K.K. and E.W. led the study. F.E.Y., G.K., K.M.F., K.K. and E.W. conceptualised the study. G.K., L.V.I., K.M.F., L.O., R.N., M.S., K.K. and E.W. supervised the research. G.K., K.M.F., R.N., K.K. and E.W. acquired funding for research. F.E.Y., S.S., M.E.A., F.D., I.M., A.M.M., P.F., I.D., A.V., M.T., G.P., J.M.O., S.O., S.V., R.O., N.P.H., Gi.R., E.B., M.L.J., G.Pa., K.G.S., K.M.F., G.F., A.L., A.Ca., E.S., S.A., M.A.T., T.Y., A.C., C.S., L.R., J.M.M.L., Gi.R., S.C., E.B., M.C.S., F.T., A.P.P., P.C., R.P., J.F.G.B., G.Pa., P.M.L.A., B.Bu., Ch.M., C.C., A.Ca., E.S., S.A., V.C.F., A.M., C.M., M.R., L.S., M.A.T., V.P.R, F.M.G., J.A.C.S., S.J.B., T.N.M., M.O.R.A., C.G.S., A.G.M., N.C. and L.Sa. were involved in sample collection. S.S., I.M., A.M.M., P.F., I.D., A.V., M.T., D.A., G.P., S.O., K.M., S.V., R.O., N.P.H., M.L.J., K.M.F., I.V.D., G.F., A.L., V.L., J.R., L.K., L.P., T.Y., S.S., A.C., C.S., L.R., J.M.M.L., B.B., Gi.R., S.C., E.B., M.C.S., F.T., A.P.P., P.C., R.P., J.F.G.B., G.Pa., P.M.L.A., B.Bu., Ch.M., C.C., A.Ca., E.S., S.A., V.C.F., A.M., C.M., M.R., L.S., M.A.T. and L.Sa. curated bioarchaeological data. F.E.Y., H.M., G.S., L.V.I., J.C., J.S., M.E.A., F.V.S., J.T.S. and C.G. were involved in genetic data generation. F.E.Y., T.P., H.M., M.S., T.V., I.A., T.S.K., A.R., G.R. and R.N. were involved in genetic data analysis. I.M., A.B.F., K.G.S., K.M.F., S.S. were involved in other data generation and analysis (Calibrating 14C-dating, diet, Strontium etc.). F.E.Y., G.K., A.W., R.T., K.K. and E.W. drafted the main text. F.E.Y., G.K., S.S., A.B.F., A.W., R.T., I.M., S.V., R.O. and N.P.H. drafted supplementary notes and materials. F.E.Y., G.K., S.S., A.B.F., A.W., R.T., I.M., I.A., H.M., G.S., A.V., J.M.O., L.V.I., S.V., R.O., N.P.H., Gi.R., E.B., M.L.J., G.Pa., K.G.S., K.M.F., R.N., L.O., A.Ca., M.E.A., M.S., J.V.M., K.K. and E.W. involved in reviewing and editing drafts. All co-authors read, commented on and agreed on the submitted manuscript.

## Ethics declarations

The authors declare no competing interests

## Additional Information

Supplementary Information is available for this paper.

## Extended Data Figures

**Extended Data Fig.1.**
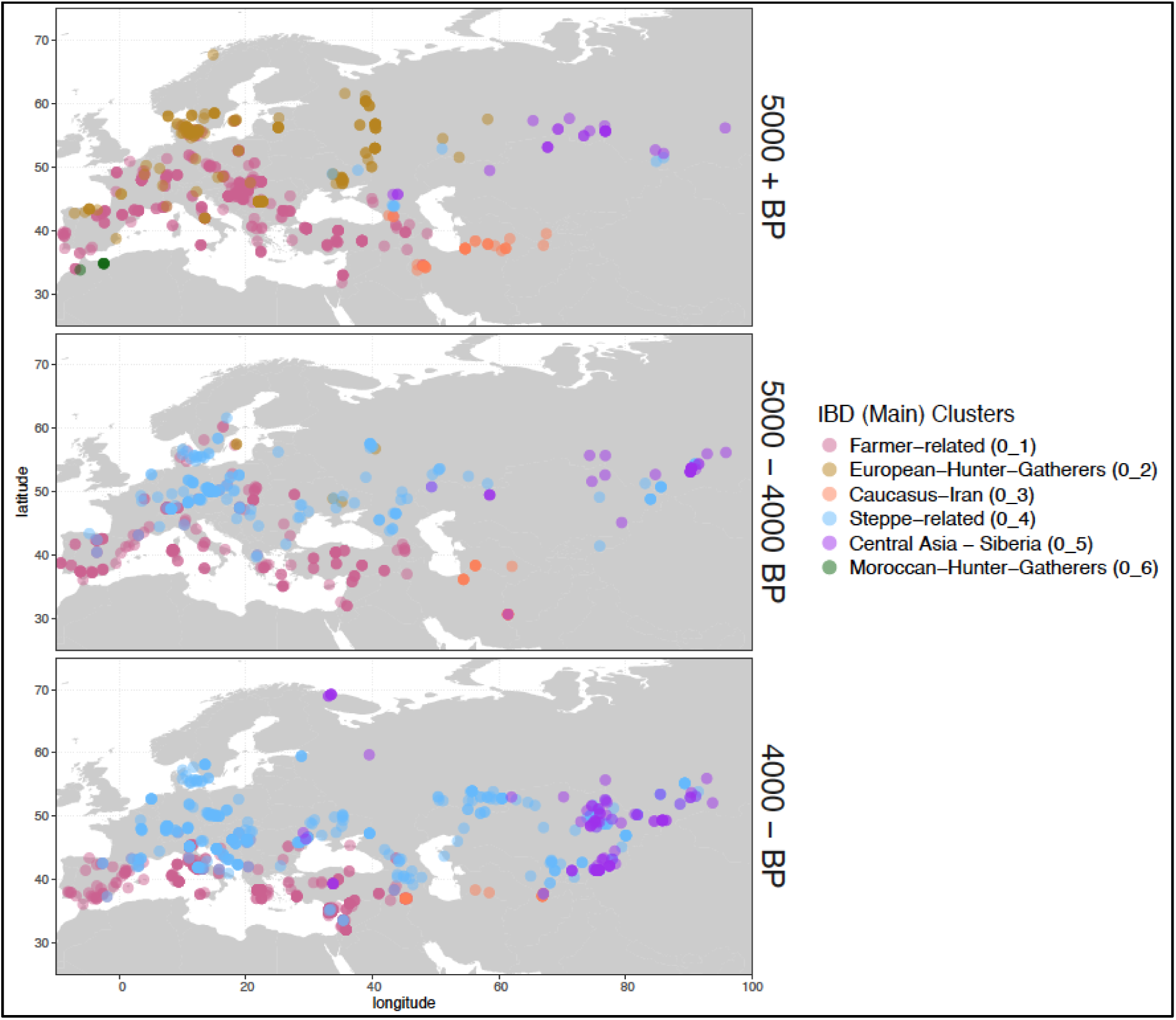
Geographical distribution of the main IBD clusters, split into time ranges, pre 5,000 BP, 5,000–4,000 BP, and post 4,000 BP.

**Extended Data Fig. 2.**
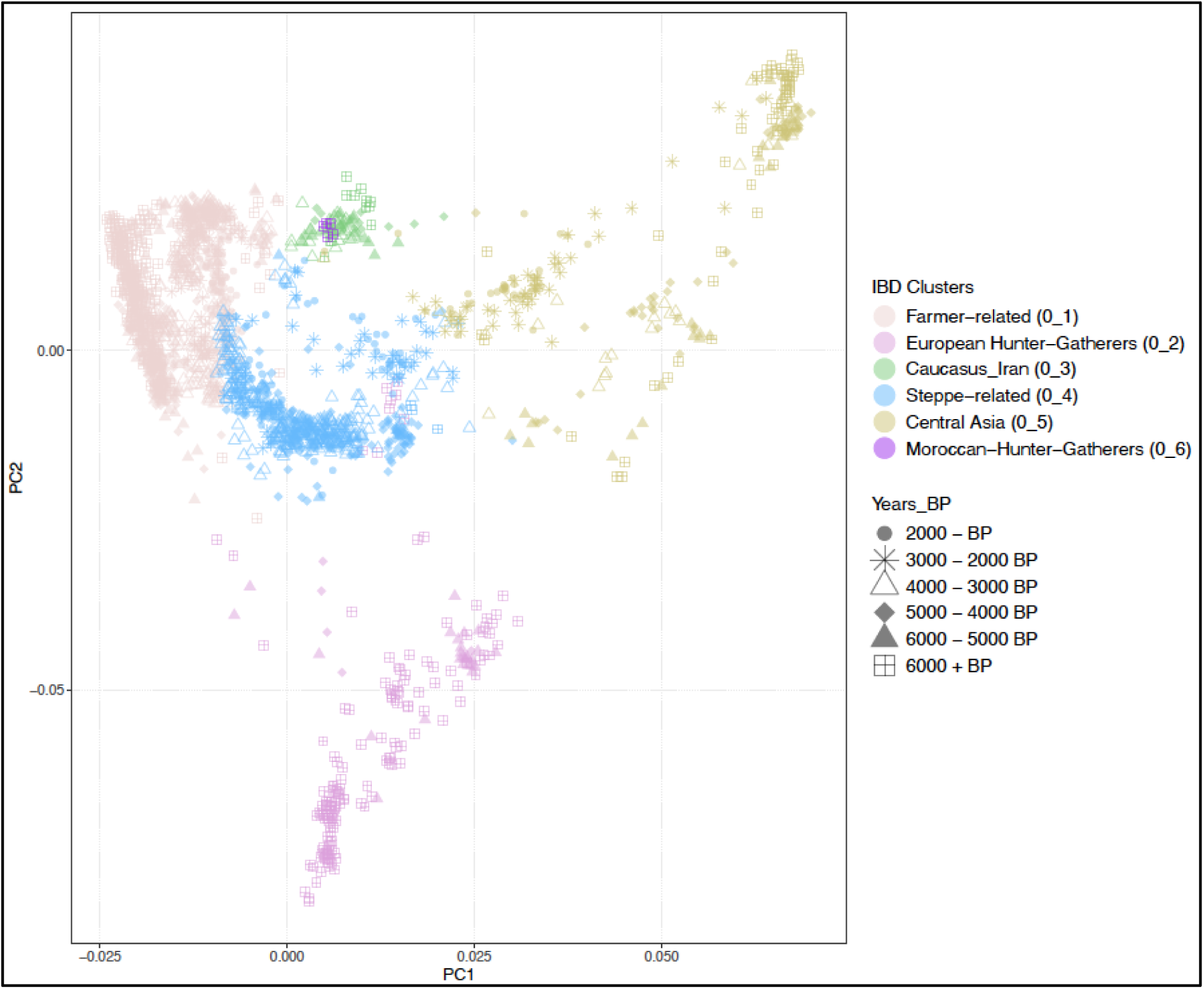
The PCA plot demonstrates the distribution of main IBD clusters on 2,228 ancient individuals.

**Extended Data Fig. 3.**
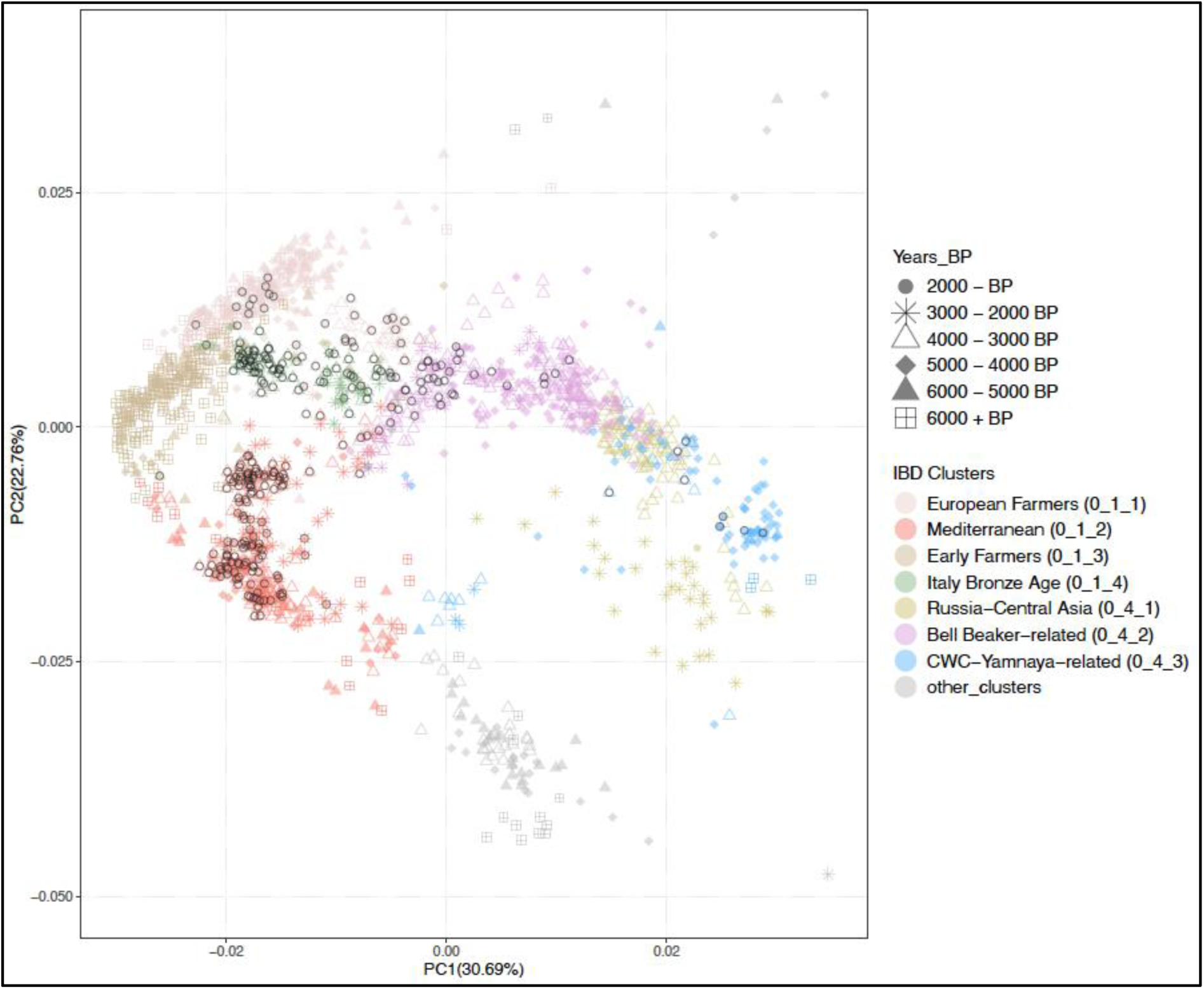
The PCA plot demonstrates the distribution of subclusters of “Farmer-related (0_1)” and “Steppe-related (0_4)”. New genomes presented in this study were marked with black circle legends.

**Extended Data Fig. 4.**
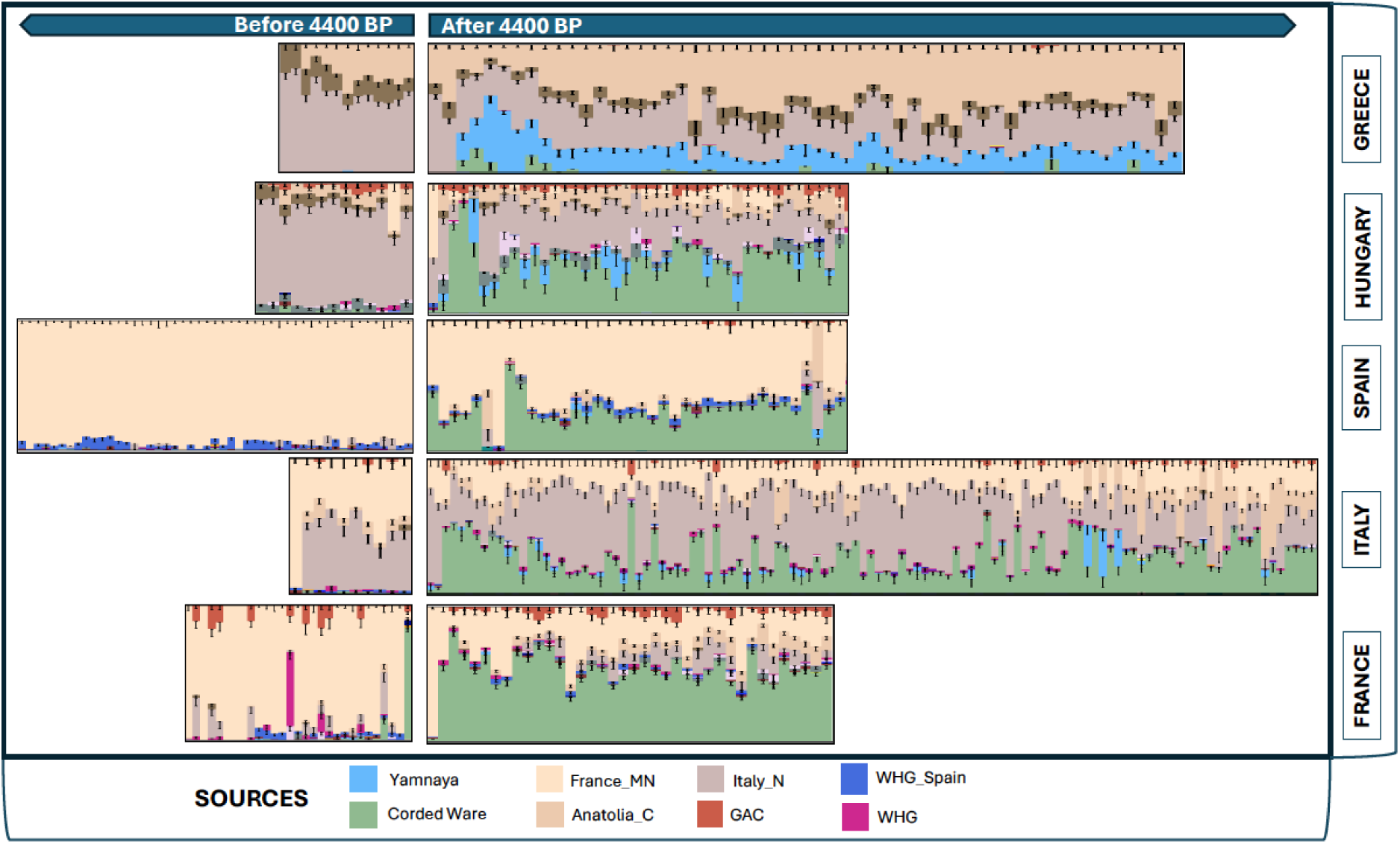
Ancestry bar plots generated for each individual using source population proportions of IBD admixture modelling sorted by time BP and divided into two time series, before and after 4,400 BP, illustrating a Southern and Central Eastern Europe split (Italy, France, Spain and Hungary vs Greece).

**Extended Data Fig. 5.**
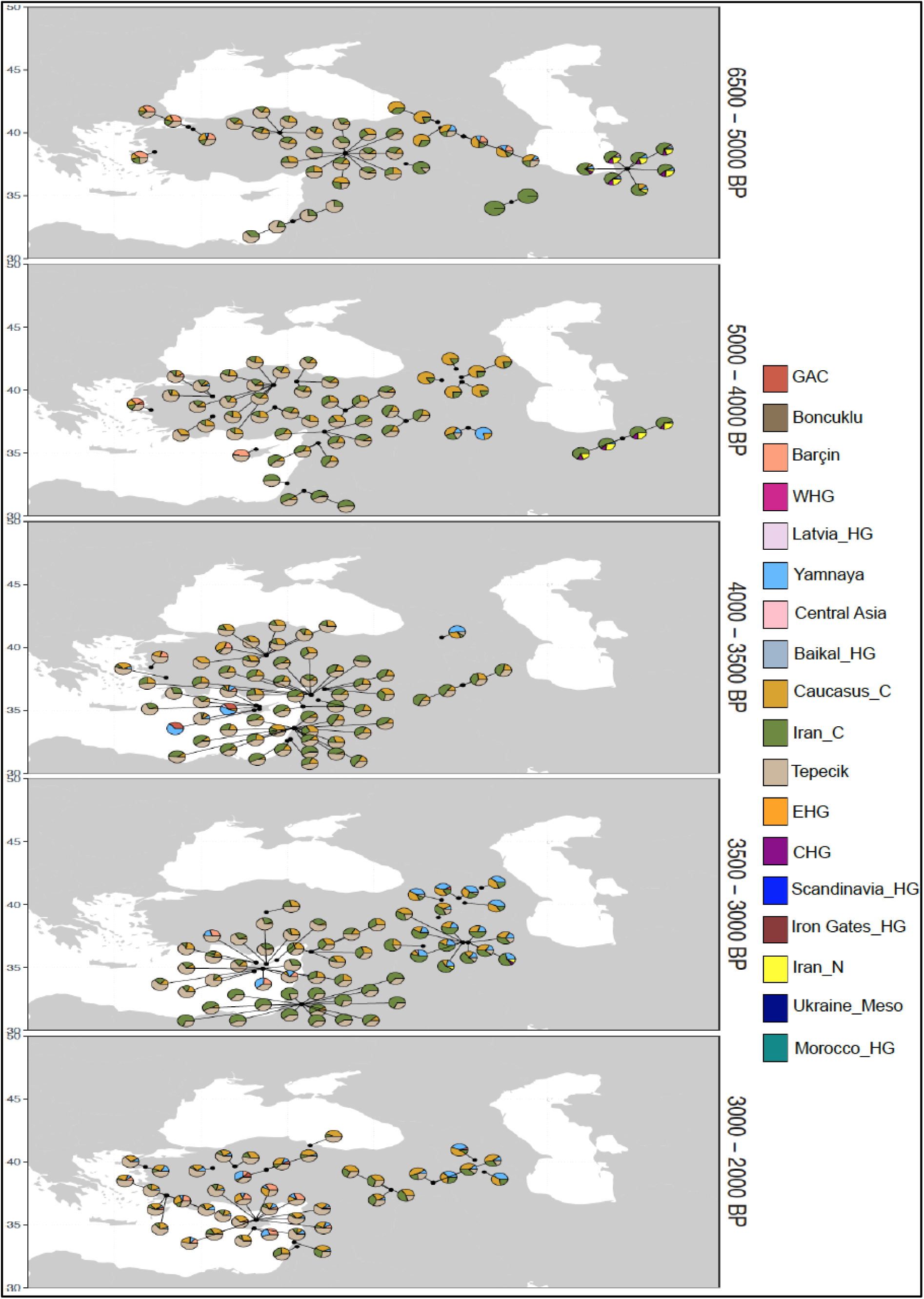
Pie charts generated by using the proportions of the applied IBD admixture model for each individual from Anatolia, Cyprus, Iran, Caucasus and Levant, divided into five time periods to avoid overlapping.

**Extended Data Fig.6.**
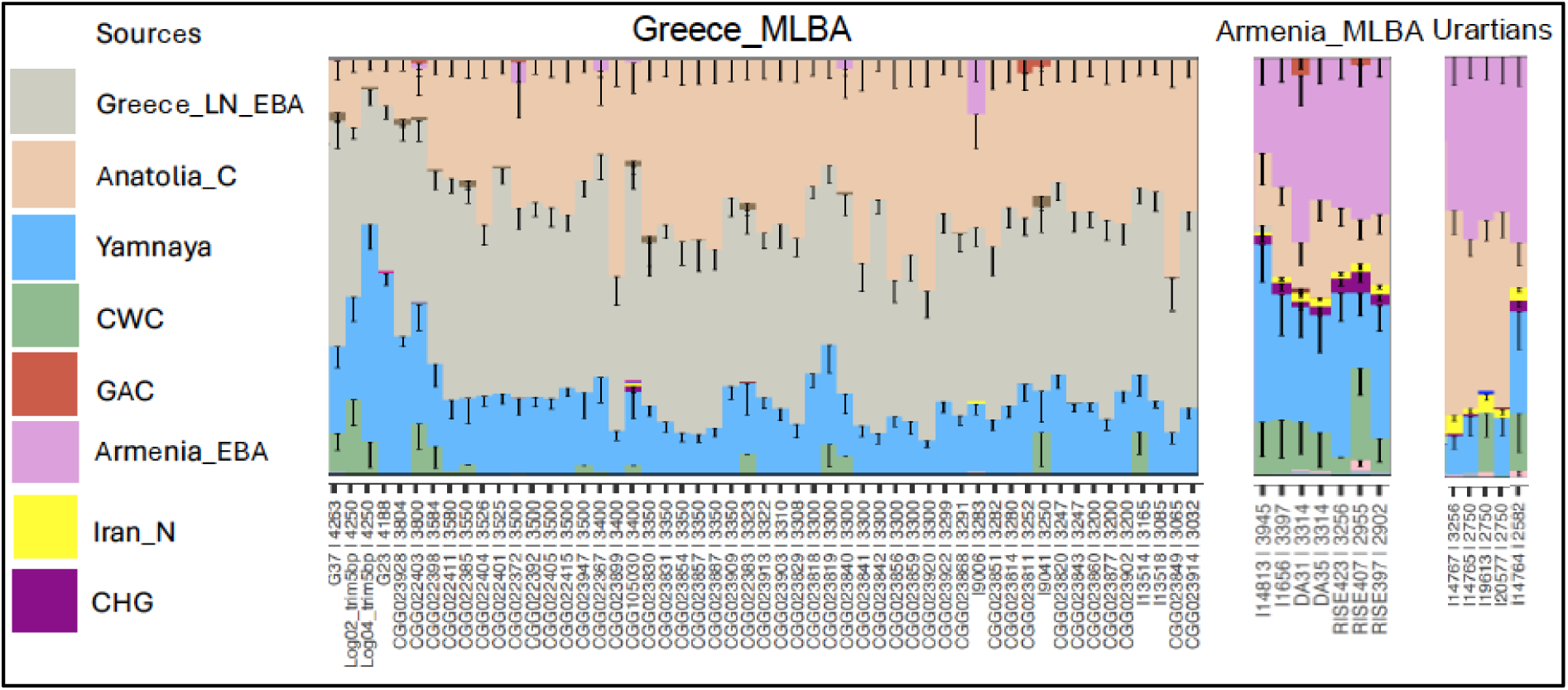
Bar plots generated using source proportions of the IBD admixture modelling that shows the similarity of steppe ancestry in Greece and Armenia Middle Late Bronze Age and Urartians modelled with Yamnaya and local populations.

