## Supplementary material for "Ancient genomics support deep divergence between Eastern and Western Mediterranean Indo-European languages": Genetics and Strontium Supplementary

**Genetics and Strontium Isotope Analysis Supplementary Material**

***S1. Data generation	1***

***S2. Bioinformatics pipeline	1***

***S3. Dataset preparation	1***

**S4. Principal component analysis (PCA)	2**

***S5. IBD (Identical by Descent) clustering	5***

***S6. IBD mixture modelling	15***

**S6.1 Defining farmer sources	16**

**S6.2 Southern Europe	23**

**S6.3 Central Eastern Europe	35**

**S6.4 Eastern Mediterranean	38**

***S7. Relatedness analysis	54***

***S8. Genetic sex determination	54***

***S9. Uniparental markers	55***

**S9.1 Y-chromosome analysis	55**

**S9.2 Mitochondrial DNA analyses	57**

***S10. Individual mobility assessment using strontium (Sr) isotopes	63***

**References	75**

### S1. Data generation

Charleen Gaunitz, Fulya Eylem Yediay, Lasse Vinner

Ancient samples in this study were generated by applying both manual and automated wet-lab procedures as detailed in Allentoft 2024 and McColl 2024^1,2^. All pre-PCR procedures were carried out in the dedicated ancient DNA clean lab facilities at the Lundbeck Foundation GeoGenetics Centre, University of Copenhagen, according to strict guidelines, and post-amplification procedures were carried out in separated post-PCR laboratories. The sequencing was done on the Illumina Hiseq 4000, as described in Allentoft et al., and the NovaSeq6000 platform at the GeoGenetics Sequencing Core, University of Copenhagen^1^.

### S2. Bioinformatics pipeline

Isin Altinkaya, Abigail Daisy Ramsøe, Fulya Eylem Yediay, Gabriel Renaud, Thorfin Sand Korneliussen

All libraries were generated by following manual and automated wet-lab pipelines as detailed in McColl et al. 2024^2^. We screened 1,929 libraries, which were produced from 380 ancient individuals. For each sample, we produced four libraries (two USER and two non-USER). Due to contamination, sample swapping and low library complexity, we excluded non-authentic libraries, so not all samples ended up with four libraries in the end (Supplementary Table S1). After assessing library quality and complexity, we merged the selected libraries using two approaches: 1) If the coverage was less than 1x, we merged both user and non-user treated libraries and trimmed the damages from 5bp from 5’ ends. 2) If the coverage was above 1x, we only merged user-treated libraries (Supplementary Table S1). Non-user libraries were used to evaluate the damage patterns^3^. To assess the mitochondrial contamination level, we ran contamMix and schmutzi, and for the X chromosome contamination level, we ran ANGSD^4–6^ (Supplementary Table S1).

### S3. Dataset preparation

Fulya Eylem Yediay

In total we screened 380 ancient individuals and retrieved DNA coverage of between 0.004x and 15.5x , 144 of which were radiocarbon-dated. However, we continued downstream analyses with 314 ancient individuals which provided an efficient amount of DNA (>0.1x) from Southern Europe and the Eastern Mediterranean dating from between 5,200 BP and 1,100 BP (Supplementary Table S1; Fig. S3.1). To generate the dataset, we first imputed ancient samples merged from 2,089 published capture and shotgun data together with 314 newly sequenced whole genome shotgun samples[^1,7–89^](https://paperpile.com/c/8RV5oM/ZBJ7t+JQFFu+Mfzu+sVibA+T0YR+lZBkR+WQhF+5yBR+e7NSh+QNIKg+4PvbP+KAWU+4qstp+2cyo+hkHT+QpAu+6M5l+CBIF+8cTF+Biks+LmXR+P8bm+DdxuO+gAUOG+3oRU+1vTL+CPRo+7qrt+74fT+qoBR+iaQIC+OhnV+fqYqX+P8ff7+nXLCp+NXnx+JV3A+TUbtX+Dnjdp+ZthJt+Bx78c+Fwh5+psBV+0Iqpk+6QN92+TPhf+OxWB+yXG5+0pqpx+7ePx+1RMa+qvOQ+XSbD+ItXS+KEh5+gIjBt+Rurc+axcw+s2cE+PJv4+EDPj+5ol5+3pWy+c6ME+J3TF+dbst+ieur+okuL+spNO+dlQF+vp6J+Icjy+21PN+rfxc+Dsl8+Kl6v+d5pB+gGJC+MLVn+hPmG+I21F+cjRU+f0Me+Jtnu) (Supplementary Table S2), filtered the merged VCF files INFO>0.5, MAF (minor allele frequency) 0.01 and restricted to 1,240K capture SNP sites, and finally applied 1000G strict mappability mask. To run IBDseq, we included the samples with >0.1x coverage for shotgun, >1x coverage for capture, and >0.90 average genotype probability across 643,430 SNP (single nucleotide polymorphism) sites^90^ (Supplementary Table S2). We ran IBDseq following the implementation in McColl et al. 2024 on 2,403 ancient samples within a broad geographical area and period (300 BP–45,000 BP)^1,2^. After removing close relatives, we ran IBD clustering with a total of 2,228 ancient samples, of which 274 were newly sequenced Bronze Age Mediterranean individuals (Fig. S3.2, Supplementary Table S2; S3).

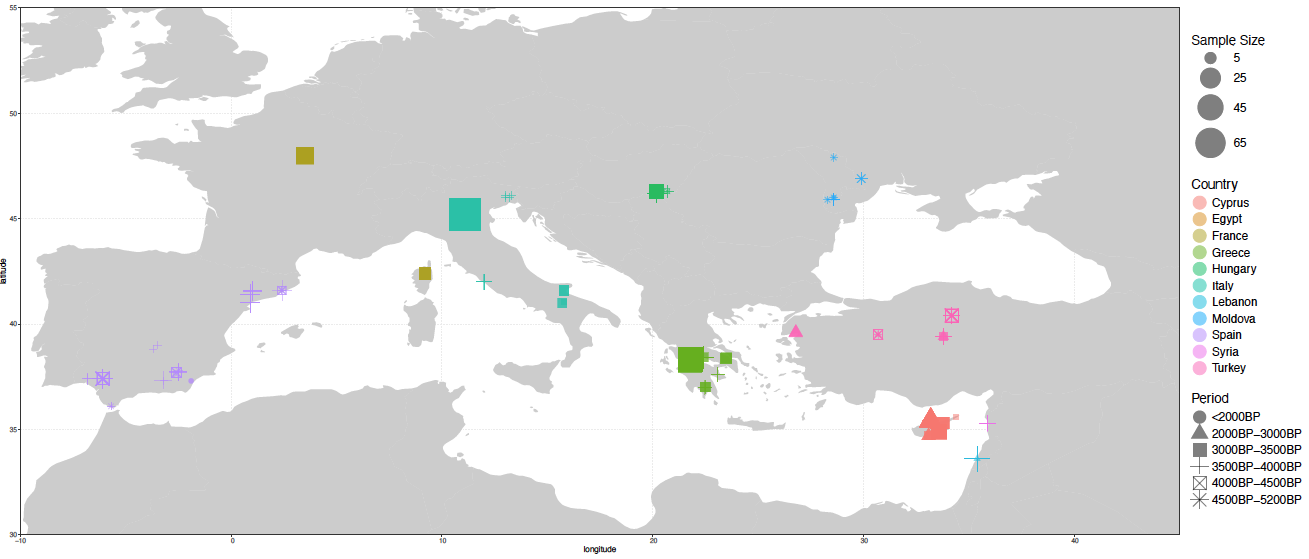

Fig. S3.1. Map of all samples that have been processed for the first time in this study.

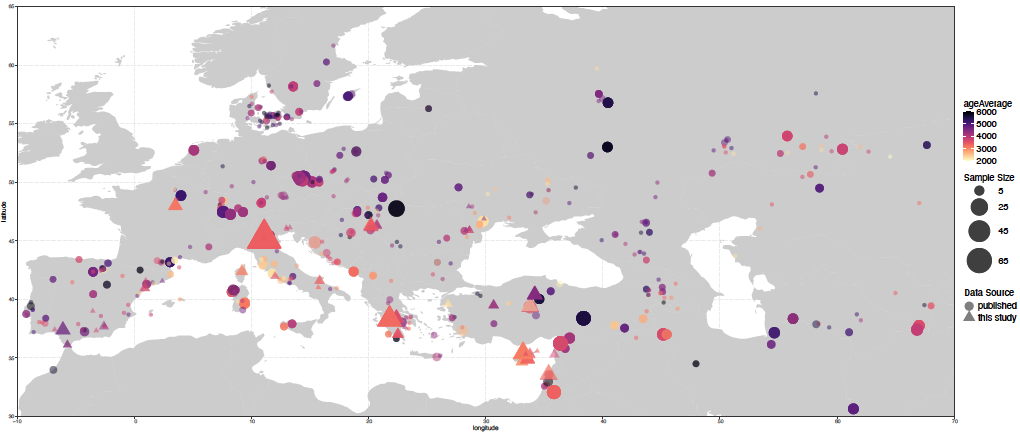

Fig. S3.2. The map represents the geographical distribution of individuals from each archaeological site limited to an average age of between 6,000 BP and 2,000 BP.

### S4. Principal component analysis (PCA)

Fulya Eylem Yediay

To get an overview of the basic structure of our data, we carried out a principal component analysis (PCA) on an imputed dataset, comprising 1,837 ancient individuals from both shotgun and capture (1240K) data using plink (v1.90b6.21). We first restricted the SNPs to capture sites, and applied MAF (0.05) filtering, resulting in a dataset with 580,130 SNPs. To visualize our data clearly, we plotted the individuals by splitting them into two periods, the first between 5,200 BP and 3,500 BP, and the second between 3,500 BP and 2,000 BP, then coloured them by countries of origin. The European Bronze Age individuals are referred to individuals who carry steppe-related ancestry and are placed between Farmer and Steppe clusters (Fig. S4.1). We observed an emerging Balkan cline between the European Bronze Age and the Aegean cluster towards the East Mediterranean cluster (Fig. S4.1). The individuals from Hungary in our dataset scattered on this cline from the edge of Europe to the East Mediterranean, together with one individual from Greece (CGG_2_022403) and three Italy Middle Bronze Age individuals from Northeastern Italy (CGG_2_022923, CGG_2_022253, CGG_2_022653). The Chalcolithic individuals from Spain clustered with other Iberian farmers, except for one individual (CGG_2_023951) who falls on the Sardinian farmers cluster (orange colour) together with one sample from Italy (CGG_022655, Lucone). However, Spain Bronze Age individuals are shifting to the European edge of the Balkan cline from local Spain farmers. We also found two individuals from Central Italy (CGG_2_101266, CGG_2_101264, Coppa Navigata) within this cluster. In the Aegean cluster, Greece Middle Bronze Age individuals fall between Anatolia and the Balkan cline, except one Chalcolithic individual from Kalyvia who clusters with Greece farmers. The individuals from Cyprus fall on the Anatolian Bronze Age cluster, except one individual (CGG_2_022488) shifting towards Greece Bronze Age. In the Steppe cluster, we found Early Bronze Age individuals from Moldova, while the Middle Bronze Age individuals clustered with Corded Ware (CWC) individuals. We also found one outlier (CGG_2_103667) slightly shifting towards the Caucasus from the CWC individuals.

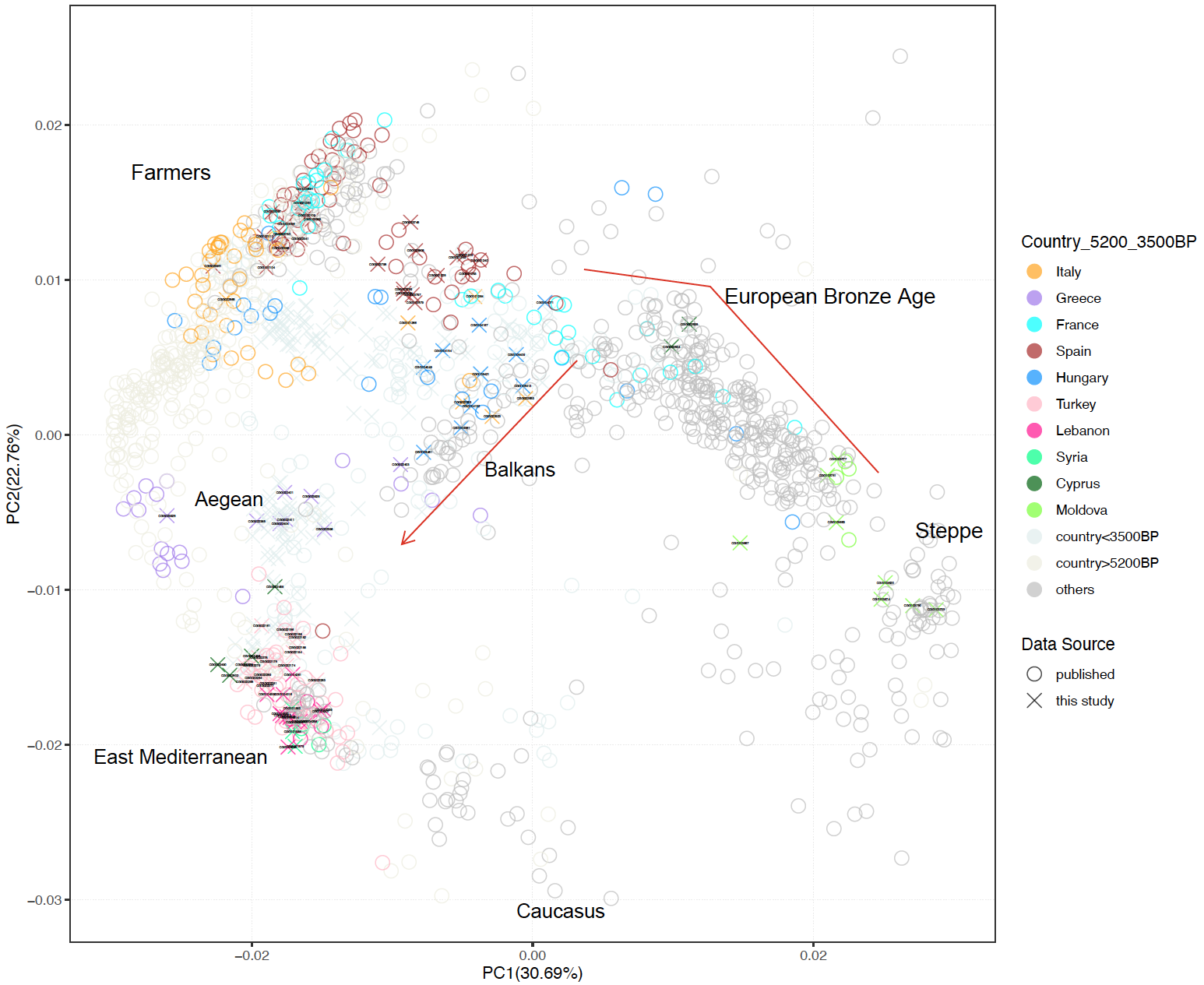

Fig. S4.1. PCA plot generated using a subset of data. Only individuals from the listed countries and dated between 5,200 BP and 3,500 BP were coloured, labelled by sample ID.

After 3,500 BP, we detected that most Italy Bronze Age individuals are located between Sardinian farmers and the Balkan cline, with some of them shifting to the European edge of the Balkan cline, and a few shifting towards the Aegean edge (CGG_2_100646, CGG_2_022640, CGG_2_022641) (Fig. S4.2). The Italy Bronze Age individuals in the European edge of the Balkan cline cluster with Late Bronze Age individuals from France (Migennes) together with some of the Hungarian Bronze Age individuals. Among the individuals from Italy, we also detected one outlier who falls in the European Bronze Age cluster (CGG_2_022591) further from the Balkan cline together with one individual from Hungary (CGG_2_103857). On the contrary, Late Bronze Age individuals from Corsica cluster with Italy Late Bronze Age individuals from Olmo who are placed between Sardinia farmers and the Balkan cline.

In Cyprus, the shift from the East Mediterranean to the Aegean pool became more extensive compared to earlier periods (before 3,500 BP), and moreover, some of the individuals clustered with Greece Bronze Age individuals while some fell between Anatolia and Greece Bronze Age. Among those individuals, we observed two Iron Age individuals from Western Anatolia also slightly shifting to Greece Bronze Age (CGG_2_023618, CGG_2_022162) (Fig. 4.2).

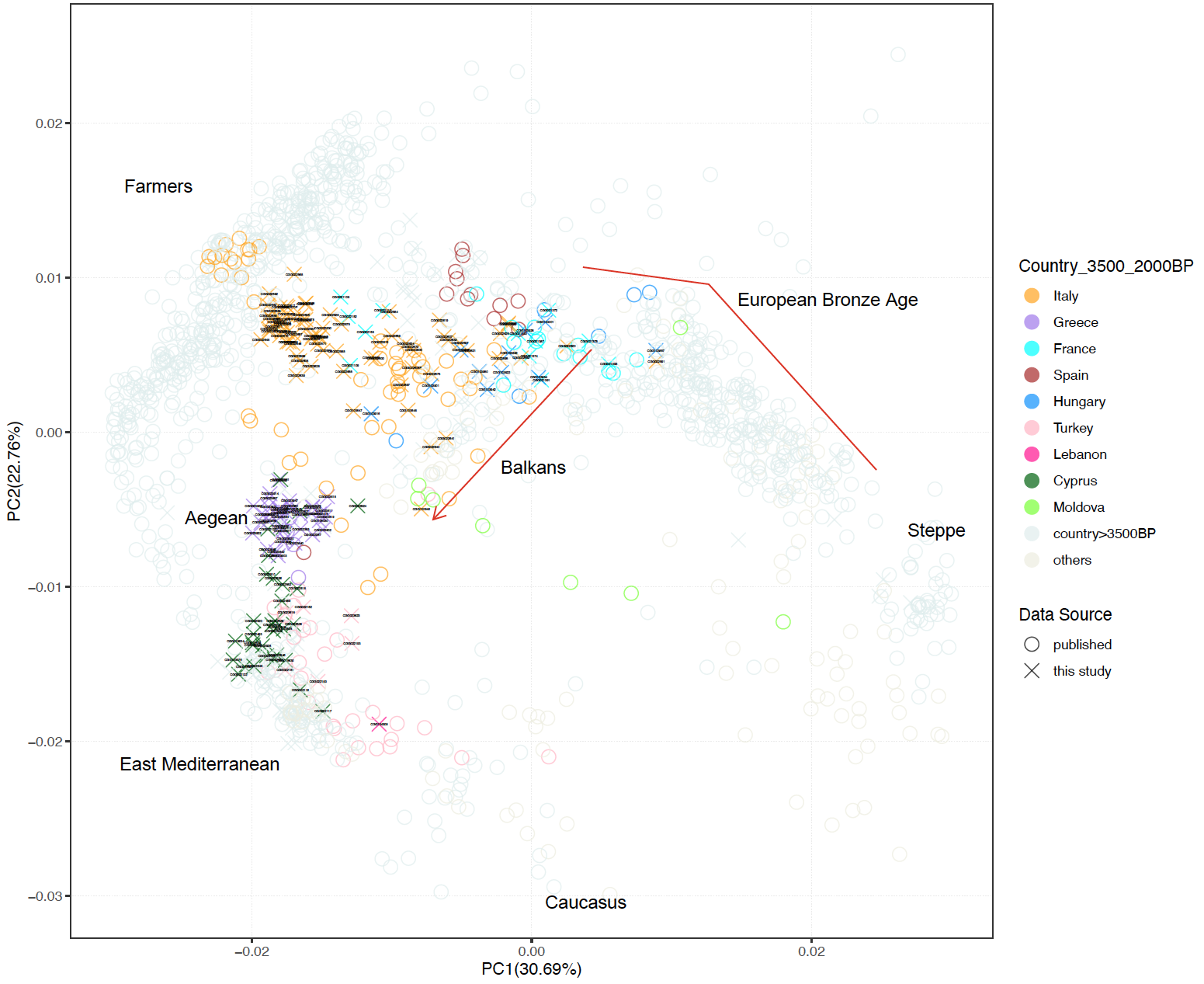

Fig. S4.2. PCA plot generated using a subset of data. Only individuals from the listed countries and dated between 3,500 BP and 2,000 BP were coloured, labelled by sample ID.

### S5. IBD (Identical by Descent) clustering

Fulya Eylem Yediay, Hugh McColl, Martin Sikora

Prior to running Leiden network-based hierarchical clustering, we applied filters by removing the IBD segments of less than 1 cM, LOD score of less than 3, and hotspot regions. We also removed one pair of the first and second-degree relatives to minimize small clusters formed only with close relatives. We then ran the clustering with 2,228 ancient individuals by setting a minimum total shared IBD of 5 cM and a permutation of 200.

Among the IBD clusters, we detected six main clusters that represent a geographical and temporal distinction and contain many subclusters (Supplementary Table S3; Table S5.1; Fig. S5.1). We visualized the PCA plot of 2,228 ancient individuals by colouring the main clusters (Fig. 5.2). Here we scrutinize the two main clusters (0_1 and 0_4) and their subclusters, within which 314 individuals of this study fell (Table S5.2; Fig. S5.3).

| **Deep clusters** | **Period** | **Ancestry** |
| --- | --- | --- |
| 0_1 | 300 BP–26,000 BP | Farmer-related |
| 0_2 | 300 BP–38,000 BP | European Hunter-Gatherers |
| 0_3 | 3,000 BP–14,000 BP | Caucasus – Iran |
| 0_4 | 400 BP–6,900 BP | Steppe-related |
| 0_5 | 500 BP–45,000 BP | Central Asia – Siberia |
| 0_6 | 7,000 BP–15,000 BP | Morocco-Hunter-Gatherers |

Table S5.1. Ancestry groups of deep clusters and time ranges.

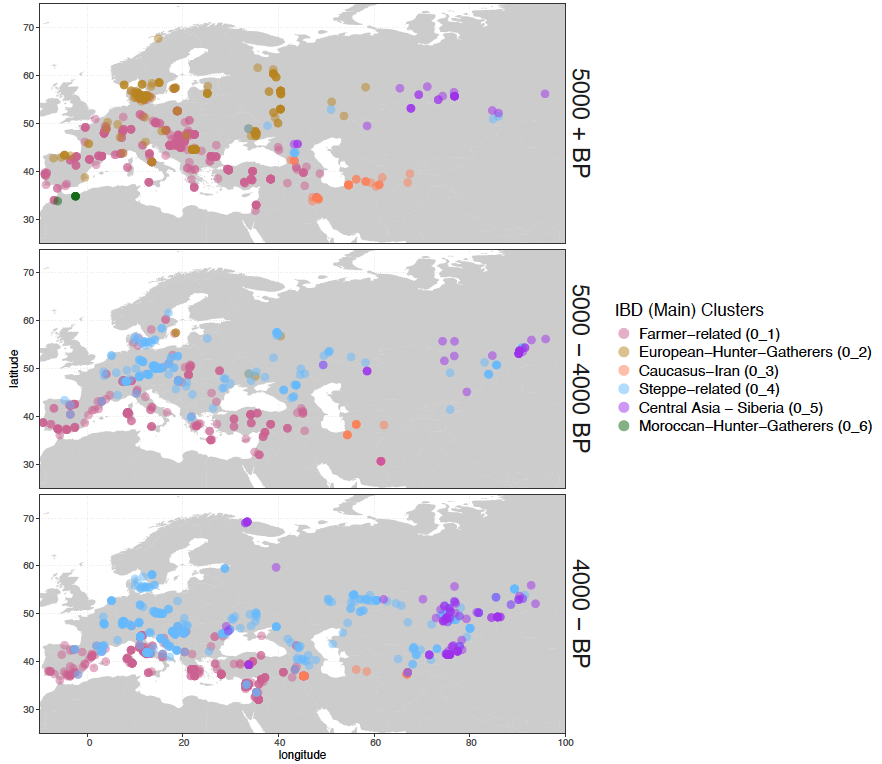

Fig. S5.1. Geographical distribution of the main IBD clusters, split into time ranges, pre 5,000 BP, 5,000–4,000 BP, and post 4,000 BP.

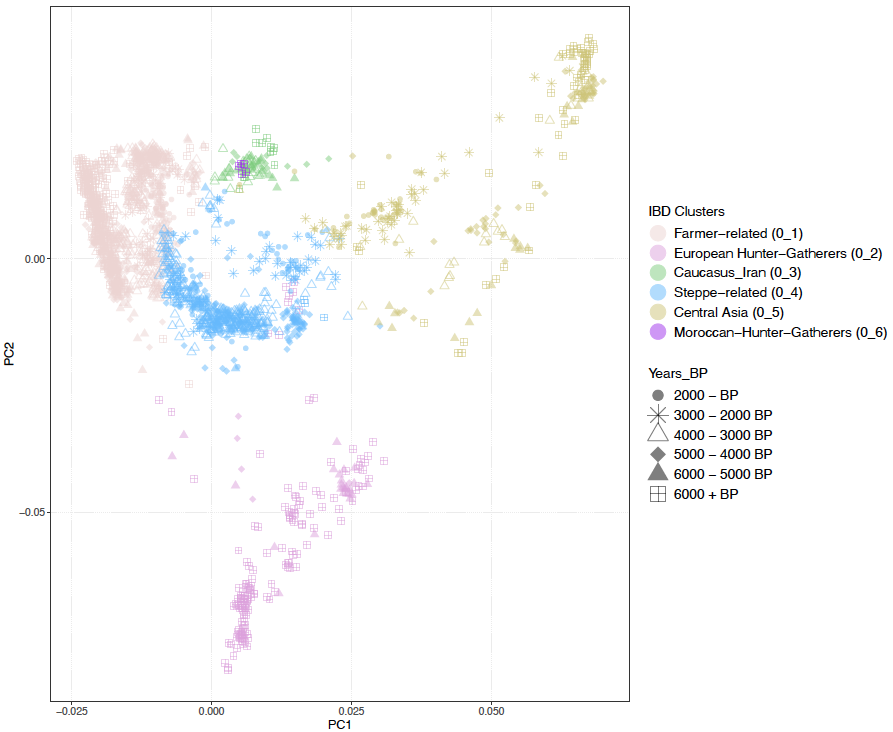

Fig. 5.2. PCA plot generated with 2,228 individuals that were included for IBD clustering (after removing close relatives), coloured by the main clusters.

| **The Subclusters** | **Period** | **Ancestry** |
| --- | --- | --- |
| 0_1_1 | 500 BP–7,400 BP | European Farmers |
| 0_1_2 | 300 BP–26,000 BP | Mediterranean |
| 0_1_3 | 3,100 BP–16,000 BP | Early Farmers |
| 0_1_4 | 4,200 BP–3,000 BP | Italy Bronze Age |
| 0_4_1 | 1,500 BP–5,600 BP | Russia-Central Asia |
| 0_4_2 | 400 BP–4,700 BP | Bell Beaker-related |
| 0_4_3 | 800 BP–6,900 BP | CWC-Yamnaya-related |

Table S5.2. Ancestry groups of the subclusters that the individuals of this study fall within and time periods.

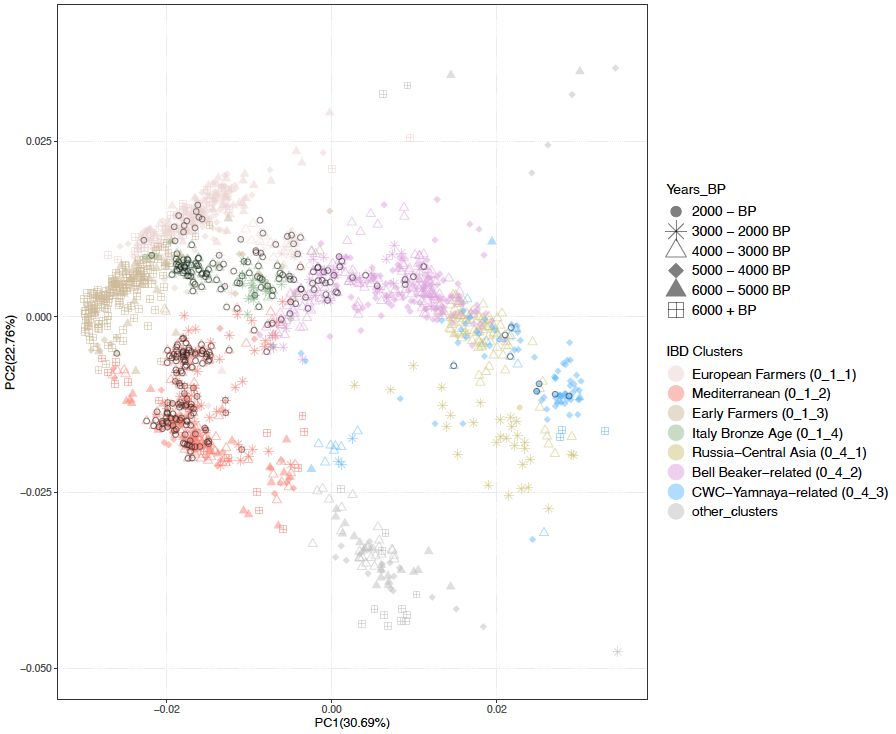

Fig. 5.3. PCA plot generated by a subset of individuals (n=1,837), coloured by subclusters. Black circles indicates individuals presented in this study.

The “Farmer-related (0_1)” cluster, mainly consisting of farmer-related individuals and later period individuals who carry more farmer ancestry. This cluster contains four subclusters: 0_1_1, which represents European farmers; 0_1_2, which covers individuals from a broad geographical region from Western Asia to Southern Europe; 0_1_3, which is the Early Farmer cluster; and 0_1_4, which stands out as containing mainly Italy Bronze Age individuals (Fig. S5.4). We identified a phenomenon in cluster 0_1_1 whereby, while the European farmers in this cluster had a wider geographical distribution, after ~4,000 BP only individuals from Southwestern Europe remained in this subcluster. This shows that the expansion of steppe ancestry is later or has lower impact in this area (Fig. S5.4). In the five subclusters of 0_1_1, we find our newly sequenced Spain Chalcolithic/Bronze Age individuals within subcluster 0_1_1_2, and our newly sequenced Late Bronze Age individuals from Corsica and two individuals from Central Italy (Coppa Navigata and Pian Sultano), along with previously published individuals from this area, within subcluster 0_1_1_5.

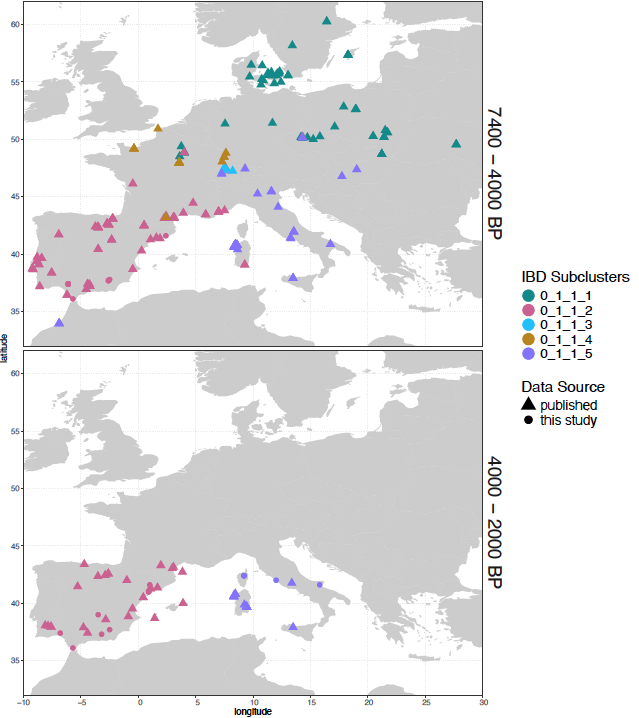

Fig. S5.4. Geographical distribution of the individuals within subclusters of 0_1_1, before/after 4,000 BP.

The subcluster 0_1_2 represents the East Mediterranean populations, containing individuals from Southern Europe and Western Asia. It is split into six subclusters. We show the distribution of individuals in these subclusters in three different time periods in Fig. S5.5. The earliest individuals found in this subcluster are from Anatolia, the Levant, Azerbaijan, and Iran (Tepecik, Buyukkaya, Israel_PPNB, Haji Firuz) before ~7,000 BP (Fig. S5.5). The individuals dating to between 7,000 BP and 4,000 BP are from Greece, Hungary, and Western Asia, and within the subclusters they show a geographical differentiation between Anatolia, Europe, and the area from the Caucasus to the Levant. When we look at the individuals in the later periods (2,000 BP to 4,000 BP), the subcluster 0_1_2_3 shows a wider distribution including individuals from Italy, the Balkans, Cyprus, and Anatolia, including our new data from Cyprus, Italy, and Western Anatolia (Fig. S5.5).

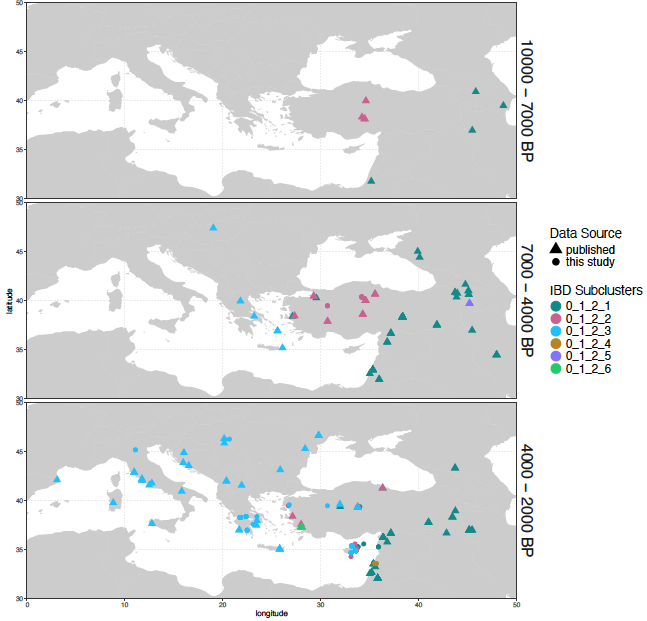

Fig. S5.5. Geographical distribution of the individuals within the subclusters of 0_1_2.

Another Farmer-related subcluster, 0_1_3, covers early farmer individuals mainly from Central Eastern Europe and Anatolia, along with exclusively LBK-related farmers from Western Europe before ~6,400 BP (Fig. S5.6). After this time, only Central Eastern and Southeastern Europe farmers are found within subcluster 0_1_3 (Fig. S5.6). In this subcluster, our newly sequenced Early Bronze Age individual from Greece (CGG_2_023925, Kalyvia) clusters with the Central Eastern European farmers and one individual from Cyprus (CGG_2_02253, Karavas) clusters with Anatolian farmers, as do Greece Early Bronze Age individuals from the Peloponnese. Given that Neolithic and Early Bronze Age individuals from Greece fell within various Farmer-related subclusters, this was a dynamic period connecting the Eastern Mediterranean and the Balkans.

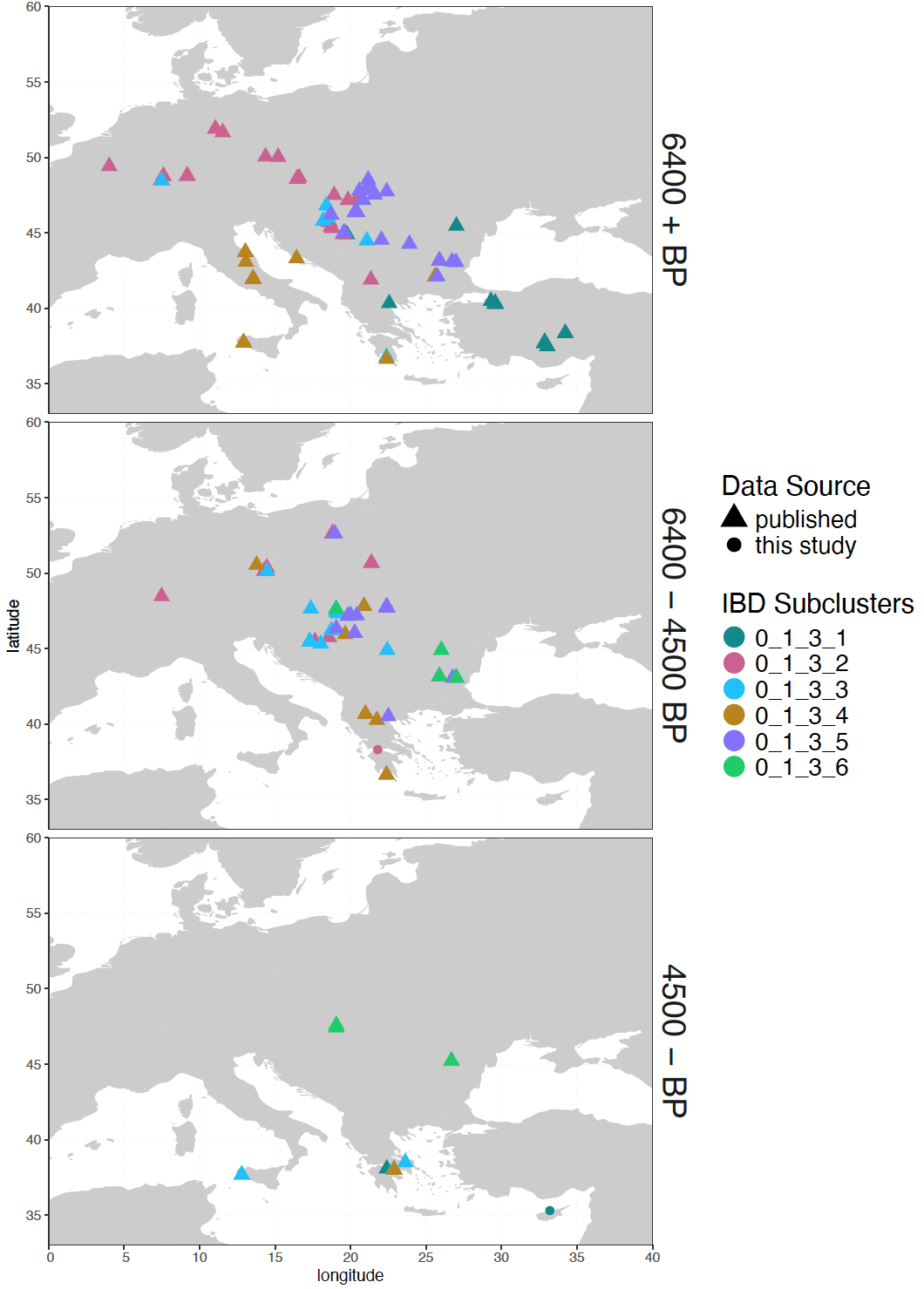
Fig. S5.6. Geographical distribution of the individuals within subclusters of 0_1_3, divided into three time ranges.

Due to dense sampling of Bronze Age individuals from Italy, they split into the subcluster 0_1_4 along with Corsica Late Bronze Age individuals, as well as a few individuals from Southern Germany, Hungary, and Switzerland dated to between 1,400 BP and 4,300 BP, showing a possible Balkan connection during the Bronze Age (Fig. S5.7).

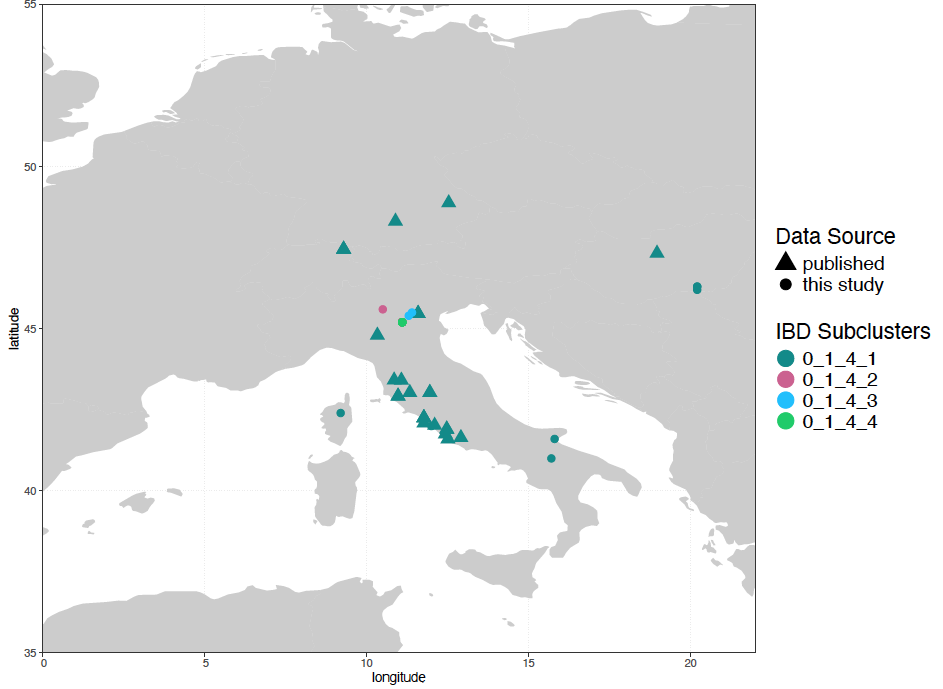

Fig. S5.7. Geographical distribution of the individuals within subclusters 0_1_4, between 1,400 BP and 4,350 BP.

While most individuals from Italy fell within the Farmer-related cluster, some fell within the “Steppe-related (0_4)” cluster. This main Steppe cluster is divided into three subclusters (0_4_1, 0_4_2, and 0_4_3) containing individuals from Siberia to Western Europe dated to between 500 BP and 6,900 BP, and it shows a clear geographical distinction especially after 4,000 BP (Fig. S5.8). The individuals from Italy, Hungary, and Moldova reported in this study fell within subclusters 0_4_3 and 0_4_2, except one newly sequenced Moldovan individual (CGG_2_103781, ~3,700 BP) that falls within the subcluster 0_4_1 together with one published Ukraine Chalcolithic individual (I6561 from Mathieson et al., 2018)^46^ and some individuals from Russia and Central Asia Bronze and Iron Age. Interestingly, Fatyanovo related individuals are also found within two other subclusters (0_4_2 and 0_4_3). The subcluster 0_4_1 spread eastward into Central Asia and Siberia after 4,000 BP (Fig. S5.8).

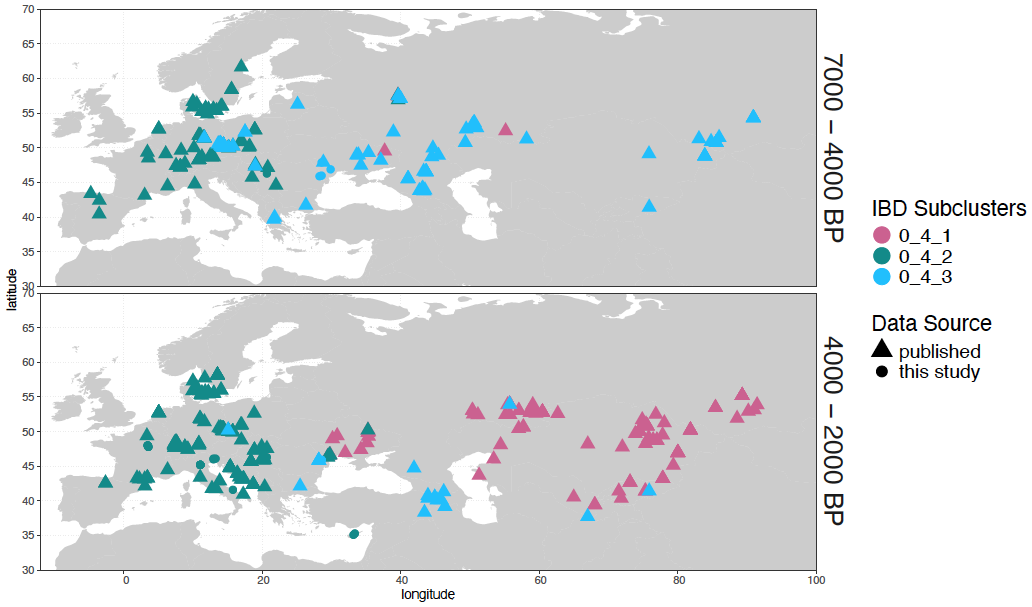

Fig. S5.8. Geographical distribution of the individuals within subclusters of the main Steppe cluster 0_4, divided into two time periods.

Even though all Yamnaya and early Corded Ware-related individuals fall within the subcluster 0_4_3, we see a distinction of these populations into three subclusters (Fig. S5.9). Moreover, we observe that Bronze Age individuals from Middle Bronze Age Greece, individuals from Hungary associated with the Bell Beaker complex, and individuals from Early Bronze Age Bulgaria also fell within this subcluster. The subcluster 0_4_3_1 contains individuals from Yamnaya (Ukraine, Samara, Kalmykia), Afanasievo, Catacomb, and Caucasus Yamnaya contexts. Here we report new data from the Moldova Bronze Age, associated with Late Yamnaya people and clustering with Yamnaya individuals along with the published Greece Middle Bronze Age sample Log04, showing the steppe connection^14^. The Early Corded Ware individuals, together with Bronze Age individuals from Hungary and Bulgaria and Moldova Middle Bronze Age individuals, fell into subcluster 0_4_3_2 (Fig. S5.9). Whereas the Early Bronze Age individuals from Moldova clustered with Yamnaya, two of the newly sequenced Moldovan individuals from *Balabanu* and *Traclia* also belong to subcluster 0_4_3_2. Another subcluster is 0_4_3_3, which contains Steppe Eneolithic individuals from Piedmont and individuals from Ukraine, Greece, and Bulgaria dated to before 4,000 BP. After this time, Armenia Middle/Late Bronze Age individuals and one Urartian individual are found within the same cluster^9,15,36,40,46^.

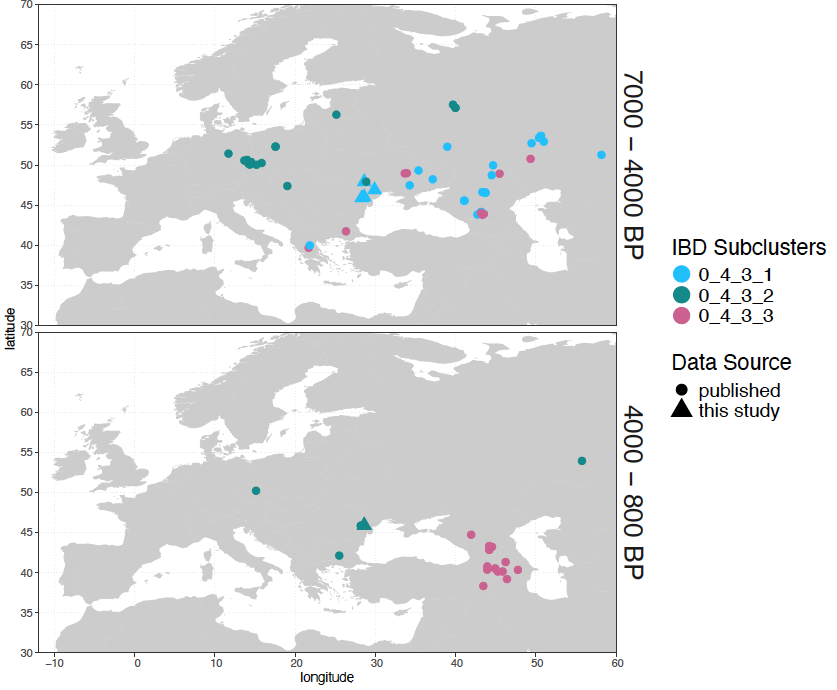

Fig. S5.9. Spatial and temporal distribution of the individuals within subclusters of 0_4_3.

The steppe-related European Bronze Age individuals are included in subcluster 0_4_2 regardless of their cultural context (CWC, BB, Únětice etc.), however, the majority of individuals are Bell Baker-related (Fig. S5.10). We found many subclusters within 0_4_2, but note that two subclusters (0_4_2_6 and 0_4_2_7) consist of only a few samples that could be due to third and further degree relatives clustering separately. To show the distribution of the subclusters under 0_4_2, we split individuals into two periods: before and after 3,000 BP (Fig. S5.10). This plot shows that two subclusters (0_4_2_2 and 0_4_2_3) are found to be widespread in Western and Southern Europe. Moreover, the Southern European subcluster 0_4_2_3 shows an expansion along the Adriatic coast and Central Eastern Europe. The Scandinavian Bronze Age individuals, mainly represented by subcluster 0_4_2_5, spread into the Balkans and Southern Europe after 3,000 BP, and a few Lebanon Medieval individuals are also found in this subcluster.

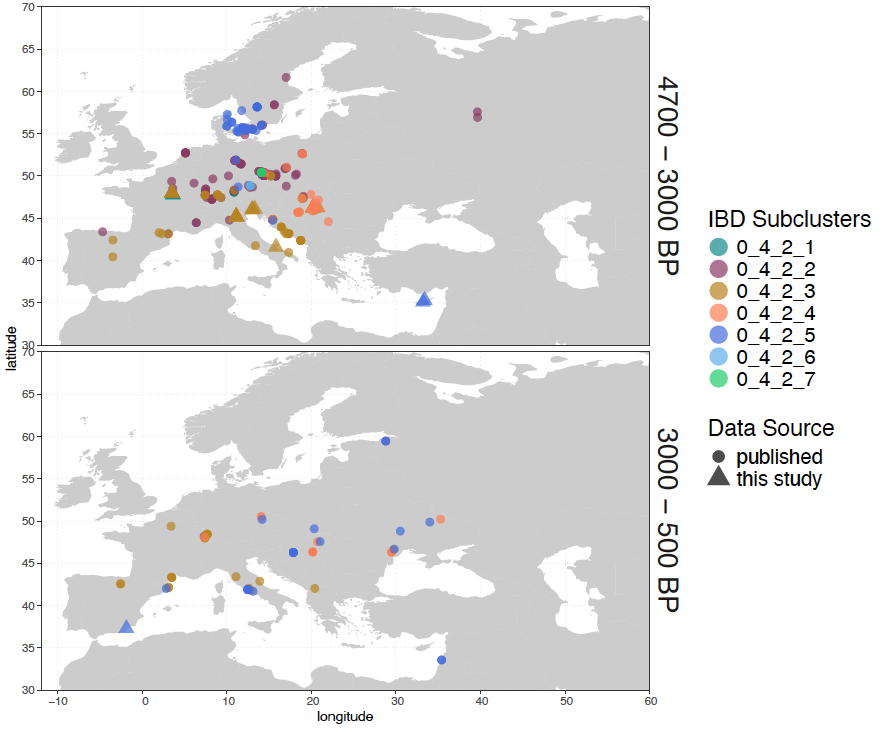

Fig. S5.10. Spatial and temporal distribution of the individuals within subclusters of 0_4_2.

### S6. IBD mixture modelling

Fulya Eylem Yediay, Hugh McColl, Martin Sikora

IBD admixture modelling is a method combining allele-matching profiling^29^ and chromopainting^91^ using the shared IBD length instead of allele frequency to get a better resolution for distinguishing especially genetically similar populations. Before aggregating the IBD sharing, we filtered out IBD segments of less than 1 cM, with no limit to the upper bound of total shared segments^1,2^.

For modelling, we first generated a palette with different source populations and outgroups based on shared IBD length^2^. With this palette, we can model the ancestry of target groups as a mixture of putative sources. As explained in Section S4, ancient individuals are grouped with the use of network clustering to select the individuals within the shared clusters for both deeper temporal (outgroups) and proximal populations. To model target individuals, we use a base model, we include all deep ancestral source populations (Supplementary Table S4; Fig. S6.1), and then add more proximal source populations.

#### S6.1 Defining farmer sources

As shown by the distribution of the individuals based on IBD clusters in section S4, we detected that all individuals from farmer groups split into three different subclusters (0_1_1, 0_1_2, 0_1_3). In this base model, we only have the Pre Pottery farmer group from Boncuklu as a deep ancestral population to all Europeans. We added more farmer sources to investigate the differentiation of farmer ancestry in the later populations. With the base model (Set1; Table S6.1), the farmer groups in subcluster 0_1_1, which represents European Middle and Late Neolithic individuals, are modelled with Anatolian farmers (Boncuklu) and elevated Western-Hunter-Gatherers (WHG) compared to farmer groups of subcluster 0_1_3, which are modelled either only with Early Anatolian farmers or a small proportion of Balkan Hunter-Gatherers (Fig. S6.3; S6.4; Supplementary Table S5). In Central Anatolia, we replicated the small proportion of CHG proportion found as early as ~9,800 BP (Asikli and Musular)^36^, and also detected in a few individuals from Barçın (I1103, I1098, I0708)^45^. Apart from those with a small CHG proportion, all Barçın individuals were fully painted with Boncuklu as in early Greek farmers and LBK-related (Linearbandkeramik) farmers. Subsequently, we added a second Anatolian farmer source from Barçın (Northwestern Anatolia) on top of our base model (Set2; Table S6.1; Supplementary Tables S4 and S5) and observed that the farmer proportion in Early Europeans and Central Anatolian farmers was replaced with Barçın. However, there was some noise of a small proportion of Boncuklu ancestry in individuals from the Chalcolithic and Bronze Age Anatolian, which could suggest that another farmer source would better fit. Similar noise was also detected in the European farmers after ~7,000/6,500 BP from the “European farmer (0_1_1)” cluster (Fig. S6.3; Supplementary Table S5). We established another model (Set3; Supplementary Table S4) with the Central Anatolian farmer group from Tepecik which already carry CHG, and this model replaced the farmer proportion in Anatolian Chalcolithic and later period individuals, as well as the region between the Southern Caucasus/Zagros and the Levant (Fig. S6.5; Supplementary Table S5). Additionally, a mixed component of two farmers (Barçın and Tepecik) is first observed in Neolithic Greece in 6,500 BP, and this pattern is similar to Early Central Anatolian farmers from Çatalhöyük and Musular, either pointing to another farmer population formation or indicating that the model cannot differentiate between these two closely related sources.

In the model Set4 (Supplementary Table S4), we started including European farmers with Italy Early Farmers, revealing that the farmer proportion of the European Middle Late Neolithic (in cluster 0_1_1) was completely replaced with the Italian Neolithic source (Fig. S6.6; Supplementary Table S5). Lastly, we added Globular Amphora and France Middle Neolithic Farmers, which fall within cluster 0_1_1 (Set5; Supplementary Table S4). This model distinguishes the southern-western farmer groups from Italian farmers. A similar pattern was observed in Central Eastern and Northern Europe after ~6,500 BP, suggesting a new formation of farmer populations (Fig. S6.7; Supplementary Table S5).

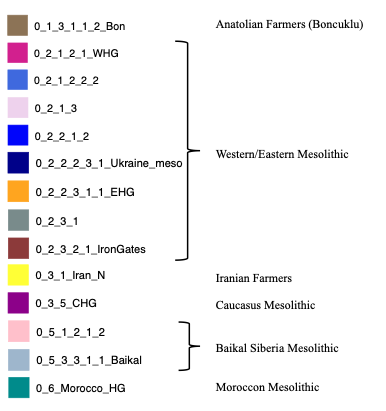

Fig. S6.1. The cluster of the base source populations and represented colours. Individuals in these clusters are given in Supplementary Table S4.

| Set1 | Base |
| --- | --- |
| Set2 | Base + Barçın |
| Set3 | Base + Barçın + Tepecik |
| Set4 | Base + Barçın + Tepecik + Italy_N |
| Set5 | Base + Barçın + Tepecik + Italy_N + Globular Amphora + France_MN |

Table S6.1. Farmer models set populations.

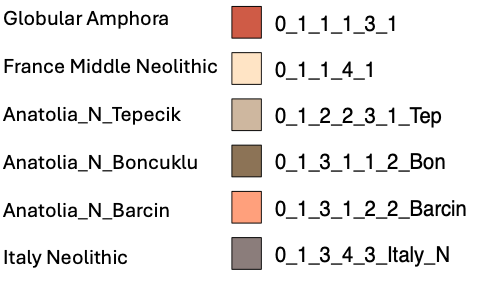

Fig. S6.2. The cluster of Farmer source populations. Individuals in these clusters are given in Supplementary Table S4.

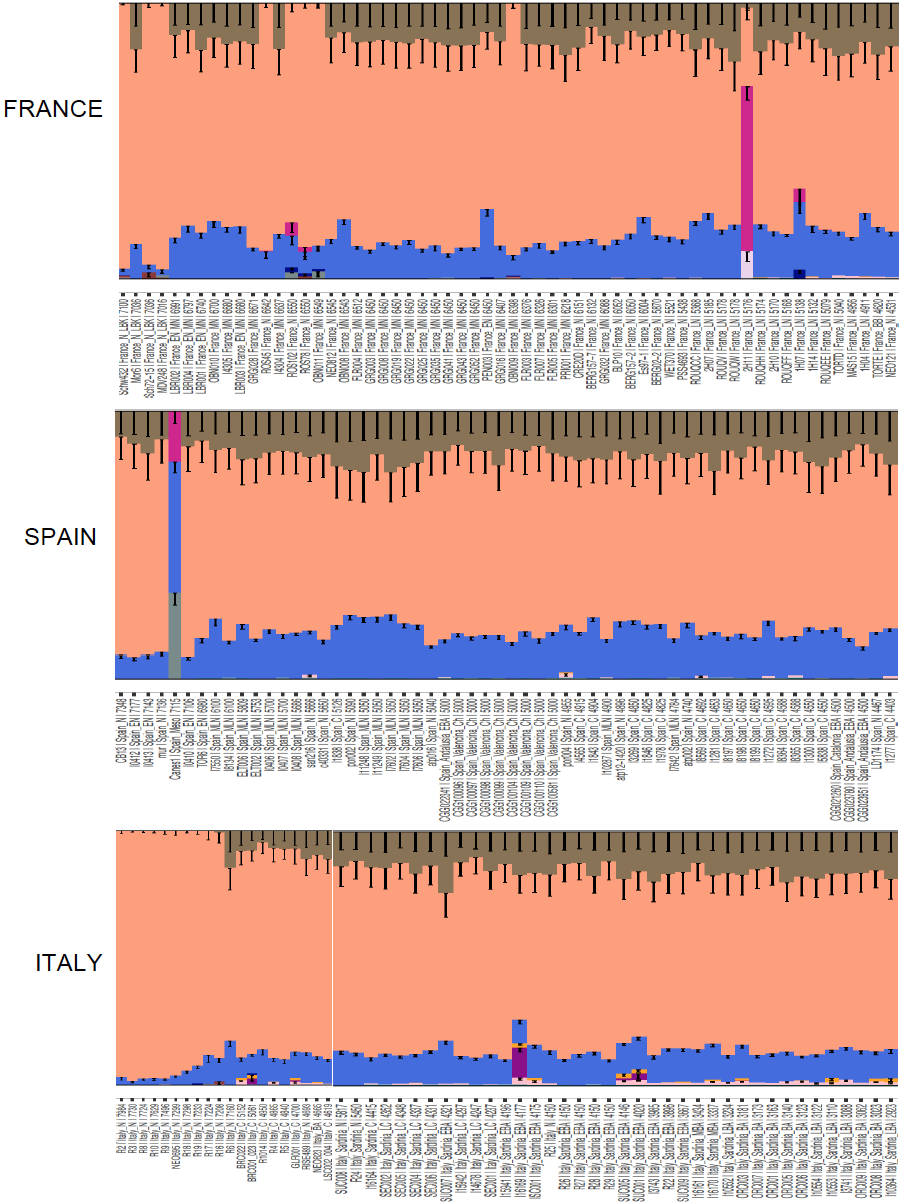

Fig. S6.3. Bar plot generated for each individual from Southwestern Europe farmers using source proportion from Set2.

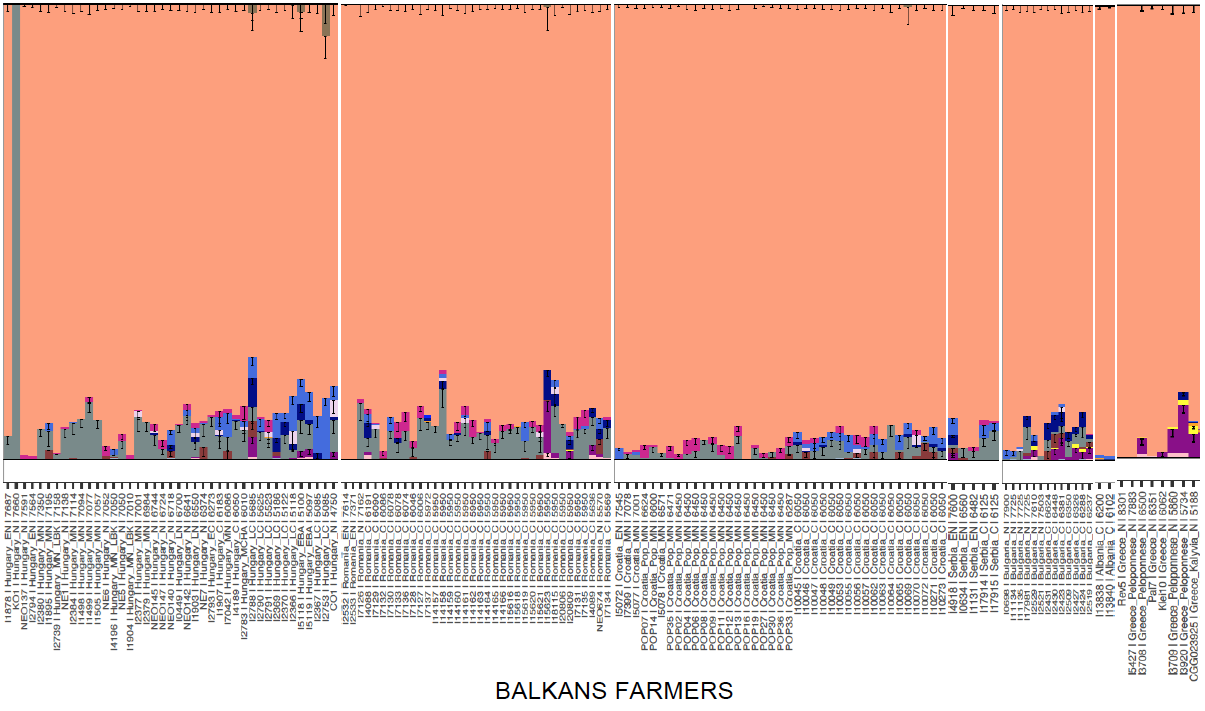

Fig. S6.4. Bar plot generated for each individual from Balkan farmers using source proportion from Set2.

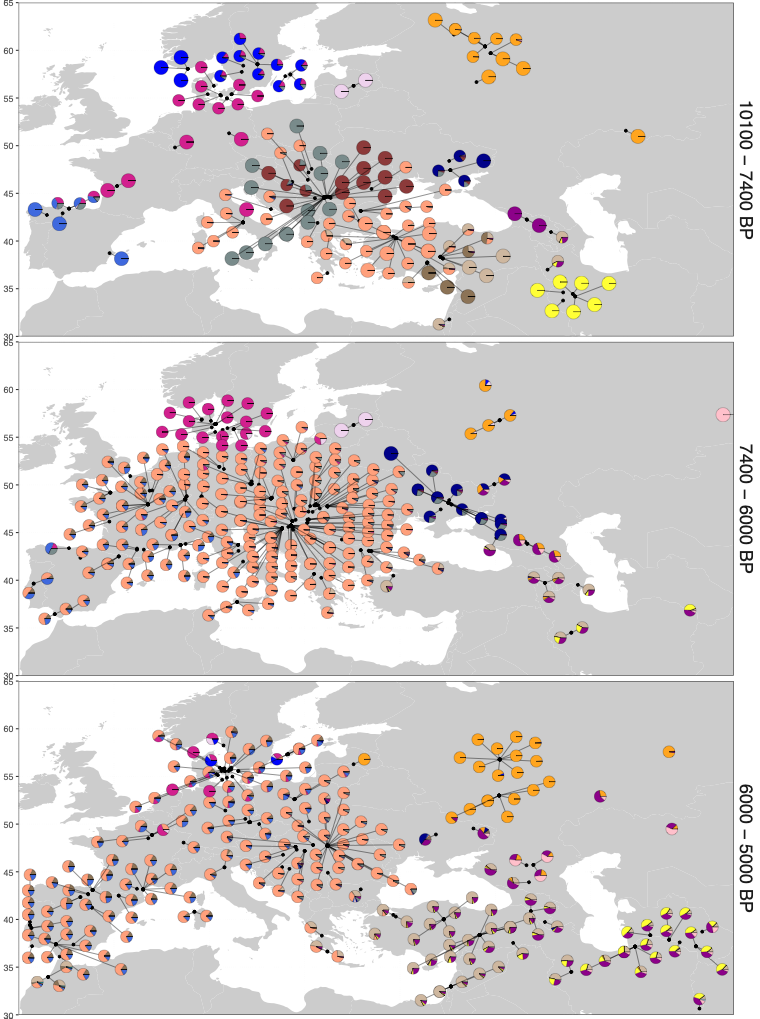

Fig. S6.5. Pie chart generated for each individual by using the proportion of sources from the Set3 admixture model, divided into three time periods; 10,100 BP–7,400 BP, 7,400 BP–6,000 BP, 6,000 BP–5,000 BP.

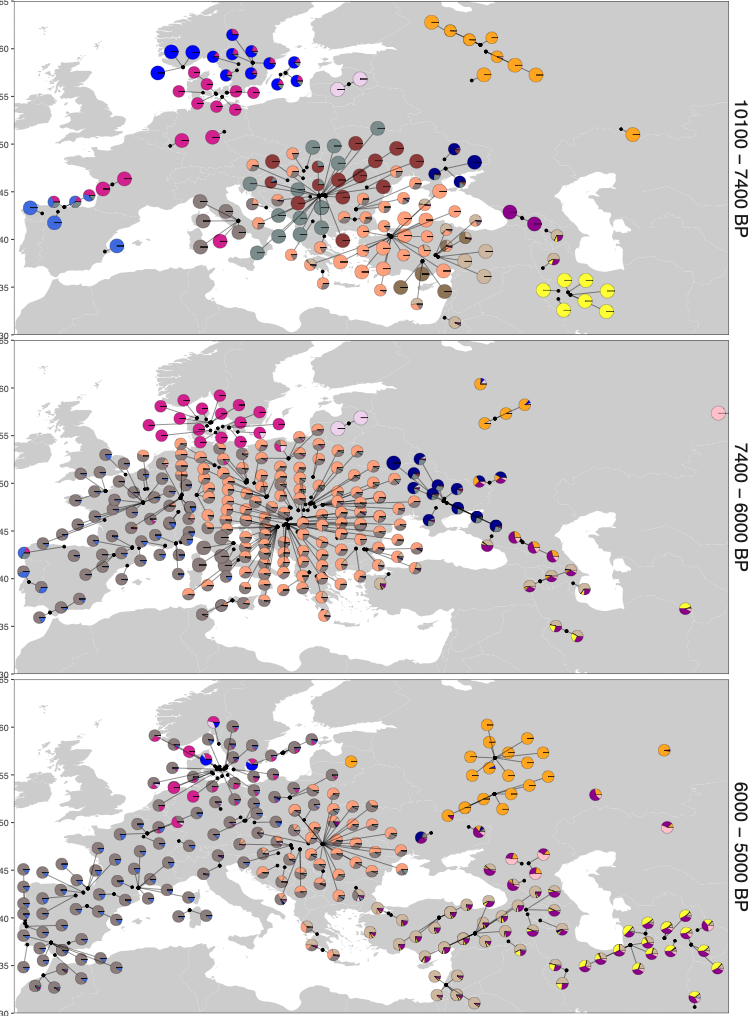

Fig. S6.6. Pie chart generated for each individual by using the proportion of sources from the Set4 admixture model, divided into three time periods; 10,100 BP–7,400 BP, 7,400 BP–6,000 BP, 6,000 BP–5,000 BP.

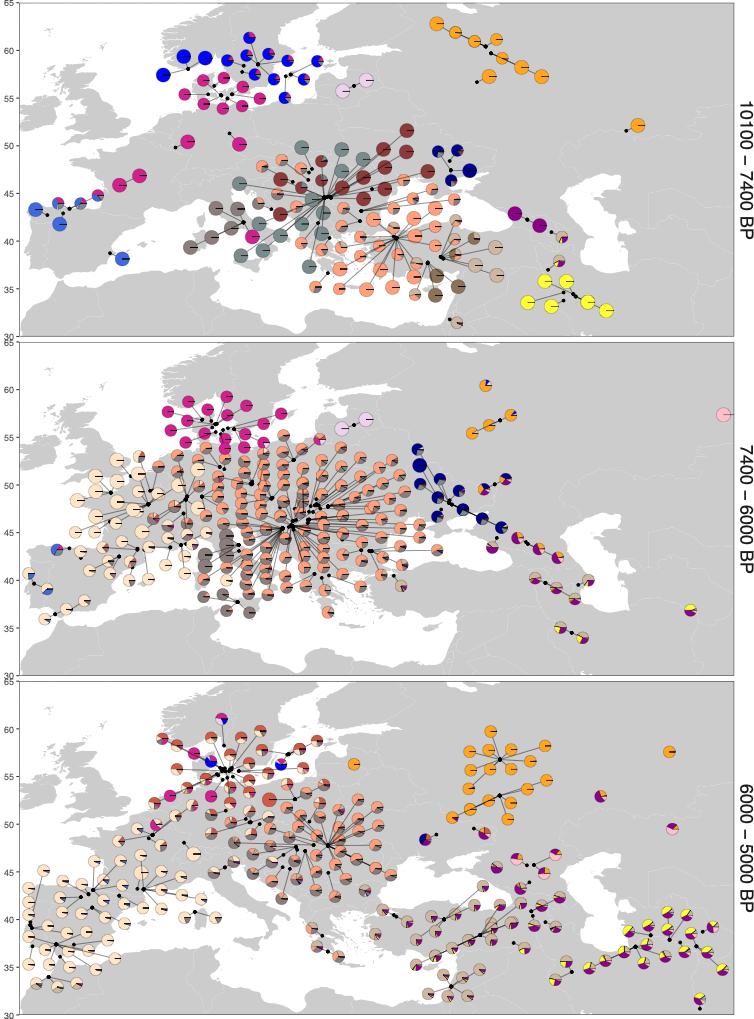

Fig. S6.7. Pie chart generated for each individual by using the proportion of sources from the Set5 admixture model, divided into three time periods; 10,100 BP–7,400 BP, 7,400 BP–6,000 BP, 6,000 BP–5,000 BP.

#### S6.2 Southern Europe

To model the European Bronze Age individuals, we selected a set of source populations (EU_Set1: Supplementary Table S4) that included three European farmer population groups: The Globular Amphora Complex, Italy Neolithic, and France Middle Neolithic. Additionally, we incorporated one eastern source, Anatolia Late Chalcolithic, and two steppe sources, Yamnaya and Early Corded Ware individuals (CWC). The selected Yamnaya cluster comprises individuals from Samara and Poltavka cultural context (Supplementary Table S4). The model EU_Set1 revealed that the steppe ancestry in Greece differs from Italy, France, and Spain by carrying a Yamnaya component. In contrast, Italy, Spain, and France, as well as most Western Europe Bronze Age individuals, are primarily modelled with Corded Ware ancestry (Fig. S6.8; S6.9; Supplementary Table S5). Additionally, we observed mixed CWC and Yamnaya ancestry in some Late Bronze Age individuals from Italy and the Balkans. This pattern of mixed CWC/Yamnaya composition might indicate steppe-related sources different from these two, or it could be the result of an admixture event between two populations who carry these ancestries.

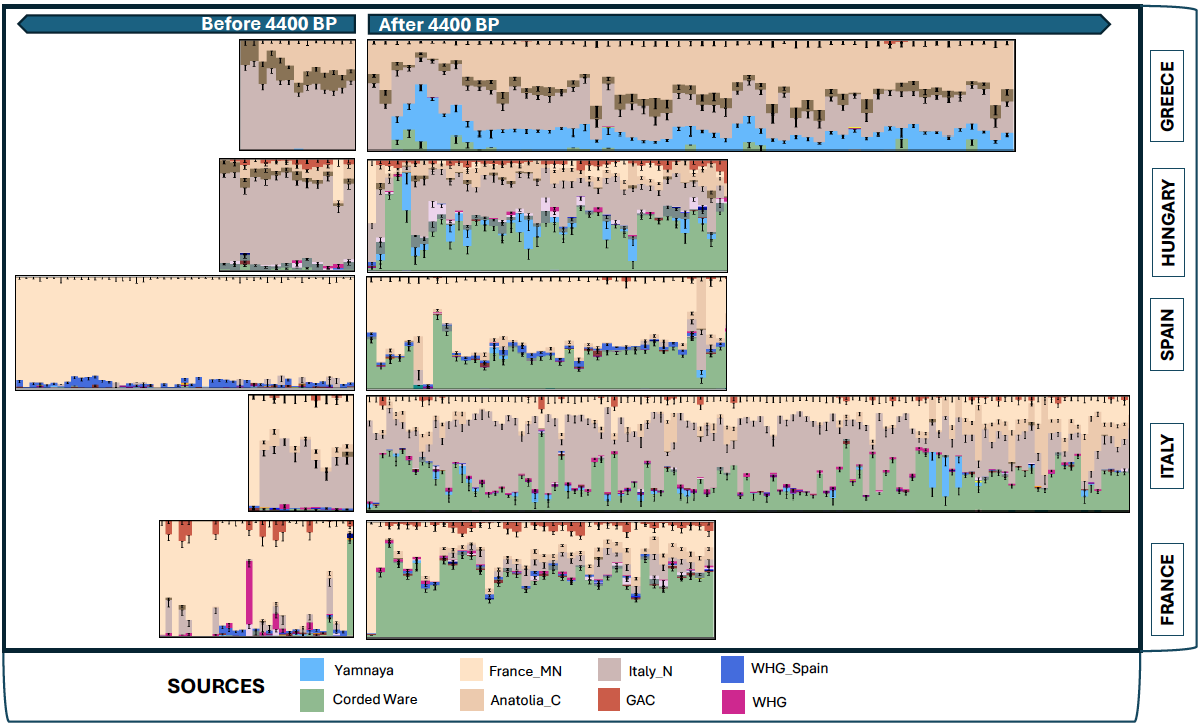

Fig. S6.8. Ancestry bar plots generated for each individual using source population proportions of IBD admixture modelling sorted by time BP and divided into two time series, before and after 4,400 BP, illustrating a Southern and Central Eastern Europe split (Italy, France, Spain and Hungary vs Greece).

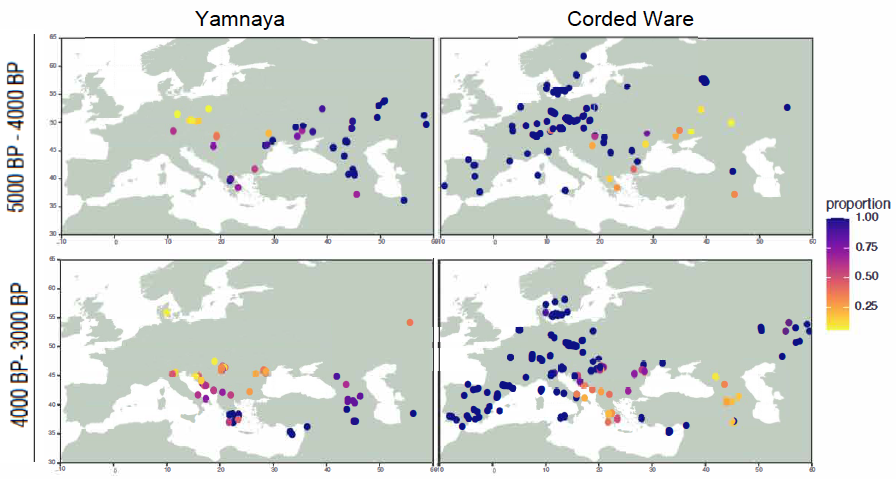

Fig. S6.9. Distribution of CWC-derived and Yamnaya-derived ancestry proportions obtained from the IBD admixture model. The standardized proportion presents the ratio of each ancestry component relative to total steppe ancestries.

**Italy**

The Italian Peninsula has a complex history with diverse populations characterized by unique cultural practices, even during the Neolithic period^11,54^. Genetic and cultural variation increases during the Bronze Age as a result of significant advancements in technology, economy, and the development of long-distance trade networks. Considering the diverse genetic make-up of Italian populations, we collected samples from Northern, Central and Southern Italy spanning the Middle and Late Bronze Age (3,800 BP to 2,900 BP) to shed light on the formation of the population structure and development of the Italic languages. The majority of Italian Bronze Age individuals analysed in this study were from Northern Italy and most densely sampled from Olmo di Nogara in the Po Valley. To answer the following questions, we set various admixture models:

1) Are there any regional differences regarding steppe ancestry throughout Italy, indicating migration from the Mediterranean, Western Europe or the Balkans into Italy?
2) Was the arrival/spread of steppe ancestry a population replacement or local admixture events with newcomers from CWC/Bell Beaker-related populations?
3) Is there any connection between mainland Italy and Western Mediterranean islands in terms of the spread of steppe-related ancestry through the Mediterranean?

We modelled Italy Bronze Age individuals by applying the following sets of source groups (source individuals are given in Supplementary Table S4):

EU_Set1; Base + Italy Neolithic, Anatolia Chalcolithic, GAC, Yamnaya, and CWC.

EU_Set2; Bell Beaker (BB) groups.

EU_Set3; France Middle Neolithic.

EU_Set4; Middle Bronze Age individuals from Northern Italy.

In EU_Set1, the steppe ancestry proportion was predominantly represented by CWC with the exception of a few individuals from Central Italy who displayed a different ancestral composition (Fig. S6.10; Supplementary Table S5). This composition was also observed in Balkan/Aegean Bronze Age populations. We also detected a small proportion of Globular Amphora Complex (GAC) ancestry present in European Middle Late Neolithic and Bronze Age populations. Interestingly, the steppe ancestry proportion varied significantly, indicating a diverse population structure even within a single site, Olmo di Nogara, during the Late Bronze Age. In a few individuals, we also detected a signal of Mediterranean admixture, indicated by the proportion of Anatolian Chalcolithic ancestry.

EU_Set2: Since we observed an additional GAC component in the first model, similar to that found in Bell Beaker-related groups, we included a BB source, mainly from Germany, which replaced all CWC components in the analysis. However, this model did not fully fit for some individuals from Northeastern Italy (CGG_2_022923, CGG_2_022253_022566, CGG_2_022653) and from Central-East Italy (CGG_2_100646, NEO806, R1), as shown in the PCA plot shifting to the Balkan cline (Fig. S4.1; S4.2; Supplementary Table S5). These individuals from Northeastern Italy carry an additional eastern component (the proximal source is Anatolia Chalcolithic) along with Yamnaya/CWC ancestry proportions, suggesting another steppe component. Meanwhile, three individuals from Central-Eastern Italy exhibit an increased proportion of Yamnaya and eastern sources, indicating potential gene flow possibly from Greece or the Balkans (Fig. S6.11; Supplementary Table S5). We also detected the presence of increased eastern ancestry in two undated (14C) individuals (CGG_2_022640, CGG_2_022641), possibly from later periods, and two individuals dated to a more recent period (~1200 BP) from Olmo (CGG_2_022616, CGG_2_022580). However, no differences were observed in the type of steppe ancestry, which is similar to the BB-related type. This ancestry composition with increased eastern ancestry suggests gene flow from the Eastern Mediterranean, particularly observed in the later periods^11^. We detected this eastern proportion in Central Italy Bronze Age individuals who carry BB ancestry, however we do not observe this proportion in two individuals from Coppa Navigatta (CCG_2_101264, CGG_2_101266), similar to individuals from Northern Italy.

EU_Set3: As some individuals still exhibit GAC ancestry, as well as small proportions of WHG ancestry, we introduced another farmer source group from France Middle Neolithic. This model supports the connection between individuals from the Adriatic coast and the Aegean/Balkans, as these individuals exhibit no proportion of France Middle Neolithic ancestry. Additionally, three individuals from Northeastern Italy, mentioned in EU_Set1, do not fit the model that includes the France Middle Neolithic source (Fig. S6.12; Supplementary Table S5). We also observed little or no proportion of Italy Neolithic ancestry in some of the genetic outliers, contrasting the rest of Italy Bronze Age individuals, conversely increasing the proportion of France Middle Neolithic ancestry (CGG_2_022591, CGG_2_022625, CGG_2_022595, CGG_2_022583, CGG_2_022569, GCP002). This could suggest that these individuals may have originated from other regions associated with BB-related populations with different farmer combinations.

EU_Set4: Finally, we selected two Middle Bronze Age individuals from Northeastern Italy as the local steppe sources (CGG_2_022253_022566 and CGG_2_022653). The steppe source in the model replaced the BB component in most of the Late Bronze Age individuals, but it did not fit for some individuals and rejected one outlier (CGG_2_022591), confirming that this individual has steppe ancestry of a different origin. Additionally, we observed that some individuals exhibited both components, suggesting that the model did not fit for those individuals and distinguishing them from the rest of the Italian individuals (Fig. S6.13; Supplementary Table S5). Moreover, this local source replaced the BB component in individuals from Corsica, Sicily, Albania, Croatia Middle and Late Bronze Age, and a few Late Bronze Age Hungarians (Fig. S6.14; Supplementary Table S5), suggesting shared ancestry with the source individuals from Northeastern Italy. These individuals were modelled either with BB or a mix of BB/Yamnaya components in the model EU_Set3, suggesting that they share a common steppe ancestry in comparison to other regions in Europe. Since we already detected these source individuals shifting towards the Balkan cline in the PCA, this could explain its consistency with Balkan/Adriatic individuals.

As we have already detected a lower proportion of steppe ancestry in most Late Bronze Age individuals from Olmo, this model might suggest local admixture events involving this source group or interactions between local farmers and the groups that share this steppe ancestry. Notably, this model did not fit for individuals from Central Italy except one (CGG_2_101266), a pattern similar to that observed in the previously published Bell Beaker individuals, Spain and France Bronze Age individuals (Fig. S6.14; S6.15, S6.17; Supplementary Table S5).

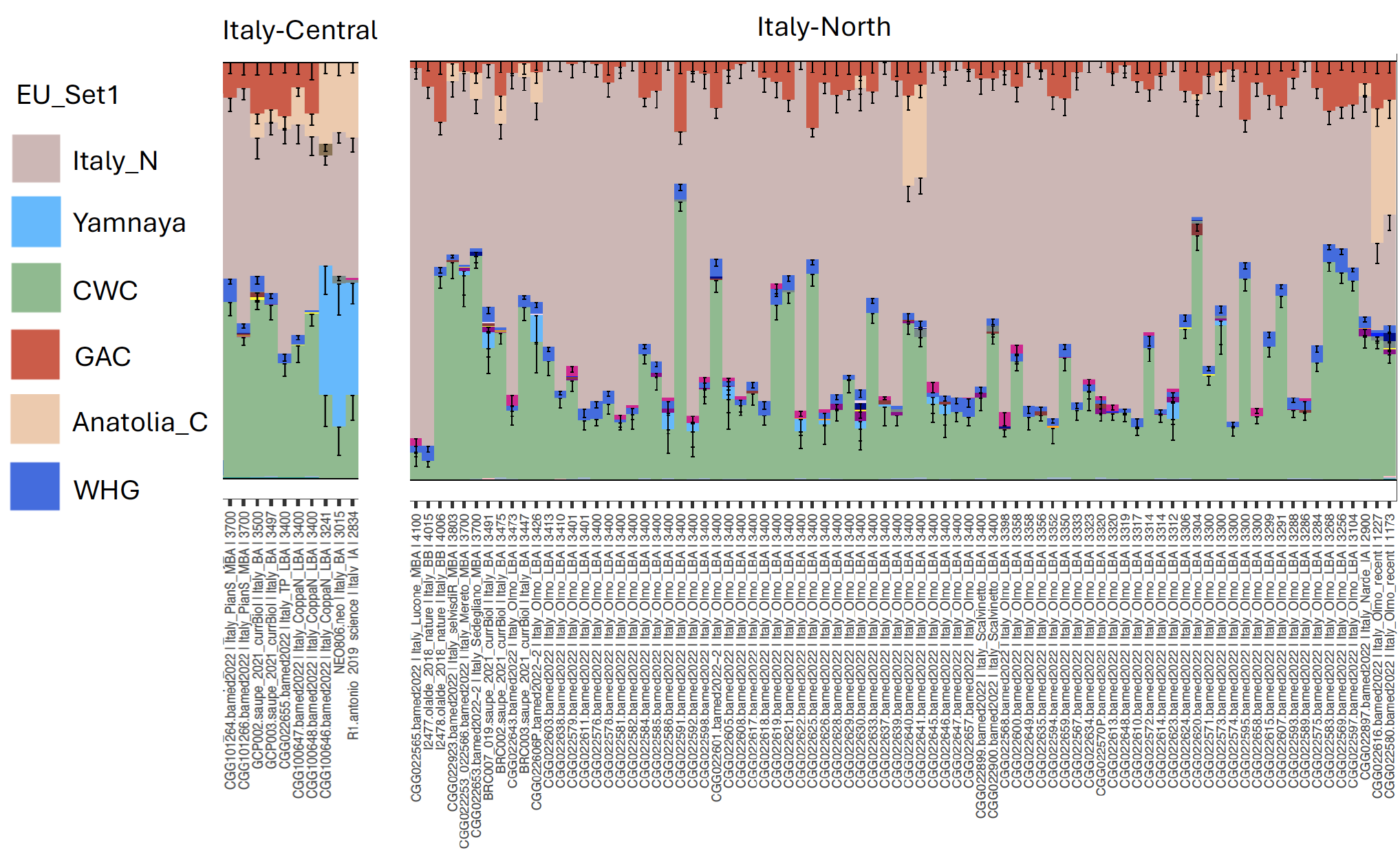
Fig. S6.10. Bar plots generated by EU_Set1.

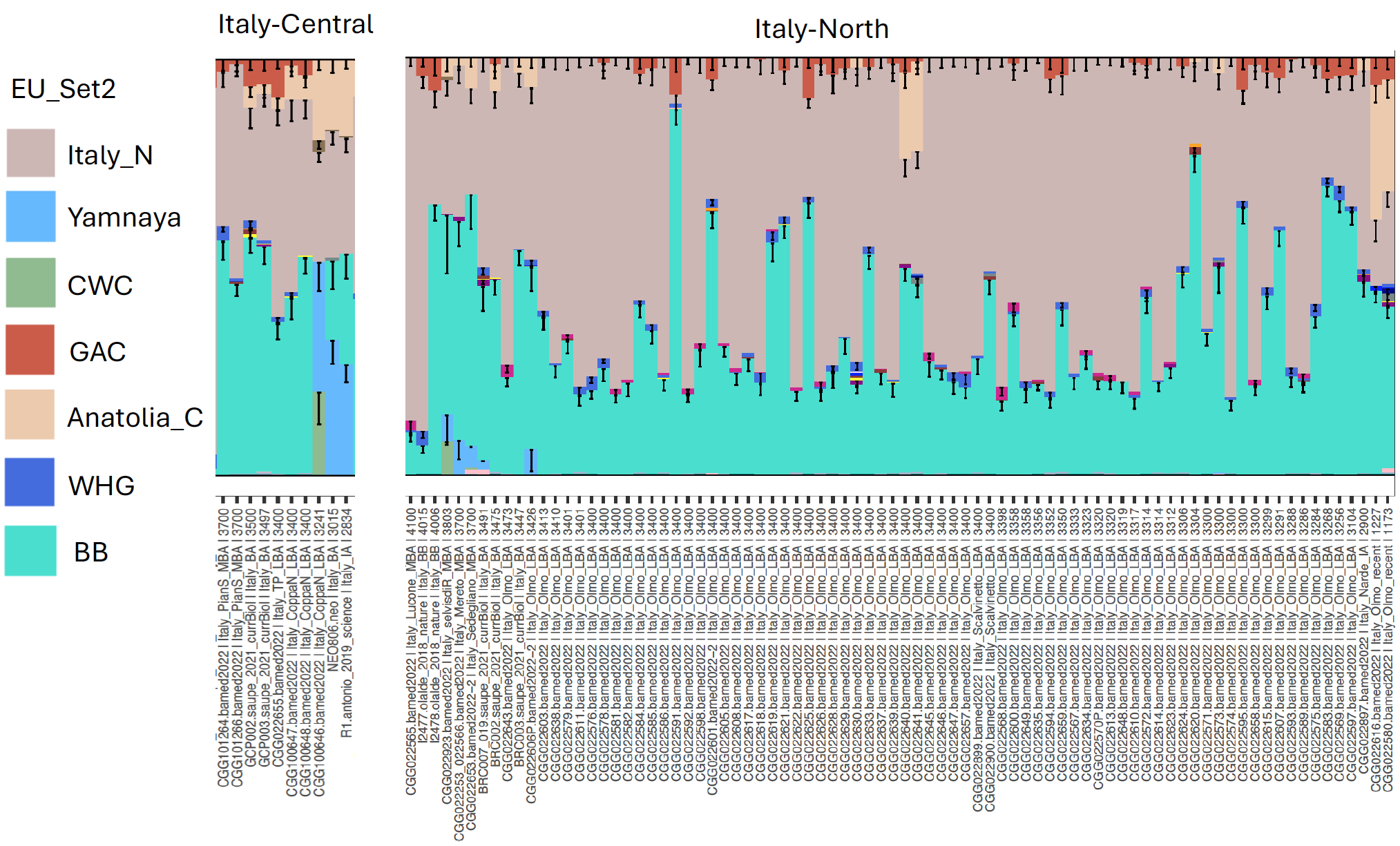

Fig. S6.11. Bar plots generated by using ancestry proportions obtained from EU_Set2, ordered by time.

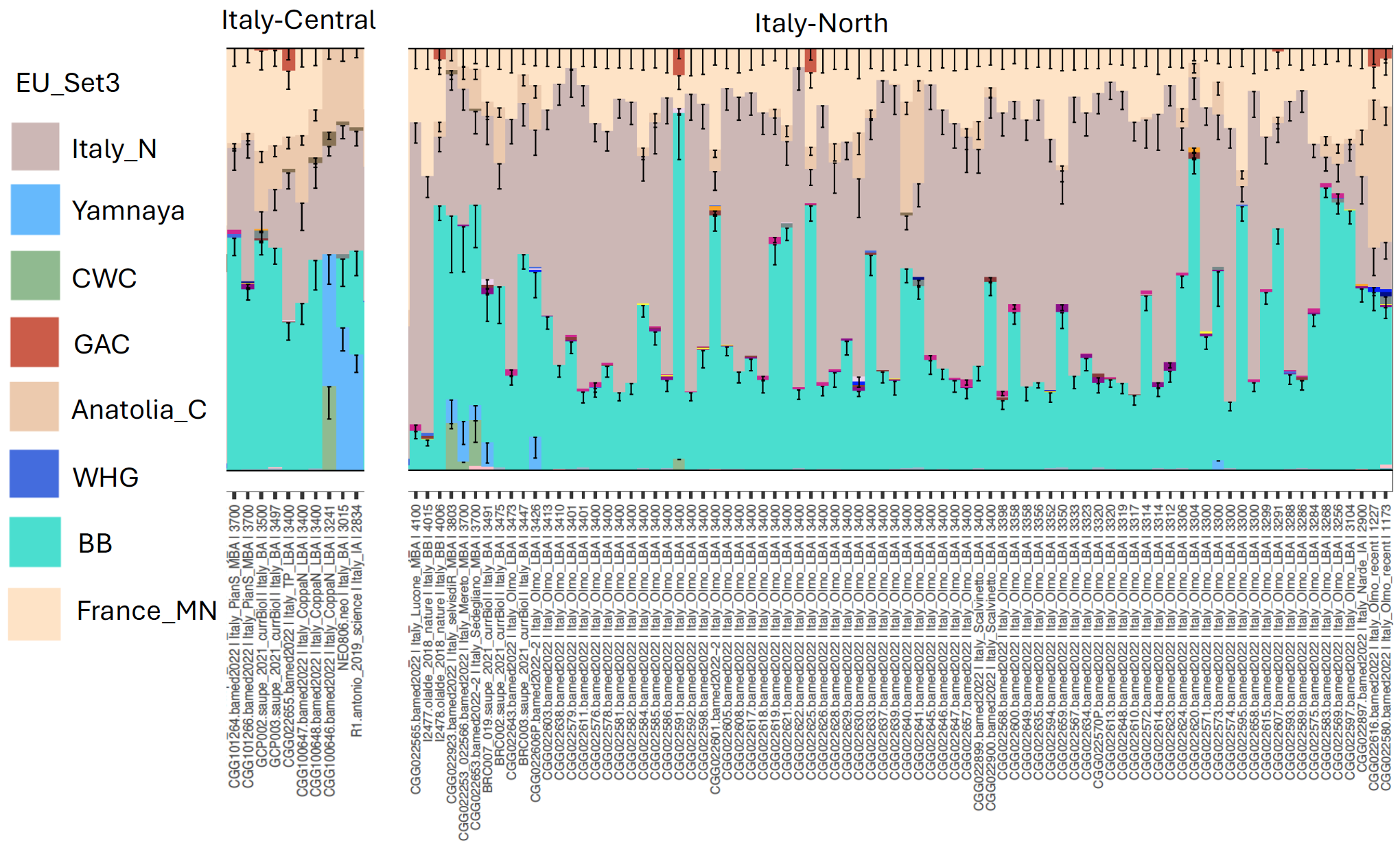

Fig. S6.12. Bar plots generated by using ancestry proportions obtained from EU_Set3, ordered by time.

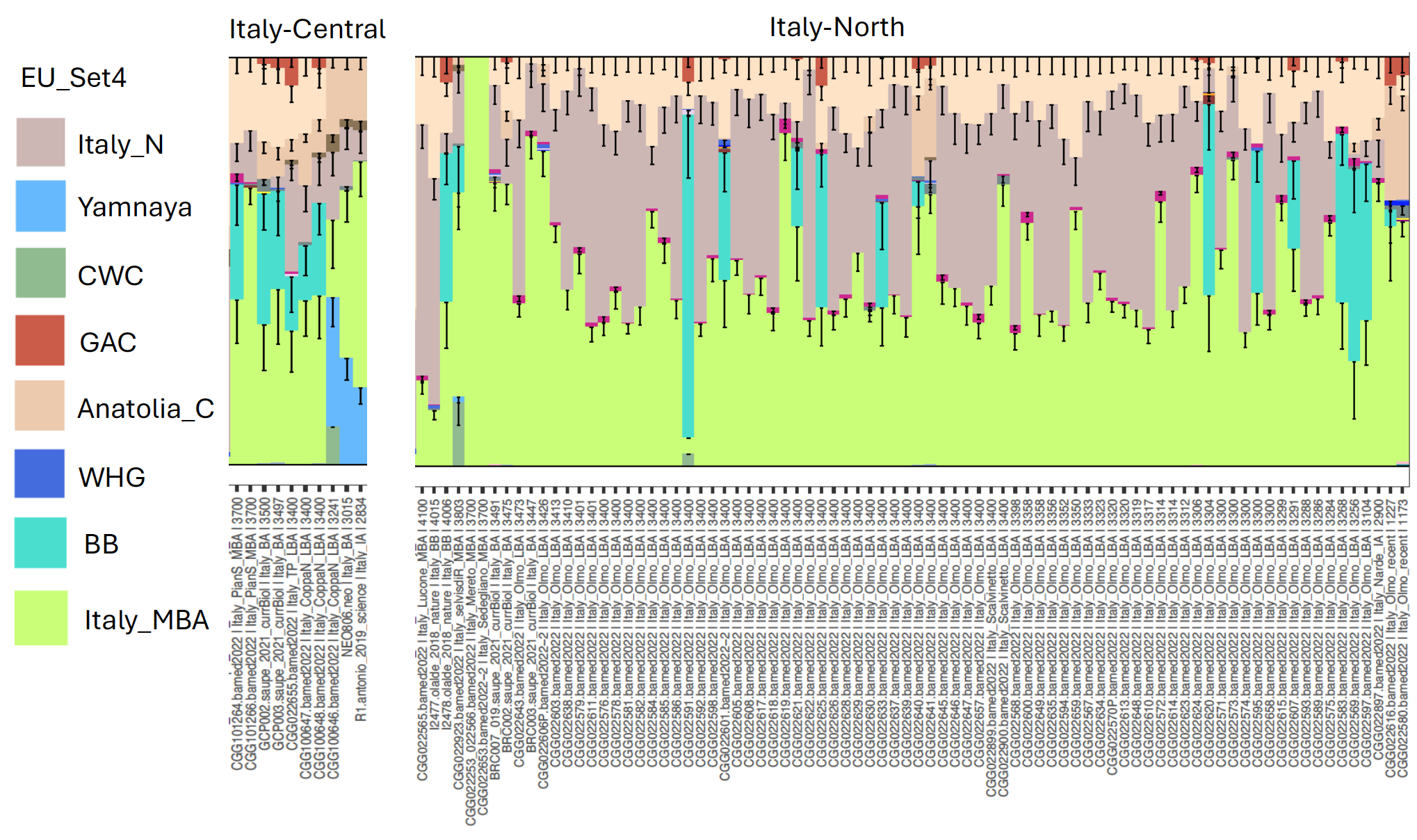

Fig. S6.13. Bar plots generated using proportion of source populations obtained in EU_Set4.

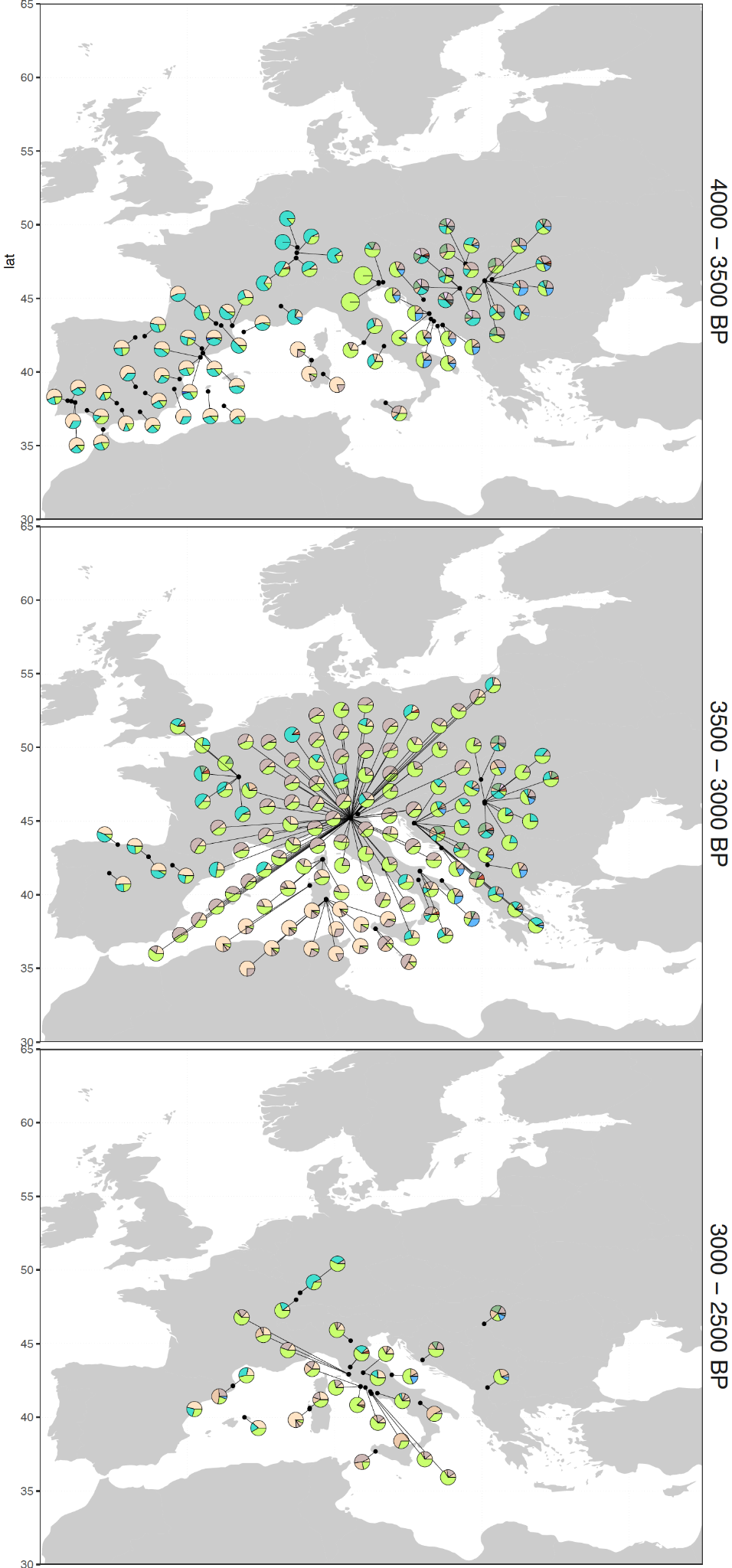
Fig. S6.14. Pie chart representing all individuals from Southern Europe and the Balkans generated by using source proportions in EU_Set4.

**Spain, France**

The arrival of steppe ancestry and its regional variations in Spain has been previously documented and associated with the Bell Beaker culture^44,49,50^. Similar to Spain, steppe ancestry in France shows higher levels in the north compared to southern regions^49^. To investigate the differences of the steppe ancestries and their connection to the Mediterranean, we analysed 17 Early Middle Bronze Age individuals across the entire coastal region of Spain (including nine Chalcolithic individuals from Valencina) and 19 Late Bronze Age individuals from Central France (Migennes) and Corsica (Supplementary Table S1).

As we have already created four model sets in Section S6.2, we focus on the model EU_Set3 to investigate the population structures of Spain and France. In this model, Spain and France show a similar steppe ancestry, and both modelled with Middle Neolithic France farmers and Bell Beaker groups (Fig. S6.15; S6.16; Supplementary Table S5). However, the steppe proportion is higher in France compared to Bronze Age individuals from Spain. Additionally, some Early Bronze Age individuals from France along with one individual from Spain (NEO649) show an additional CWC proportion, suggesting a link to other Bell Beaker groups. Since these individuals are from the northern region of both countries, this ancestry may be linked to Bell Beaker groups from the Netherlands. In the Late Bronze Age, the individuals from Migennes differ slightly from the earlier individuals by displaying an extra/another farmer component similar to Italy Bronze Age individuals. It is evident with the model EU_Set4 that the Italian source proportion increases in the Late Bronze Age; moreover, one individual (CGG_2_021374) carries the highest proportion of this source. In contrast, the Late Bronze Age individuals from Corsica show a similar pattern by exhibiting both farmer ancestry from Italy and France, and completely replacing the Bell Beaker component with Italy Bronze Age ancestry in EU_Set4 (Fig. S6.15). The model EU_Set4 did not fit for Spain Bronze Age individuals, although it replaced the steppe proportion of the previously published Aegean outlier (I8215)^50^ (Fig. S6.17), which is consistent with the Balkan/Adriatic connection shown in Fig. S6.14. On the other hand, in the newly sequenced Chalcolithic individuals from Spain, we do not observe a significant difference from the Spain farmer populations; however, they have less or no WHG component compared to other Chalcolithic individuals from the region (Fig. S6.16; Supplementary Table S5).

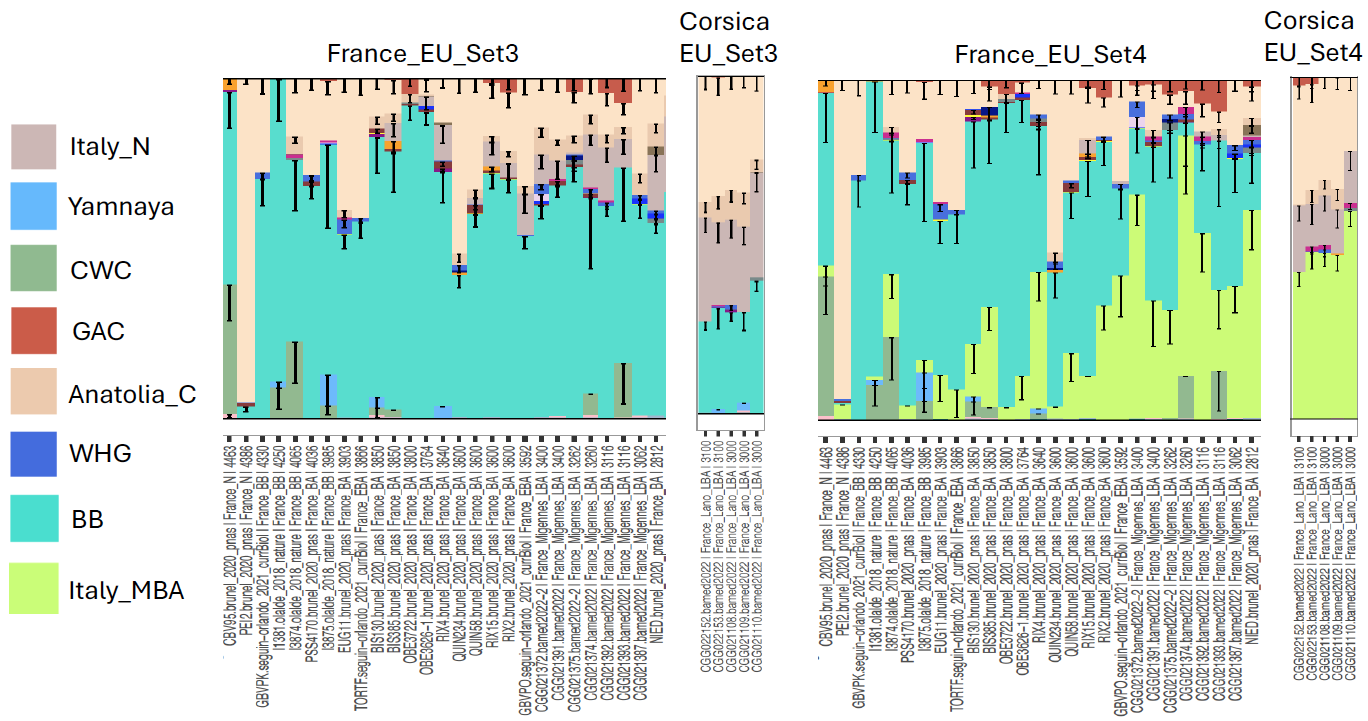

Fig. S6.15. Bar plot generated by using the source proportions obtained from EU_Set3 and EU_Set4 for France individuals.

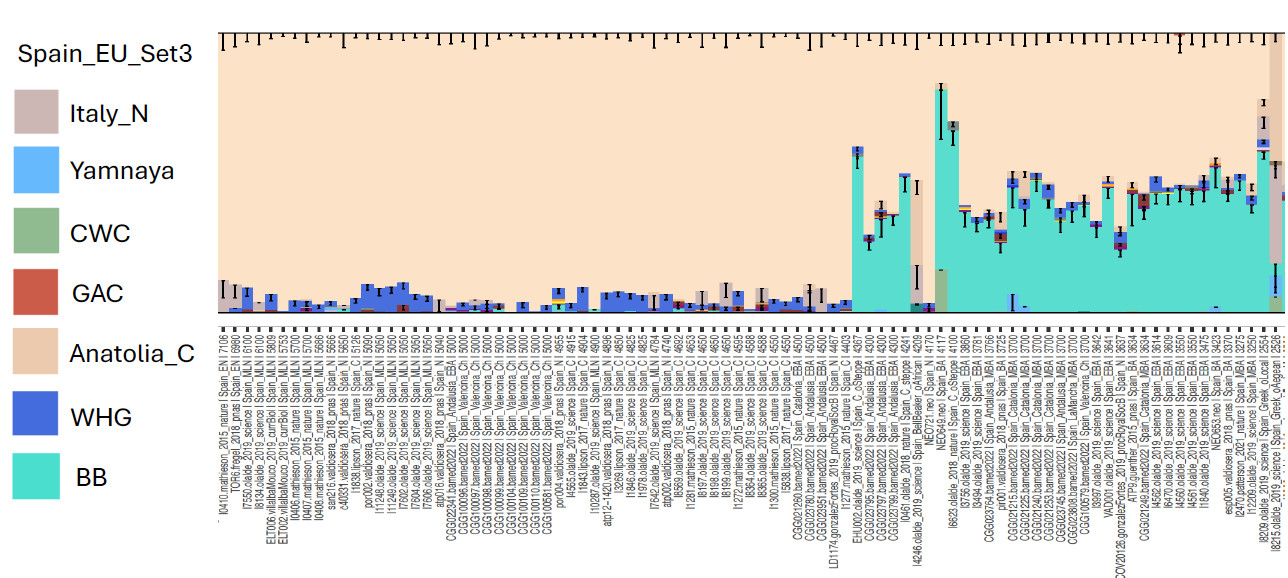

Fig. S6.16. Bar plot generated by using the source proportions obtained from EU_Set3 for Spain individuals.

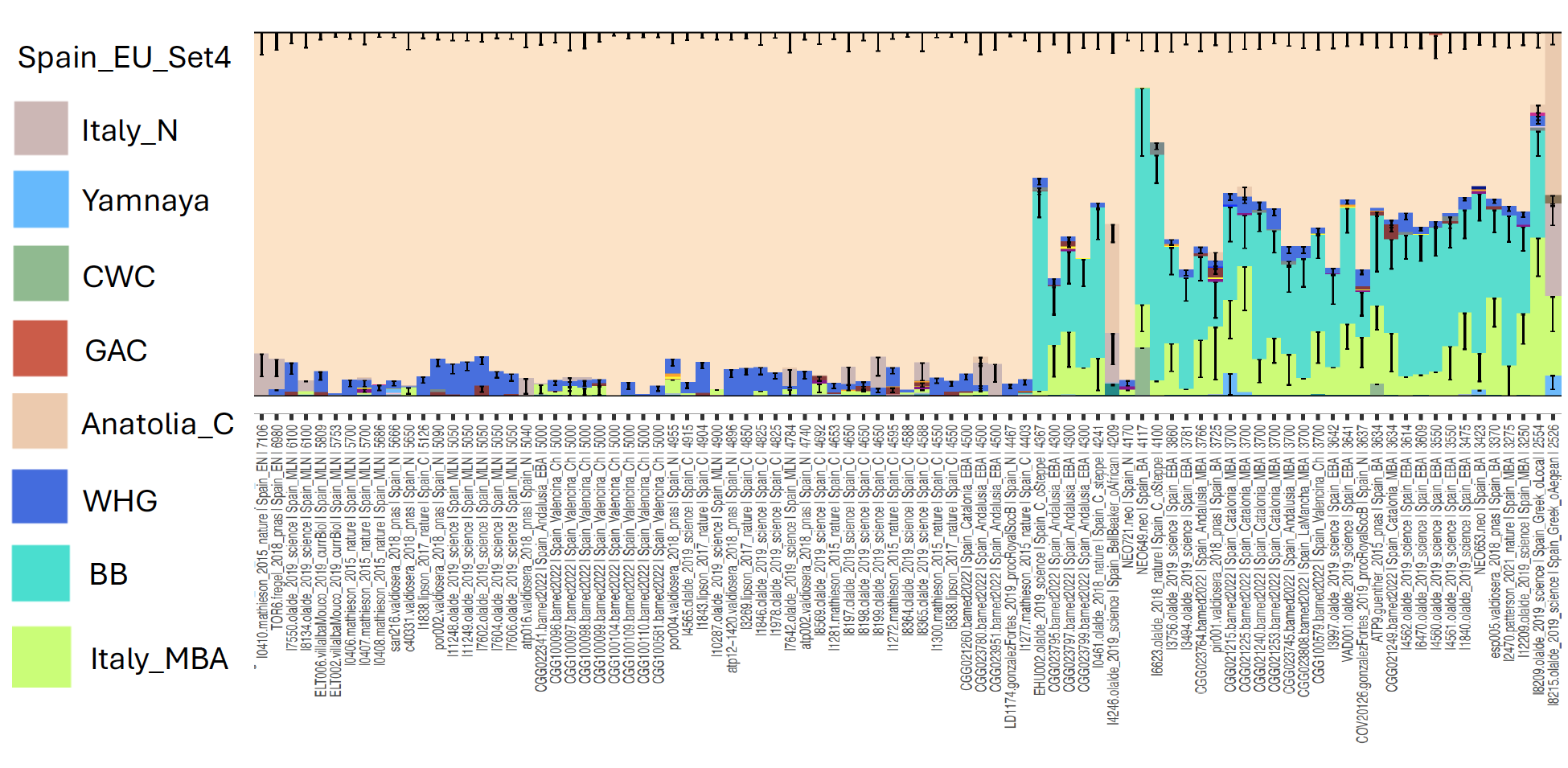

Fig. S6.17. Bar plot generated by using the source proportions obtained from EU_Set4 for Spain individuals.

**Greece**

Steppe ancestry arrived in Northern Greece in ~4,200 BP^14^ and reached Central Greece and Crete^36,92^ in the Middle and Late Bronze Age. Here, we present dense population data from Central Greece and the Peloponnese covering this time period to better understand population formation. Following the basic admixture model EU_ Set1 (Section S6.2; Fig. S6.8), the steppe ancestry in Bronze Age Greece (both Middle Helladic and Mycenaean periods) stands out from the European Bronze Age by showing the pattern of another steppe source, in this model by carrying Yamnaya (Samara) ancestry. Moreover, we observed a sharp change in the Bronze Age in which the introduction of steppe ancestry in individuals between 4200 BP and 3800 BP (Greek Middle Helladic and Middle Bronze Age) differs from the later period samples corresponding to the Mycenaean period (LBA: 3,700 BP–3,200 BP) that have a higher steppe proportion.

In addition to model EU_Set1, we built source sets to model Greece Bronze Age specifically. In Greece_Set1, we replaced the farmer source with Late Neolithic and Early Bronze Age populations from the Peloponnese and kept only Yamnaya as steppe source and also added GAC (Fig. S6.18; Greece_Set1; Supplementary Tables S4 and S5). In this model, we observed an elevated CHG proportion after ~3,600 BP, then we added an eastern source which is Anatolian Chalcolithic (Çamlibel) who already possessed the CHG proportion. The Greece_Set2 revealed that the Anatolian Chalcolithic source completely replaced the CHG proportion in Greece Bronze Age individuals (Fig. S6.18). Then we added CWC individuals and observed the same pattern with Greece_Set2 except with a few individuals (Fig. S6.18). In the model with two steppe sources (Yamnaya, CWC in Greece_Set3; Fig. S6.18; Supplementary Tables S4 and S5), we also detected a small proportion of CWC in a few individuals with lower error bars. Potential explanatory scenarios for this could include another steppe source from an unsampled steppe population or interaction with populations from Central Eastern Europe. Another explanation is that we could not find the right farmer source for those individuals. However, when we added BB groups, a small BB proportion in one individual from Kirrha (CGG_2_022403, ~3,800 BP) indicates a contact with the populations from the Balkan Bronze Age that exhibit a mix of BB and Yamnaya proportions (Fig. S6.19; Supplementary Table S5). This individual is also shown in the PCA plot shifting towards the Balkan cline, and clustered with Hungarian Bronze Age individuals.

To investigate Greece more deeply, we also tested local Early Bronze Age populations as sources (Fig. S6.20; Supplementary Table S5). Among the Early Bronze Age Greece sources, one group from island sites (Koufonisia and Petras) (Greece_Set 5; Supplementary Table S4) modelled the Late Bronze Age individuals especially from Voudeni better than the rest by replacing the farmer proportion. In Greece_Set6, we included the Early Bronze Age from the mainland that showed a better source for almost all individuals, but could not replace the CHG proportion as in Greece_Set5. Lastly, we added Minoans as a source and obtained a result with a slightly higher CHG proportion and high error bars (Greece_Set7; Supplementary Table S4). The CHG component in Greece would be replaced by either Anatolian Chalcolithic or populations from the northeast of the Aegean Sea.

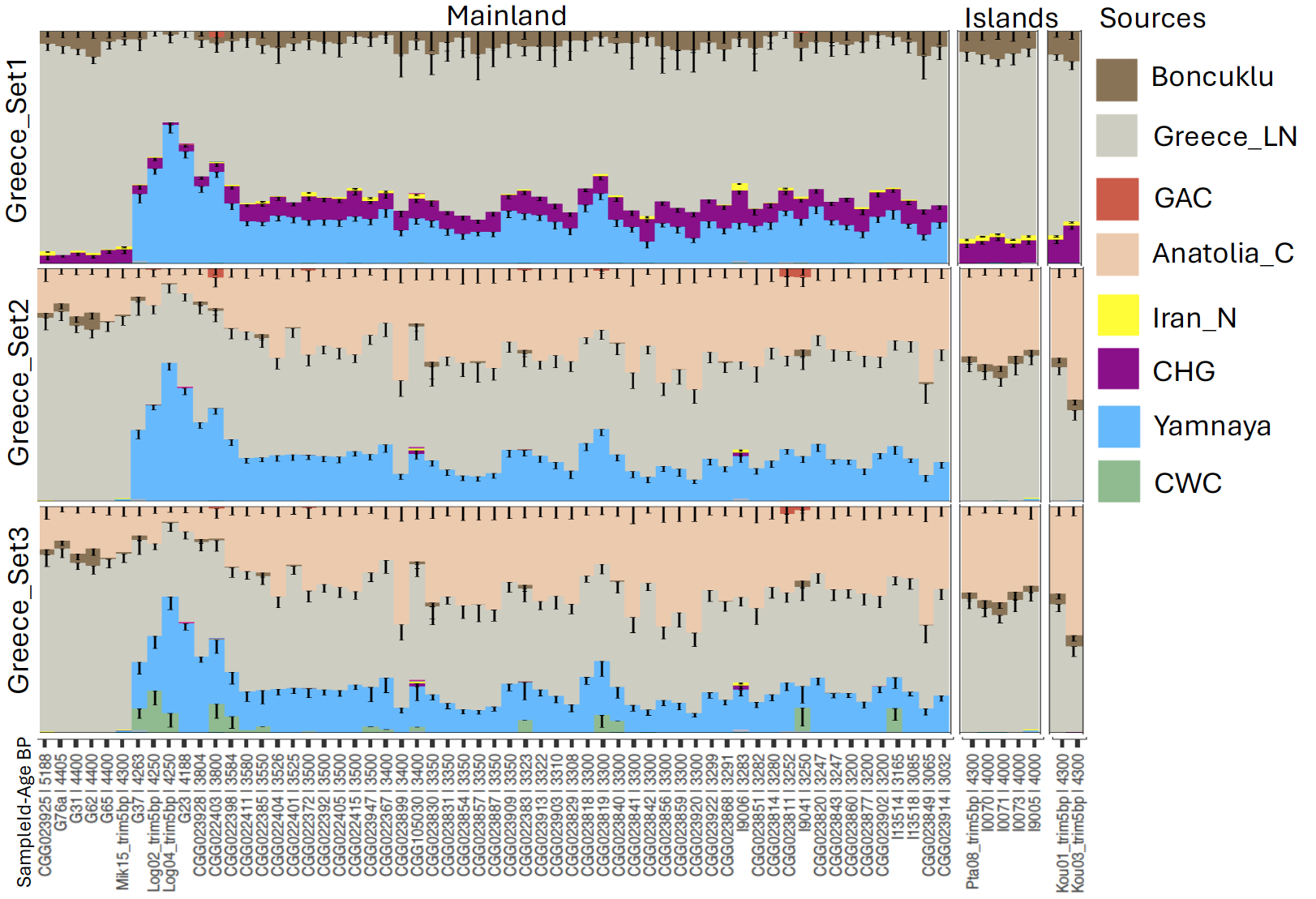

Fig. S6.18 Bar plots generated by using source proportions of Greece_Set1, Greece_Set2, and Greece_Set3 and sorted by Age BP.

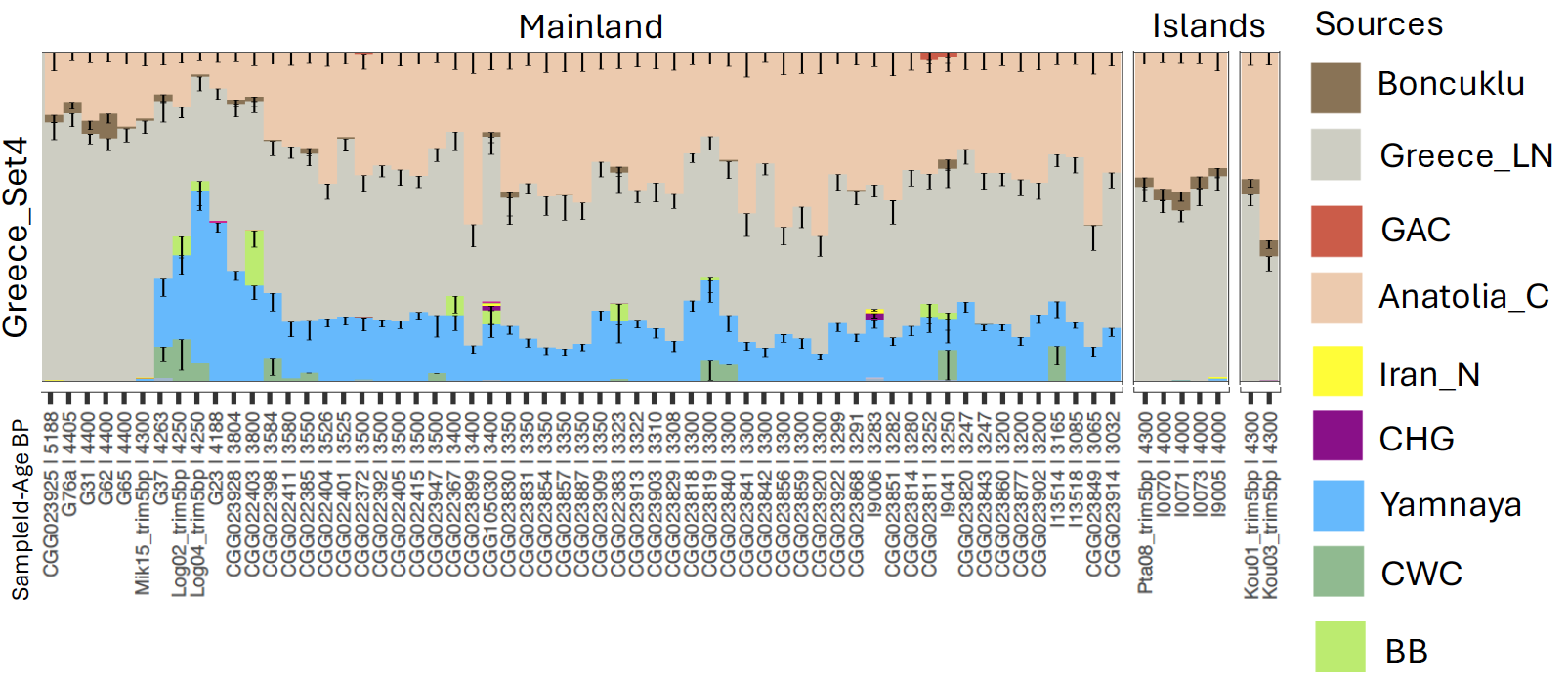

Fig. S6.19. Bar plots generated by using source proportions from Greece_Set4.

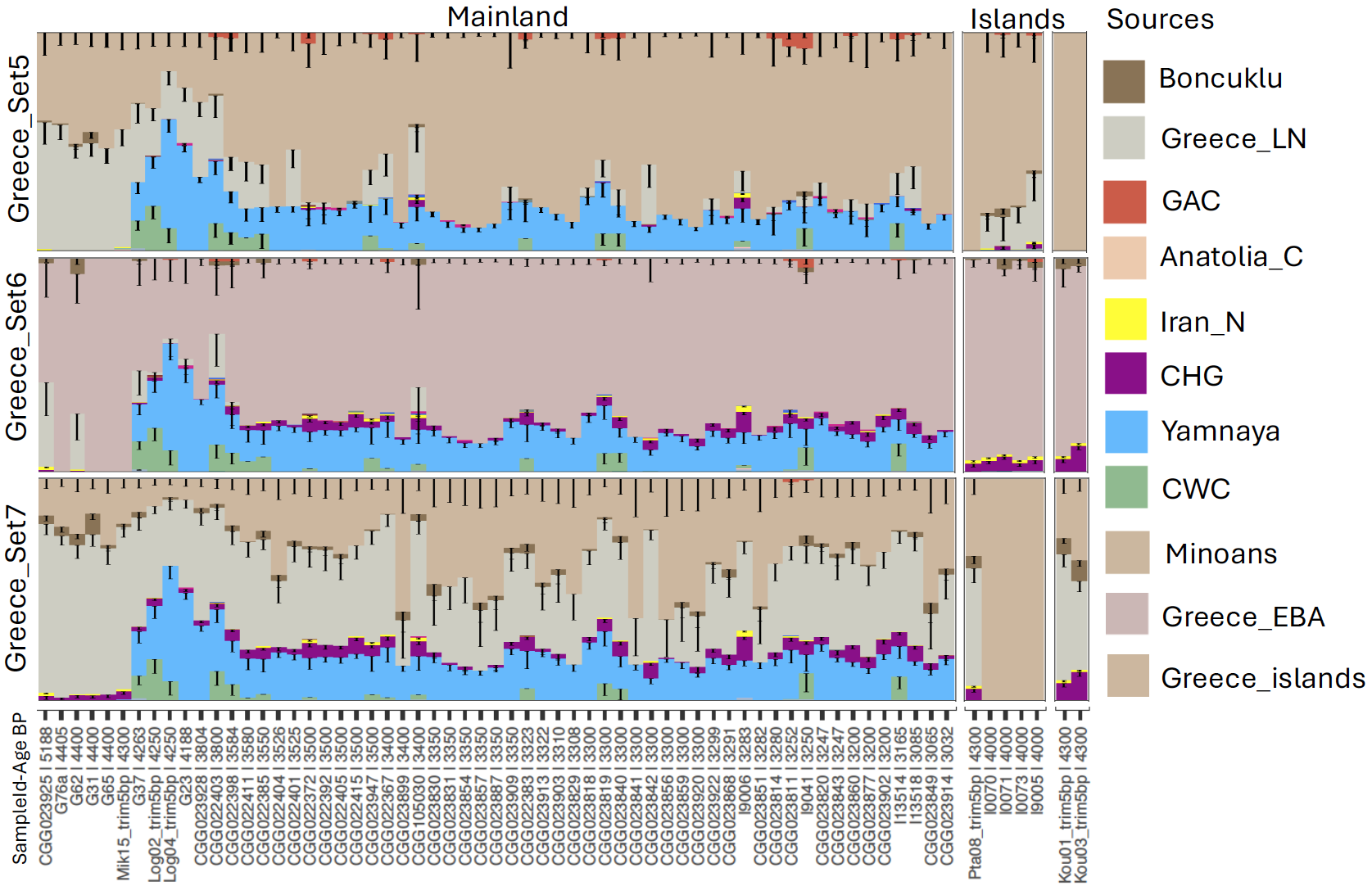
Fig. S6.20. Bar plots generated by using source proportions from Greece Early Bronze Age source population sets and sorted by Age BP.

Since two Middle Bronze Age individuals clustered with Yamnaya groups (Log04, ~4,200 BP) and with Armenia Middle Late Bronze Age individuals (G23, ~4,200BP), we tested all population groups within subcluster 0_4_3 as a potential representer of source from the Pontic steppe populations (regardless of whether labelled as Poltavka, Catacomb, Srubnaya, Ukraine_Yamnaya, etc.). None of these source groups replaced the Yamnaya proportion, but newly sequenced Early Bronze Age Moldovans (Late Yamnaya) within this cluster replaced the Yamnaya proportion in the Late Bronze Age individuals (Fig. S6.21; Supplementary Table S5). This could suggest that the Moldovans are a more proximal source for the later individuals in Greece. The earlier individuals (before ~3,800 BP) and some of the Late Bronze Age individuals were not modelled with Greece_Set8 (Supplementary Table S4), and their pattern is similar to Middle Late Bronze Age Armenia. Then, we added Armenian populations in Greece_Set9 and Greece_Set10 (Supplementary Table S4). In Greece_Set9 (Supplementary Table S4), we did not see a significant Armenia Early Bronze Age component except for a weak signal in a few individuals (pink peaks in Fig. S6.21). When we added the Middle Late Bronze Age individuals from Armenia in Set9, this weak signal became clearer for those individuals that have a pink proportion (I9006, CGG_2_023840, CGG_2_022372), (Fig. S6.21; Greece_Set10). Overall, steppe sources in Greece are compatible with individuals from the Pontic Steppe, more specifically those derived from Yamnaya.

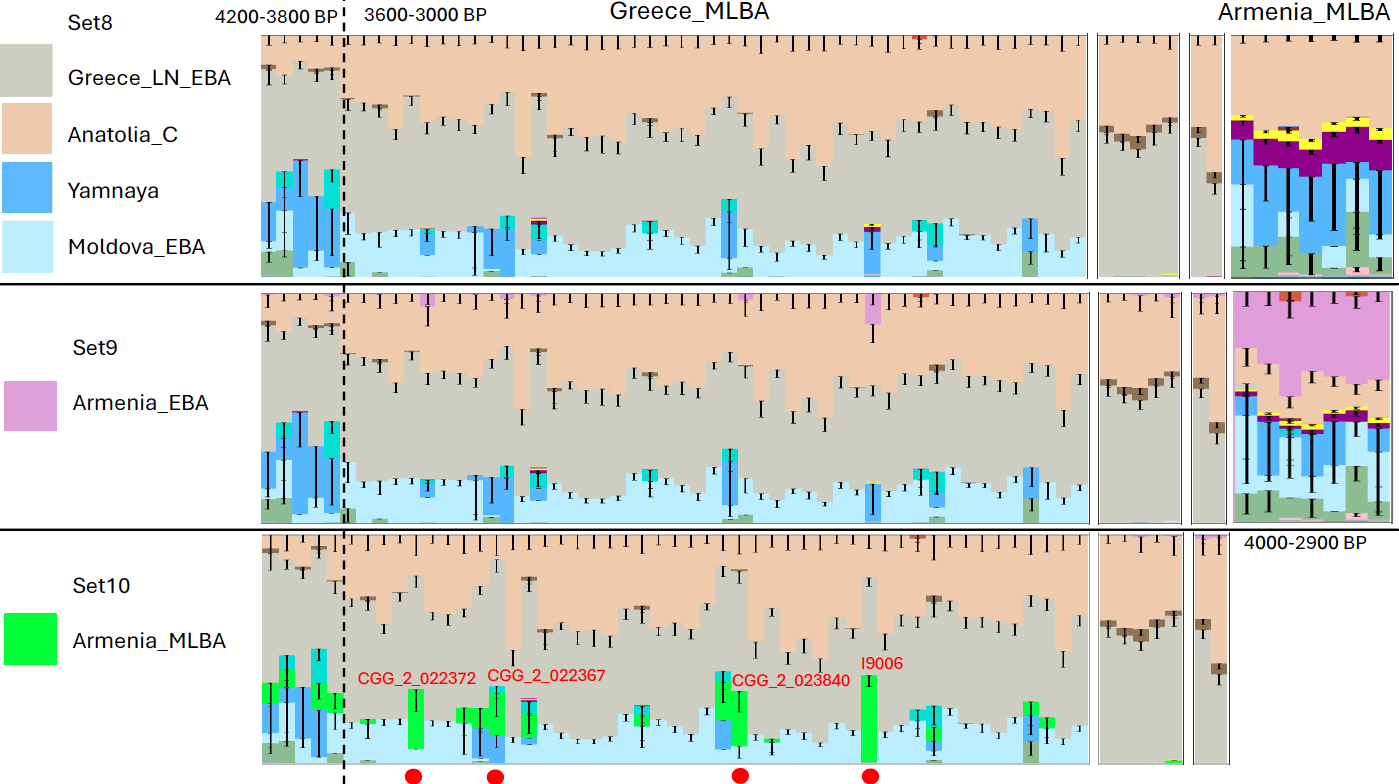

Fig. S6.21. Bar plot showing admixture for each set with source populations from Moldova and Armenia.

#### S6.3 Central Eastern Europe

Hungary and Moldova

To explore the connection of Hungary and Moldova with Greece and Southern Western Europe, we sequenced 22 individuals from Late, Middle, and Early Bronze Age Hungary and six individuals from Early and Middle Bronze Age Moldova. Hungary Bronze Age individuals sequenced in this study show a complex population structure falling within three different clusters which are Aegean/Balkan, Italy Bronze Age, and Bell Beaker. To model individuals from Hungary, Croatia, Montenegro, Serbia, and Romania, we modified our base model by adding a farmer source group from Balkan Neolithic/Chalcolithic individuals (Supplementary Table S4). In the model CE_Set1, some individuals exhibit a CWC proportion, while some have an additional Yamnaya proportion. We also detected Iron Gates and Latvia Mesolithic proportions in some of the Hungarian Bronze Age individuals similar to Croatia (Jag), Serbia, and Bulgaria Bronze Age individuals. When we added BB source (individuals from Poland and Czech Republic) in the model CE_Set2 (Supplementary Table S4), CWC proportion was replaced with BB in most individuals, however two published Hungary BB individuals still show CWC, one (I2787) is from steppe-related (0_4_3) cluster model as a mix of CWC and Yamnaya, the other is from BB cluster (0_4_2) modelled as a mix of BB and CWC. Overall, Balkan Bronze Age presents a similar signature as a mix of Yamnaya and BB components (Fig. S6.22; Fig. S6.23; Supplementary Table S5). This could suggest a connection with Greece which only carries the Yamnaya component or another migration from the Pontic Steppe. The individuals with an elevated Latvia Mesolithic proportion modelled better with Bell Beaker individuals from the Czech Republic without Yamnaya component.

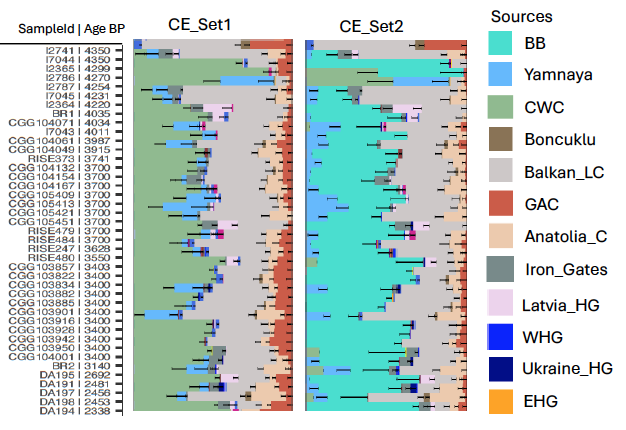

Fig. S6.22. Admixture plots of Hungary Bronze Age individuals obtained from Central Eastern Europe sets (CE_Set1 and CE_Set2).

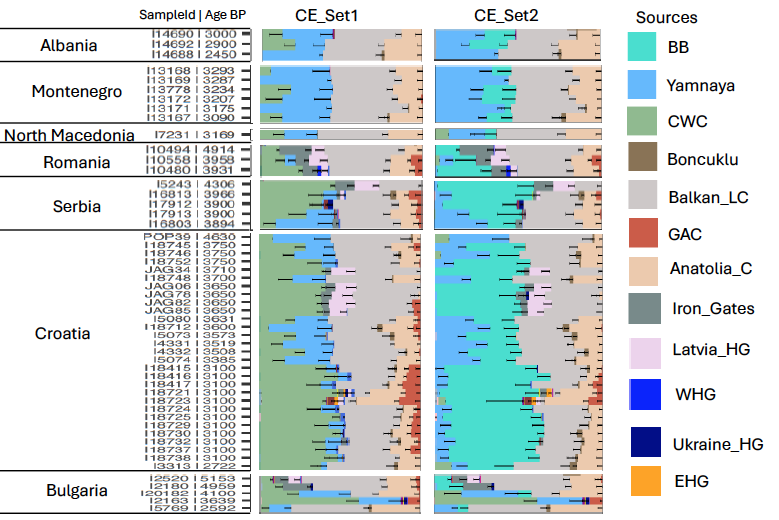

Fig. S6.23. Admixture plots of Balkan Bronze Age individuals obtained from Central Eastern Europe sets (CE_Set1 and CE_Set2).

In individuals from Moldova, we observed two patterns: one group from the Early Bronze Age modelled with only Yamnaya, whereas a group from the Middle Bronze Age modelled with CWC/Yamnaya as in previously published individuals^40^. Since one of the Early Bronze Age individuals (CGG_2_103674) is clustered with Yamnaya cluster which is a source population group, we excluded this individual from the modelling (Fig. S6.24; Supplementary Table S4; S5).

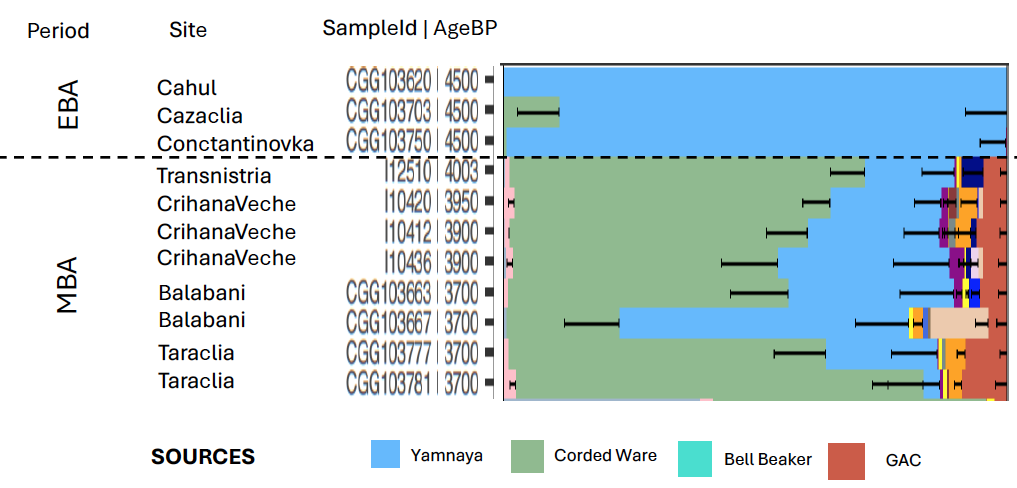

Fig. S6.24. Admixture plot of the Moldovan individuals from Early and Middle Bronze Age (shown as EBA, MBA) obtained from CE_Set2.

#### S6.4 Eastern Mediterranean

Anatolia and Levant

During the Late Neolithic and Chalcolithic, various regions bordering the Greater Caucasus Mountain range saw an increase of Caucasus Hunter-Gatherer (CHG) ancestry. Towards the end of the Neolithic period, the genetic make-up of the Eastern Mediterranean also began to shift towards elevated Caucasus-related gene flow^16,40,61^. It continued into a homogenization process during the Bronze Age, a period characterized by increased cultural and human interaction and expanding trade networks. Here, we report new data from 25 individuals from Central-northern and Western Anatolia. The Early Bronze Age individuals come from Resuloglu Hoyuk, which is associated with the Hattian culture and Kulluoba dated from ~4,300 BP to ~4,900 BP (see Archaeological supplement). We included individuals from the Middle and Late Bronze Age in Central Anatolia from the Kalehoyuk site within the Hittite core region and also added a few Iron Age individuals from Antandros, Kecicayiri, Kulluoba, and Kalehoyuk (Supplementary Table S1, Archaeology Supplementary 2.12). In addition to Anatolia, we included a total of 24 Middle Bronze Age samples from Lebanon, the Canaanite city of Sidon and coastal Syria, the site of Tall Sūkās, and one Sidon individual from the Iron Age (Supplementary Table S1, Archaeology Supplementary 2.8; 2.11). As it is well documented, the genetic make-up of the populations from the Caucasus and Levant within the Kura-Araxes Cultural area is less distinguishable during the Bronze Age^61^. To deeply investigate the populations from the region, we set various models by using eastern source populations, focusing on two geographical areas, the Caucasus and Iran, and subsequently introducing steppe source populations into our models (Table S6.2; Supplementary Table S4).

| East_Set1 | Base + Tepecik + Barçın |
| --- | --- |
| East_Set2 | Base + Tepecik + Barçın + Iran_C |
| East_Set3 | Base + Tepecik + Barçın + Israel_C |
| East_Set4 | Base + Tepecik + Barçın + Armenia_C |
| East_Set5 | Base + Tepecik + Barçın + Caucasus_C |
| East_Set6 | Base + Tepecik + Caucasus_C + Iran_C |
| East_steppe_Set1 | Base + Tepecik + Caucasus_C + Iran_C + Yamnaya + GAC |
| East_steppe_Set2 | Base + Tepecik + Caucasus_C+ Iran_C + Yamnaya + GAC + Piedmont |
| East_steppe_Set3 | Base + Tepecik + Caucasus_C+ Iran_C + Yamnaya + GAC + Piedmont + CWC |

Table S6.2. Source populations of East Mediterranean models.

**East_Set1:** We kept three farmer groups from Boncuklu, Tepecik, and Barçın as sources in our base model before adding eastern sources. The model revealed that all Anatolian Chalcolithic and later period samples are modelled as a mixture of Tepecik and additional eastern components including CHG and, to a lesser extent, Iran Neolithic, along with a small proportion of EHG. The proportion of EHG increases in the Iron Age, similar to the pattern observed in the Levant during this period (Fig. S6.25; S6.26; Supplementary Table S5). Moreover, we detected this small EHG proportion in a Barçın Chalcolithic individual (I1584), although this individual and some of the Western Chalcolithic/Bronze Age individuals exhibit an extra Barçın proportion. An explanation for this inconceivable observation could be that we do not have the right farmer source for these individuals or we are missing an outgroup that shares the EHG proportion. Alternatively, as previously reported, this small EHG proportion, together with CHG in Armenia Chalcolithic (Areni)^38^, could be the potential origin of the EHG proportion in the individual from Barçın. The overall population structure of Anatolia and the Levant Chalcolithic/Bronze Age is very similar, with the exception of the increased Iran Neolithic component in Eastern Anatolia and the Levant compared to Western and Central Anatolia. To investigate the origin of CHG and Iranian components in the Levant and Anatolia, we further modelled Anatolian and Levantine populations with selected source populations from the Caucasus (Armenia_C and Caucasus_C), Iran (Iran_C_Seh Gabi), and the Levant (Israel_Chalcolithic) (Supplementary Table S5). First, we ran each source separately without incorporating any secondary sources, and then we included one or more additional sources to distinguish between them.

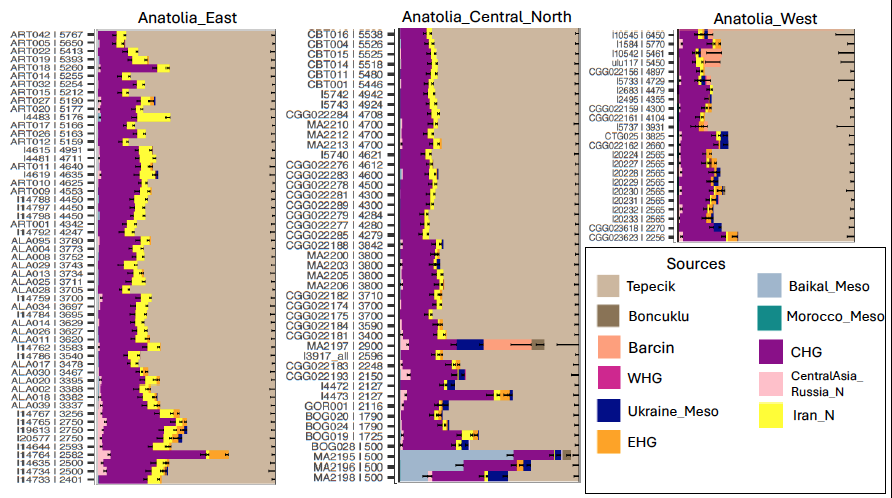

Fig. S6.25. Bar plots generated by using source proportions from East_Set1 in Anatolia presented in three geographical groups (Anatolia-East, Anatolia-Central-North, and Anatolia-West).

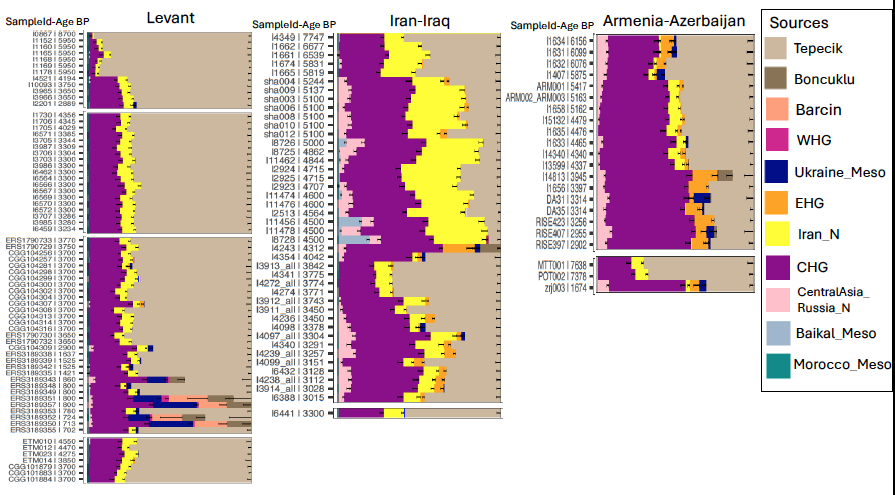

Fig. S6.26. Bar plots generated by using source proportions obtained from model East_Set1 for Levant, Iran-Iraq, and Armenia-Azerbaijan.

**East_Set2:** First, we added Iran Chalcolithic individuals (from Seh Gabi), and observed that the model pulled out the CHG proportion in most of the Levant Chalcolithic/Bronze Age individuals. However, Anatolian individuals required additional or alternative CHG sources except for a few samples from Arslantepe and Alalakh (Fig. S6.27; Supplementary Table S5). The model replicates the previously shown admixture model as a mix of ~30% Iran Chalcolithic and local farmer^28^ for Israel Chalcolithic. Since we do not have Levant Neolithic in our source groups, they were modelled with Tepecik and Iran Chalcolithic. In Middle Bronze Age Israel and Jordan, the Iranian proportion doubled. However, Sidon-Lebanon shows a mixed pattern similar to Arslantepe; while the model fits for some individuals, others still exhibit an additional CHG proportion (Fig. S6.28; Supplementary Table S5).

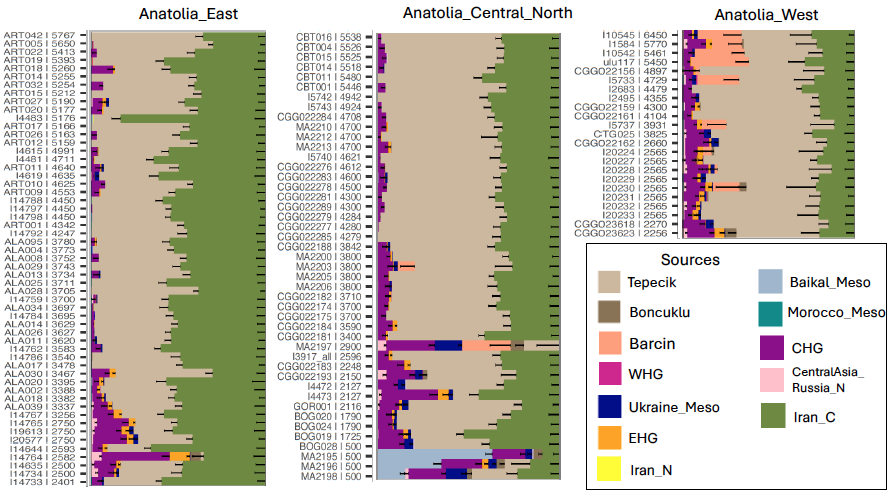

Fig. S6.27. Bar plots generated by source populations of East_Set2 in Anatolia presented in three geographical groups (Anatolia-East, Anatolia-Central-North, and Anatolia-West).

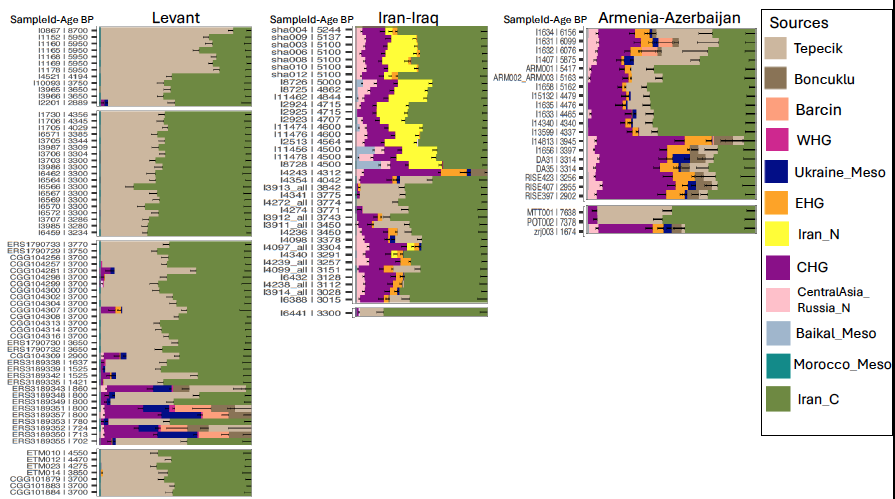

Fig. S6.28. Bar plots generated by using source proportions obtained from model East_Set2 for Levant, Iran-Iraq, and Armenia-Azerbaijan.

**East_Set3:** In this model we replaced the Iran Chalcolithic source with Levant Chalcolithic. The model did not show any differences in the CHG proportion for either Anatolia or Levant Bronze Age individuals (Fig. S6.29; Supplementary Table S5). However, the Levantine proportion completely replaced the farmer proportion in Jordan compared to the rest (Fig. S6.30; Supplementary Table S5). We also observed that this proportion increased in East Anatolia, was lowest in Western Anatolia, and was intermediate in Central Anatolia.

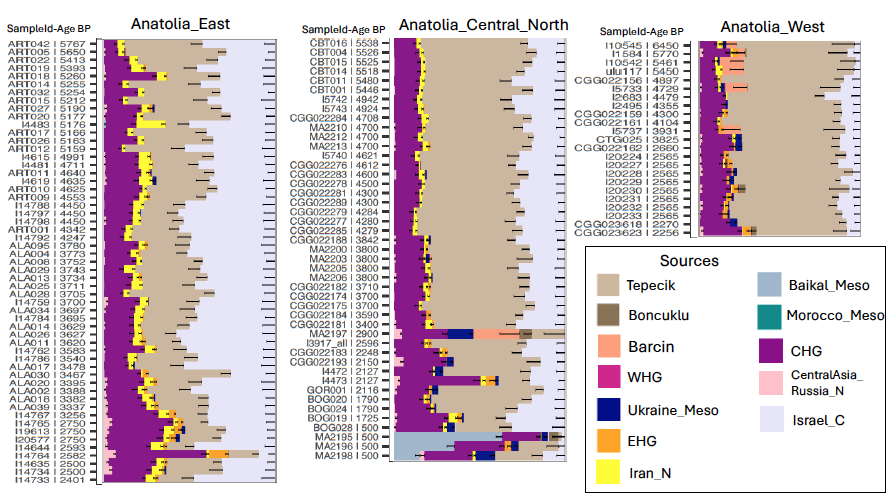

Fig. S6.29. Bar plots generated by source populations of East_Set3 in Anatolia presented in three geographical groups (Anatolia-East, Anatolia-Central-North, and Anatolia-West).

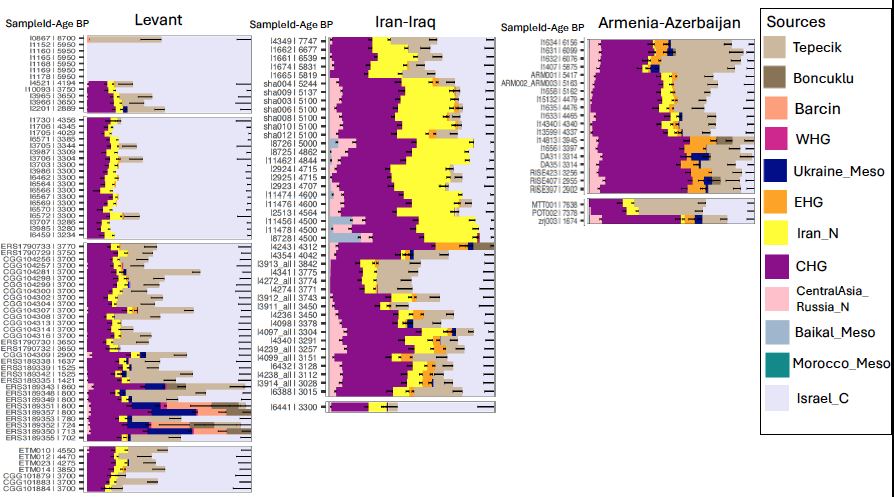

Fig. S6.30. Bar plots generated by using source proportions obtained from model East_Set3 for Levant, Iran-Iraq and Armenia-Azerbaijan.

**East_Set4:** In this model we introduced a Caucasus source by adding Armenia Chalcolithic individuals. The model revealed that while the small proportion of Armenian ancestry was detected in all individuals, it did not replace the CHG proportions in Anatolia and Levant (Fig. S6.31). However, it increases in Armenia Early, Middle and Late Bronze Age individuals (Fig. S6.32).

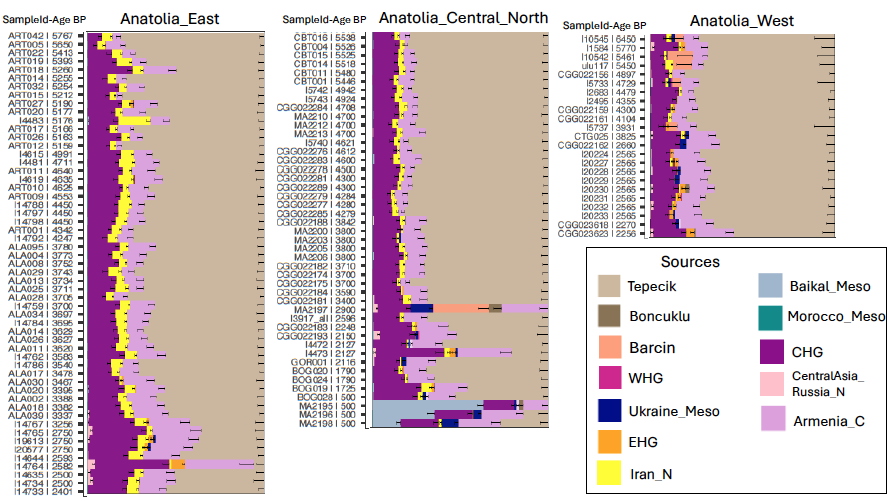

Fig. S6.31. Bar plots generated by using source populations of East_Set4 in Anatolia presented in three geographical groups (Anatolia-East, Anatolia-Central-North, and Anatolia-West).

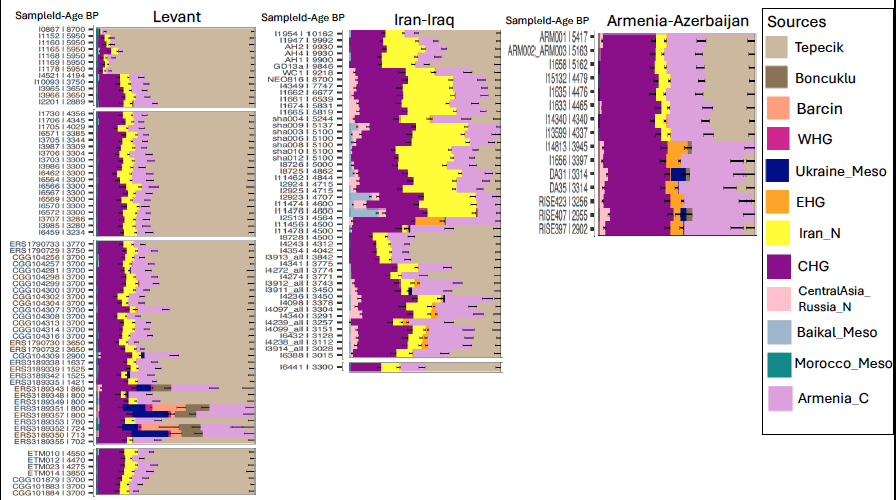

Fig. S6.32. Bar plots generated by using source proportions obtained from model East_Set4 for Levant, Iran-Iraq and Armenia-Azerbaijan.

**East_Set5:** In our clustering, we identified two individuals from the Caucasus within the same cluster. Even though they are labelled with different cultural contexts, given their temporal and spatial proximity, we used them as a single source. We added two individuals (I2056, SA6002) associated with Late Maikop and the Caucasus Eneolithic. This source replaced all CHG proportions in most Anatolian and Levantine individuals, however some still required an additional Iranian component particularly from Eastern Anatolia and the Levant. Interestingly, a small EHG proportion in some Bronze/Chalcolithic individuals from Central and Western Anatolia was also replaced by this source (Fig. S6.33; S6.34). Considering the Caucasus Chalcolithic individuals already carry Iran Chalcolithic ancestry with additional CHG ancestry and a small proportion of EHG, also shared ancestry with steppe-related populations, this small proportion steppe ancestry might be arrived by the southern Caucasus populations.

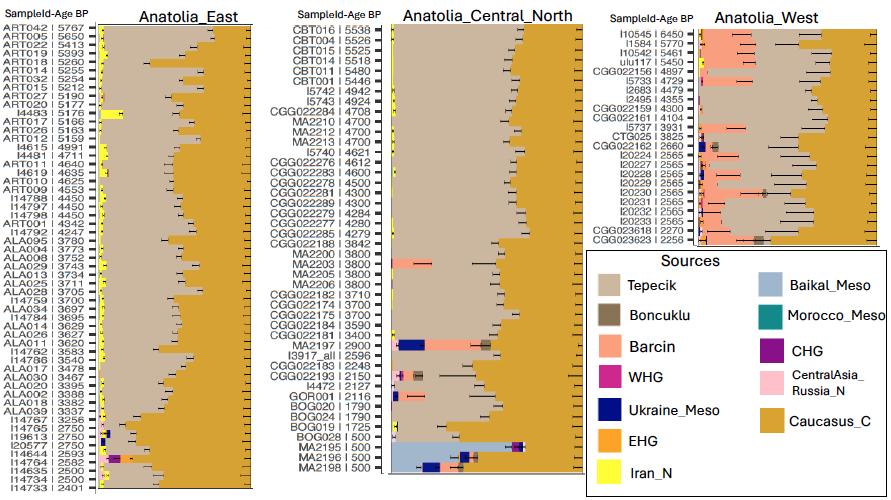

Fig. S6.33. Bar plots generated by using source populations of East_Set5 in Anatolia presented in three geographical groups (Anatolia-East, Anatolia-Central-North, and Anatolia-West).

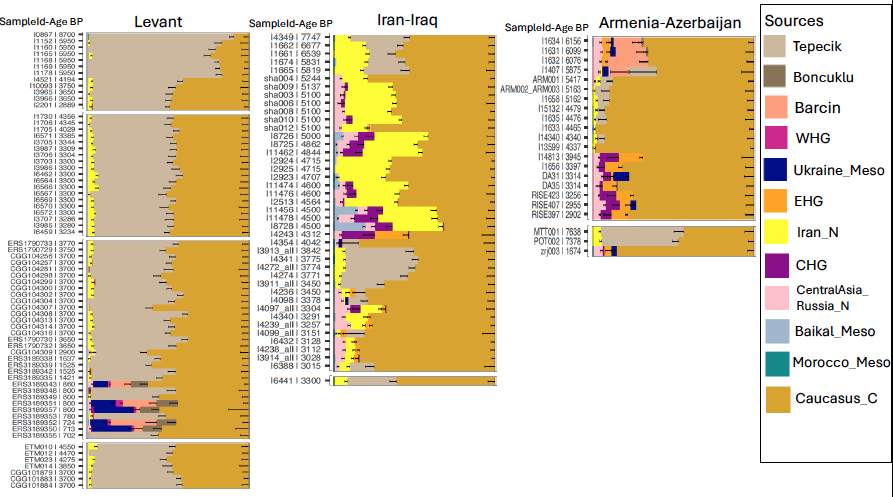

Fig. S6.34. Bar plots generated by using source proportions obtained from model East_Set5 for Levant, Iran-Iraq, and Armenia-Azerbaijan.

**East_Set6**: Following Set2 and Set5, we added both Iran and the Caucasus Chalcolithic groups. By comparing these two sources, we identified three patterns: 1) Individuals from Israel Chalcolithic, Jordan, and a few from Eastern Anatolia are modelled with Iran Chalcolithic and Tepecik; 2) Bronze Age individuals from coastal Levant and most Chalcolithic/Bronze Age individuals from Anatolia show both ancestries in varying proportions; 3) Some individuals from Central and Western Anatolia display only Caucasus ancestry (Fig. S35; Fig. S36; Supplementary Table S5). This Caucasus ancestry increases in Central Anatolia (Kalehoyuk) after ~3800 BP compared to the earlier Central Anatolian individuals. Additionally, some Hittite-associated individuals (MA2200, MA2203, CGG_2_022184) show almost no Iranian ancestry. Moreover, the Western Anatolian individuals appear to carry another farmer source, in this case mix of Barçın and Tepecik, and two individuals from Kaman previously shown to have a small proportion steppe ancestry exhibit this farmer proportion as well (MA2200 and MA2197)^16^. In the east models we only added Anatolian farmers, observing Barçın proportion in some individuals suggesting another farmer population likely from Balkan or Greece as we showed Barçın ancestry widespread in these regions in S6.1.

Since we detected EHG/Ukraine Meso proportions in Anatolia as early as ~5,700 BP in Barçın and in some individuals from Iron Age periods, to clarify the connection between the steppe and Anatolia, we added steppe sources on top of the model East_Set6.

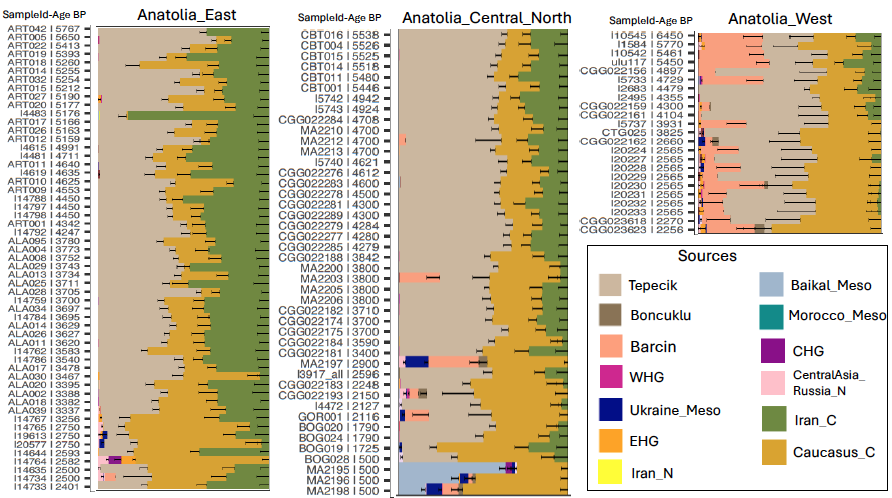

Fig. S6.35. Bar plots generated by using source populations of East_Set6 in Anatolia presented in three geographical groups (Anatolia-East, Anatolia-Central-North, and Anatolia-West).

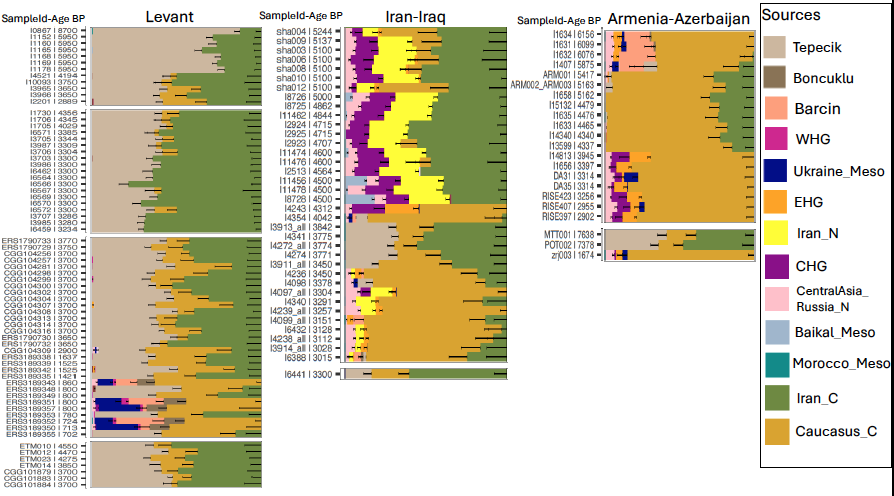

Fig. S6.36. Bar plots generated by using source proportions obtained from model East_Set6 for Levant, Iran-Iraq, and Armenia-Azerbaijan.

**East_steppe_Set1:** We continued adding source populations to East_Set6 (Supplementary Table S4), starting with Yamnaya along with individuals from the Globular Amphora Complex (GAC) who already carried steppe ancestry. The small proportion of EHG/Ukraine_Meso mentioned above is replaced with Yamnaya, though with a high margin of error bar, suggesting it might only be noise or represent another steppe source (Fig. S6.37; Supplementary Table S5). However, this signal becomes clearer in the Iron Age individuals, with increasing Yamnaya proportion.

**East_steppe_Set2:** When we added an early steppe group from Piedmont as a source, the steppe proportion of the Chalcolithic/Bronze Age individuals from Western Anatolia was replaced by the Piedmont source. A small proportion of this ancestry was observed in a newly sequenced individual from Küllüoba (CGG_2_022159). We also detected a mix of Yamnaya and Piedmont components in published Urartian individuals and a newly reported Iron Age Kalehoyuk individual (CGG_2_022193), suggesting other steppe sources (Fig. S6.38).

**East_steppe_Set3:** To distinguish the steppe component in the Bronze Age and later period populations, we added CWC as a representative of a steppe source for European Bronze Age individuals. The Piedmont proportion was replaced by CWC in a Chalcolithic individual from Northwestern Anatolia (I1584) and Central Anatolia Bronze Age individuals mentioned above (Fig. S6.39). However, Piedmont proportion remained in Armenia Chalcolithic and Iran Bronze Age individuals dated before ~4,300 BP (Fig. S6.42). Additionally, we observed a mixed pattern of CWC and Yamnaya in Iron Age individuals from Keçiçayiri (CGG_2_022162), Gordion (GOR001) and Kaman-Kalehoyük (MA2197) similar to the pattern in Balkan Bronze Age individuals (Fig. S6.39).

In summary, we performed the IBD admixture modelling using various eastern sources, as this approach effectively distinguishes geographically distinct populations. To trace the origin of the CHG component in Anatolia and the Levant, we tested sources from the Caucasus and Iran, comparing multiple source populations to evaluate their contributions. In Central Anatolia, individuals from the Chalcolithic and Early Bronze Age show similar ancestry profiles, but this structure changes after ~4,000 BP with an increase in Caucasus ancestry. In Eastern Anatolia, there is significant variation in ancestry, with Arslantepe playing an intercrossing role between Iran and the Caucasus, showing both Iran and Caucasus ancestry and a similarity in structure to coastal Levant Bronze Age. Additionally, population structure was changing in the Iron Age with a peak of Caucasus proportion with steppe ancestry. In contrast, Western Anatolia shows a higher farmer component compared to other regions and with almost no Levantine or Iranian ancestry. The Chalcolithic individuals from Western Anatolia show a distinct mixture of farmer components, such as those from Tepecik and Barçın, differing from the rest of Anatolia. This suggests a potential additional farmer source, possibly from the Balkans or unsampled farmer populations from Anatolia. Previously, we detected a small proportion of CHG and EHG ancestry in one Chalcolithic individual from Ilıpınar (I1584). We also identified Barçın ancestry in one Middle Bronze Age individual from Kalehöyük (MA2203), associated with a Hittite context, as well as in Iron Age individuals, indicating western connections. However, while our eastern source populations from Iran and the Caucasus were unable to fully account for this proportion, steppe sources from CWC populations replaced it. This might suggest either a speculatively early steppe signal or the absence of the correct farmer source causing this analysis noise.

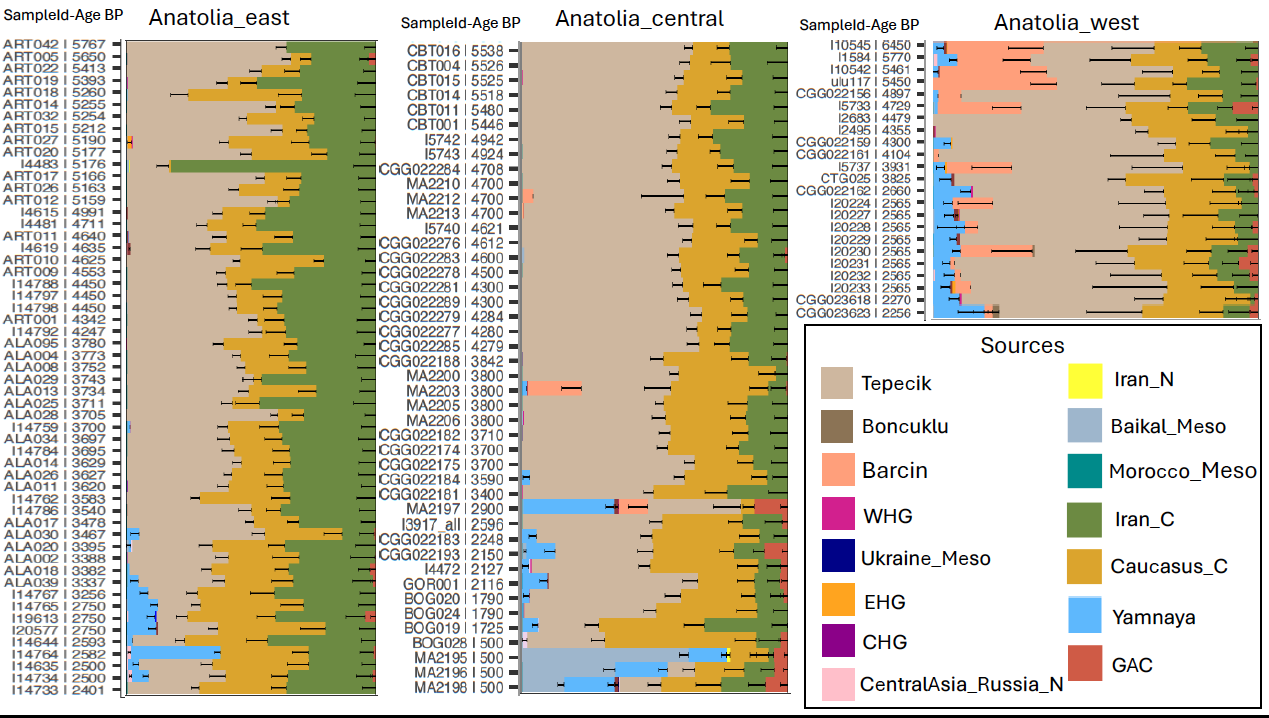

Fig. S6.37. Bar plots generated by using the proportion of source populations of East_

steppe_Set1 in Anatolia.

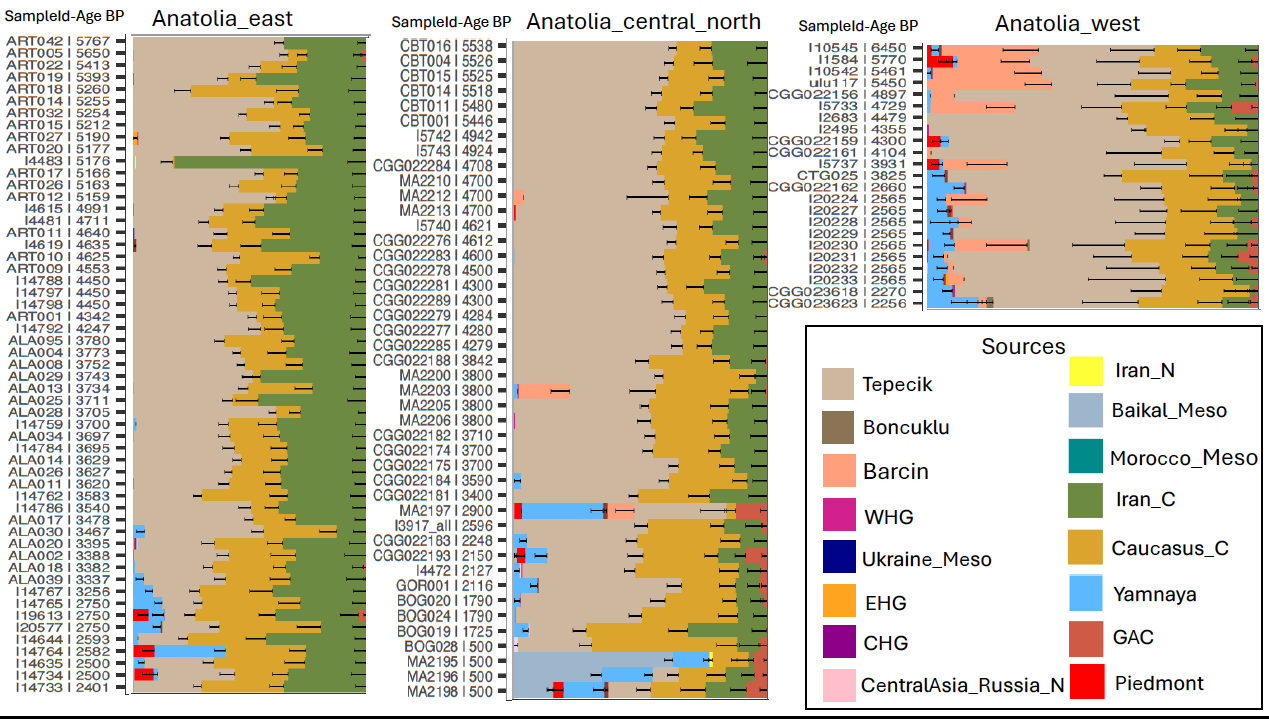

Fig. S6.38. Bar plots generated by using the proportion of source populations of East_

steppe_Set2 in Anatolia.

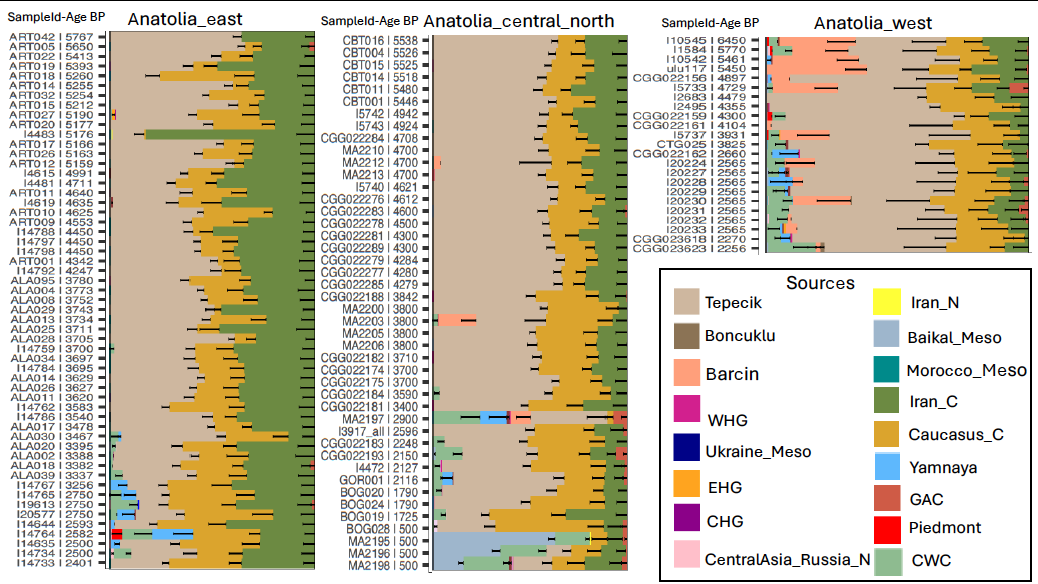

Fig. S6.39. Bar plots generated by using the proportion of source populations of East_

steppe_Set3 in Anatolia.

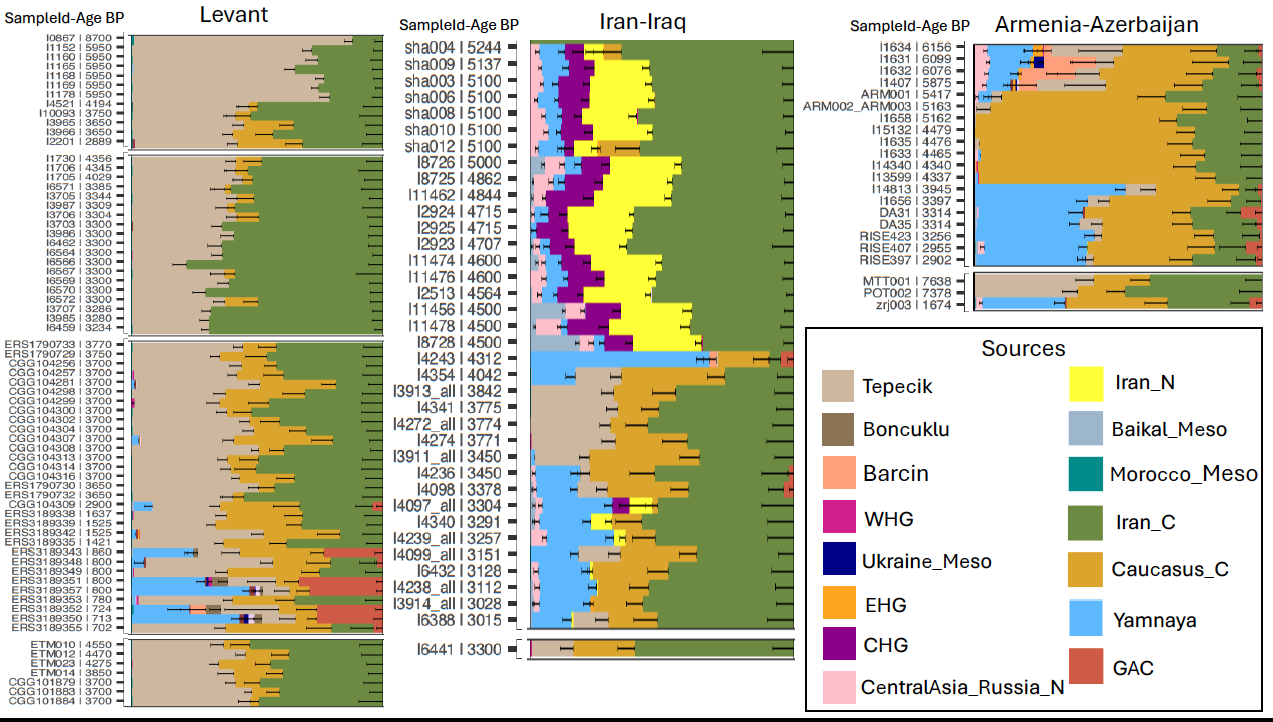

Fig. S6.40. Bar plots generated by using the proportion of source populations of East_

steppe_Set1 in Levant, Iran-Iraq, and Armenia-Azerbaijan.

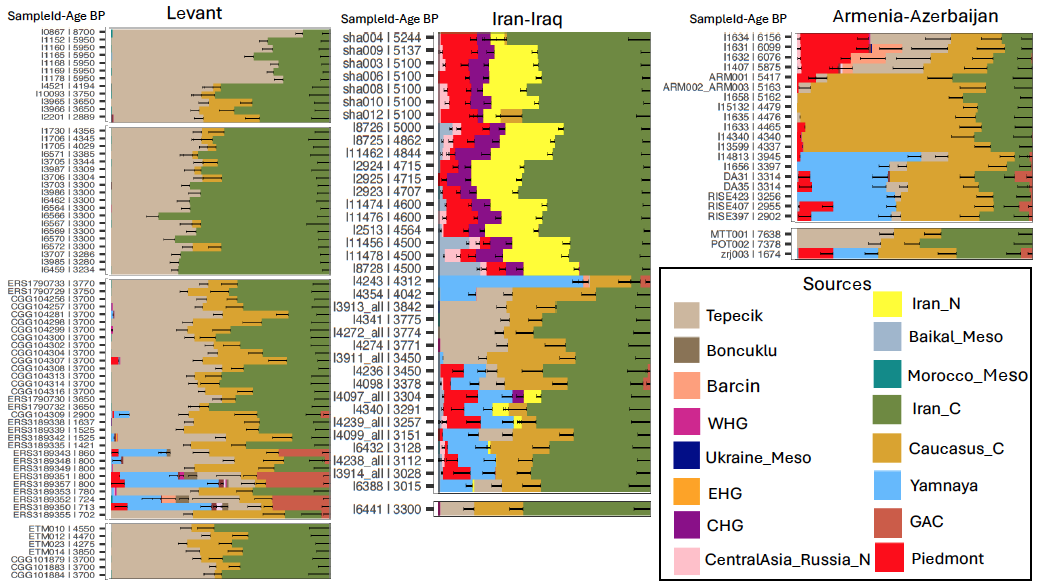

Fig. S6.41. Bar plots generated by using the proportion of source populations of East_

steppe_Set2 in Levant, Iran-Iraq, and Armenia-Azerbaijan.

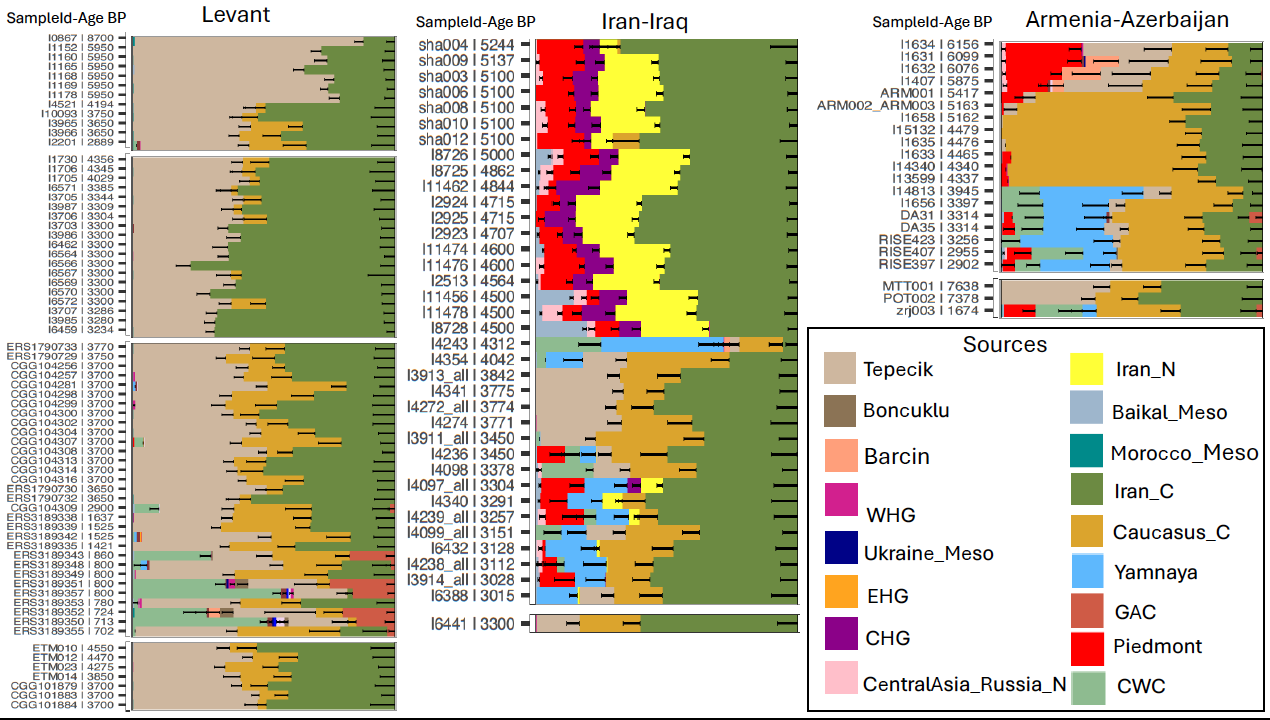

Fig. S6.42. Bar plots generated by using the proportion of source populations of East_

steppe_Set3 in Levant, Iran-Iraq, and Armenia-Azerbaijan.

To look closer at the population structure of Anatolian individuals, we modelled them by using local Chalcolithic and Early Bronze Age populations and three steppe-related populations (BB, CWC, Yamnaya), and replaced the Barçın and Tepecik individuals with Greece Late Neolithic and Early Bronze Age individuals (Supplementary Table S4).

**Anatolia_S1**: In this model, we included Chalcolithic Arslantepe individuals (Eastern Anatolia) that fit for the most eastern populations, and Alalakh individuals from the Northern Levant but not for the other regions of Anatolia (Fig. S6.43;S6.44; Supplementary Table S5).

**Anatolia_S2**: We replaced Arslantepe with Çamlibel Chalcolithic individuals (Central Anatolia). This modelled a few individuals from Arslantepe, one individual from Southwestern Anatolia (Harmanoren), and some newly sequenced individuals from Resuloglu (Fig. S44; Supplementary Table S5).

**Anatolia_S3**: In this model, we added individuals from Central and Western Anatolia Bronze Age, and this replicated the results in Anatolai_S2 (Fig. S44; Supplementary Table S5). However, this source modelled three Resuloglu individuals which did not model with Anatolia_S2. Perhaps Resuloglu received a genetic contribution from both earlier individuals and central-western populations.

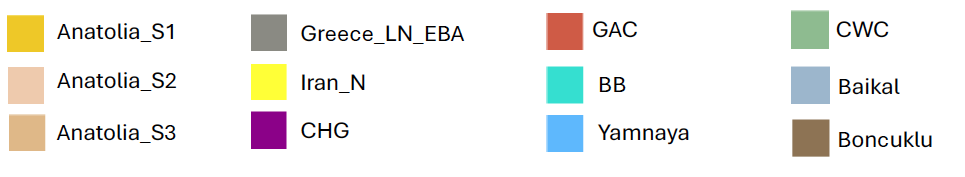

Fig. S6.43. Colours indicate source populations for Anatolian models (S1, S2, S3).

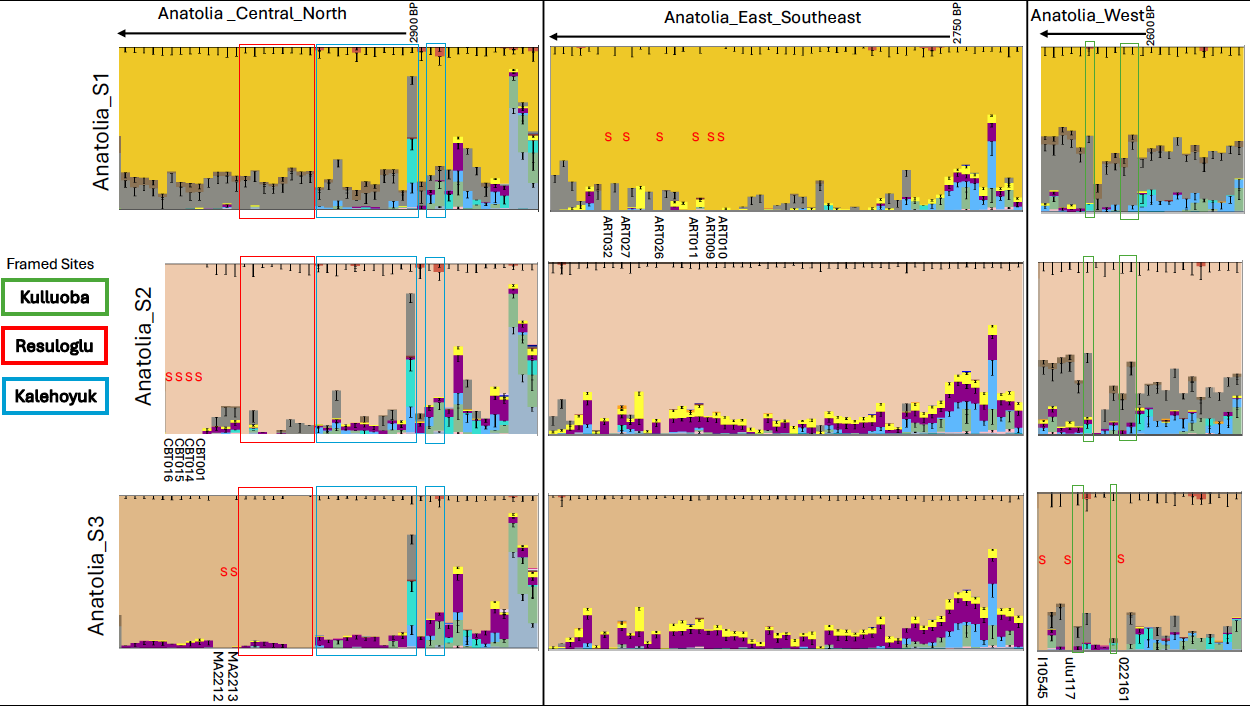

Fig. S6.44. Bar plot generated by modelling Anatolian individuals with local Chalcolithic/Early Bronze Age source populations.

**Cyprus**

To model individuals from Cyprus, we used East_Set6 and Yamnaya (East_steppe_Set1; Supplementary Table S4) and CWC (East_steppe_Set3; Supplementary Table S4). The model with Iran and Caucasus populations revealed that the earliest individual from Cyprus dated to 4,300 BP showed farmer ancestry while the rest of the individuals showed multiple population structures. One group is similar to Anatolia and Levant Bronze Age populations (Tepecik+Caucasus/Iran), while the second group shows the same pattern with Western Anatolian Iron Age individuals (Tepecik+Caucasus+Barçın+Yamnaya), and a few individuals carry a higher proportion of Ukraine Meso/Neolithic, which indicates steppe-related ancestry potentially from Europe (Fig. S6.45). When we added two steppe sources, Yamnaya and CWC respectively, this variation became clearer, showing two steppe outliers (CGG_2_022534; CGG_2_022535) carrying CWC ancestry (Fig. S6.45; Supplementary Table S5). These individuals are also clustered with Scandinavian Bronze Age individuals. Another group shows the Aegean/Balkan signature by showing a mixed pattern of both steppe sources, and the other group resembles Eastern Anatolian/Levantine Bronze Age, showing increased Iran ancestry. Among two outliers with the highest steppe proportion, we cautiously report that one of them (CGG_2_022534) has no reliable context information, and might be from somewhere else. The other one has no direct carbon dating (Archaeology Supplementary 2.2). Interestingly, the Bronze Age individuals from Hala Sultan Tekke show three different admixture patterns; similar to Balkan/Greece, Anatolia Bronze Age, and Lebanon Bronze Age (Fig. S6.42). Further, we investigated Anatolian contribution to Cyprus and Levant Bronze/Iron Age by applying Anatolia Chalcolithic/Bronze sources populations (Fig. S6.46; S6.47). The model with Arslantepe individuals completely modelled Levant populations except the Iron Age and later period individuals who received steppe proportion (Fig. S6.47; Supplementary Table S5). The model with Central and Western Anatolian Chalcolithic/Bronze modelled Cyprus Bronze Age individuals better than Arslantepe and Çamlibel sources (Fig. S6.46; Supplementary Table S5). The individuals with steppe ancestry modelled both with Yamnaya and Bell Beaker. Whereas two Scandinavian outliers are modelled with Bell Beaker, the rest are modelled with Yamnaya, similarly Greece Bronze Age individuals (Fig. S6.46; Supplementary Table S5).

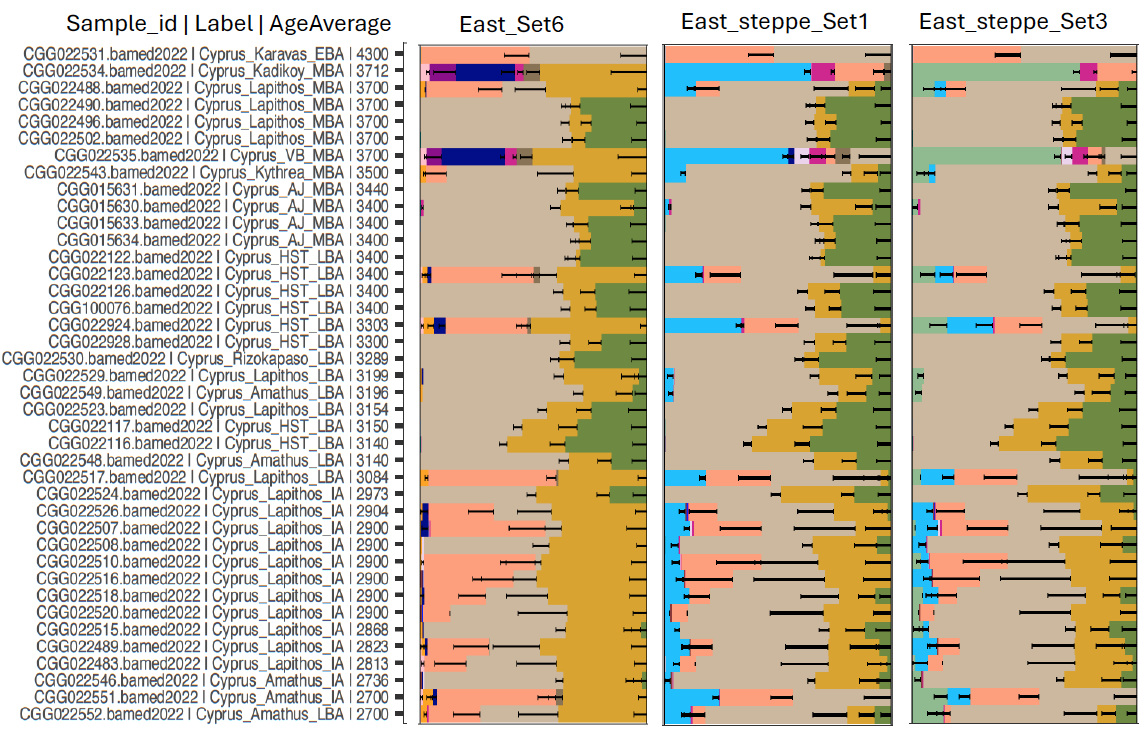
Fig. S6.45. Bar plots generated using eastern and steppe sources to model Cyprus populations.

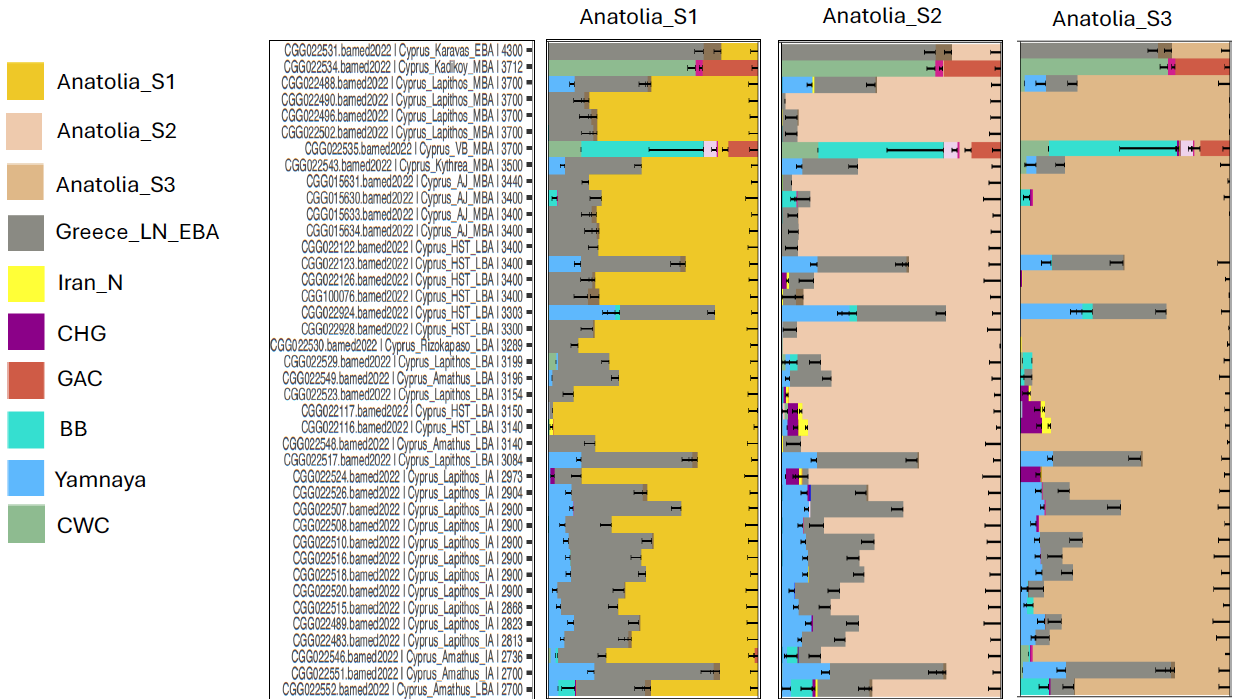

Fig. S6.46. Admixture models for Cyprus populations with Anatolian Chalcolithic/Bronze Age sources.

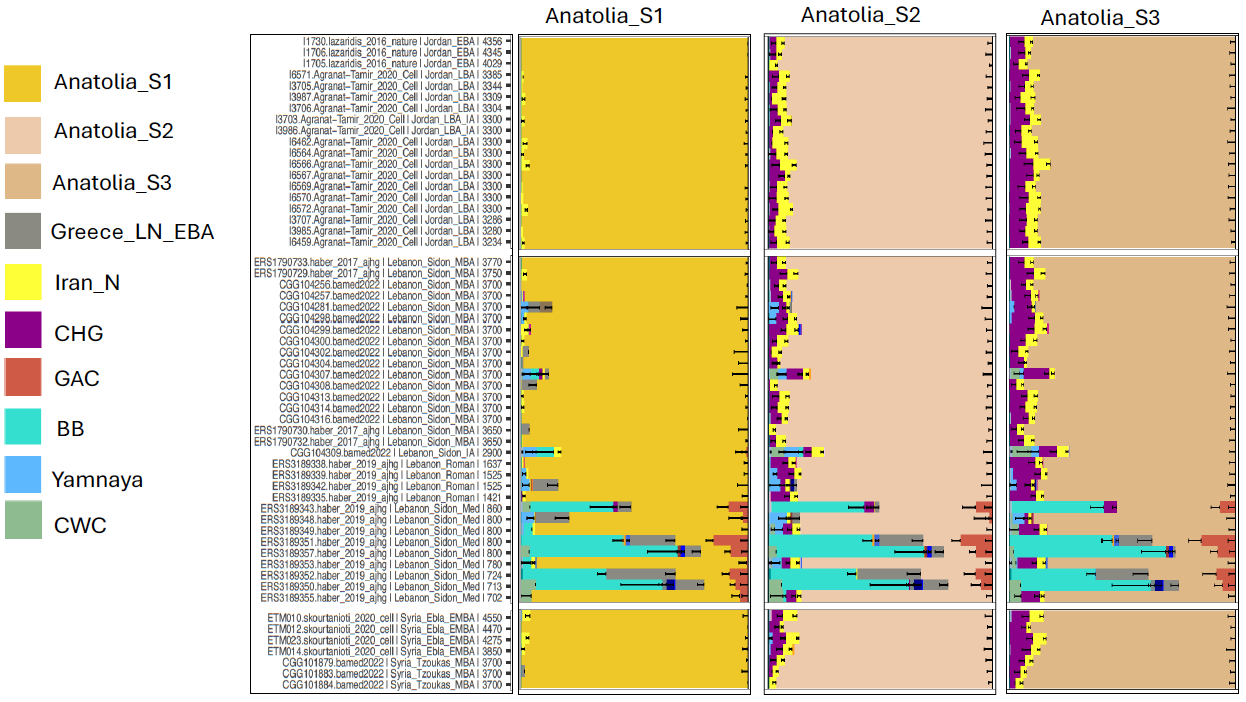

Fig. S6.47. Admixture models for Levant populations with Anatolian Chalcolithic/Bronze Age sources.

### S7. Relatedness analysis

Fulya Eylem Yediay

To identify relatedness, we ran NGSRelate (v2)^93^ on the imputed dataset, calculating allele frequency by using only our samples since we have a dense population structure from Eurasia. We presented here only twin/duplicates, first and second-degree relatives and defined PO (parent-offspring), FS (full-siblings), and HS (half-siblings/grandparent-grandchild/avuncular) by combining rab, R0, R1, KING-robust values from NGSRelate results (Supplementary Table S6). To evaluate those values, we use the following rab value cut-offs; for twin/duplicate 1, 0.75), for the 1st degree [0.75, 0.375), 2nd degree [0.375, 0.1875). To estimate PO and FS, we use R0, R1 and KING values; the range is close to 0, 0.5, and 0.25 for PO, and it is above 0.02, 0.6, and 0.20 for FS, respectively[^94^. We also reported the second-degree relatives as HS using KING range between 0.2 and 0.1, R0 is between 0.14 and 0.3, and R1 is less than 0.5.

Among those first and second-degree pairs of individuals, we found 14 PO, 15 FS and 36 HS relationships. We also identified nine pairs of individuals as twins or duplicates. In the first run of contamination checking, we found CGG_2_022570 to be contaminated, then we split the bam file into two files with separated petrous and tooth data. It turned out that the tooth samples from CGG_2_022570 belonged to individual CGG_022571 (Supplementary Table S6). Therefore, we only used data from the petrous bone and labelled it as CGG_2_022570P.

### S8. Genetic sex determination

Thomaz Pinotti

We estimated the depth of coverage for the individuals with newly generated data using pysam, by counting and measuring the length of the reads (MQ > 30) and dividing the sum by the reference contig length of chromosomes 1–22, X and Y. Because the Y-chromosome presents large regions of repetitive sequence not mappable using short-read sequencing technologies^95–97^, we restricted all analyses to the 10 Mb single-copy region defined in^96^. We called chromosomal sex for all individuals in the dataset by calculating the ratio of the depth of coverage of X to the autosomes, Y to the autosomes, and Y and X chromosomes (Fig. S8.1). We found one individual (CGG_2_104299, DoC: 2.4572) to have a chromosome dosage compatible with a XXY karyotype (Supplementary Table S7). This individual, from Bronze Age Lebanon, represents one of the oldest cases of this karyotype in the archaeogenetics record^4^.

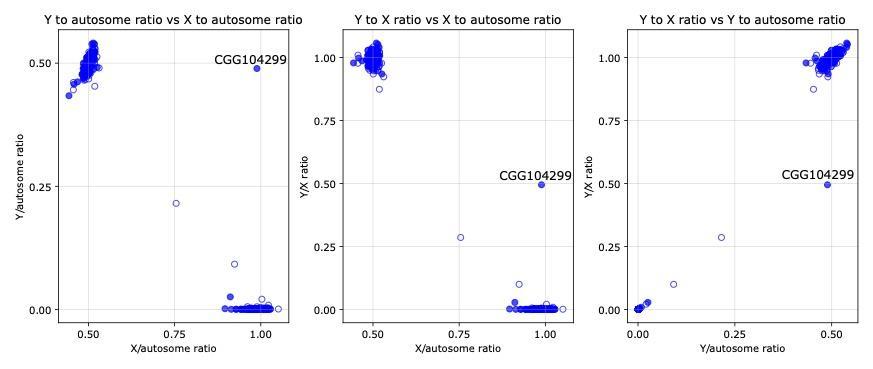

Fig. S8.1. Chromosomal sex of individuals in the dataset. We plotted different ratios of the depth of coverage of autosomes, X and Y chromosomes to identify chromosomal sex among individuals. Individuals clustering on the top of the plot are karyotypically males, while individuals on the bottom are karyotypically female. Unfilled circles represent low-coverage individuals or presenting signals of present-day contamination.

### S9. Uniparental markers

#### S9.1 Y-chromosome analysis

Thomaz Pinotti

We used bcftools^98^ mpileup and call functions to call genotypes within the 10 Mb accessible region of the Y-chromosome^96^. We excluded indels, triallelic positions, and genotypes that were not called in more than 95% of the population of non-clonal reads. To determine haplogroups, we matched ancestral and derived calls to the ISOGG 2019–2020 database using an in-house script that generates haplogroup paths in a root-to-tip manner. Those paths are ranked by the number of supporting variants – while also distinguishing C to T in forward and G to A in reverse strands – and then manually verified. Full results can be found in Supplementary Table S7.

**Discussion**

Haplogroup R is the most common male lineage found in Europe today, and it was initially thought to represent a major European Palaeolithic haplogroup, with its distribution representing patterns of human dispersal after the Last Glacial Maximum (LGM)^99,100^. Strikingly, the ancient DNA record has shown that the two most prevalent subhaplogroups downstream of R, R1a-M417 (R1a1a1) and R1b-M269 (R1b1a1b), are entirely absent in the region until around 5,000 years ago, and their dispersal instead could be linked with the spread of Yamnaya-related autosome ancestry in Western Eurasia during the Bronze Age^9,45^.

Despite this strong correlation, not all reported males from Yamnaya burials carry the dominant lineages found in Europe today. No R1a Yamnaya has been sequenced, and all R1b individuals fall on the R-Z2103 (R1b1a1b1b) branch, while over 90% of R1b males in present-day Europe are in its sister branch, R-L51 (R1b1a1b1a)^101^. This is in contrast with ancient individuals from the Corded Ware and Bell Beaker cultures, which carry matching Y-chromosome lineages to later Europeans, a pattern which may indicate another layer of complexity to the spread of steppe ancestry in the Bronze Age.

**Greece**

Previous works have shown a small pulse of steppe ancestry in Bronze Age Greece among Mycenaeans and in some Late Minoan individuals^14,39,40,92^. However, no clear male lineage contribution from the steppe has been found so far in those populations, owing to both small sample size and the fact that almost all the data have been generated using hybridization technology (“1240K” capture), making high-resolution definition of haplogroups difficult.

We reanalysed the previously published 73 Bronze Age Greece male individuals for their Y-chromosome^14,39,40,92^ – seven from the Early Bronze Age, 38 from Minoan sites in Crete, 28 from the Mycenaean period – and compared them to the haplogroups from the 28 newly generated data from Mycenaean males. We found a relatively high haplogroup overlap between the groups, with most lineages occurring among all populations, with mismatches occurring mostly at the subhaplogroup level. Haplogroups J-P58 (J1a2a1a2) and J-Z2229 (J2a1a1a2) were only present among Minoans, both which seem to have an Anatolian and Levantine association^102,103^, including in our dataset, where it occurs in Turkey, Lebanon, and Cyprus. Mycenaeans, instead, also had a high frequency of J, but most of them fall on the J-Y7011 (J2a1a2b2a~) branch. With rho statistics^104,105^ using the Y-chromosome mutation rate from^19^ (Fu, 2014), we date this lineage to have split 10.97 (95% confidence interval, 9.69–12.45) kya, which, together with the presence of this lineage in an individual from the Neolithic in Croatia^46^, makes it likely farmer-associated.

Crucially, haplogroup R1b-M269 (R1b1a1b) or downstream only appears in individuals from Mycenaean contexts, with a single exception: a Late Minoan individual who is an outlier due to high steppe ancestry^92^. Despite this, only the very rare subhaplogroup R-PF7558 (R1b1a1b2) has been found so far in Bronze Age Greece. This is a sister lineage of dominant R-L51 and the R-Z2103 found in Yamnaya, and is with all likelihood also connected with the Bronze Age spread of steppe ancestry. Among the newly generated individuals, however, we report here the occurrence of four individuals from three different sites in Bronze Age Greece carrying the R-Z2103 lineage. All four of them can be placed in the R-Z2110 (R1b1a1b1b3a1, Z2103 > M12149 > Z2106 > Z2108 > Z2110) branch; similar to previously published Yamnaya individuals, who belong to R-Z2108. Additionally, one Early Bronze Age individual (CGG_2_103620) from Moldova reported in this study whose autosomes can be modelled totally with Yamnaya source also carries the same subhaplogroup (R-Z2108, R1b1a1b1b3a). We date the R-Z2108 lineage to coalesce at 5.70 (95% confidence interval, 5.03–6.49) kya, which is consistent with a dispersion during the Bronze Age.

Haplogroup J-L283 (J2b2a1) is found among Mycenaeans but not Minoans, and its presence in both Bronze Age Caucasus^106^ and a 4.5 kya individual from Moldova, the latter of which can be modelled as 100% from an Yamnaya source, may point to a spread concurrent with the dispersal of steppe ancestry. We date the lineage to be 5.81 (95% confidence interval, 5.13–6.59) kya, which may also support this hypothesis. However, it also occurs among Nuragic age individuals in Sardinia without steppe ancestry^43^. Another ancient DNA study on Sardinia^18^, however, detected the same lineage particularly in an outlier individual, who can be modelled as carrying either extra steppe (Bell Beaker) or Eastern Mediterranean ancestry (Mycenaean or Bronze Age Jordanian). Therefore, currently it is hard to pinpoint its exact route for dispersal.

**Anatolia**

In line with previously published results, we did not find classical steppe uniparental markers in Bronze Age Anatolia, even among Indo-European speakers^16,40,61^. Two individuals from the Bronze Age found in Hittite contexts, however, were I-L699 (I2a1b1a2a2a~), one of which can be confidently placed in I-Y5669 (I2a1b1a2a2a2), the same subhaplogroup as one Yamnaya individual from Kalmykia^8^. One sample from Küllüoba CGG_2_022159 also carries the same haplogroup as published Western Anatolian I5737.

#### S9.2 Mitochondrial DNA analyses

Tharsika Vimala

**Methods**

We re-aligned the newly sequenced ancient DNA reads to the revised Cambridge Reference Sequence (rCRS) for the human mitochondrial DNA sequence using *bwa aln v. 0.7.17**^107^* *(options: -l10000)* and filtered for reads with a mapping quality of minimum 30 using SAMtools v. 1.17^98^*.* We then reconstructed consensus sequences of the mitogenomes with *bcftools**^108^* *v. 1.18* using *mpileup* (*options: --no-BAQ*) to obtain read pileups along the reference sequence, which were then inputted to *bcftools call (options: --multiallelic-caller -- ploidy 1)* for haploid genotype calling. We kept variants covered by at least five reads and a genotype quality above 25. We generated the final consensus sequences with *bcftools consensus.* The reconstructed mitogenomes were aligned with *mafft**^109,110^* *v.7.490,* while we restricted the phylogenetic analysis to the coding region located at 577–16,023 base pairs (rCRS coordinates). We carried out a Maximum Likelihood (ML)-based phylogenetic tree analysis with *RAxML-NG**^111^* *v. 1.2.2* under the substitution model GTR+I+G4 *(options: --all --bs-trees 100).*

**Results**

Haplogroups U and K

We identify individuals from all six main IBD clusters which are given as IBDGlobalCluster (Supplementary Table S3) represented by U and K haplogroups (Fig. S9.1). The most diverse sub-haplogroup is U5, which can further be divided into sub-groups U5a and U5b with their respective sub-clusters. The earliest individuals in sub-group U5b are from two farmer-related (IBD cluster: 0_1_1) individuals from Spain, Valencia 5,000 BP, clustering closely with another farmer-related individual from Italy 3,700 BP. This cluster diverges from a set of mitogenomes carried by individuals from Moldova (n=1) (IBD cluster: 0_4_3 steppe-related) and two Mediterranean individuals from Greece and Syria ~3 ,800–3,700 BP, respectively. The Greek individual carries a mitogenome falling closely to the genetic variation of the early farmers from Spain/Italy. Despite the limited sample size within each of the sub-haplogroups, we find a wide representation within U5a of the main IBD clusters. Most interestingly we find three Mediterranean (IBD: 0_1_2) individuals clustering closely with a Hungarian individual (IBD: 0_1_4 European Bronze and Iron age). Diverging from this cluster we observe two steppe-related individuals from Moldova indicating migrations between Western Europe through Hungary to Greece/Cyprus. A similar pattern is identified for U1 represented by three Mediterranean individuals from 3,700–2,700 BP diverging from a mitogenome carried by a 37,00-year-old Hungarian individual falling within IBD cluster 0_4_2 representing European Bronze Age-related ancestry. Within haplogroup U4, we find an example of an individual from Lebanon 3,700 BP diverging from a steppe-related Moldovan-related individual from 4,500 BP as well as two Italian individuals clustering with a Hungarian and Greek individual, respectively, indicative of the pattern of a West-to-East migration previously mentioned.

When we consider haplogroups U8 and K diverging haplogroup, we mainly find a higher representation of individuals with Mediterranean-related ancestry (IBD 0_1_2). For instance, the sub-haplogroup K1a is dominated by Mediterranean individuals except for a number of Italian (Olmo) genomes from 3,400 BP whose genomic variation falls within the IBD clusters for European Bronze Age/Iron Age (0_1_4). We additionally identify a single ancestrally Western Asian farmer-related individual (IBD 0_1_3) from Cyprus from 4,300 BP in a cluster with another individual from Cyprus from ~3,000 BP falling within IBD clusters for Mediterranean 0_1_2. We have three individuals represented in haplogroup K1b from Spain and Italy >3,300 BP.

Haplogroups W, X, N and I

The haplogroups directly descending from main haplogroup N are mainly carried by individuals with Mediterranean, European Bronze Age, and European Bronze Age/Iron Age ancestry (Fig. S9.2). Within haplogroup W, we observe the three main ancestries, closely clustering according to their genomic ancestries, while a single individual from Cyprus (IBD: 0_1_2) clusters closely with two earlier Bronze Age/Iron Age-related Italian Olmo individuals (IBD: 0_1_4). We find a similar clustering pattern for a sub-cluster of haplogroup C2, while another sub-cluster, haplogroup X2b, is mainly represented by Italian individuals carrying Bronze Age and Iron Age-related IBD sharing. For haplogroup N and I we find more examples of closely related mitogenomes between individuals from Italy (Olmo) and Mediterraneans, respectively (Fig. S9.2).

Haplogroups J and T

Sub-haplogroup J1 is mainly influenced by individuals from Italy Olmo from IBD cluster 0_1_4 (European Bronze Age/Iron Age) and Mediterraneans (Fig. S9.3), while J2 is mainly influenced by Mediterraneans as well but also a few individuals from Spain >4,000 BP from whom we identify mitogenomes of later Italian and Anatolian individuals diverging. Haplogroup T1a is likewise represented by Mediterranean individuals from Greece, Cyprus, and Anatolia, however we also identify a single Hungarian individual belonging to IBD cluster 0_1_2 (Mediterranean) and a Moldovan individual from 3,700 BP assigned to IBD cluster 0_4_1 (Central Asia/Russia). Haplogroup T2 represents the widest range of genomic variation in terms of IBD clusters.

Haplogroups R0, H, V

Haplogroups H and V are represented by the largest number of newly sequenced individuals, while only two individuals, from Anatolia and Greece respectively, carry the ancestral haplogroup R0. Haplogroup HV is mainly represented by Mediterranean individuals in this dataset, while haplogroup V is represented by individuals from Italy, Olmo assigned to the European Bronze Age/Iron Age IBD cluster. We find that the remaining sub-haplogroups within H are influenced by a mix of ancestries.

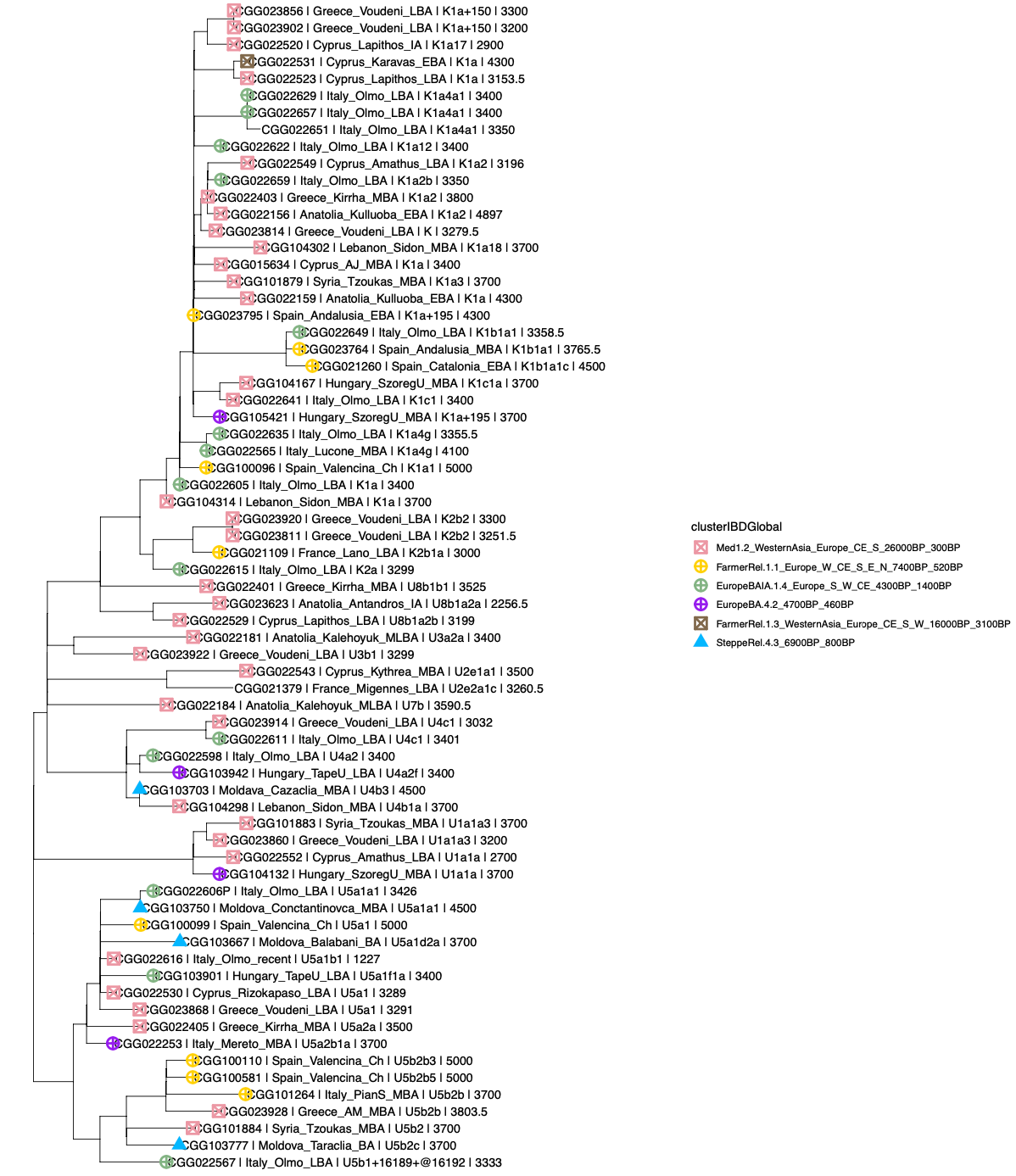

Fig. S9.1. Phylogenetic tree of haplogroups U and K.

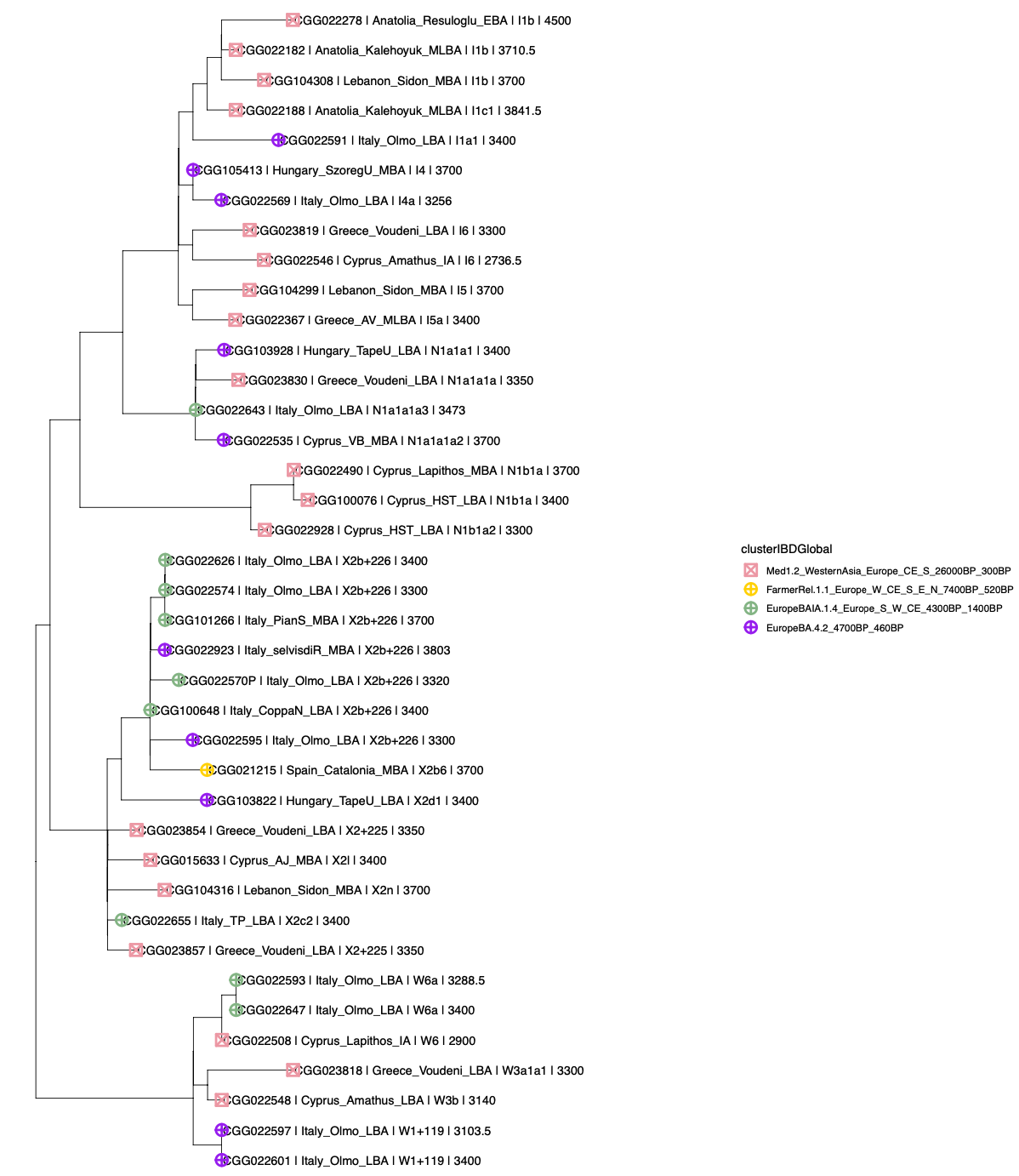

Fig. S9.2. Phylogenetic tree of haplogroups W, X, N, and I.

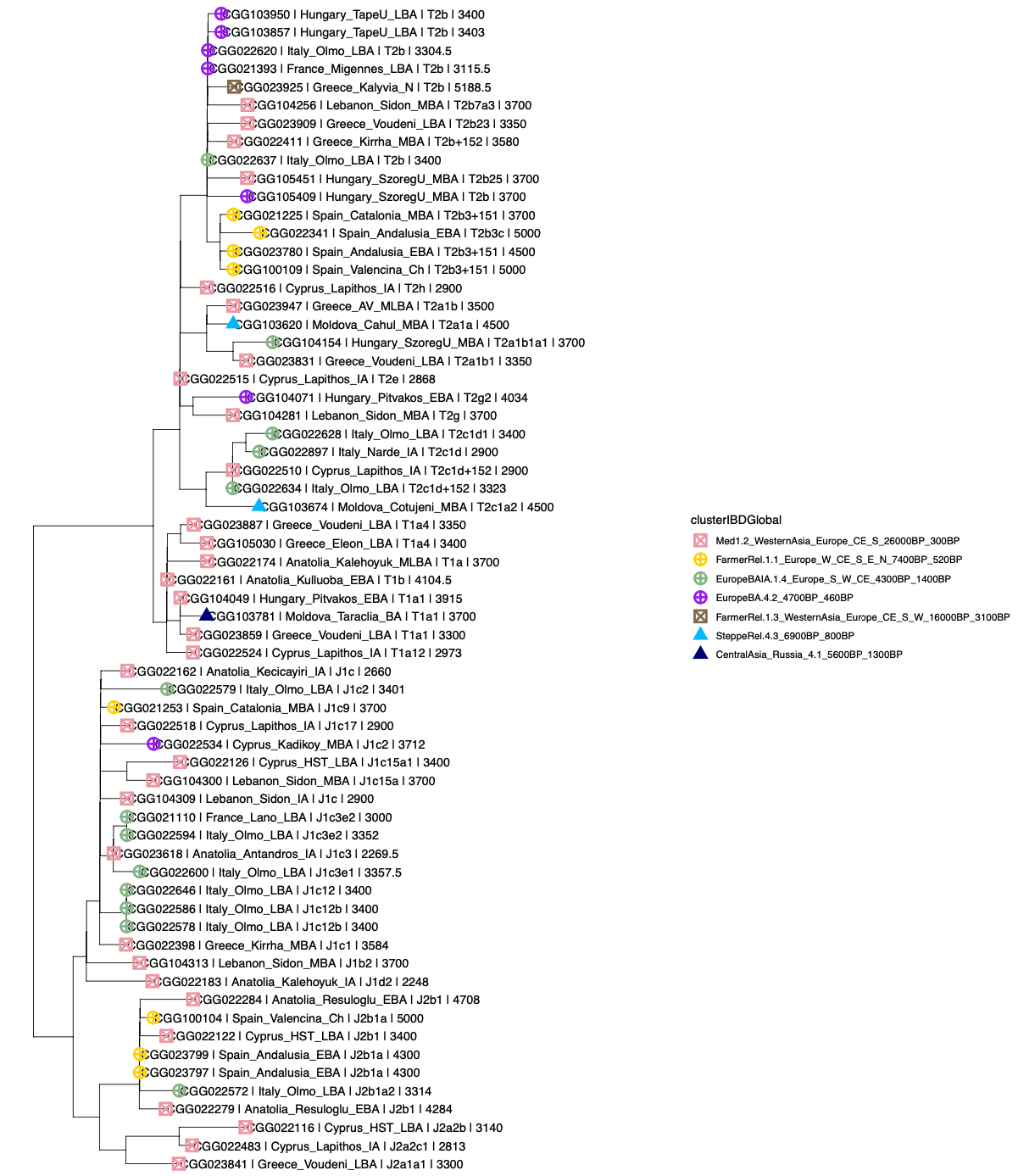

Fig. S9.3. Phylogenetic tree of haplogroups J and T.

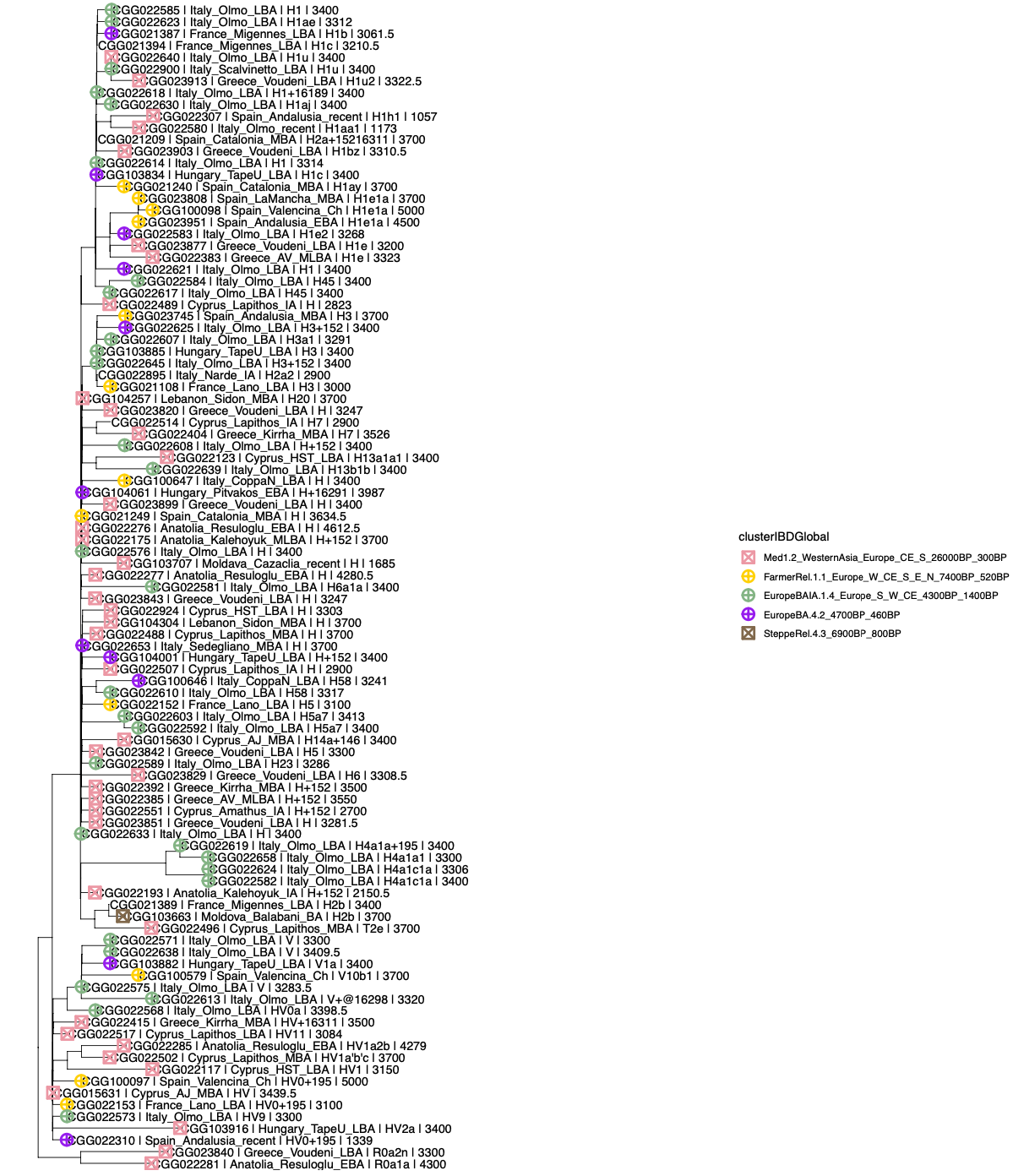

Fig. S9.4. Phylogenetic tree of haplogroups R0, H and V.

### S10. Individual mobility assessment using strontium (Sr) isotopes

Anja Frank, Karin Margarita Frei

Sr isotopes are a useful tool to determine the mobility of ancient individuals on an individual scale. It is possible to determine whether an individual originated from a different place than their burial ground by comparing their skeletal Sr isotope signatures to the bioavailable ⁸⁷Sr/⁸⁶Sr signature at the excavation site and surrounding area. Further, by comparing ⁸⁷Sr/⁸⁶Sr signatures of multiple skeletal parts of the same individual, such as different teeth, or teeth and hair, a timeline of the individual’s movement can be created, as different skeletal parts are (re-)formed at different life stages^112^. However, a meaningful interpretation of ⁸⁷Sr/⁸⁶Sr signatures hinges on the availability of extensive bioavailable ⁸⁷Sr/⁸⁶Sr data of the area of interest, which serves as comparative material for interpreting the human data.

In this study we present skeletal ⁸⁷Sr/⁸⁶Sr of 224 ancient individuals from Cyprus, Greece, Italy, and Spain and compare it with previously published bioavailable ⁸⁷Sr/⁸⁶Sr isotope signatures of these areas to identify and characterize ancient mobility within the Mediterranean during the Bronze Age.

**Methods**

*Sr isotope analysis*

We performed Sr isotope analysis on 232 skeletal samples (139 teeth and 93 petrous bones) from 224 ancient individuals from Cyprus, Greece, Italy, and Spain (Supplementary Table S8). A diamond-tipped dental drill was used to cut a clean enamel sample (1-2 mg) from the tooth samples and to drill 1-2 mg of sample from the densest part of the otic capsule of the petrous bones. The tooth and bone samples were dissolved using a 1:1 solution of 0.5 ml 6M HCl and 0.5 ml 30% H_2_O_2_. Selected samples were spiked with a ^84^Sr-enriched tracer to determine Sr concentration via isotope dilution (ID).

The Sr column separation was done according to the methods of Frei et al^112^, using disposable 1 ml pipette tips fitted with pre-cleaned filters and charged using 200 µl pre-cleaned SrSpec™ resin (50–100 mesh; Eichrome Inc./Tristchem) as disposable extraction columns. The prepared samples were dissolved, loaded onto the columns, and washed using 3M HNO_3_, before the Sr was collected using mq. All Sr concentrations and isotope measurements were performed at the University of Copenhagen using A VG Sector 54 IT mass spectrometer equipped with eight Faraday detectors.

*Sr isotope baseline calculation*

We compiled published bioavailable ⁸⁷Sr/⁸⁶Sr data from archaeological, palaeontological, agricultural, and baseline studies to calculate Sr isotope baselines specific to our investigated excavation sites in Cyprus, Greece, Italy, and Spain (Table S10.1). The data included various proxies for bioavailable Sr, ranging from biomineral data from human or faunal remains^113–115^, over food products^116,117^ to environmental samples^118–120^. Some studies reported several ⁸⁷Sr/⁸⁶Sr values for the same site^118,121^. In such cases, the average ⁸⁷Sr/⁸⁶Sr value was used to avoid biasing the baselines towards an over-represented location.

We based our baselines on modern political boundaries (Table S10.1) for easy understanding. Whether the baselines were calculated at a country, region or province level was based on data availability, as we aimed for at least five baseline values and, if possible, a sampling density of >0.001 sites/km^2^ for a statistically relevant baseline. We calculated two baselines each: 1) as the average bioavailable ⁸⁷Sr/⁸⁶Sr signature of the area ± its single standard deviation (x̅± σ), and 2) as the average bioavailable ⁸⁷Sr/⁸⁶Sr signature of the area ± its double standard deviation (x̅± 2σ).

| **Country** | **Region** | **Province** | **km^2^** | **n** | **n/km^2^** | **x(^87^Sr/^86^Sr)** | **1σ** | **Archaeological sites** | **References** |
| --- | --- | --- | --- | --- | --- | --- | --- | --- | --- |
| Cyprus |  |  | 9,251 | 37 | 0.0040 | 0.70791 | 0.00079 | Ajios Jakovos, Hala Sultan Tekke, Karavas, Kythrea, Lapithos, Rizokarpaso, Vounous-Bellapais | (Hoogewerff et al., 2019; Ladegaard-Pedersen et al., 2020; Voerkelius et al., 2010) |
|  | Limassol |  | 1,396 | 7 | 0.0050 | 0.70816 | 0.00109 | Amathus | (Hoogewerff et al., 2019; Ladegaard-Pedersen et al., 2020; Voerkelius et al., 2010) |
| Greece | Peloponnese | Laconia | 3,636 | 16 | 0.0044 | 0.70968 | 0.00207 | Ayios Vasileios | (Frank et al., 2021b; Hoogewerff et al., 2019; Richards et al., 2008) |
|  |  | Argolis | 2,154 | 17 | 0.0079 | 0.70833 | 0.00041 | Apollo Maleatas | (Frank et al., 2021b; Hoogewerff et al., 2019; Nafplioti, 2008, 2011) |
|  | Central Greece | Phocis | 2,120 | 9 | 0.0042 | 0.70850 | 0.00028 | Kirrha | (Frank et al., 2021a; Hoogewerff et al., 2019) |
|  |  | Boeotia | 3,211 | 6 | 0.0019 | 0.70848 | 0.00033 | Eleon | (Frank et al., 2021a; Prevedorou, 2015; Wang et al., 2019) |
|  | Western Greece | Achaea | 3,272 | 9 | 0.0028 | 0.70829 | 0.00022 | Voudeni | (Frank et al., 2021b; Hoogewerff et al., 2019; Prevedorou, 2015) |
|  |  | Elis | 2,618 | 5 | 0.0019 | 0.70845 | 0.00028 | Kalyvia | (Frank et al., 2021b; Hoogewerff et al., 2019) |
| Italy | Lazio | Rome | 5,363 | 11 | 0.0021 | 0.70967 | 0.00059 | Pian Sultano Crepaccio | (Hoogewerff et al., 2019; Killgrove, 2013; Killgrove and Montgomery, 2016; Lugli et al., 2022; Marchionni et al., 2013; Palombo et al., 2005; Pellegrini et al., 2008; Voerkelius et al., 2010) |
|  | Apulia | Foggia | 7,008 | 31 | 0.0044 | 0.70847 | 0.00021 | Coppa Nevigata | (Hoogewerff et al., 2019; Lugli et al., 2019; Tafuri et al., 2016) |
|  | Basilicata | Potenza | 6,594 | 21 | 0.0032 | 0.70701 | 0.00105 | Toppo Daguzzo | (Frijia et al., 2015; Hoogewerff et al., 2019; Marchionni et al., 2013; Voerkelius et al., 2010) |
|  | Friuli-Venezia Giulia |  | 7,924 | 8 | 0.0010 | 0.70978 | 0.00317 | Mereto, Sedegliano, Sant Osvaldo, Selvis di Remanzacco | (Hoogewerff et al., 2019; Lugli et al., 2022; Petrini et al., 2015; Voerkelius et al., 2010) |
|  | Veneto | Rovigo | 1,819 | 12 | 0.0066 | 0.70920 | 0.00039 | Narde | (Aguzzoni et al., 2020; Cavazzuti et al., 2019; Hoogewerff et al., 2019; Lugli et al., 2022; Marchina et al., 2018) |
|  |  | Verona | 3,121 | 21 | 0.0067 | 0.70883 | 0.00081 | Olmo di Nogara, Scalvinetto | (Cavazzuti et al., 2019; Francisci et al., 2020; Hoogewerff et al., 2019; Ladegaard-Pedersen et al., 2022; Nava et al., 2020) |
|  | Lombardy | Brescia | 4,786 | 15 | 0.0031 | 0.70995 | 0.00336 | Lucone | (Aguzzoni et al., 2020; Cavazzuti et al., 2019; Hoogewerff et al., 2019; Ladegaard-Pedersen et al., 2022; Petrini et al., 2015; Voerkelius et al., 2010) |
| Spain | Andalusia |  | 87,599 | 52 | 0.0006 | 0.71073 | 0.00211 | Castellon Alto, Cerro de la Virgen, Cuesta del Negro, Cueva Artificial la Carada, Los Torcales, Necropolis de los Algarbes, Valencia de la Conception | (Frank et al., 2022; Hoogewerff et al., 2019; Voerkelius et al., 2010) |
|  |  | Almeria | 8,775 | 30 | 0.0034 | 0.71081 | 0.00168 | Argar, Fuente Alamo | (Frank et al., 2022; Hoogewerff et al., 2019) |
|  | Castilla La Mancha |  | 79,463 | 23 | 0.0003 | 0.71104 | 0.00309 | Motilla del Azuer, Terrera del Reloj, La Encantada, | (Chiquet et al., 1999; Díaz-del-Río et al., 2017; Hoogewerff et al., 2019; Voerkelius et al., 2010) |
|  | Catalonia | Barcelona | 7,726 | 12 | 0.0016 | 0.71078 | 0.00208 | Can Martorell | (Valenzuela-Lamas et al., 2018, 2016; Voerkelius et al., 2010) |
|  |  | Lleida | 12,150 | 7 | 0.0006 | 0.70983 | 0.00246 | Minferri, Cantorella | (Hoogewerff et al., 2019; Valenzuela-Lamas et al., 2018; Voerkelius et al., 2010) |

Table S10.1. Overview of baselines calculated for this study. n is the number of used baseline sample points, which were sourced from the listed references, x is the average bioavailable ⁸⁷Sr/⁸⁶Sr composition and 1σ it’s double standard deviation.

**Results**

The skeletal Sr concentrations and ⁸⁷Sr/⁸⁶Sr signatures are given in Supplementary Table S8. The skeletal samples from Cyprus have an average ⁸⁷Sr/⁸⁶Sr signature of 0.7089 ± 0.0023 (σ, n=46) with the lowest value of 0.7079 measured for CGG_2_022116 (third molar) from Hala Sultan Tekke and the highest value of 0.7243 for CGG_2_022535 from Vounous-Bellapais. For the individual CGG_2_022116, three different molars were analysed, which returned a narrow range of 0.7079-0.7080. The average ⁸⁷Sr/⁸⁶Sr signature of the Greek skeletal samples is 0.7085 ± 0.0003 (σ, n=56). The lowest ⁸⁷Sr/⁸⁶Sr signature of 0.7081 was measured for CGG_2_023859 from Voudeni and the highest for CGG_2_023933 from Apollo Maleatas (0.7093). The 95 skeletal samples from Italy have an average ⁸⁷Sr/⁸⁶Sr signature of 0.7095 ± 0.0010 (σ) with a range from 0.7078 for CGG_2_022655 from Toppo Daguzzo to 0.7173 for CGG_2_022620 from Olmo di Nogara. For CGG_2_022606 from Olmo di Nogara and CGG_2_101264 from Pian Sultano Crepaccio two skeletal samples were measured, which ranged from 0.7092–0.7108 and 0.7082–0.7094, respectively. The average ⁸⁷Sr/⁸⁶Sr signature of the Spanish skeletal samples is 0.7089 ± 0.0009 (σ, n=35), with the lowest value of 0.7079 measured for CGG023951 from Cueva Artificial la Carada and the highest of 0.712107 for CGG_2_022341 from Valencina de la Concepcion. For the individuals CGG_2_023764 from Cerro de la Virgen, CGG_2_021225 from Los Torcales, and CGG_2_023793 from Terrera del Reloj, multiple skeletal samples were analysed. While CGG_2_023764 has a similar ⁸⁷Sr/⁸⁶Sr signature for both samples (0.7081), CGG_2_021225 and CGG_2_023793 returned varying ⁸⁷Sr/⁸⁶Sr values between 0.7096 and 0.7100 and 0.7084 and 0.7086, respectively.

The average bioavailable Sr isotope composition and standard deviation for the political regions of the investigated excavation sites are given in Table S10.1. The average ⁸⁷Sr/⁸⁶Sr signature varies between 0.7070 and 0.7110, with standard deviations between 0.0002 and 0.0034. The resulting calculated bioavailable Sr isotope baselines for Cyprus, Greece, Italy, and Spain are shown in Fig. S10.1.

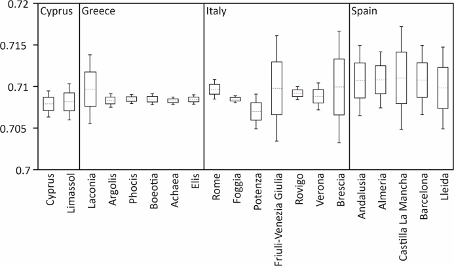

Fig. S10.1. Bioavailable Sr isotope baselines for the investigated areas in Cyprus, Greece, Italy, and Spain. The red dotted lines give the average ⁸⁷Sr/⁸⁶Sr signature of the investigated country, region, or province, the boxes the single standard deviation, and the whiskers the double standard deviation.

**Discussion**

*Sr isotope baselines*

The calculated bioavailable Sr isotope baselines of the investigated regions vary in their average ⁸⁷Sr/⁸⁶Sr composition and range, but are also characterized by significant overlap (Fig. S10.1; Table S10.1). The Sr isotope baseline for all of Cyprus and for Cyprus’s region of Limassol are almost identical, while the investigated political regions of Italy show large differences both in average ⁸⁷Sr/⁸⁶Sr and range of their respective bioavailable isotope baseline. With the exception of Laconia, all of the Greek provinces are characterized by very narrow baselines (σ≤0.0004) close to an average ⁸⁷Sr/⁸⁶Sr signature of 0.07084, while the Spanish baselines are characterized by more radiogenic averages and higher variabilities (σ≥0.0017). The large overlap in bioavailable Sr isotope signatures between many of the investigated areas demonstrates the main limitation of Sr isotopes as a provenancing tool, as mobility between areas with overlapping Sr isotope baselines cannot be identified. For example, all the Greek individuals identified as within the local baseline for Argolis, Phocis, Boeotia, Achaea, and Elis could also come from Laconia, due to the latter’s wider baseline range. Further, these individuals could also come from any of the other investigated countries. Hence, only the mobility of individuals falling outside the local baseline can be unequivocally identified using Sr isotopes alone.

However, identifying non-locals can be difficult in regions characterized by heterogeneous ⁸⁷Sr/⁸⁶Sr signals, such as Friuli-Venezia Giulia and Brescia in Italy or Castilla La Mancha in Spain, as our methods of calculating baselines using the average ⁸⁷Sr/⁸⁶Sr signature and single or double standard deviation resulted in very wide baselines for such regions. For example, in Brescia, the north is characterized by significantly more radiogenic ⁸⁷Sr/⁸⁶Sr values than the south^119^, so that the Sr isotope baseline range based on the double standard deviation covers the ⁸⁷Sr/⁸⁶Sr signatures of most of the other investigated regions. Thus, not only individual CGG_2_022565 from Lucone, Brescia, falls within its local double standard deviation baseline, but also all but two individuals investigated in this study despite the wide variety of localities investigated. To account for this to some extent, our interpretations are mainly based on the less conservative baselines using only the single standard deviation.

Finally, it is important to note that, for some of the investigated regions, very little bioavailable Sr isotope data was available, resulting in baseline sample densities as low as 0.0003 sample baseline points per km^2^. For Spain in particular, ⁸⁷Sr/⁸⁶Sr baseline data is lacking, which is why we had to base our interpretations on poorly covered regional baselines for the majority of the investigated sites. Hence, our findings are preliminary and should be refined using more site-specific baselines once data availability permits it.

*Individual mobility*

Of the 224 investigated individuals/232 skeletal samples, 53 individuals/54 skeletal samples returned ⁸⁷Sr/⁸⁶Sr signatures outside one or both of their respective baselines (Table S10.2), suggesting that these individuals moved at least once within their lifetimes. However, most of the non-local signatures fall only outside their single standard deviation baseline, and some with a minimal difference as low as 0.000003. Due to the uncertainties associated with the ⁸⁷Sr/⁸⁶Sr the baselines, we interpreted our ⁸⁷Sr/⁸⁶Sr results as follows: 1) ⁸⁷Sr/⁸⁶Sr signatures falling less than 0.0001 above or below their single standard deviation baseline are considered potentially non-local; 2) signatures off-set by 0.0001 or more from their single standard deviation baseline, but within their double standard deviation baseline are considered likely non-local; and 3) signatures outside the range of both of their baselines are considered definitely non-local.

With this in mind, we were able to group the 54 ⁸⁷Sr/⁸⁶Sr signatures outside their respective baselines in 12 potentially non-local, 32 likely non-local, and 10 definitely non-local ⁸⁷Sr/⁸⁶Sr signatures. When considering all of these non-local ⁸⁷Sr/⁸⁶Sr signatures, about 24% of the investigated individuals likely moved at least once during their lifetime. As our calculated baselines are based on modern political lines and not site-specific, they are likely biased based on the baseline data availability and distribution (see also discussion above). Hence, we consider only the likely and definitely non-local ⁸⁷Sr/⁸⁶Sr signatures reliable, which suggests that mobility was slightly lower at ~18%. However, mobility was not evenly distributed across the investigated countries. The percentage of mobile individuals for Cyprus and Greece is rather low with only ~7% and ~11% mobile individuals, respectively. On the other hand, about a quarter of the investigated individuals from both Italy and Spain were likely mobile.

The non-local ⁸⁷Sr/⁸⁶Sr signatures from Greece (two definitely and four likely non-local) and Cyprus (one definitely and two likely non-local) all fall above their respective baselines suggesting that these individuals spent time in an area with a more radiogenic baseline. However, while the Greek individuals could have moved from other parts of Greece, such as the Southern Peloponnese or northern mainland^122^, the definitely non-local individual CGG022535 from Vounous-Bellapais falls far outside the typical Sr isotope range of Cyprus^119,121^. The ⁸⁷Sr/⁸⁶Sr signature of CGG_2_022535 is even more radiogenic than the bioavailable Sr isotope data reported for well-known trading partners of Cyprus, such as Greece or Anatolia^119,122–125^. Within Europe, such radiogenic ⁸⁷Sr/⁸⁶Sr values have mainly been reported for Northern Scandinavia ^119^, which fits in well with the aDNA findings of this study [FA1].

Of the non-local signatures identified for Italy, the ones from Pian Sultano Crepaccio (CGG_2_101264 and CGG_2_101266) fall below their baselines, while the remaining non-locals are all more radiogenic than their respective baselines. However, just like in Greece, all of the non-local individuals could have moved within Italy, due to its wide range in bioavailable ⁸⁷Sr/⁸⁶Sr^126^. For the women CGG_2_022606 from Olmo di Nogara and CGG_2_101264 from Pian Sultano Crepaccio, a petrous and a tooth sample were investigated, and both returned one definitely non-local ⁸⁷Sr/⁸⁶Sr signature and one signature within their respective baselines, enabling us to better identify their origin and movement. The petrous ⁸⁷Sr/⁸⁶Sr signature of CGG_2_101264 from Pian Sultano Crepaccio falls outside the baselines of Rome, clearly indicating that this woman did not originate from Pian Sultano Crepaccio, as the petrous already starts forming in utero. The ⁸⁷Sr/⁸⁶Sr signature of her tooth, however, falls within Rome’s baselines, suggesting that she moved to Pian Sultano Crepaccio during her early childhood/adolescence, when the tooth enamel mineralizes. For CGG_2_022606 from Olmo di Nogara, the petrous ⁸⁷Sr/⁸⁶Sr signature falls within the baselines of Verona, indicating that this woman likely originated there, potentially even in Olmo di Nogara itself. Her tooth, on the other hand, falls above Verona’s baselines, suggesting that the individual moved outside of Verona during the molar’s formation in early childhood, before eventually returning to Olmo di Nogara where she was buried.

The non-local ⁸⁷Sr/⁸⁶Sr signatures from Spain all fall below their respective baselines, suggesting that they spent time in an area with less radiogenic ⁸⁷Sr/⁸⁶Sr signatures. Such signatures have also been reported within Spain^118,119^, suggesting that the individuals might have only travelled within Spain. For the likely non-local individual CGG_2_023764 from Cerro de la Virgen, two molar samples were analysed, which returned similar ⁸⁷Sr/⁸⁶Sr signatures. As both fall outside the single standard deviation of the (very preliminary) baseline of Andalusia, no movement during her childhood could be detected. Instead, the woman might have moved to Cerro de la Virgen, where she was buried, during her adolescence or later. However, considering the poor baseline coverage of Andalusia, this finding should be verified with additional baseline data at a later time.

In summary, this study shows that a significant percentage of ancient individuals were mobile within the Mediterranean region during the Bronze Age. However, most of the movement could have been in-country and over short distances. Individual CGG_2_022535 from Cyprus, however, reveals potential travel from as far away as Scandinavia, showing the inter-connectivity of Europe during that time. However, our study also revealed a lack of suitable bioavailable ⁸⁷Sr/⁸⁶Sr baseline data. For a more robust interpretation of our skeletal ⁸⁷Sr/⁸⁶Sr, more extensive and comprehensive baseline studies of the investigated areas are needed.

| **Country** | **Sample ID** | **Sex** | **Sample type** | **^87^Sr/^86^Sr** | **2SE (ppm)** | **Δ_1σ_** | **Δ_2σ_** |
| --- | --- | --- | --- | --- | --- | --- | --- |
| Cyprus | CGG022543 | XY | Petrous | 0.70871 | 9 | 0.000011 |  |
|  | CGG022524 | XY | Molar | 0.70890 | 9 | 0.000199 |  |
|  | CGG022526 | XY | Molar | 0.70871 | 13 | 0.000006 |  |
|  | CGG022504 | XY | Tooth | 0.70891 | 9 | 0.000212 |  |
|  | CGG022535 | XY | Tooth | 0.72431 | 7 | 0.015610 | 0.014824 |
| Greece | CGG023933 | XX | Molar 1 (upper) | 0.70935 | 15 | 0.000607 | 0.000200 |
|  | CGG105030 | XY | Incisor (upper) | 0.70882 | 15 | 0.000010 |  |
|  | CGG022400 | XY | Molar 1 (upper) | 0.70907 | 16 | 0.000289 | 0.000012 |
|  | CGG022404 | XY | Canine (lower) | 0.70883 | 18 | 0.000046 |  |
|  | CGG023835 | XX | Incisor 1 (upper) | 0.70865 | 15 | 0.000126 |  |
|  | CGG023846 | XX | Molar 1 (lower) | 0.70852 | 11 | 0.000003 |  |
|  | CGG023851 | XX | Canine (lower) | 0.70866 | 14 | 0.000139 |  |
|  | CGG023884 | XX | Molar 1 (lower) | 0.70864 | 12 | 0.000121 |  |
|  | CGG023913 | XY | Molar 1 (lower) | 0.70871 | 17 | 0.000188 |  |
| Italy | CGG100646 | XX | Petrous | 0.70906 | 16 | 0.000375 | 0.000168 |
|  | CGG022567 | XX | Molar | 0.70971 | 11 | 0.000075 |  |
|  | CGG022568 | XX | Molar | 0.71002 | 11 | 0.000389 |  |
|  | CGG022573 | XX | Molar | 0.71047 | 7 | 0.000837 | 0.000031 |
|  | CGG022574 | XY | Molar | 0.71014 | 14 | 0.000505 |  |
|  | CGG022575 | XX | Molar | 0.70991 | 7 | 0.000278 |  |
|  | CGG022578 | XX | Molar | 0.70973 | 9 | 0.000100 |  |
|  | CGG022579 | XX | Petrous | 0.70969 | 24 | 0.000054 |  |
|  | CGG022582 | XY | Petrous | 0.70964 | 12 | 0.000007 |  |
|  | CGG022586 | XX | Molar | 0.70967 | 11 | 0.000032 |  |
|  | CGG022591 | XX | Petrous | 0.70968 | 28 | 0.000049 |  |
|  | CGG022602 | XX | Molar | 0.71282 | 12 | 0.003186 | 3.000000 |
|  | CGG022603 | XY | Tooth | 0.70996 | 12 | 0.000323 |  |
|  | CGG022606 | XX | Tooth | 0.71081 | 17 | 0.001174 | 0.000368 |
|  | CGG022606 | XX | Petrous | 0.70921 | 23 |  |  |
|  | CGG022619 | XX | Petrous | 0.70995 | 8 | 0.000317 |  |
|  | CGG022620 | XY | Molar | 0.71729 | 15 | 0.007659 | 0.006853 |
|  | CGG022623 | XY | Molar | 0.71019 | 12 | 0.000557 |  |
|  | CGG022630 | XX | Molar | 0.70987 | 18 | 0.000238 |  |
|  | CGG022635 | XY | Molar | 0.71012 | 13 | 0.000490 |  |
|  | CGG022636 | XX | Molar | 0.70974 | 14 | 0.000106 |  |
|  | CGG022639 | XY | Tooth | 0.70964 | 10 | 0.000008 |  |
|  | CGG022646 | XY | Molar | 0.70985 | 12 | 0.000222 |  |
|  | CGG022647 | XY | Molar | 0.71015 | 14 | 0.000513 |  |
|  | CGG022648 | XY | Molar | 0.70983 | 14 | 0.000193 |  |
|  | CGG022657 | XY | Molar | 0.70986 | 17 | 0.000225 |  |
|  | CGG101264 | XX | Petrous | 0.70818 | 15 | -0.000902 | -0.000315 |
|  | CGG101264 | XX | Tooth | 0.70936 | 14 |  |  |
|  | CGG101266 | XX | Petrous | 0.70809 | 16 | -0.000987 | -0.000400 |
|  | CGG022898 | XX | Petrous | 0.71003 | 14 | 0.000401 |  |
|  | CGG022899 | XX | Petrous | 0.70993 | 54 | 0.000296 |  |
|  | CGG022900 | XX | Petrous | 0.70993 | 25 | 0.000299 |  |
| Spain | CGG022310 | XY | Tooth | 0.70855 | 30 | -0.000575 |  |
|  | CGG023795 | XY | Molar 2 (lower) | 0.70809 | 20 | -0.000532 |  |
|  | CGG023797 | XY | Molar 3 (lower) | 0.70794 | 20 | -0.000690 |  |
|  | CGG023799 | XY | Molar 3 (upper) | 0.70802 | 23 | -0.000601 |  |
|  | CGG023764 | XX | Tooth | 0.70808 | 15 | -0.000543 |  |
|  | CGG023764 | XX | Tooth | 0.70806 | 24 | -0.000569 |  |
|  | CGG023951 | XY | Molar 3 (upper) | 0.70793 | 12 | -0.000694 |  |
|  | CGG022307 | XX | Tooth | 0.70828 | 13 | -0.000845 |  |
|  | CGG100104 | XX | Molar | 0.70851 | 35 | -0.000114 |  |
|  | CGG023780 | XY | Petrous | 0.70860 | 19 | -0.000022 |  |

Table S10.2. Individuals who have a skeletal ⁸⁷Sr/⁸⁶Sr signature outside their respective baselines. Δ is the difference between the skeletal ⁸⁷Sr/⁸⁶Sr signature and upper (positive values) or lower (negative values) baseline range using the singe (1σ) or double standard deviation (2σ). Samples marked in green are multiple samples from one individual.
