## Supplementary material for "Ancient genomics support deep divergence between Eastern and Western Mediterranean Indo-European languages": Linguistic Supplementary

Linguistic Supplementary Information

**Table of contents**

**1. The Indo-European languages: Introduction	1**

**2. Languages of Italy	1**

**2.1 Italic	1**

**2.2 Etruscan	2**

**2.3 Messapic	3**

**3. Languages of Iberia	3**

**3.1 Celtiberian	3**

**3.2 Lusitanian	4**

**4. Languages of the Eastern Mediterranean and the Caucasus	4**

**4.1 Greek	4**

**4.2 Phrygian	5**

**4.3 Armenian	6**

**4.4 Anatolian	6**

**5. Conclusion	8**

**References	9**

### 1. The Indo-European languages: Introduction

The Indo-European language family is among the largest in the world and is spoken by ca. 44% of the global population^1^. It derives from a prehistoric and extinct language, Proto-Indo-European, which diverged into a multitude of subgroups that dispersed across Eurasia in prehistoric times^2–5^. At their earliest attestations, the branches Italic, Celtic, Germanic, Balto-Slavic, Albanian, Greek, Anatolian, Armenian, Indo-Iranian, and Tocharian already covered a large area across Eurasia, stretching from Atlantic Europe to Xinjiang West China. Not least because of its far-reaching impact, the origin of the Indo-European languages has traditionally been and remains one of the prime questions on West Eurasian prehistory^6^. Here we highlight the Indo-European subgroups of the wider Mediterranean region, including Italic, Messapic, Celtiberian, Lusitanian, Greek, Phrygian, Armenian, and Anatolian. We conclude with a discussion of the question on the subgrouping of the Indo-European languages.

### 2. Languages of Italy

#### 2.1 Italic

The Italic family of Indo-European languages survives today exclusively in the form of the Romance languages, all having descended from Latin. But the Italic family originally comprised a second branch apart from Latin (or Latino-Faliscan), namely the Sabellic (or Osco-Umbrian) languages. The Sabellic languages included South Picene, Oscan, Umbrian, and some smaller languages usually classified as dialects of the latter two. Together, the Italic branch is attested in inscriptions from the 7th century BCE onwards.

The Venetic language, surviving at the north end of the Adriatic Sea in ca. 450 inscriptions attesting to fewer than 100 non-onomastic lexemes ^7^, is not universally considered to be an Italic language. But even the extremely fragmentary data attest to phonological developments shared with the Italic languages^8,9^ such that it may have been the first to branch off from this Indo-European subgroup^10^.

Archaeological investigation has identified several points in time at which new populations may have entered the Italian Peninsula, leading to a number of hypotheses about its Italicization, including via the Remedello, Rinaldone, Gaudo, Terramare, Proto-Villanovan, and Villanovan cultures^11^. These have included proposed arrivals from the North^5,12–14^, the North and East^15,16^, and even exclusively the East, i.e. trans-Adriatic^17^.

Purely linguistic lines of evidence highlight the importance of the Bronze Age-Iron Age transition in the Italicization question. Considerations on the difference between Latin and Sabellic suggest that the Proto-Italic linguistic community likely remained intact until the end of the Bronze Age^2,5^. The Latin word for ‘iron’, *ferrum*, reconstructs to quasi PIE **bʰersom*^15^. This development of PIE **bʰ* is an Italic one, suggesting that familiarity with iron technology reached a still intact Proto-Italic-speaking population.

Previous genetic studies on modern Italian populations had reported a decreasing North-to-South cline in Y-haplogroup R1b^18,19^, and aDNA sequencing identified Steppe ancestry reaching north Italy (in individuals buried with Bell Beaker cultural materials) during the latter half of the 3rd millennium and central Italy ~ 1,600 BCE^20–22^. Meanwhile, archaeological links between northern Italy and the Hungarian plain have long been suspected^23–25^ and more recently elaborated upon^5,26–28^.

The present IBD results are consistent with either of two scenarios for the introduction of the Proto-Italic language: 1) Via the arrival of the Steppe-related Bronze Age cluster with links to Central Europe and the Adriatic (Genetics and Strontium Supplementary Fig. S6.13; S6.14), consistent with the archaeological links between Northern Italy and Hungary; 2) Via the arrival of a Bell Beaker sub-population with links to populations in France, the Alps, and some Únětice sites. Both scenarios offer a genomic analogue, in the form of Bell Beaker-derived ancestry, for the Proto-Italo-Celtic linguistic subnode suggested by shared innovations between the Italic and Celtic branches^29–31^. Some studies (cf. Cardarelli 2009) have suggested that southward population movements subsequent to the ca. 1,200 BCE collapse of the Terramare culture could have served as the demographic vector for the spread of the Italic languages into Italy. Our results provide evidence that the peninsula was Indo-Europeanized before this date. Whether the earliest bearers of Steppe ancestry in Italy brought Proto-Italic with them cannot be determined with certainty, and we may be dealing with a situation similar to the arrival of Bell Beaker ancestry in the British Isles, which is generally agreed by linguists to have arrived too early to represent speakers of Celtic^32–34^. It is however certainly not impossible that Proto-Italic arrived in Central Italy ca. 1,700 BCE, first diverging into the no longer mutually intelligible branches of Latin and Sabellic in the Early Iron Age.

Northern Italy had evidently served as a cultural and genetic melting pot, as demonstrated by sites like Olmo di Nogara. Whether the Italic branch of Indo-European entered Italy from an eastern or western source, the residence of its speakers in this diverse region must have in part resulted in the body of prehistoric loanwords that can be demonstrated to exist within the Latin lexicon^35–39^.

#### 2.2 Etruscan

The Etruscan language is not Indo-European. Attested from c. 700 BCE to the middle of the first century CE, it was at home in ancient Etruria, modern Tuscany. Despite the existence of around 12,000 inscriptions, only around 9 documents (inscriptions and fragments of a linen book) are of any length. Despite a relatively good understanding of its morphology, much less is understood of its lexicon^10,40^. Etruscan’s only relatives are Lemnian, attested as a handful of inscriptions on the Aegean island of Lemnos, and Rhaetic, attested in several inscriptions from the Roman Alpine province of Rhaetia^41,42^, together forming the Tyrsenian (or Tyrrhenian) language family. Some scholars^43–45^ have adduced arguments in support of Herodotus’ account that Etruscan originated in Anatolia whereas others argue that it is autochthonous to Italy, the position taken by Dionysius of Halicarnassus. Linguistically, Etruscan and Lemnian appear more closely related to each other than either is to Rhaetic, suggesting that Rhaetic split from the family first and consequently that the split likely occurred in Italy^46,47^. Posth et al. (2021) found that, by 800 BCE, populations in Western Central Italy were genetically homogenous and could not distinguish between individuals from Etruscan and successive contexts. The IBD methods applied to the Posth et al. data furthermore show that individuals in Italy (including from Etruscan contexts) generally derive their farmer ancestry from a predominantly Italian farmer subcluster (Genetics and Strontium Supplementary Fig. S6.12).

#### 2.3 Messapic

Messapic is a fragmentarily attested language of the Salento peninsula of Italy surviving in ca. 600 inscriptions from the 6th c. BCE until the Roman conquest of the region^48^. Though an Indo-European language, Messapic does not belong to the Italic subgroup. Its further classification is hampered by its fragmentary attestation, but lexical correspondences with modern Albanian, visible despite the 1,500 years separating their attestation, point to a trans-Adriatic origin^49^. Three individuals from Coastal/Southern Italy indicate a gene flow from Balkans, particularly from Montenegro and Greece (Genetic Supplementary Fig S6.11). The number is small, and these individuals are from disparate contexts and therefore do not seem to be members of a single community. Some of them could, e.g., represent Mycenaean immigrants. But the presence of Balkan ancestry components in BA Southern Italy is not inconsistent with the possibility of Messapic having a Balkan origin.

### 3. Languages of Iberia

#### 3.1 Celtiberian

Celtiberian was a Celtic language of Pre-Roman Iberia. It is predominantly recorded in inscriptions found in Northeastern Spain from the 2nd century BCE to the 1st century CE^50^. The inscriptions are written in the native Celtiberian script. This script derives from the Northeastern Iberian semi-syllabary used for the unclassified (Paleo-)Iberian language, which is documented between the 6th and 1st centuries BCE.

Celtic language varieties were spoken across much of Iron Age Europe. Apart from Celtiberian, important Celtic language groups are Goidelic, Brittonic, Gaulish (including Cisalpine Gaulish and Balkan Celtic), Lepontic, and Galatian^51^. The Tartessian language, known from inscriptions from Southwestern Iberia between the 8th and 5th centuries BCE, has also been classified as a Celtic language^52^, but is widely taken as a linguistic isolate^32^.

Published and unpublished genomes included in the present study show that much of Iberia, including the south, was impacted by Bell Beaker-mediated Steppe ancestry in the second half of the 3rd millennium BCE. The Bronze Age individuals, including the MBA Andalusian individuals in the future Tartessian sphere of influence as well as the MBA Catalonian individuals in the future Iberian language area, it appears to have become homogenized, implying that genetic makeup in this period did not correlate with linguistic affiliation. This is underlined by the genetic makeup of modern speakers of Basque, a non-Indo-European language^53^. More generally, the sampled BA populations of Iberia are too early to confidently correlate them with any of the known IA language communities.

The emergence of Steppe ancestry can theoretically be correlated with the arrival of an early form of Celtic. However, there is a time gap of over two millennia between the emergence of this ancestry type and the first Celtiberian inscriptions, leaving room for more recent language intrusions. For instance, the Urnfield culture has previously been proposed as a vector for the introduction of Celtiberian to Spain, even though the area in which it appears rather correlates with the historical distribution of the unclassified (Paleo-)Iberian language^54^.

The question of the origin of Celtiberian depends on the more fundamental debate on the origin of the Celtic languages. Contrasting hypotheses exist on the location of the Celtic homeland, ranging between Central Europe^55^, Italy^30^, France^32^, and the Iberian Peninsula^56^. A recent palaeogenomic investigation of the historically Celtiberian region finds admixture from Central Europe between 1,300 and 800 BCE^57^, matching the timing of mainland ancestry, and potentially also Celtic, appearing in Britain^58^.

#### 3.2 Lusitanian

Lusitanian is a sparsely documented Pre-Roman Indo-European language of West-Central Iberia^59^. It is known from a small number of Lusitanian inscriptions, as well as from Latin inscriptions containing Lusitanian elements^60^. Due to its fragmentary nature, it is difficult to establish the exact linguistic affiliation of the language. It has been classified as an archaic form of Celtic^61,62^, but is more generally held to be non-Celtic and potentially closer to Italic than to Celtic^63^. The presence of a non-Celtic language in Iberia thus further complicates the identification of the mid-3rd millennium wave of Steppe ancestry with Celtic. Instead, this wave could have been associated with a largely undifferentiated set of Italo-Celtic dialects. But if Celtic is a more recent intrusion (see above), Lusitanian might have developed among the local populations resulting from this initial wave^64,65^.

### 4. Languages of the Eastern Mediterranean and the Caucasus

#### 4.1 Greek

Mycenaean Greek is the earliest known form of the Greek language, written in the Linear B syllabic writing system. It is attested between the 14th and 12th centuries BCE in inscriptions found mainly on Crete and the Peloponnese^66^. After the so-called Dark Ages, Greek appears in alphabetic writing again from the 8th century BCE onwards, now having split into several dialects, including Attic, Aeolic, Doric, Ionic, and Northwest Greek. Mycenaean is not ancestral to these dialects but instead close to the South Greek ancestor of Arcadocypriot, the dialect group of the central Peloponnese and Cyprus^67,68^. Arcadocypriot is native to the Greek mainland but spread to Cyprus during the Bronze Age, where it is documented in the Cypriot syllabary between the 11th and 4th centuries BCE^67^. A few individuals from Hala Sultan Tekke and Lapithos have a genetic pattern similar to Greece Bronze Age individuals, potentially revealing an affiliation with Mycenaean/Late Helladic settlers (Genetics and Strontium Supplementary Fig. S6.45; S6.46).

As to when the earliest Greek speakers may have entered the Greek mainland, our results suggest that it must have happened at the beginning of the 2nd millennium BCE, several centuries before the emergence of written Mycenaean Greek. By the Late Mycenaean period, the Steppe component is generally diluted to low proportions compared to Northern and Central Europe. This harmonizes with the observation that Greek contains a large amount of loanwords from one or more “Pre-Greek” languages, which cannot be equated with any known language^69,70^. Some of these loanwords are attested already in Mycenaean, e.g., *da-pu_2_-ri-to* ‘maze’, corresponding to Greek *labýrinthos*. However, one individual from Apollo Maleatas in the Peloponnese, dated to 1,933–1,774 calBCE, modeled as an admixture of Neolithic and Late Yamnaya ancestries with a slightly higher steppe proportion than Late Bronze Age individuals, suggesting that he might have belonged to the earliest generations of Indo-European, or more specifically, Greek-speaking people in the Peloponnese (Genetics and Strontium Supplementary Fig S6.18).

The closest known relative of Greek is the Phrygian language (see the following section), but the position of Greek in the higher order phylogeny of Indo-European remains a matter of controversy. It is most often grouped with Armenian and Albanian^71–74^, a subgrouping which has also found support by computational studies that rely on Bayesian inference^75–77^. This so-called “Graeco-Armenian” hypothesis has, however, been met with criticism too^68,78^, and it has been argued that Armenian is equally close, in a phylogenetic sense, to the Indo-Iranian clade^79,80^. Alternatively, Graeco-Phrygian has been grouped with Albanian^81^, in a node which connects to Armenian only at a higher (“Balkan Indo-European”) level.

Previous palaeogenomic studies have modeled LBA Greeks as admixed with either Yamnaya individuals or LBA Armenians^82,83^. On the basis of this evidence, the hypothesis of an arrival of the Greeks from the Caucasus^84^ could still not be excluded. Later studies^85,86^ have demonstrated a flow of Steppe ancestry in the MBA, but without narrowing down a more proximate source of this admixture. In this study, we show that MBA individuals from both Greece and Armenia are best modeled as having shared ancestry with previously unpublished samples from Moldova, associated with the Late Yamnaya culture (2,600–2,200 BCE). These results are therefore not inconsistent with the Graeco-Armenian primordial grouping, and they suggest that independent migrations on both sides of the Black Sea provide the best explanation for the arrival of the Greek and Armenian languages in their respective regions.

#### 4.2 Phrygian

Phrygian is a fragmentarily attested language. It is known through alphabetic inscriptions found in central Anatolia, primarily within the borders of the ancient Kingdom of Phrygia, whose capital was Gordion^87^. The oldest of these inscriptions make up the Old Phrygian corpus. They are dated between the 8th and 4th centuries BCE and written in a native script. New Phrygian emerges in inscriptions in the 2nd and 3rd centuries CE and is written in the Greek alphabet^88^.

Traditionally, Phrygian has been grouped with the poorly understood Thracian language, as part of a single “Thraco-Phrygian” ethnolinguistic entity^89,90^, largely based on statements by Herodotus and other ancient authors^91^. This implies that, at the time of their earliest attestation, the Phrygians were relatively recent migrants to Anatolia. However, it is now widely agreed that Phrygian is most closely related to Greek^88,92–94^.

The hypothesis that the Phrygian language is the result of a migration from the Balkans has hitherto shown difficult to support with archaeological or genetic evidence. Previous genetic studies were only able to demonstrate a proportion of approximately two percent EHG ancestry in samples from IA Phrygia^86^. Newly sampled individuals from IA Kalehöyük, Antandros, and Keçiçayırı all show low amounts of Steppe ancestry. One individual from Keçiçayırı is most relevant to the Phrygian question as it is found in an elite burial context and shares ancestry with Greek LBA individuals. This suggests that at least some individuals among the Phrygian ruling class descended from IA migrants from the Balkans, and it is consistent with the hypothesized Graeco-Phrygian clade. The two other individuals, however, have Steppe ancestry of the Lower Volga type, which appears in Armenia at the end of the 4th millennium BCE and may ultimately be associated with the Anatolian branch (see below). Once again, this underscores the high ethnolinguistic complexity of IA Anatolia and calls for additional data if firmer conclusions are to be made.

#### 4.3 Armenian

Armenian is currently spoken in the Republic of Armenia and by a worldwide diaspora, but it has historically formed a patchwork of dialects across large parts of Anatolia and the South Caucasus^95^. Its first substantial attestation is the Classical Armenian literature appearing from the 5th century CE. Traditionally, it is considered an independent branch of the Indo-European family tree^96^, but is frequently placed in a higher-order subgroup with Greek^72,74^. As previously mentioned, our new IBD analyses show that BA individuals from both Greece and Armenia are best modeled as having shared ancestry derived from a population closely related to previously unpublished MBA samples from Moldova, associated with the Late Yamnaya culture (Genetics and Strontium Supplementary Fig. S6.21; S6.42). This contrasts with, e.g., individuals associated with Italic languages, who derive their Steppe ancestry by a vector of Corded Ware and Bell Beaker individuals. These results are consistent with the assumption of a primordial Graeco-Armenian subgroup that started diverging during the middle of the 3rd millennium at the latest. And thus, the rather sudden replacement of the previously widespread Transcaucasian Kura-Araxes culture by the Trialeti culture by the end of the 3rd millennium BCE^97^, with certain similarities to early Mycenaean culture^26^, probably represents the first tangible sign in the region of an Indo-European element that can be ancestral to the Armenian branch (Anthony 2024; Lazaridis et al. 2022a).

From the Iron Age, samples with Urartian and pre-Urartian contexts show a similar proportion of ancestry associated with the western Steppe, which is consistent with the existing view that the Urartian population was multiethnic^99^ and multilingual^100,101^, and it yields support for the hypothesis that it may have contained an Armenian-speaking component^102,103^. Moreover, as mentioned earlier, Steppe ancestry emerges in the South Caucasus already in the MBA, with no significant later input, and it is only a marginal ancestry component in Central Anatolia. This means that the traditional hypothesis of a migration of Armenian speakers across Anatolia after 1,200 BCE^11,104^ is increasingly doubtful.

Many scholars have assumed a particularly close relationship between (Thraco-)Phrygian and Armenian as well^94^, even closer than that of Greek and Phrygian^105,106^. However, more recent progress in the study of Phrygian has revealed a poverty of *exclusively* shared features with Armenian, which makes such a hypothesis difficult to support^94^. Likewise, our IBD results yield no support for assuming a common migration of Armenians and Phrygians through Anatolia, but rather suggest that the shared innovations of Greek, Phrygian, and Armenian are attributable to a higher-order subgroup (or linguistic area) connected with the Late Yamnaya culture of the 3rd millennium BCE.

#### 4.4 Anatolian

The Anatolian branch is an extinct branch of the Indo-European language family, attested from at least the 20th century BCE onwards, consisting of Hittite (known 20th–12th centuries BCE), Luwian (known 20th–7th centuries BCE), and a number of less well-attested members, such as Carian, Lycian, Lydian, and Palaic.

The position of Anatolian within the Indo-European family tree is debated^107^. Some treat it on par with the other branches^108–112^, but it is widely held to be the branch that diverged from the proto-language first^107,112–114^. According to the latter view, commonly referred to as the Indo-Hittite or Indo-Anatolian hypothesis, Anatolian is paraphyletic to Indo-European^115^ and Proto-Anatolian contemporaneous with the Yamnaya culture^116^. An archaic feature of Anatolian is the lack of the core Indo-European terminology related to plow agriculture^117^ and wheel and wagon terminology^118^.

The Indo-European Anatolian branch is central to the debate on the Indo-European linguistic homeland, on which two main rival hypotheses exist: the Steppe hypothesis and the Anatolia hypothesis. According to the Steppe hypothesis, the Anatolian branch originates in a Pre-Yamnaya Steppe population that migrated to Anatolia between the late 5th and 4th millennia BCE, after which the remaining Indo-European languages dispersed with the expansion of the mobile pastoralists of the Yamnaya culture. Within this hypothesis, two main views exist on the point of entry into Anatolia^119^. A western dispersal through the Balkans has been supported with archaeological evidence, i.e. the movement of the Suvorovo-Novodanilovka group (4,400–4,200 BCE) into the Balkans and the subsequent rise of the Cernavodă culture (4,000–3,200 BCE)^5^, as well as with the center of diversity of the Anatolian languages being in Western Anatolia^120^. The alternative consists of an eastern dispersal through the Caucasus in the third millennium BCE. Alongside archaeological and linguistic evidence^121–123^, it has been supported with Hittite legendary evidence: an address of King Muwatalli II (±1,320–±1,294 BCE) to the Sun-God depicts the sun as rising from the sea, which has been interpreted as reflecting a memory of an Anatolian presence on the eastern flanks of the Caucasus^124–127^. But even when taken at face value, this depiction at best implies a memory of an “eastern litoral”^128^, either on the Caspian or the Black Sea.

The contrastive Anatolia hypothesis rather sees Anatolia as the cradle of the Indo-European languages, from which all Indo-European branches except Anatolian dispersed in various directions following the expansion of farming from the 7th millennium BCE^129^. While now largely abandoned, a related, recently proposed model claims Anatolia or the Greater Caucasian region as the primary homeland, but allows for secondary dispersals through the Yamnaya culture^77,86,130^. This third model recalls an older mixed Anatolia/Steppe model^131^ as well as the traditionally marginal homeland hypothesis known as the Near Eastern model^14,132^.

Subtle amounts of Steppe ancestry appear in Armenia (Main Text, Fig. 6) before the 3rd millennium BCE. Despite justified doubts about the relevance of some of the cultural contexts^120^, Steppe ancestry additionally appears in previously reported genomes of 2nd millennium BCE Anatolia (Main Text, Fig. 6). This recalls and complements recent findings of ancestry from an Eneolithic Lower Volga source entering Anatolia, presumably through the Caucasus, appearing in individuals from EBA Ovaören and MBA Kalehöyük, and possibly being related to the dispersal of the Anatolian Indo-European branch^133^.

### 5. Conclusion

Italy and Spain received a significant amount of their Steppe ancestry from Bell Beaker-derived populations. Greeks and Armenians, on the other hand, derive their Steppe ancestry directly from a western Yamnaya subpopulation. Previous studies have revealed Corded Ware ancestry among Balts, Slavs and Indo-Iranians^82,134,135^. It is now increasingly feasible to hypothesize the demographic channels through which Indo-European spread and split. The precursors of Italic and Celtic, as well as Lusitanian, were possibly mediated by the Bell Beaker population that genetically formed in Central Europe. A shared ancestor of Indo-Iranian and Balto-Slavic (i.e. “Indo-Slavic”) evolved among populations of the Corded Ware in East Europe. Both of these populations were formed by admixture with European farmers. In contrast, the predecessor of Armenian and Greek evolved among Yamnaya populations that had remained on the Steppe until the Middle Bronze Age. Finally, however, it does not yet seem feasible to identify an archaeologically defined vector for a Steppe intrusion of Anatolian into Anatolia.

**Proto-Indo-Anatolian (basal IE)**

︱	︱

*Yamnaya*	???

︱	︱

**Proto-Indo-European (core IE)	Proto-Anatolian**

︱	︱	︱

*Bell Beaker	Corded Ware	Late Yamnaya*

︱	︱	︱

**Italo-Celtic	Indo-Slavic	Graeco-Armenian**

**Figure 1**: Highly simplified Indo-European phylogeny correlated with the archaeological macrostructure of 4th and 3rd millennium BCE European Steppe-derived populations.
