## Supplementary material for "Ancient genomics support deep divergence between Eastern and Western Mediterranean Indo-European languages": Archaeological Supplementary

**Archaeological Supplementary Information**

Serena Sabatini^1^

^1^Department of Historical Studies, University of Gothenburg, Sweden.

INTRODUCTION	4

**1. ARCHAEOLOGICAL OVERVIEW AND DISCUSSION	5**

1.1. Main chronological phases of the Mediterranean Bronze Age	5

1.2. Anatolia and the Levant	7

1.2.1. The Levantine coast	8

1.2.2. Anatolia	8

1.2.3 Conclusive reflections	9

1.3. Cyprus	9

1.3.1 Third millennium BCE Cyprus	9

1.3.2 Second millennium BCE Cyprus	10

1.3.3 Conclusive reflections	12

1.4. Continental Greece	12

1.4.1 The Early and Middle Helladic Period	12

1.4.2 The Late Helladic Period	13

1.4.3 Conclusive reflections	14

1.5. Italy	14

1.5.1. Northern and continental Italy	15

1.5.2. Peninsular Italy	16

1.5.3 Conclusive reflections	17

1.6. Spain	17

1.6.1. Catalonia, north-eastern Spain	17

1.6.2. Andalusia, south-eastern Spain	18

1.6.3. Castilla y La Mancha, central Spain	18

1.6.4. Conclusive reflections	18

**2. CATALOGUE OF THE ARCHAEOLOGICAL SITES	19**

2.1 Introduction	19

**2.2. Cyprus	20**

2.1.1. Agios Iakovos necropolis, (de facto) İskele District, Northern Cyprus	20

2.2.2. Amathus necropolis, Limassol district	24

2.2.3. Karavas, (de facto) Kyrenia district, Northern Cyprus	27

2.2.4. Kythrea, (de facto) Lefkoşa district, Northern Cyprus	28

2.2.5. Lapithos necropoleis, (de facto) Kyrenia district, Northern Cyprus	29

Lapithos, Vrysi tou Barba cemetery	30

Lapithos, Kastros cemetery	32

Lapithos, Plakes cemetery	34

2.2.6. Rizokarpaso, (de facto) Famagusta district, Northern Cyprus	38

2.2.7. Vounos-Bellapais, (de facto) Kyrenia district, Northern Cyprus	39

2.2.8. Hala Sultan Tekke, Larnaca district	40

**2.3. Egypt	43**

2.3.1. Maasara, Helwan, Cairo Governorate	43

**2.4. France	44**

2.4.1. Lano Cave, Lano commune, Haute-Corse Department of France, Corsica	44

2.4.2. Migennes, Yonne department in Bourgogne-Franche-Comté	45

**2.5. Greece	49**

2.5.1. Apollo Maleatas, Argolis, Peloponnese	49

2.5.2. Ayios Vasileios, Laconia, Peloponnese	51

2.5.3. Kalyvia, Elis, Western Greece	54

2.5.4. Kirrha, Phocis, Central Greece	55

2.5.5. Voudeni, Achaea, Western Greece	58

2.5.6. Eleon, Boeotia, Central Greece	69

**2.6. Hungary	71**

2.6.1. Pitvaros, Csongrád County	71

2.6.2. Szőreg C cemetery, Szeged, Csongrád County	73

2.6.3. Tápé – Széntéglaégető cemetery, Szeged, Csongrád County, Hungary	75

**2.7. Italy	78**

2.7.1. Coppa Nevigata, Foggia province, Puglia	78

2.7.2. Lavello, Potenza province, Basilicata	79

2.7.3. Lucone, Brescia province, Lombardy	80

2.7.4. Mereto, Udine province, Friuli- Venezia Giulia	81

2.7.5. Narde, Rovigo province, Veneto	82

2.7.6. Olmo di Nogara, Verona province, Veneto	83

2.7.7. Pian Sultano, Rome province, Latium	97

2.7.8. Scalvinetto, Padova province, Veneto	98

2.7.9. Sedegliano, Udine province, Friuli- Venezia Giulia	99

2.7.10. Selvis, Udine province, Friuli-Venezia Giulia	100

2.7.11. S. Osvaldo, Udine province, Friuli- Venezia Giulia	101

2.7.12. Toppo Daguzzo, Potenza province, Basilicata	102

**2.8. Lebanon	103**

2.8.1. Sidon, Sidon district, South Governorate	103

**2.9. Moldova	111**

2.9.1. Balabanu II, Taraclia district, southwestern Moldova	111

2.9.2. Cahul/Crihana Veche, Cahul district	112

2.9.3. Cazaclia, Ceadîr-Lunga district, Southwestern Moldova	113

2.9.4. Constantinovca, Slobozia district, Transnistria, southeastern Moldova	115

2.9.5. Cotiujeni, Șoldănești district, Northern Moldova	116

2.9.6. Taraclia, Taraclia district, southern Moldova	117

**2.10. Spain	118**

2.10.1 Argar, Antas, Almería, Andalusia	118

2.10.2. Can Martorell, Dosrius, Barcelona, Catalonia	119

2.10.3. Cantorella, Maldà, LLeida province, Catalonia	120

2.10.4. Castellon Alto, Galera, Granada, Andalusia	122

2.10.5. Cerro de la Virgen, Orce, Granada, Andalusia	123

2.10.6. Cerro de la Encantada, Granátula de Calatrava, Ciudad Real	125

2.10.7. Cuesta del Negro, Purullena, Granada, Andalusia	126

2.10.8. Fuente Álamo, Cuevas del Almanzora, Almería, Andalusia	127

2.10.9. Los Torcales 6, Beas, Huelva, Andalusia	128

2.10.10. Minferri, Juneda, LLeida, Catalonia	129

2.10.11. Motilla del Azuer, Daimiel, Ciudad Real province, Castilla y La Mancha	131

2.10.12. Necrópolis de los Algarbes, Tarifa, Cádiz	132

2.10.13. Terrera del Reloj, Dehesas de Guadix, Granada province, Andalusia	134

2.10.14. Valencina de la Concepción, Calle Trabajadores 14-18, Seville, Andalusia	135

2.10.15. Valencina de la Concepción, Calle Dinamarca 3-5, Seville, Andalusia	137

2.10.16. Cueva de la Carada, Huéscar, Granada, Andalusia	139

**2.11. Syria	140**

2.11.1. Tall Sūkās, Latakia Governorate	140

**2.12. Turkey	141**

2.12.1. Antandros, North-western Anatolia	141

2.12.2. Resuloğlu, Çorum province	142

2.12.3. Kaman-Kalehoyük Kazisi, Kırşehir province	144

2.12.4. Küllüoba Kazısı, Eskişehir province	146

2.12.5. Keçiçayırı Kazısı, Eskişehir province	148

**References	149**

### INTRODUCTION

The archaeological Supplementary Information is organised in two main parts:

1. Archaeological overview and discussion
2. Catalogue of the Archaeological sites

In the first part, the reader will be presented with a critical summary of the archaeology of the investigated regions; the aim of this first block is to critically discuss characteristics, and challenges, that the contexts from which the sampled individuals were collected offer for our understanding of the Bronze Age across the Mediterranean. The first part discusses each region, separately following a geographical order from the eastern coast of the Mediterranean to the Iberian Peninsula in the west.

The second part consists of a detailed catalogue of all archaeological sites involved in this project and of all the specific archaeological contexts or graves from which the sampled individuals discussed in the main paper come from. In the catalogue, to facilitate reading, the order is alphabetical per country of provenience and then per archaeological sites (see § 2.1)

All in all the aim of this supplementary is to contribute to a deeper understanding and contextualization of the results presented in the main paper and to provide consideration which may stimulate future research.

### 1. ARCHAEOLOGICAL OVERVIEW AND DISCUSSION

#### **1.1. Main chronological phases of the Mediterranean Bronze Age**

Before going through the archaeological contexts, it is crucial to briefly address the chronology of the Mediterranean Bronze Age. The purpose of this section is to provide a simplified and yet accurate picture of the main chronological phases and periods named both in the main text of the article and in the various supplementary information files.

There is no room to thoroughly discuss issues and problems linked to the establishment of reliable chronological frames of reference for all the areas of the Mediterranean that are discussed in this work. Likewise it would be impossible to include a list of the dedicated publications. Both relative and absolute chronology of single sites or larger regions of the Mediterranean has been variously analysed and presented. The proposed chronological table (Table 1) is to be considered as a comprehensive and yet orientating chart, built on a selection of widely acknowledged chronological studies^1–6^.

The samples analysed within this project come from the whole Mediterranean region and beyond; from the Iberian Peninsula in the west, to Syria and Lebanon in the east, and from mid-west France in the north until Egypt in the south. From a chronological point of view most of the samples belong to the second millennium BCE, however in some areas a limited number of earlier and later samples has been collected as well. Our table (Table 1) has no ambition to be comprehensive for all regions included in this study; it rather provides an overall view of the main chronological phases of the areas from which most samples have been collected and analysed. More specific information about chronology can be found in the specific description of the various contexts.

| **IBERIA** | **ITALY** | **GREECE** | **CYPRUS** | **ANATOLIA** |
| --- | --- | --- | --- | --- |
| Late Chalcolithic  2550-2200 | Late Chalcolithic  2800-2150 | Early Helladic I-III  3100-2100/2050 | Late Chalcolithic /Philia phase  2700-2250 | Early Bronze  Age I-III  3000-2000 |
|  |  |  | Early Cypriot I-II 2250-2000 |  |
| Early Bronze Age  2200-1925 | Early Bronze Age  2150-1650 | Middle Helladic  2100/2050-1700/1675 | Early Cypriot III /Middle Cypriot I-II  2000-1750/1700 | Middle Bronze Age I-III  2000-1700 |
| Middle Bronze Age  1925-1600 | Middle Bronze Age 1  1650-1550 | Late Helladic I  1700/1675-1635/1600 | Middle Cypriot III  1700/1750-1650 | Middle Bronze  Age IV  1700-1600 |
| Late Bronze Age IA  1600-1550 | Middle Bronze Age 2A  1550-1500 | Late Helladic IIA  1635/1600-1480/1470 | Late Cypriot IA  1650-1550 | Late Bronze Age 1a  1600-1500 |
| Late Bronze Age IB  1550-1450 | Middle Bronze Age 2B  1500-1450 | Late Helladic IIB-IIIA1  1480/1470-1390/1370 | Late Cypriot IB  1550-1450 | Late Bronze Age 1b  1600-1400 |
|  | Middle Bronze Age 3A  1450-1400 |  |  |  |
| Late Bronze Age IC    1450-1300 |  |  | Late Cypriot IIA-IIB  1450-1300 BC | Late Bronze Age 2a    1400-1300 |
|  | Middle Bronze Age 3B  1400-1325/1300 | Late Helladic IIIA2  1390/1370-1330/1315 |  |  |
| Late Bronze Age II  1300-1050 | Recent Bronze Age 1  1325/1300-1225/1200 | Late Helladic IIIB  1330/1315-1200/1190 | Late Cypriot IIC  1300- 1200 | Late Bronze Age 2b  1300-1200 |
| Late Bronze Age III  1050-900 | Recent Bronze Age 2  1225/1200-1150/1025 | Late Helladic IIIC  1200/1190-1075/1050 | Late Cypriot IIIA-IIIB  1200-1050 | Early Iron Age I  1200-900 |
| Early Iron Age I    900-600 | Final Bronze Age  1150/1025-925/900 | Sub-Mycenaean  1075/1050-950 | Cypro-Geometric I-III  1050-750 | Early Iron Age II    900-750 |
|  | Early Iron Age  925/900-750 | Protogeometric /Geometric period  950-750 |  |  |
|  |  |  |  | Early Iron Age III    750-550 |
|  | Orientalizing Period  750-580 | Archaic period  750-480 | Cypro-Archaic  750-600 |  |

Table 1. Chronological table (all dates are BCE) with the main periods and the main regions mentioned in this work. Data from main chronological studies^1–6^.

#### **1.2. Anatolia and the Levant**

Anatolia and the Levantine coast of the Mediterranean Sea are two distinct neighbouring regions with a complex and interlaced history throughout the Bronze Age. Differently from other parts of the Mediterranean, the study of the archaeological evidence from these regions can be complemented by a diverse array of ancient written texts and documents expanding tremendously the possibility to reconstruct their Bronze Age world, despite interpretation is at times debated^7–10^.

From a historical point of view, it seems that the political economic organisation of Near Eastern/ Mesopotamian societies underwent considerable development from the early state societies of the 4^th^ millennium BCE to the mature states of the subsequent 3^rd^ millennium^11^. Such development included among other things a more direct control over craftsmanship and trade, possibly affecting individual mobility^12^. In Anatolia, the state organisations proper with urban societies and control over staple economy and extended trading systems would emerge in the 2^nd^ millennium BCE^7,9,11,13,14^.

Along the Levantine coast, corresponding approximately to modern day Lebanon and Israel, the 2^nd^ millennium is a time of flourishing trade and international relations^15^, affected to a considerable extent by the expansive power of the Egyptian, on the one hand, and of the Hittite kingdom, on the other^16,17^. The Late Bronze Age in mid-2^nd^ millennium BCE is a relatively short period of great connectivity across the Mediterranean and beyond; wide exchange networks and extensive bulk trade in metal, textiles and other raw materials reach astonishing levels^18–22^, but comes to an end at the onset of the 12^th^ century BCE, when major socio-political transformations occur both in the Aegean and the eastern Mediterranean. A complex set of largely debated critical events, including migratory fluxes, generally linked to the phenomenon of the so-called Sea people, irremediably transforms the geopolitical setting of the whole area^23–26^.

##### *1.2.1. The Levantine coast*

In our study we have been focusing on two sites, the ancient city of Sidon in today Lebanon and the burial mound of Tall Sūkās, in southern coastal Syria.

The city of Sidon is often mentioned in ancient written sources and is no doubt one of the most prominent of the Canaanite/Phoenician cities of the late second and first millennium BCE^27,28^. Since the modern town lies right above its ancient predecessor, the Phoenician city is not well-known; relatively few archaeological remains are known or have been excavated. The samples we collected are therefore of particular interest, in the first place because they come from the so-called ‘College site’ in down-town modern Sidon and thus also in the heart of the ancient city; secondly, because the excavation revealed a continuity of occupation from the Chalcolithic period until the Phoenician period and later during historical times (see § 2.8). In other words, our samples provide a unique opportunity to analyse the development of the local population through time.

During the local Middle Bronze Age (cf. Table 1) to which the levels 1-8 at the College site in Sidon correspond (see below), the archaeological evidence suggests extensive contacts with the world beyond the Levantine coast both to the west and to east. The sampled individuals have the possibility to show the possible impact that contacts, trade and exchange with the outside world could have on the local population. There is only one individual, dated to the Iron Age (CGG_2_104309) - thus after the 1200 BCE critical period at the end of the Bronze Age^23–26^; the comparison between this specific individual with those living in Sidon during the Bronze Age would provide a new piece of evidence as to the debated theories about incoming groups from the west at the end of the 13^th^ century BCE^26,29^.

The individuals from the Middle Bronze Age burial mound at Tall Sūkās^30^, in southern Syria are approximately contemporary to those of the Sidon College site levels 4-8; their characteristics also suggest that the local population along the Levantine coast prior to the crisis at the beginning of the 12^th^ centuries BCE had primarily commercial contacts with the rest of the Mediterranean region.

##### *1.2.2. Anatolia*

Our samples from Anatolia come from very different parts of the peninsula and cover a time range from the Early Bronze Age to the Hellenistic period. Given the vastness and the geographical complexity of the Anatolian Peninsula, our samples cannot be straightforwardly taken as representative for the wide population genomics of the region; however, they provide significant information about regional characteristics.

Individuals dated to the 3^rd^ millennium BCE, thus an advanced phase of Near Eastern Early Bronze Age (Table 1), have been sampled from the cemetery at Resuloğlu in Central Anatolia and from the burials at Küllüoba, in north-western Anatolia, which date from the Chalcolithic to the end of the Early Bronze Age (see § 2.12.2 and § 2.12.4). The samples from those roughly contemporary sites show slightly different archaeological evidence. Material culture from the site at Küllüoba shows that during the Early Bronze Age the local community had consistent trade with other communities of the North Aegean Sea. It is likely that site had a key role on the route between the coast and inland Northwestern Anatolia^31^. The grave goods from the cemetery at Resuloğlu show contacts with both Western Anatolia and with the Mesopotamian world^32^.

The large mound at Kaman-Kalehöyük, in central Turkey was in use for a long time from the Early Bronze Age to the Ottoman period^33^. Our samples come from Stratum IIIb and IIIc, which are dated respectively to the so-called Assyrian Colony period (Karum) and the subsequent Old kingdom/Hittite period, thus to the first half of the 2^nd^ millennium BCE. One could consider those samples to be representatives of Hittite populations. As known, the origin of the Hittites is debated^34^; the sampled individuals from Kaman-Kalehöyük therefore provide a fundamental contribution to this long-lasting debate. The limited number of sampled Iron Age individuals does not allow drawing general conclusions about the development of Anatolian populations after the great political transformations that occurred at the end of the 2^nd^ millennium BCE with the collapse of the Hittite kingdom. Our samples from the 1^st^ millennium BCE include one member of the community at Kecicayiri (CGG_2_022162) on the so-called Phrygian plateau, and several individuals from the Hellenistic phase of the large and complex necropolis at Antandros in the Troas.

##### *1.2.3 Conclusive reflections*

The limited number of samples from this region significantly hampers the possibility to get a complete picture of the local population during the Bronze Age. Like in the case of Cyprus (see chapter 2.1), during the 2^nd^ millennium BCE the archaeological evidence suggests that the region was embedded in a complex web of long-distance exchanges with the entire eastern Mediterranean and the Near East. Such prosperous trade underwent a period of crisis and of considerable transformation around 1200 BCE, with the collapse of the Hittite kingdom.

#### **1.3. Cyprus**

Cyprus lies in the heart of the Eastern Mediterranean. It is home to incredibly rich and distinctive archaeological cultures through time. The local archaeological record suggests great creativity and complex cultural expressions throughout prehistory^2,35^. Due to its strategic position and its rich mineral resources, it has been involved in long distance exchanges between the Levant, the Near East and Egypt, Crete, the Aegean, Sardinia, and the central Mediterranean; the remarkably rich copper deposits of the Troodos mountains dominate the economy of the island throughout the Bronze Age^36^.

##### *1.3.1 Third millennium BCE Cyprus*

The so-called Philia phase at the end of the Cypriot Chalcolithic (around 2400/2350- 2300/2250 BCE) is generally considered a sort of cultural turning point and a period of important transformations, possibly as the outcome of a colonisation process^4,37^, and possibly with a relevant influence from Anatolia^9^. The individuals from Karavas (CGG_2_022531) and Kythrea (CGG_2_22543) were sampled with the aim to shed new light on this early migration into the island; unfortunately, the context from which the sample from Kythrea was collected is uncertain (see § 2.2.4) and the sole evidence from Karavas is too scanty to be able to draw any firm conclusion.

| **Cypriot** | **Absolute dates** | **Near East** |
| --- | --- | --- |
| **Ceramic Neolithic** | 5500 – 3900/3700 BCE | Chalcolithic |
| **Chalcolithic** | 3900/3700 – 2500/2300 BCE | Early Bronze Age I-III |
| **Early Cypriot (EC)** | 2500/2300 – 2200/1900 BCE | Intermediate Bronze Age |
| **Middle Cypriot (MC) I-II** | 2200/1900 – 1750 BCE | Middle Bronze Age I |
| **Middle Cypriot (MC) III** | 1750 – 1650/1550 BCE | Middle Bronze Age II-III – Late Bronze Age |
| **Late Cypriot (LC) IB** | 1550-1450 BCE | Late Bronze Age |
| **Late Cypriot II** | 1450 – 1200/1150 BCE |  |
| **Late Cypriot (LC) III** | 1200/1150 – 1050/1000 BCE | Late Bronze Age – Iron Age I |
| **Cypro-Geometric I-II** | 1050/1000 – 900 BCE | Iron Age I |
| **Cypro-Geometric III** | 900 – 750 BCE | Iron Age IIA-B |
| **Cypro-Archaic – Cypro-Classical** | 750 – 330 BCE | Iron Age IIC-Persian Period |

Table 2 Main chronological periods in Cyprus archaeology^36^ compared with main periods of the Near East^38^.

##### *1.3.2 Second millennium BCE Cyprus*

All the other samples from Cyprus are dated to the 2^nd^ millennium BCE or later and include several periods (Table 2). The most intriguing individual from Cyprus in our study is probably the young men from Vounous-Bellapais (CGG_2_022535), who seem to have originated in a region corresponding to modern day Sweden confirmed by the evidence of the Strontium isotopes (Genetics and Strontium Supplementary S10). The results of our study unveil an extraordinary journey from Scandinavia to Cyprus terminated with death at Vounous Bellapais, by the northern shores of Cyprus. It has been discussed how Vounous Bellapais likely held a dominant commercial role at the beginning of the 2^nd^ millennium BCE. Our results suggest that the extensive exchange network of the site might have attracted travellers and merchants from far more distant regions than Anatolia and the eastern Mediterranean. Although not abundant, archaeological evidence suggesting long-distance contacts with the rest of the European continent at the beginning of the 2^nd^ millennium BCE includes the distribution of toggle pins (*schleifennadel*) which span from Central Europe to the East Mediterranean^39^. From the archaeological point of view, it is very interesting to consider that the context (grave 69) from which the Vounous Bellapais non-local individual was recovered does not provide any hint as to his far away origin^40^. The individual was not radiocarbon dated, thus room for some uncertainty about this individual should be left open; however, the evidence shows that genetic studies may unveil mobility patterns that the archaeological material at times does not seem to reveal.

It has been proposed that the dominant commercial role of Vounous-Bellapais lasted for a relatively short period from the Cypriot Early Bronze Age (EC) until the MC II (Table 2) and was later taken on by the neighbouring community established in Lapithos^41^ from which a large number of samples have been collected for this project. Laphitos, strategically positioned on the northern coast of Cyprus, facing Anatolia, became one of the main MC harbours for trade in copper^41^. The sizable degree of wealth in some of the burials suggest that a rich community was living there. Three different cemeteries have been sampled in Lapithos: *Vrysi Tou Barba*, in use approximately between 2050 and 1650 BCE; *Plakes* and *Kastros* dated to the Cypro-Geometric period (Table 1). The individuals from those different cemeteries give the outstanding possibility to analyse the development of the population living along the northern coast of the island throughout the entire 2^nd^ millennium and the first quarter of the 1^st^ millennium BCE.

The remaining sampled individuals from Cyprus come from the southern (Hala Sultan Tekke) and eastern part of the island (Agios Iakovos and Rizokarpaso). They are generally dated to the 2^nd^ half of the 2^nd^ millennium BCE (see chapter 2.1), thus to the LC period (see Table 2), a time of great connectivity with extraordinary archaeological evidence for long distance exchanges across the Mediterranean. The archaeological evidence from the well-excavated harbour site of Hala Sultan Tekke suggest in facts that it was home to a complex urban community where merchants, travellers, and possibly workers from the entire eastern Mediterranean including Crete, the Aegean, Anatolia, Egypt, the Levant and also Sardinia^42–44^. Hala Sultan Tekke seems abandoned at the end of the LC period in a time (after 1200 BCE) of crisis all over the eastern Mediterranean.

Our dataset includes a number of Cypriot samples from the early 1^st^ millennium BCE; thus, giving the possibility to discuss population transformation across time and most importantly after the 1200 BCE crisis at the end of the Bronze Age. Together with the individuals from the Cypro-Geometric tombs at the Lapithos cemeteries of Kastros and Plakes (see above), we also included a few individuals from the Cypro-Archaic site of Amathus in southern Cyprus. During the 1^st^ millennium BCE, the premises for long distance trade and exchange with the rest of the Mediterranean changed considerably; although debated, the role of Cyprus as a copper producing region remains significant^45^.

##### *1.3.3 Conclusive reflections*

Cyprus is an island rich in copper, the most fundamental component of bronze. The archaeological evidence from the island shows that the copper mineral was abundantly exploited and that local communities were able to engage in large scale trade and long-distance exchanges with the whole eastern Mediterranean and beyond. Harbour sites such Hala Sultan Tekke must have enjoyed privileged access to the world overseas, most likely hosting multicultural communities. The dramatic crisis of the 1200 BCE, is particularly evident on Cyprus and major transformations occur on the island at the end of the 2^nd^ millennium BCE.

#### **1.4. Continental Greece**

Modern Greece consists of a continental part at the very end of the Balkan Peninsula and an insular world, which is a complex constellation of archipelagos and single islands stretching from the Ionian Sea to the west until the Aegean Sea reaching the coast of modern Turkey. To the south, modern Greece includes the fifth largest island of the Mediterranean, Crete, home to the spectacular Minoan culture in the first half of the 2^nd^ millennium BCE. Traditionally the archaeology of those different parts of the country are considered separated both culturally and chronologically (Table 3), although clearly interacting with each other^3^. In our project we focused on continental Greece, to understand the genetic make-up of the classical Mycenaean population. To do that we included in our dataset samples from several cemeteries across western and central Greece and the Peloponnese. We also included samples from earlier phases than the LH/Mycenaean period proper to unveil transformations in a long-term perspective.

##### *1.4.1 The Early and Middle Helladic Period*

Our oldest individual comes from the Final Neolithic/EH cemetery of Kalyvia (CGG_2_002395), which is one of the few known burial grounds of such an early date (4^th^ millennium BCE) from the Peloponnese.

| **Mainland Greece** | **Crete** | **Cyclades** | **Absolute dates (BCE)** |
| --- | --- | --- | --- |
| **Early Helladic (EH) I-III** | **Early Minoan (EM) I-III** | **Early Cycladic (EC) I-III/Middle**  **Cycladic (MC) I** | 3100-2100/2050 |
| **Middle Helladic (MH)** | **Middle Minoan (MM) I-III** | **Middle Cycladic (MC) I-III** | 2100/2050-1700/1675 |
| **Late Helladic (LH) I** | **Late Minoan (LM) IA** | **Late Cycladic (MC) I** | 1700/1675-1635/1600 |
| **Late Helladic (LH) IIA** | **Late Minoan (LM) IB** |  | 1635/1600-1480/1470 |
| **Late Helladic (LH) IIB** | **Late Minoan (LM) II** |  | 1480/1470-1420/1410 |
| **Late Helladic (LH) IIIA1** | **Late Minoan (LM) IIIA1** | **Late Cycladic (MC) II-III** | 1480/1470-1390/1370 |
| **Late Helladic (LH) IIIC** | **Late Minoan (LM) IIIC** |  | 1200/1190-1075/1050 |

Table 3 Main chronological phase of the Aegean Bronze Age^3^.

The next oldest sampled individuals are indeed over a millennium younger than the lady from Kalyvia. They come from Argolis also in the Peloponnese and specifically from the site of Apollo Maleatas. The three individuals from the site come from three isolated pit graves on the northern edge of the local hill close to the remains of an EH village. Because of that they were dated to the EH period (see Table 3). However, different sets of radiocarbon analyses (see § 2.5.1) dated them to 1900-1750 BCE or an early part of the MH period (Table 3). By that time the neighbouring settlement had been abandoned and the hill had become the theatre of cultic activities, which intensified with time until the establishment of the Apollo sanctuary in the classical period. In critically assessing the results of the genetic studies, one might consider the personal history of the individuals from Apollo Maleatas possibly linked to the sacred nature of the site.

##### *1.4.2 The Late Helladic Period*

The LH period (see Table 3) in mainland Greece is a complex period, which sees what we could call the exponential development and climax of the Mycenaean world and its dramatic end at the end of the 12^th^ century BCE. Despite well studied archaeological and textual evidence, little is known about the characteristics of the local population. The individuals sampled for this project provide the unprecedented possibility to document possible migratory events that contributed to the shaping of Myceanean communities. Our samples (see chapter 2.3) have been in fact collected from core regions of the Greek mainland such as Phocis (Kirrha) and Boeotia (Eleon) in central Greece, Achaea (Voudeni) and Laconia (Ayios Vasileios) in the Peloponnese . All those sites are archaeologically speaking classical Mycenaean contexts and the obtained results will undoubtedly be of great value for many future studies.

##### *1.4.3 Conclusive reflections*

As to the archaeology of mainland Greece, it has been discussed how profound changes can be observed for example in architecture and settlement organisation, but also material culture at the end of the of 3^rd^ millennium BCE and in particular the end of the EH II and the beginning of the EH III^46^; it has been proposed, on the base of the archaeological evidence that small scale migrations from Anatolia occurred and contributed to transform the Aegean world with the exception of Crete, which seems to follow a different trajectory. Indeed, there are also significant connection with the Balkans and the Adriatic world^47^^,^^48^, while connections to the Carpathian region are clear at the beginning of LH I, linked to horse equipment and the possible arrival of chariotry and steppe horses^39,49,50^. The results obtained by the project provide a significant set of new data to be contrasted to the existing knowledge of the archaeological record.

#### **1.5. Italy**

Present day Italy is a long country stretching from the Alps in the north to the middle of the Mediterranean Sea in the south. It is crossed north-south by the Apennine Mountain range, which divides the country in a Tyrrhenian and an Adriatic side, considerably affecting communication, and contributing to cultural and economic differentiation. The sea, due to c. 8000 km of coastline, has dominated life and possibilities for communication with the outside world throughout history.

In archaeological literature, northern/continental, peninsular, and insular Italy are often presented and considered as different territorial units^51^. In our project we included samples exclusively from Northern/continental Italy, and from the peninsular part of the country. Although differences occur and local regional chronologies have been developed and fine-tuned through the years, one could say that the Italian Bronze Age ranges from the end of the 3^rd^ millennium to the beginning of the 1^st^ millennium BCE (Table 4).

| **Period** | **Sub-Period** | **Absolute dates (BCE)** |
| --- | --- | --- |
| Early Bronze Age (EBA) | - | 2150-1650 BCE |
| Middle Bronze Age (MBA) | MBA 1 | 1650-1550 BCE |
|  | MBA 2A | 1550-1500 BCE |
|  | MBA 2B | 1500-1450 BCE |
|  | MBA 3A | 1450-1400 BCE |
|  | MBA 3B | 1400-1325/1300 BCE |
| Recent Bronze Age (RBA) | RBA 1 | 1325/1300-1225/1200 BCE |
|  | RBA 2 | 1225/1200-1150/1125 BCE |
| Final Bronze Age (FBA) | - | 1150/1125-925/900 BCE |

Table 4. Main chronological phase of the Italian Bronze Age^1^.

##### *1.5.1. Northern and continental Italy*

Northern Italy is that part of the country that goes from the Alps in the north to the Padan plain in the south. It is a highly connected area, crossed by an imposing water system that brings water from the Alps and from the Northern Apennines (Tosco-Emilian Apennine range) down to the Padan plain, the Po River, and finally to the Adriatic Sea. The Po River is a determinant feature representing a geographical, cultural, economic, and political border between the two sides of the plain. The Northern Italian individuals sampled in this project are all from the territories north of the Po and come from the north easternmost region of Friuli-Venezia Giulia (Mereto, Sedegliano, Selvis, S. Osvaldo) and from the Padan plain (Lucone, Narde, Olmo di Nogara, Scalvinetto).

Three of the four individuals sampled from Friuli Venezia Giulia region in north-eastern Italy (Mereto, Selvis, and S.Osvaldo, see Catalogue of the archaeological sites) were buried in EBA (see Table 4) monumental barrows. During the Bronze Age barrows or tumuli appeared across Europe^52^. North-eastern Italy is a region where an unusual concentration of such tumuli is to be found^53^. They typically contain a single central grave, where individuals with a special social or cultural position were deposited; the ritual activities that took place along with the construction of those burial monuments suggest the importance of those individuals^53^. The careful excavation at Mereto^54^ unveiled a particularly complex monument, grown progressively above the tomb, with a large stone platform in the beginning where multiple ritual activity took place over a long time to be finally covered by a mound by the MBA.

The sample from Lucone in the Lombardy region (see § 2.7.3) is roughly contemporary to those from Friuli, but comes from a completely different archaeological and cultural context. The sample is from a child whose remains seem to have been offered and thus belong to what has been interpreted as a ritual deposition. The remains were found in a settlement context and more specifically in a typical south Alpine pile dwelling; such settlements are widely spread across the whole Alpine area^55^ and have a complex historical trajectory and undergo a period of decline and abandonment at the onset of the MBA (see Tab. 4). It is suggested that a consistent part of the Alpine population from those sites moved down to the Padan plain and contributed to the rise of the so-called Terramare culture^51,56^.

The Terramare is the name given to the Bronze Age populations of the Padan plain^57^. They are characterised by dense settlement patterns, agro-pastoral economic base, elaborated handcraft, intense textile production, cultural and commercial contacts with both neighbouring and distant populations. A fundamental difference as to burial practices distinguish the Terramare south of the Po River and west of the Mincio, and those north of the Po and between the Mincio and the Adige rivers. The former groups practise a crematory burial ritual, genetic information about those populations is therefore completely absent from our records; the latter, living north of the Po, practise a mixed burial ritual and their cemeteries include both inhumation and cremation graves^58^. In our project we provide a deep close-up on one such community from the modern Veneto region, Olmo di Nogara^59^, complemented by three individuals from a nearby cemetery at Fondo Paviani/Scalvinetto^60,61^.

Scholars consider that the imposing demographic growth observable on the plain from the MBA 1 (Table 4), is the likely result of a consistent process of colonisation of the plain^58^. It is also suggested that some of the populations that might have moved into northern Italy would come from the cremation-practising tell communities of the Hungarian plain^62^ and from the Alpine lake dwellings (see above). The large number of individuals sampled from the necropolis of Olmo di Nogara and the few individuals from Scalvinetto provide for the first time the possibility to test existing hypotheses based on archaeological evidence using palaeogenomic data. Terramare communities see a rather abrupt ending at the end of the RBA (see Table 4). The reasons for such a dramatic transformation are widely debated^1^, among our samples we have individuals from the FBA (see Table 4) cemetery at Narde. Narde is one of the two large cemeteries of the impressive settlement at Frattesina di Fratta Polesine, which blossom after the end of the Terramare at the edge of the Padan plain close to the Adriatic coast. Conspicuous archaeological evidence of craft production and large-scale exchange has been detected at Frattesina. The dominating burial ritual at Frattesina is cremation, but a handful of inhumations exist; their limited number suggest that those individuals are probably not representative for the whole community, nonetheless they give the opportunity. at least to some extent, to discuss the local population after the crisis at the end of the Terramare period.

##### *1.5.2. Peninsular Italy*

Our samples from peninsular Italy are from very diverse sites. The total number of samples from this region is limited, providing just a glimpse of what can possibly have been the underlying genetic complexity of the region.

For a full understanding of the results it is important to consider that the individuals from Pian Sultano, mid-Tyrrhenian Italy (modern Latium region, see § 2.7.7) belong to the same chronological horizon as the northeastern individuals from Friuli-Venezia Giulia and that from the lake dwelling at Lucone (see above).

From the southern-easternmost part of the Italian Peninsula we have sampled individuals from Toppo Daguzzo, tomb 3 layer 2, and Lavello, tomb 743 layer 6, both dated to the MBA 1 (1650- 1550 BCE) and those from Coppa Nevigata dated to the second half of the 2^nd^ millennium BCE (c. 1500-1200 BCE). This area of the peninsula is characterised during the Bronze Age by specific cultural traits and at the same a long tradition of contacts with the Adriatic, the Aegean, and the Maltese world^63^. Additionally starting approximately, from the 13^th^ century BCE a local production of the so-called Italo-Mycenaean pottery is well attested in this part of the Italian Peninsula and it is generally hypothesised that Aegean potters moved and established their own ateliers in southern Italy^64^.

##### *1.5.3 Conclusive reflections*

All in all, the individuals sampled in this project allow approaching with new data the complex archaeological and cultural diversity evident across the Italian Peninsula during the Bronze Age. Additionally, we are also able to discuss in detail the internal dynamic of one single community (Olmo di Nogara), strategically positioned between the Padan plain and the Prealps, thus with access not only to the rich fertile landscape of the plain, but likely also to the abundant Alpine copper ores^65^.

#### **1.6. Spain**

All our samples from the Iberian Peninsula come from modern day Spain. Our samples come from different regions: Catalonia, in the north-east, Andalusia in the south-east, and Castilla y la Mancha in central Spain. Despite only a limited number of the numerous samples that were initially collected provided genetic information, our dataset is very interesting as it includes several politically and culturally prominent archaeological contexts dated to distinct periods, from the 3^rd^ and the 2^nd^ millennium BCE. Interestingly, most of our samples come from collective graves but we have not been able to detect significant kin-relation among the analysed Iberian individuals from the same contexts. Probably, a higher number of individuals would be necessary to have information about kinship; for the time being, data suggests that collective graves might have not responded to kin-related necessities rather they might have represented other social constellations or bonds.

##### *1.6.1. Catalonia, north-eastern Spain*

The three archaeological sites from Catalonia provide a long-term perspective on the population of the region as our samples are dated to different periods: the individuals from the burial cave of Costa de Can Martorell fall approximately within the second half of the 3^rd^ millennium BCE. The samples from the settlement at Cantorella are dated to c. 1880-1680 BCE; while those from the settlement at Miniferri are dated to the first half of the 2^nd^ millennium BCE (c. 2130-1450 BCE).

Costa de Can Martorell is a large and complex Neolithic hypogeum grave, whose characteristics suggest that at least part of the deceased were buried there in the aftermath of one single episode of violence (see § 2.10.2). The settlements at Cantorella and Miniferri on the other hand, come from the Early Bronze Age phases and consist of smaller groups of individuals buried in silos from the Neolithic period, which are typically reused as graves during the Bronze Age.

##### *1.6.2. Andalusia, south-eastern Spain*

Andalusia is a vast region in southern Spain. The warm climate of the region did likely compromise the preservation of the DNA and despite the large number of sites included in our dataset only a few individuals per each site could be studied. Most of our samples from Andalusia are dated to the Argaric period (c. 2000-1600 BCE), with the notable exception of the burials from the prominent Chalcolitic site of Valencina de la Concepción dated approximately between 3200 and 2400 BCE and the late Chalcolitic/Early Bronze Age cemetery at Los Algarbes (c. 2500-2000 BCE).

To fully understand the results of this project, it is important to consider that Valencina de la Concepción is the largest Copper Age settlement not only in the Iberian Peninsula, but also in Western and Central Europe covering an area of approximately 200 ha.

The cemetery de los Algarbes has been interpreted as a funerary site likely functioning as a sort of territorial central place for different groups burying their members together in the same place. The site is dated from Iberian Late Chalcolithic to the Early Bronze Age (Table 1) and the individuals analysed in this study fall into Early Bronze Age phases of the site. All the remaining individuals from southern Iberia are also from the Early Bronze Age and provide the possibility to closely study the development of the local population during a time that is also known as the Argaric period (c. 2200-1500 BCE).

##### *1.6.3. Castilla y La Mancha, central Spain*

Our dataset from central Spain includes two prominent Bronze Age fortified settlements with intramural burials, dated to the first half of the 2^nd^ millennium BCE and thus contemporary to the individuals from the Argaric population in Andalusia.

##### *1.6.4. Conclusive reflections*

The individuals sampled from Iberian Peninsula provide the possibility to further develop earlier studies^66,67^ and to expand our understanding of the consistent transformations at the onset of the Early Bronze age when the distinctive Argaric culture rises in the southern part of the Iberian Peninsula.

### 2. CATALOGUE OF THE ARCHAEOLOGICAL SITES

#### **2.1 Introduction**

In this document we present critical information about the archaeological contexts where each of the individuals analysed in the paper was found. The vast majority of the samples come from regular excavations and abundant data about the contexts of provenience and their chronology are available. All sample providers have been committed to produce abundant bibliographical references for further reading.

The analysis of the archaeological contexts is arranged geographically per modern country of provenance. The countries where the samples have been excavated are listed in alphabetical order. Within each nation the various sites are also listed in alphabetical order.

The description of each site is preceded by technical information such as:

- author of the text^[1]^
- geographical coordinates
- the name of the person or the institution that kindly provided the samples
- when possible, a short general description of site is provided before the detailed description of the single contexts from the site
- When available the radiocarbon dating from the sites or the analysed samples is given.

The specific context or grave where the single samples come from, are thereafter analysed in detail. Samples are identified in the text by their laboratory number (CGG number) to allow cross-referencing with the rest of the supplementary material and the main paper. When existing, the RISE nr is also provided. The RISE nr is the number given to the samples by the other researchers of the so-called Rise II project (<https://www.gu.se/forskning/the-rise-ii-en-ny-europeisk-forhistoria-att-integrera-dna-isotoper-sprakhistoria-och-arkeologi-for-en-nytolkning-av-grundlaggande-forandringar-i>) that co-funded the research along with Lundbeck Foundation GeoGenetics Centre in Copenhagen (<https://globe.ku.dk/research/lundbeck-foundation-geogenetics-centre/>). The RISE nr has been used by the facilities of the National Museum in Copenhagen to identify samples that underwent strontium isotope analyses. The RISE nr might therefore help to cross-reference the samples discussed here in future publications as well.

We are deeply grateful to all sample providers and the authors of the various texts for their collaboration.

#### **2.2. Cyprus**

Please note, much of the information provided below about the various Cypriot contexts can be consulted in the online database (Carlotta database) of the Swedish Museum of Mediterranean and Near Eastern Antiquities, where the sampled material is currently preserved. Carlotta database can be consulted at: <https://collections.smvk.se/carlotta-mhm/web>

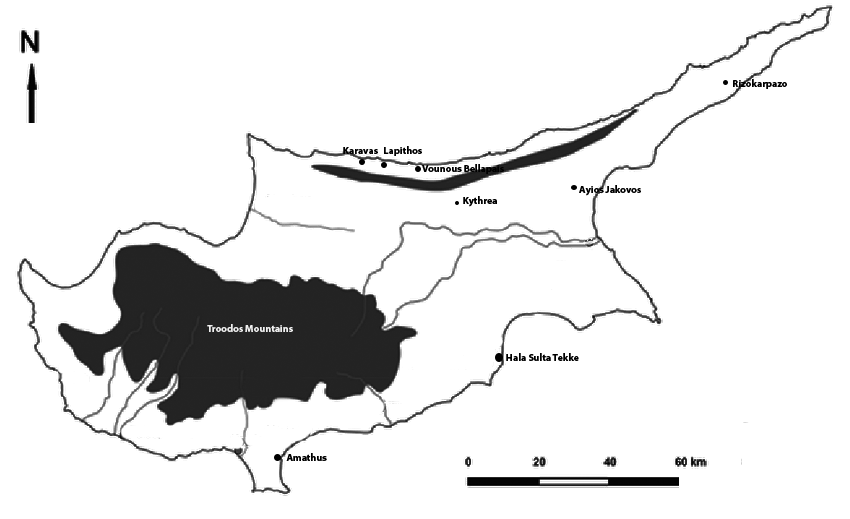

Map 1. Map of Cyprus with the sites sampled for this project (© authors)

##### ***2.1.1. Agios Iakovos necropolis, (de facto) İskele District, Northern Cyprus***

The exact Latitude and Longitude of the site is missing, but the modern village of Ayιος Iάκωβος/Altinova has Latitude: 35.315892006041665 and Longitude: 33.82316519397129.

Sample provider: Christian Mühlenbock, Mia Broné

Serena Sabatini

The cemetery at Agios Iakovos (published by Einar Gjerstad and his collaborators as Aijos Jakovos, or the Necropolis of Melia)^68^ is situated c. 1.5 km east of the village of Agios Iakovos (today Ayιος Iάκωβος/Altinova) and 1.8 km south of the village of Mandres (today Mάντες/Ağillar) at the foot of the Kyrenia Mountains (see Map 1). The closest major town is Famagusta c. 22.5 km to the south. The Swedish Cyprus Expedition worked at the site for seven weeks starting in late June 1929. They excavated 14 graves (Fig. 1). All graves are chamber tombs dug out in the local sedimentary limestone.

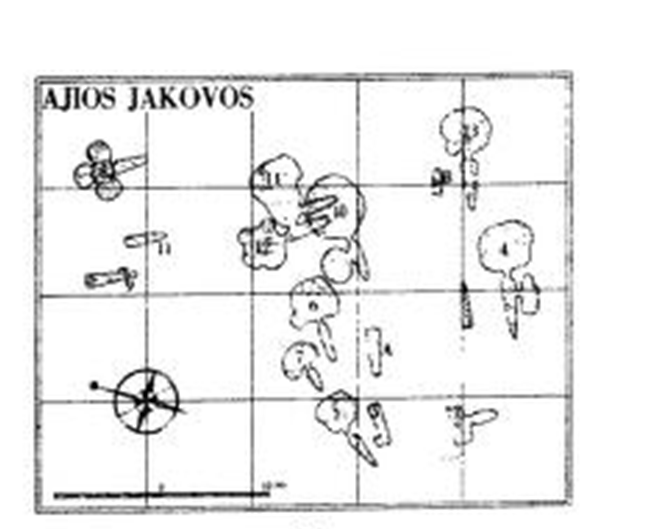

Fig. 1. Plan of the necropolis at Agios Iakovos^69^.

***Agios Iakovos Tomb 8***

Tomb 8^68^ is maybe the most impressive of the excavated tombs, in the sense that it has a central pit and three large lateral chambers (Fig. 2), while the other tombs from the same cemetery consist of only one main chamber and eventually a second smaller one. The grave was well preserved and excavated in its entirety. The excavators could distinguish several layers and three main burial periods. To the oldest deposition (Layer 8) belong 9 individuals (IV-V, VII-IX and XIX-XXII), which were poorly preserved. Judging from the grave goods this layer was dated to the Middle Cypriot III (c. 1750-1650 BCE). When the tomb was to be used for a second deposition phase, the human remains from these 9 individuals were swept down to the bottom of the main chamber together with their grave goods. They were covered with a layer of white clay.

The second deposition phase appeared more complex (maybe more sub-phases); the excavators collected the rests of (they believed) 35 individuals (I-III, VI, XIII-XVIII, XXIII-XXV, XXVII-XXVIII, XXXIII-XXXV, XXXVIII-XLIII, XLV-XLIX, LII-LVII) and very little grave goods. To this second phase belong all the individuals sampled in this project (individual II [**CGG_2_015634/Rise 614]**; individual XXIIIa [**CGG_2_015631/Rise 611**]; individual XXIIIb [**CGG_2_015630/Rise 610**]). The excavators considered this second deposition layer as a sort of mass-grave. This second burial phase was dated archaeologically to the Late Cypriot I (c. 1650-1450 BCE). Cranium XXIIIa has been 14C dated to 3235±15 (cal.BP to 3481-3396, see Suppl. Tab. S9), which confirms the archaeological chronology proposed by the excavators.

It is important to mention that two of the analysed samples (**CGG_2_015630** and **CGG_2_015631**) belong to individuals that have not been recognized during the excavations^68^; the excavators collected only one individual XXIII; later on in the laboratory of the Medelhavsmuseet in Stockholm, they realised that the box with the individual XXIII actually contained the rest from three different skeletons which were therefore numbered as XXIIIa, XXIIIb and XXIIIc (Carlotta database). XXIIIa (**CGG_2_015630**) and XXIIIb (**CGG_2_015631**) were sampled in this project. There is no reason to doubt that all three individuals (XXIIIa-c) belonged to the same second depositional layer of the grave (it seems to us that the excavators just collected the bone fragments underestimating the number of skeletons; thus, the depositions from the second layer should be changed to 37 bodies). According to a note found with Cranium XXIIIa, the latter was excavated “*on 16/7/1929, from layer 71*”, this information might be the same also for XXIIIb and XXIIIc, but we haven’t been able to find any further information about layer 71.

The grave contained a third and last burial phase that was better preserved than the previous ones. The grave was in fact probably closed and left undisturbed once the last body from phase III was buried. The skeletons belonging to this last phase (X-XII, XXVI, XXIX-XXXII, XXXVI-XXXVII, XLIV, L-LI, LVIII-LIX, LXII) were apparently in situ, lying in a outstretched dorsal position. No samples were taken from this last phase.

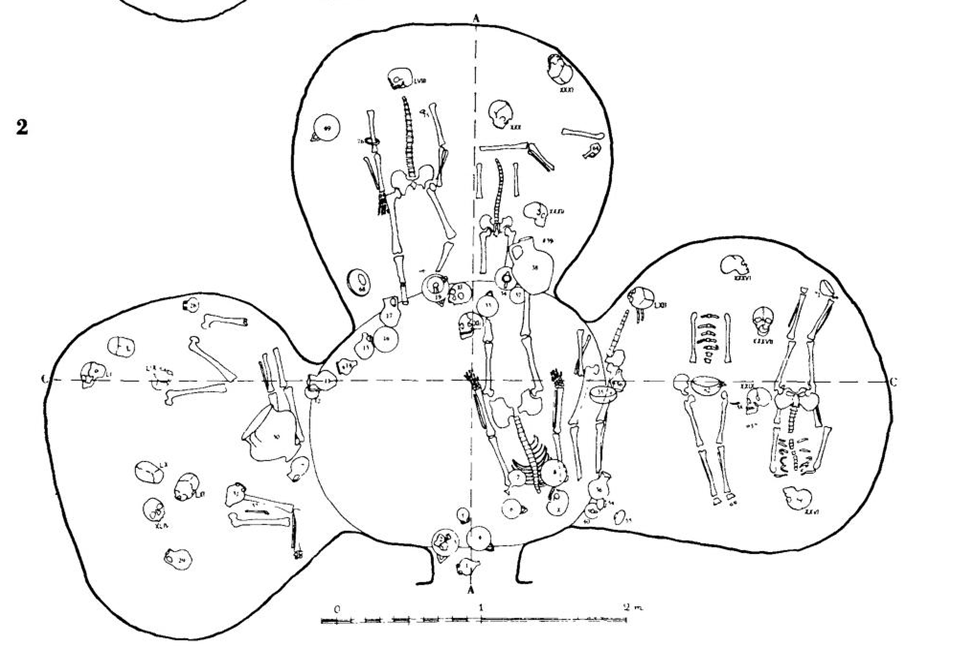

Fig. 2. Agios Iakovos, plan of phase II in tomb 8, fig. 127.2^69^

***Aijos Iakovos Tomb 14***

Tomb 14^68^ has an unusual shape with a relatively large niche on the eastern side of the elongated chamber. The entrance to the chamber is not perpendicular to the main axis of the chamber, it opens laterally and in correspondence with the narrowest part of the chamber. This most likely happened because the grave was built very close to two earlier chamber tombs (10 and 12) and had to literally squeeze in between them. Why it was decided to build the grave in this limited space cannot be understood. The grave was well preserved and had two clear burial phases separated by artificial filling strata. The earlier burial stratum (6) contained 35 individuals. 25 of those were practically still in situ, deposited in a dorsal outstretched position or in a slightly lateral position. The bodies were put on top of each other and the excavators suggested it was a sort of mass grave. The second more recent burial phase had only five skeletons, deposited one in each side of the chamber.

The first burial phase is dated to the late Middle Cypriot III-early Late Cypriot IA (c. 1700-1550 BCE), while the second burial phase is dated to the Late Cypriot IIA (c. 1450-1375 BCE).

According to the archive of the Medelhavsmuseet in Stockholm (Carlotta database) Cranium XX (**CGG_2_015633/Rise 613**) belongs (osteological observations) to an adult, possibly a male, which is confirmed by the genetic sex. It is not possible to say whether the skeleton belonged to the first mass deposition or to the later one. If one considers the number of the cranium, it is likely that it was part of the earlier deposition phase with 35 individuals. A second individual from this grave was also sampled (Cranium IIX(sic) **CGG_2_015632/Rise 612**). The number of crania is obviously not correct, but it is not possible to understand the original one (cf. Carlotta database); it is also not possible to say whether the skeleton belonged to the first or the second layer.

##### ***2.2.2. Amathus necropolis, Limassol district***

Latitude: 34.71037010782082; Longitude: 33.1424284473686

Sample provider: Christian Mühlenbock, Mia Broné

Serena Sabatini

The site of Amathus lies on a hill on the south coast of Cyprus, about 10 km east of Limassol on the road to Nicosia (see Map 1). It was excavated during April/May 1930^68^ and 26 rock-cut and/or built shaft graves with a dromos leading to the chamber were excavated (Fig. 3). The samples presented in this study come from tomb 11 (**CGG_2_022546/Rise 1689, CGG_2_022547/Rise 1690**), tomb 21 (**CGG_2_022548/Rise 1691, CGG_2_022549/Rise 1692**), and tomb 23 ( **CGG_2_022551/Rise 1694, CGG_2_022552/1695**).

The shape of the Amathus graves is somehow unusual and it has been discussed whether there is a connection between this unusual shape and the so called Eteo-Cypriots. The term “Eteo-Cypriot” is largely disputed, but it is used to define a Cypriot language written in alphabetic script, which might be connected to the original language still spoken by the autochthonous inhabitants of Cyprus after the Hellenization of the island at least in some places. Amathus, is the city from where almost the totality of the “Eteo-Cypriot'' corpus of written documents is known so far^70^. Nonetheless, it has been also pointed out how the local stone is of a very brittle quality so that the usual Cypriot chamber graves would not be possible to build at Amathus. As a matter of fact the architecture of the local tombs could then just be enforced by the local geology^71^.

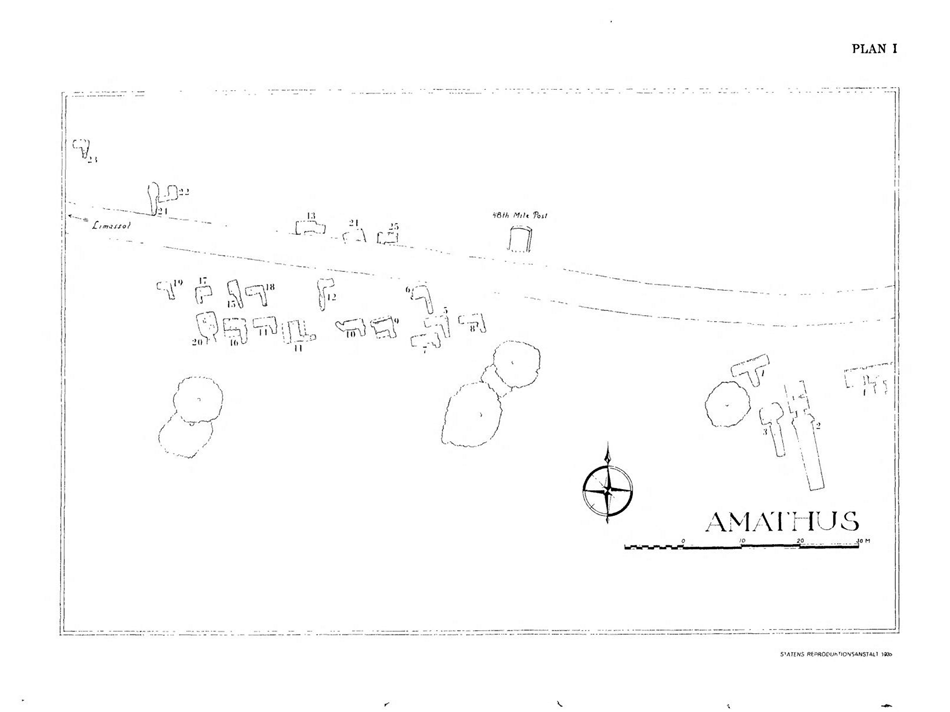

Fig. 3 Plan of the cemetery at Amathus, Cyprus, Plan 1^68^.

***Amathus, Tomb 11***

The tomb is a large shaft grave, which revealed several depositional events^68^. The oldest burial at the bottom was probably one male individual. Above him in the second layer two more individuals -a male and a female- were buried. These first two burial events have been dated by the excavators to the Cypro-geometric III and the Cypro-Archaic I periods (c. 900-600 BCE). Above them, in the most recent stratum there were 3 more depositions. Cranium 1 (**CGG_2_022546/Rise 1689**) and cranium 2 (**CGG_2_022547/Rise 1690**) sampled by the project, belong to this most recent level. The remains were apparently found along the eastern wall and had no clear connection to any grave goods; for stratigraphical reasons they have been dated around the end of the Cypro-Archaic II period (c. 600-475 BCE).

Cranium 1 (**CGG_2_022546/Rise 1689**) has been 14C dated to 2580±15 and 2σ cal. to 786-773 BCE. Such chronology does not correspond to the one proposed by the excavators on the basis of the stratigraphy. It rather suggests that the upper and most recent stratum belongs to a central phase of the Cypro Archaic I period, like the other two earlier phases; the grave was then probably used during a relatively short period of time and the various layers possibly were just accurately separated and prepared for the new depositions.

According to the osteological observations available in the Medelhavsmuseet database (Carlotta database), both crania belong to adult individuals. For both of them, the genetic sex results to be male.

***Amathus, Tomb 21***

Tomb 21^68^ was of considerable size when compared to the other graves from the same cemetery. It was only partly excavated since its front and central parts lie under the modern road that cut the cemetery in two parts. Only the innermost part of the grave could therefore be investigated. The sides of the shaft were lined with walls of ashlar. It seems that the interior part of the tomb was used to push back the earlier depositions, when it was time to put new individuals. This might explain why no entire skeletons were found and why the lowest stratum was the one where bones were least well-preserved and most mixed up. The different burial layers seemed otherwise separated by pebble floors. Two crania came from layer 2 (right above the oldest deposition layer). Each cranium had been placed inside a large amphora together with other scattered human bones. Both crania have been sampled by our project (**CGG_2_022548/Rise 1691** and **CGG_2_022549/Rise 1692**). There was a third quite disturbed deposition layer of much later date (Hellenistic/Roman).

Layer 2 (from which our samples come from) was dated by the excavators to the end of the Cypro-Geometric II period (c. 950-900 BCE). However, and quite surprisingly, both samples have been radiocarbon dated to an earlier date than the Cypro-Geometric II.

**CGG_2_022549/Rise 1692** is now dated to 2965±15BP, cal. BP 3315-3075 (see Suppl. Tab. S9) and **CGG_2_022548/Rise 1691** is now dated to 2990±20BP, cal. BP 3208-3070 (see Suppl. Tab. S9). Generally the earliest dates for the cemetery at Amathus are considered the beginning of the Cypro-Geometric period (c. 1050 BCE); the results of our 14C analyses show that this grave contained individuals dated to Late Cypriot II-III. Therefore it was likely used for a longer time period than expected or (since the two crania were found inside an amphora) it is possible that the sampled individual were re-buried in this tomb under specific circumstances from elsewhere.

According to the osteological observations available in the Medelhavsmuseet database (Carlotta database), both crania belong to adult individuals and cranium 2 (**CGG_2_022549/Rise 1692**) possibly to a female individual, now confirmed by the genetic sex.

***Amathus, Tomb 23***

Tomb 23 was a shaft grave of rectangular shape. It revealed several deposition layers^68^. The pottery and the other finds from the grave date all the layers to the Cypro-Archaic I period (c. 750-600 BCE). As the excavators pointed out^68^, the stratigraphy was very clear and the grave had 4 different depositional phases, all very close in time.

The samples presented in this study come from Cranium 11 (**CGG_2_022552/Rise 1695**) and Cranium 15 (**CGG_2_022551/Rise 1694**).

There was at least one deposition in the first and oldest burial layer, but no samples were taken from it. Cranium 15 (**CGG_2_022551/Rise 1694**) comes from the second burial layer and was found close to another skull (Cranium 14), which likely fell from a body in a sitting position from the corner of the shaft. It is not clear whether any other individual was also deposited in this second layer. The layer was archaeologically dated by the excavators to an early phase of the Cypro-Archaic I period (maybe c. 750-700 BCE).

In the third burial layer at least two individuals were identified, but no samples were taken from it. In the fourth and last burial layer, which was dated archaeologically to the end of the Cypro-Archaic I period (maybe c. 650-600 BCE?), there were 11 skulls (crania 1-11). None of the bodies was in situ and all the bones were mixed together. Cranium 11 (**CGG_2_022552/Rise 1695**) comes from this last phase. No 14C analyses were run on these samples.

According to the osteological observations available in the Medelhavsmuseet database (Carlotta database), both cranium 11 and 15 belong to adult individuals.

##### ***2.2.3. Karavas, (de facto) Kyrenia district, Northern Cyprus***

The exact Latitude and Longitude of the site is missing; however the modern village of Karavas, today Καραβάς/*Alsancak* (see Map 1) has Latitude: 35.347222 and Longitude: 33.207778.

Sample provider: Christian Mühlenbock, Mia Broné

Serena Sabatini

**Karavas, Tomb 1**

According to the digital archive of Medelhavsmuseet (Carlotta database) the tomb was excavated by P. Dikaios in 1933-1934^72^. The antiquarian of the Cyprus Museum in Nicosia. P. Dikaios and the French archaeologist C.F.A Schaeffer, made some inquiries to C.-M. Fürst in Sweden concerning osteological analysis and the material was thereafter sent to Sweden. Since Fürst passed away before further investigations, the material was analysed by C.-H. Hjortsjö^73^. Hjortsjö^73^ reports a description of the site at Karavas. It is clearly a Neolithic settlement at c. 1 mile from the northern coast of Cyprus. The pits were discovered within the settlement area and the situation looked very much like the one that Dikaios had already excavated at Erimi.

According to Fischer^74^ it is written by Dikaios and Stewart^75^ that the skeleton was found in one of the burial pits. The diameter of the pit varied from 0.6 to 1 m, while the depth was approximately constant at 1 m. The skeleton was in the doubled-up position and stag antlers were found under the bones. The context is dated archaeologically to the Chalcolitic period I (c. 3000-2500 BCE).

According to Hjortsjö^73^ the skull from the sample (**CGG_2_022531/Rise 1674**) belongs to an adult woman.

##### ***2.2.4. Kythrea, (de facto) Lefkoşa district, Northern Cyprus***

The exact Latitude and Longitude of the site is missing. The modern village of Kythrea is at Latitude 35.250000 and Longitude 33.483333.

Sample provider: Christian Mühlenbock, Mia Broné

Serena Sabatini

Kythrea (Turkish Değirmenlik) lies c. 10 km northwest of Nicosia (see Map 1). It is the site of a Neolithic settlement excavated by the Swedish Cyprus Expedition in 1930^68^. In the excavation report nothing is written about possible graves. Given that burial pits have been found in Neolithic settlements at Karavas and Erimi, it is not impossible that human bones were collected also here, but not recognized as such. It is in any case not possible to know whether the find comes from the Swedish Cyprus Expedition campaigns or from other excavations in the area. According to the digital archive of Medelhavsmuseet (Carlotta database) on the note found together with the osteological material is written: “M1-2, ytan [*ytan is the Swedish word for surface*] -0,55”. 1B was added by the museum intendant”. It is clear from the report of the excavation that the Neolithic settlement at Kythrea was documented within a grid where each square is named with a letter and a number (298 plan XII 1-2)^69^; our sample (**CGG_2_022543/Rise 1686**) could therefore possibly come from squares M1-2, which would be from the northeastern corner of hut III. We were not able to confirm this hypothesis; however the settlement at Kythrea was dated by Gjerstad to the end of the 4^th^ millennium BCE

##### ***2.2.5. Lapithos necropoleis, (de facto) Kyrenia district, Northern Cyprus***

The exact Latitude and Longitude of the site is missing. The modern village of Lapta is at Latitude: 35.336667 and Longitude: 33.174167.

Sample provider: Christian Mühlenbock, Mia Broné

Serena Sabatini

The site of Lapithos is located close to the village with the same name on the north coast of Cyprus, today the name of the village is Λάπηθος/*Lapta* (see Map 1)*.* The area is very rich in archaeological sites and has been repeatedly excavated by several missions. Lapithos is located strategically in many ways and it has been considered as one of the main harbours for the Middle Bronze Age trade in copper^41^. The wealth of the grave goods in some of the Middle Bronze Age tombs indicate that there was a rich community living there.

The Swedish Cyprus Expedition excavated a settlement there during the months of October and November 1928^69^. The settlement was dated by the finds to the 4th millennium BCE. In the same area where the Neolithic settlement is located, a small group of later tombs (dated to the Cypro-Geometric I and II, c. 1050-900 BCE) were individuated and excavated, those tombs are known as coming from the cemetery of ***Plakes*** (Fig. 4). Along the shore west of the modern village at a locality called ***Vrysi tou Barba*** (Fig. 4) 23 chamber tombs were excavated in 1927 (September-November), the known cemetery today is bigger as many more tombs have been excavated there in 1913, 1917 and also after the work of the Swedish-Cyprus today^76^. Most of the tombs from Vrysi you Barba were richly furnished and dated between the Early Cypriot II and Middle Cypriot III (c. to 2150-1650 BCE). It is possible that the Early/Middle Cypriot Lapithos community (*Vrysi tou Barba*) gained its political and economic importance thanks to aggressive politics against neighbouring sites^77^. From the end of 1927 until spring 1928, in the locality of ***Kastros*** (Fig. 4), another cemetery was excavated revealing 30 tombs of different types. Also here some of the contexts were richly furnished. The Kastros necropolis was dated to the Cypro-Geometric period (c. 1050-750 BCE) and were therefore roughly contemporary to those from ***Plakes***.

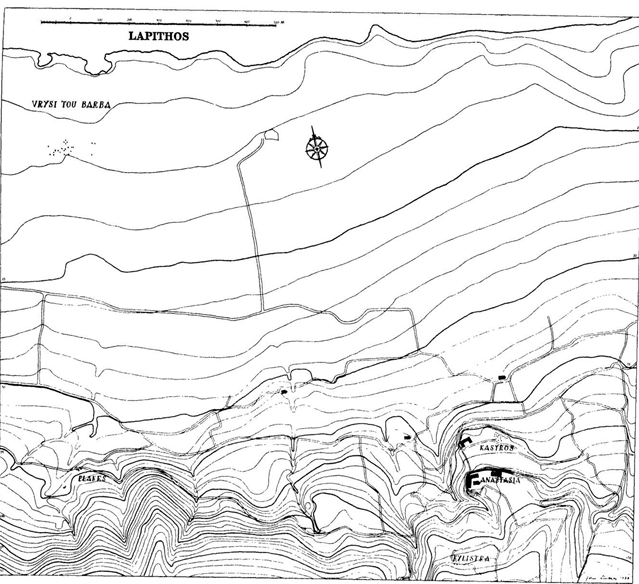

Fig. 4 The area of Lapithos, with the three cemeteries of Kastros, Plakes and Vrysi tou Barba (from Gjerstad 1934, Pl.III.1)^69^.

###### Lapithos, Vrysi tou Barba cemetery

**Lapithos Vrysi tou Barba, Tomb 315.**

Tomb 315 is a large tomb and it has 3 chambers^68^. The tomb was heavily disturbed by modern work, so that a proper understanding of the internal stratigraphy and the characteristics of the burial became impossible. In chamber A the rest of at least six individuals were found. Cranium V (**CGG_2_022497/Rise 1640**) is to be considered part of this group. In chambers B-C modern water and mud infiltration has created much disorder and it is not clear how many depositions there were. Gjerstad et al. (1934, 108)^69^ could see at least three depositions, but the samples from this chambers called 315c/4 (**CGG_2_022502/Rise 1645**) suggest that there might have been at least a fourth skull comes from this chamber. None of the skeletons excavated in grave 315 was found in situ or in its original position.

Judging from the finds that could be retrieved in the different chambers the tomb was probably cut at the beginning of Middle Cypriot I (c. 2000-1850 BCE) and used until the beginning of the Middle Cypriot II period (c. 1850-1750 BCE).

According to the catalogue of the Medelhavsmuseet in Stockholm (Carlotta database), **CGG_2_022497/Rise 1640** belongs to an adult individual, possibly male; according to Fischer (1986, 30) this individual is a 30 years old male. Also the genetic sex resulted to be male According to the Carlotta database **CGG_2_022502/Rise 1645** is an adult individual.

**Lapithos Vrysi tou Barba, Tomb 322.**

Tombe 322^68^ is exceptionally large with 5 chambers. According to the excavators the exceptionality of the context can be explained by the shape of the dromos. They believed it could have been two different tombs, whose dromoi met and became one. Additionally, J. Webb's^78^ essay on this specific grave shows that there is overlooked data/material and that it would deserve renewed attention. All in all the tomb is usual and impressive in many ways. The digital archive of Medelhavsmuseet in Stockholm, says that the sample comes from “*Tomb 322B; Cranium: I. The present skeleton designation is in agreement with the numbers used by Fürst (1933)*”. It is therefore likely that **CGG_2_022496/Rise 1639** belongs to individual I from Chamber B. The excavators write that the tomb and in particular chamber A and B had a magnificent construction. The tomb was richly furnished with unusually many bronzes and also gold and a lump of iron. There was evidence for impressive ceremonies taking place at the burial as many animal bones and other material suggest.

From the chronological point of view the grave was in use from the Early Cypriot III to the Middle Cypriot I (c. 2100-1850 BCE). Chamber B was, according to the excavators^69^ the oldest chamber.

According to the catalogue of the Medelhavsmuseet in Stockholm (Carlotta database), **CGG_2_022496/Rise 1639** belongs to an adult individual, possibly female; according to Fischer (1986, 30) this individual is a 18-24 years old female.

**Lapithos Vrysi tou Barba, Tomb 323.**

The sample (**CGG_2_022488/Rise 1631**) from this grave is -according to Medelhavsmuseet’s archive (Carlotta database)- from grave 23; however the initial numbers 1-23 given by the excavators of the Swedish Cyprus Expedition to the graves they investigated in 1927 were changed prior to publication to numbers 301-323 to be sure to distinguish those graves from the many others already excavated and that were left to excavate. Grave 323^69^ is a large grave with four chambers (A-D). Human remains were badly preserved and apparently only in Chamber C. The documentation about this context is somewhat contradictory as in the publication the authors mention long bones, but not skulls; however skull bones might have just been in fragmentary conditions and thus not worth mentioning.

The finds from this grave date the tombs as a whole between the Early Cypriot II and the Early Cypriot III period (c. 2150-1950 BCE). According to the catalogue of the Medelhavsmuseet in Stockholm (Carlotta database), the sample analysed in this project **(CGG_2_022488)** belongs to an adult individual.

###### Lapithos, Kastros cemetery

**Lapithos Kastros, Tomb 403**.

Tomb 403^69^ was well preserved and showed the presence of three burial layers. A total of eight individuals were buried there. To the earliest burial layer belonged skeletons V-VII. This burial phase was dated to the beginning of the Cypro-Geometric III (approximately 900-850 BCE). Skeletons I, III and VIII belonged to the second burial phase, which was dated to a later stage of the first part of the Cypro-Geometric III (c. 850-800 BCE). Skeletons II and IV were the last two depositions dated to the mid Cypro-Geometric III (around approximately 800 BCE). We sampled skeleton VIII (**CGG_2_022516/Rise 1659**) from the second burial phase and skeleton II (**CGG_2_022510/Rise 1653**) from the third and last phase.

It is interesting to note that the grave was richly furnished and that several of the skeletons were adorned with golden jewellery. According to Fischer^74^ skeleton VIII could belong to a female individual of 18 years; also the genetic sex resulted to be female.

**Lapithos Kastros, Tomb 406**.

Tomb 406^69^ is a typical shaft grave as others from the site. It contained three individuals and no evident stratigraphy. Considering that the grave goods are very homogeneous, the excavators suggest that probably a very short time, if any, passed between the three depositions. The finds date the tomb to an early part of the Cypro-Geometric I (c. 1050-1000 BCE). According to the database of the Medelhavsmuseet in Stockholm (Carlotta database), both samples (**CGG_2_022517/Rise 1660** and **CGG_2_022523/Rise 1666**) belong to adult individuals. The skull of our sample **Rise 1660** was analysed also by P. Fischer^74^.

Both samples from grave 406 were radiocarbon dated to 2935±15BP (cal. BP 3162-3004) and 2990±15BP (cal. BP 3229-3076), respectively (see Suppl. Tab. S9). The calibration suggests a slight difference between the archaeological chronology and radiocarbon dates, possibly due to dietary habits.

**Lapithos Kastros, Tomb 408**.

During the excavation in Tomb 408^69^ not all the finds appeared in situ. However there was no complex stratigraphy in the tomb, so the excavator concluded that it had only one main burial phase and that the material moved due to soil and water infiltrations through the years. Three bodies came from this tomb, two from the main chamber and one from the niche carved on the side of the chamber. The two bodies in the chamber were probably in an outstretched dorsal position, while the body in the niche was probably in a sitting position with the back lining on the wall. The characteristics of the grave goods suggest that the body in the niche might have been deposited some time after the first two in the main chamber. All the finds belong to the early part of the Cypro-Geometric II (c. 950-900 BCE). Looking at the drawing of the tomb^69^ it seems possible to state that cranium I (**CGG_2_022507/Rise 1650**) belongs to the individual that was put in a longitudinal position at the centre of the chamber with the head towards the entrance. This skeleton was furnished among other things with a golden ring.

According to the catalogue of the Medelhavsmuseet in Stockholm (Carlotta database), the individual sampled in this study is an adult, possibly a female. According to P. Fischer^74^ the bones of this same individual belong instead to a male of 36 years.

**Lapithos Kastros, Tomb 420**.

The way in which grave 420^69^ was found suggests that it had a single deposition phase. All the finds appeared more or less in situ. Inside the tomb there were four bodies, three adults and one child. All and in particular skeleton 1 were richly furnished with many pieces of golden jewellery. The Skeletons had been buried in a dorsal position with their head towards the entrance. Outside the grave in the dromos (access corridor) two skeletons also in dorsal position, and with their head towards the entrance were found without any grave good. The sample **CGG_2_022508/Rise 1651** seems to belong to one of these two individuals in the dromos. The excavators suggested that these could have been slaves sacrificed at the moment of the burial. The tomb is dated to an early part of the Cypro-Geometric I period (c. 1050-1000 BCE).

According to the catalogue of the Medelhavsmuseet in Stockholm (Carlotta database), the individual sampled in this study is an adult . According to P. Fischer (1986, 39) it was a male individual of 30 years.

**Lapithos Kastros, Tomb 421**.

Tomb 421^69^ is a shaft grave different in shape from those of the other tombs and rather small in size. Only two individuals were buried in this grave and in the same burial phase. The material dates the context to the end of the Cypro-Geometric III (c. 850-750 BCE). According to the catalogue of the Medelhavsmuseet in Stockholm (Carlotta database) it is likely that the bones sampled in this study (**CGG_2_022518/Rise 1661**) belong to individual II and that it was an adult individual. The genetic sex of the sample resulted to be female.

**Lapithos Kastros, Tomb 422**.

Tomb 422^69^ is a shaft grave with long dromos. According to the information collected from the digital archive of the Medelhavsmuseet (Carlotta database) this sample (**CGG_2_022520/Rise 1663**) comes from the dromos (access corridor) of the grave. This is very interesting, because the complex situation excavated in the dromos suggested to the excavators that human sacrifices were performed there; thus this sample likely belongs to an individual that has been killed ritually. The observations made by the excavators are precise and convincing^69^. The grave contained two burial phases, of the earliest one very little was left (bones and objects were crossed and swept away). This first layer was dated to an early part of the Cypro-Geometric I (c. 1050-1000 BCE). The second layer belonged to one individual. At the same stratigraphic level as this body, but outside the grave in the dromos, three individuals were found piled up one on top of the other. The position of their bones suggested that their hands and feet were tied. On top of the uppermost skeleton two stone idols were laid down; below the bottom individual there was a peculiarly carved stone which has been interpreted as a sacrificial table. From the available documentation it is not given to understand which of the three individuals has been sampled. This layer was dated by the find to the later part of the Cypro-Geometric III (c. 850-750 BCE) According to the archive of the Medelhavsmuseet (Carlotta database) the individual sampled in this study is an adult. The genetic sex of this sample resulted to be female.

**Lapithos Kastros, Tomb 423**.

Tomb 423^69^ was found looted and there were no intact burial remains nor many items left in it. The few finds suggested that the grave should have been dated to the end of the Cypro-Geometric II (c. 900 BCE). Nothing could be said about the number of individuals or deposition phases.

According to the digital archive of the Medelhavsmuseet in Stockholm (Carlotta database), the bones sampled in this study (**CGG_2_022515/Rise 1658**) belongs to an adult, possibly a male individual, also Fischer^74^ considers these bones as belonging to a male individual of 36 years.

The sample was radiocarbon dated to 2775±20BP (cal. BP 2948-2786, see Suppl. Tab. S9), roughly corresponding to the archaeological chronology given by the excavators.

###### Lapithos, Plakes cemetery

**Lapithos Plakes, Tomb 601**.

Grave 601 is a shaft tomb^69^. Two burial phases could be discerned in this grave. The first phase had only one individual which must be our sample **CGG_2_022527/Rise 1670**, since the label of the museum attached to this find says “Gr 601/1” (= tomb 601 cranium I as in the publication of the site^69^). This first phase is dated to the Cypro-Geometric I (c. 1050-950 BCE). The second phase contained two individuals and was dated to the Cypro-Geometric II (c. 950-900 BCE). Cranium II (**CGG_2_022509/Rise 1652**) and Cranium III (**CGG_2_022524/Rise 1667**) belong to this second deposition. The available osteological analyses suggest that both Cranium II and Cranium III are adult male individuals^74^(Fischer 1986, 40). However the genetic sex appears to be female for **022509** and male for **022524**. According to the digital archive of the Medelhavsmuseet in Stockholm (Carlotta database) **CGG_2_022527** also belongs to an adult individual, now genetically assessed as female. The kinship inference shows these three individuals share a first degree relation. It is therefore a family grave.

Cranium III (**CGG_2_022524/Rise 1667**) was radiocarbon dated to 2860±15BP (cal. BP 3060-2884, see Suppl. Tab. S9), roughly corresponding to the chronology proposed by the excavators. The date also suggests that the first and the second strata in the graves were probably closer in time than previously thought, which corresponds well to the results of the kinship inference and the close relatedness of the analysed individuals.

More difficult is to understand how a duplicate (exactly the same individual, or a twin) could be found in a different grave. However, it is hypothesised here that the bones of the CGG_2_022504 sample, which have no proper provenience (only Lapithos, Vrysi you Barba? See below) is likely not a twin rather the same individual as **CGG_2_022524** here, whose bones ended up for unknown reasons in two different boxes.

**Lapithos Plakes, Tomb 602**.

Tomb 602^69^ is a shaft grave similar to the others (601 and 603). The excavators could see two burial phases. Skeleton I in the first stratum was very well preserved and was dated to a later part of the Cypro-Geometric I period (c. 1000-950 BCE). The body was well preserved and adorned with metal items of both bronze and Iron and remarkable toe-rings. According to the digital archive of the Medelhavsmuseet in Stockholm (Carlotta database) this individual is a young adult. The excavators suggest it is male individual. The other two skeletons were found in the second earlier burial phase and were not in their original position.

Four samples were taken from this grave **CGG_2_022526/Rise 1669, CGG_2_022525/Rise 1668**, **CGG_2_022514/Rise 1657**, and **CGG_2_022528/Rise1671**. It is important to take into account that according to both the digital archive of Medelhavsmuseet in Stockholm (Carlotta database) and Fischer^74^ it is not clear if the designations of the skull currently present at the museum correspond with those given in the publication. According to Fischer^74^ sample **CGG_2_022514/Rise 1657** possibly belongs to a female individual of c. 20 years.

Sample **CGG_2_022526/Rise 1669** was radiocarbon dated to 2795±15BP (cal. BP 2952-2853, see Suppl. Tab. S9); the date corresponds to the archaeological chronology proposed by the excavators.

**Plakes, Tomb 603**.

Tomb 603 is a shaft grave, where only one burial phase with only an individual (**CGG_2_022529/Rise 1672**) could be identified. The body had a bronze fibula and grave goods nicely deposited to form a half circle. The material dates this deposition to an early phase of the Cypro-Geometric I (c. 1050-1000 BCE). According to the digital archive of the Medelhavsmuseet in Stockholm (Carlotta database) the sample belongs to an adult male individual; the data is confirmed by the genetic sex.

The sample was radiocarbon dated twice. The first result (UCIAMS 224137) gave 2995±20BP (cal. BP 3320-3076, see Suppl. Tab. S9). The second result (UCIAMS 223990) gave 2985±15BP (cal. BP , see Suppl. Tab. S9). The dates suggest that the individual was buried considerably earlier than what the archaeological chronology proposed by the excavators would suggest; however as in other cases from Lapithos, this could be possibly explained by the prevailing marine dietary habits.

**Lapithos unknown/uncertain cemetery**

**Lapithos, Tomb ?**

Sample **CGG_2_022483/Rise 1626** was brought to Stockholm by the Swedish Cyprus expedition in the 1930s^69^. The note that was found together with the material says Lapithos, but there is no tomb number nor any site specification. According to the digital archive of the Medelhavsmuseet in Stockholm (Carlotta database) this individual is an adult female.

**Lapithos, Vrysi tou Barba? (Lapithos, Plakes, tomb 601)**

Sample **CGG_2_022504/Rise 1647** was brought to Stockholm by the Swedish Cyprus expedition in the 1930s^69^. The note found together with the material says “Lapithos, Vrysi tou Barba?” but there is no tomb number or any other information. According to the digital archive of the Medelhavsmuseet in Stockholm (Carlotta database) the bones belong to an adult individual. The genetic information shows that this individual is a duplicate of CGG_2_022524 (see above). As these bones do not have a proper context it is possible that individuals **CGG_2_022504** and CGG_2_022524 were not twins rather the same individual whose bones for unknown reasons got separated somewhere and ended up in different boxes.

This sample has been radiocarbon dated to 2855±15BP (cal. BP 3058-2881, see Suppl. Tab. S9), thus contemporary to the 14C date of CGG_2_022524, confirming that the two samples are contemporary or from the same person.

**Lapithos, Tomb ?**

Sample **CGG_2_022489/Rise 1632** was brought to Stockholm by the Swedish Cyprus expedition in the 1930s^69^. According to the notes found together with the present material the origin is Lapithos. The tomb number, however, is unknown. According to the digital archive of the Medelhavsmuseet in Stockholm (Carlotta database) the bones belong to an adult individual. The sample has been radiocarbon dated to 2740±15BP (cal. BP 2865-2779, see Suppl. Tab. S9). The 14C suggests that the individual was buried during the Cypro-Geometric III period (c. 900-750 BCE) and could be therefore coming from the cemeteries at Plakes or Kastros where there are several contemporary contexts.

**Lapithos, Vrysi tou Barba? Tomb 302 or 308?**

According to the digital archive of the Medelhavsmuseet in Stockholm (Carlotta database), on the cranium from which sample **CGG_2_022490/Rise 1633** was taken is written T2. It could possibly mean Tomb 2 and thus tomb 302 in the final publications^69^. The description in the Carlotta database is not very clear: “*According to notes found together with the present material the origin is ”Lapithos T8, 14-15”. The above-mentioned tomb number is based on an inscription, T2, on the present cranium. None of the numbers, T2 or T8, however, exist in the final publication concerning Lapithos (Gjerstad et al 1934). However, in order to distinguish the various sites in Lapithos the excavators have changed the serial numbers concerning the material from Vrysi tou Barba from tomb 1-23 to 301-323. The present material consists of two individuals: one adult and one subadult respectively. The present materials relate to skeleton: FCL 1, Gr 302, Lapithos. IB: 3, 32-35, 45 and 46 is not fully understood.*”

It is interesting that in tomb 8 or 308^69^ there were four individuals and that the oldest was found together with finds 14 and 15. This might be a clue for the note found at the museum.

Tomb 302 was dated by the excavators to the beginning of the Early Cypriot III period (c. 2100-2050 BCE), while tomb 308 was dated to the end of the Early Cypriot I to the middle of the Early Cypriot II (c. 2200-2100 BCE). Unfortunately this sample did not undergo 14C analyses.

##### ***2.2.6. Rizokarpaso, (de facto) Famagusta district, Northern Cyprus***

The exact Latitude and Longitude of the site is missing. If close to Korovia (see below) possibly 35.51222948712876, 34.27521210066613

Sample provider: Christian Mühlenbock, Mia Broné

Serena Sabatini

There is very little information about this site. It seems to come from an Iron Age tomb close to Rizokarpaso probably in the vicinity of Korovia (see below and Map 1), but no closer information is available.

The sample (**CGG_2_022530/Rise 1673**) was excavated during a Cypriot rescue excavation in 1951. The osteological material was later collected by the SEAA (Swedish Expedition for Archaeology and Anthropology) in 1951-52 by Nils-Gustaf Gejvall et al. According to the digital archive of the Medelhavsmuseet in Stockholm (Carlotta database), on a note found together with the osteological material it is written: “1951/III-8/1, Iron Age”.

P.M. Fischer^74^ writes that the skull was artificially deformed; he also^74^ refers to a letter that he got from Dr. I. Nikolau in which the Cypriot archaeologist writes that the grave in question is called tomb 1951/VIII (=not III) - 8/1 and was found outside the village of Korovia (today Κορόβεια/*Kuruova*) on the Karpas Peninsula. The tomb was removed and the objects sent to Nicosia Museum (file 25/51 bl. 5). It does not seem that any closer date or observation can be established.

The sample was radiocarbon dated to 3070±20BP (cal. BP 3356-3220, see Suppl. Tab. S9). The 14C suggests that the individual was not buried during the Iron Age rather during the Late Cypriot IIIB-C periods (c. 1375-1200 BCE). A forthcoming study will investigate the effects of diet on the obtained 14C results.

##### ***2.2.7. Vounos-Bellapais, (de facto) Kyrenia district, Northern Cyprus***

The exact Latitude and Longitude of the site is missing. However the modern village of Kazafana has Latitude: 35.317222 and Longitude: 33.354167.

Sample provider: Christian Mühlenbock, Mia Broné

Serena Sabatini

**Vounous-Bellapais, Tomb 69.**

The site was found southeast of the village of Kazaphani (today, Καζάφανι/*Kazafana* or *Ozanköy*), and east of the Abbey of Bellapais (Map 1). It was excavated between 1931 and 1938, by different expeditions^40^. In 1931-1932 P. Dikaios from the Cyprus Museum excavated tombs 1-48^79^. In 1933 the Cyprus Museum and the National Museum of France, represented respectively by P. Dikaios and C.F.A. Schaeffer excavated tombs 49-79^80^. In 1937-38 the British School at Athens excavated tombs 80-164^81^. Some of the skeletons excavated by Schaeffer were sent to C.M. Fürst in Sweden. Since Fürst passed away before any investigation was done, the osteological analyses were made by C.H. Hjortsjö^73^. As far as the chronology of the context is concerned there is a letter written by R.S. Merrillees to Paul Åström (28/02/1980) which says "nothing earlier than MC I”^40^, thus c. 2000-1850 BCE.

Tomb 69 consisted of a single chamber and looked disturbed at the moment of the excavation^40^. Cranium 2 (**CGG_2_022535/Rise 1678**) was found immediately to the right of the entrance together with finds 1-10^40^. The skull was probably not in situ. According to Hjortsjö^73^ the skull belonged to a male of approximately 30 years. Fischer^74^ defined the find as belonging to a male adult of c. 18-20 years. Dunn-Vaturi questions both results, because the grave seems to have contained just one individual, who was furnished with grave goods including a spindle whorl and a pin, typical for female burials.

As a curiosity, it is interesting to mention that according to Hjortsjö's craniometrics studies^73^ (such studies were still carried out at the beginning of last century) cranium II was considered a sort of outlier, compared to the other skulls from the region.

##### ***2.2.8. Hala Sultan Tekke, Larnaca district***

Latitude: 34.887; Longitude: 33.605

Sample provider: Peter M. Fischer.

Peter M. Fischer

Hala Sultan Tekke is the modern name of the Late Bronze Age city which is located on the south-eastern coast of Cyprus along the western bank of the Larnaca Salt Lake. The lake was once linked to the Mediterranean thus creating a well-protected harbour. The ancient city, one of the largest Bronze Age sites in the eastern Mediterranean, flourished for approximately 500 years, that is between c. 1630 and 1150 BCE^42,43,82^. This period corresponds to the local Late Cypriot I–IIIA period. Fifteen seasons of excavations directed by P. M. Fischer (2010–2024) exposed four city quarters and an extramural cemetery east of the city with rich finds reflecting the cosmopolitan societies of the city. In addition to abundant mortuary goods of local origin, a high proportion of objects were imported from numerous cultures. These encompass from west to east the area from Sardinia to Afghanistan/India, and from north to south the Baltic Sea to Sudan. Dominating among the imports are those from the Mycenaean, Minoan, Anatolian, Egyptian and Levantine cultures (in that order). The three tombs (Z9, X and RR) from which the samples of this project derive are situated in the extramural southern cemetery of the site.

**Pit Tomb Z9**

This tomb was excavated in 2017. It is an abandoned well with a diameter of c. 1.1 reused for interments of likely underprivileged individuals (Fig. 5). Mortuary goods are virtually missing^44^. The pottery sherds associated with the skeletons certainly derive from the backfill thus they should not be considered as burial goods. There are five entombed individuals: two adults and three children, who were buried or rather dumped at a single occasion. Our samples derive from Skeletons 1 and 3.

Skeleton 1 (**CGG_2_022116/Rise 1417**) is estimated to be a male individual, 30–40 years of age and 166 ± 4 cm in height. He suffered from malnutrition and a poor dental status (caries, calculus, periodontal disease, periapical abscess, enamel hypoplasia), porosities in the roofs of the ocular orbits attributed to *cribra orbitalia*, osteoarthritis, periostitis, and healed fractures of a rib and fibula were identified. A perimortem trauma on the frontal bone by a pointed weapon might have been the cause of death. The age of Skeleton 3 (**CGG_2_022117/Rise 1418**) (and 5) was estimated to 34–36 weeks, indicating death at birth or possibly a fatal premature birth. The mother is likely to be the adult female (Skeleton 2) holding in her arms these two skeletons (3 and 5), most likely twins. The genetic sex of **022116** and **022117** resulted male and female, respectively.

The tomb is dated to the first half of the 12th century BCE, which corresponds to the zenith of the “crisis years”^44^ just before the city was destroyed and abandoned. One sample (**022116**) was radiocarbon dated to 2965±15BP (cal. BP 3208-3070, see Suppl. Tab. S9) which confirms the chronology provided by the archaeological material.

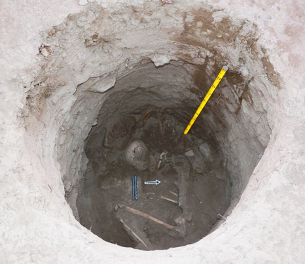

Fig. 5 Well/Tomb Z9 during excavation. Photograph P.M Fischer.

**Chamber Tomb X**

Chamber Tomb X, excavated in 2016 represents a tomb with exquisite mortuary goods and personal belongings. It consists of two interconnected chambers of figure-of-eight shape^42,43,83^. The tomb was dug into fairly soft clayish soil down to a depth of 1.35 m from today’s eroded surface. Just below colluvial soil and on top of this double chamber is a centrally placed shaft from which the tomb was entered. Despite the relatively simple architectural design, the number and quality of the finds point to individuals of high rank who seem to have belonged to the ruling class of the city. The sequence of the buried seventeen individuals, who comprised eight infants and nine adults, could not be fully established, because the remains of previously buried individuals were swept aside before each new interment. However, the position and nature of numerous finds along the periphery of the tomb enabled us to present a model of the sequence of deposition of the skeletons and burial goods. The tomb gifts of complete items are more than 100 in number. In addition to high-quality local objects, the proportion of imported items of gold, silver, precious stones and ivory from the Mycenaean, Minoan, Egyptian and Levantine cultures is high. Among the finds are Egyptian scarabs of which one shows the cartouche of Tuthmosis III, which permit *post quem* dates based on the Egyptian pharaonic genealogy.

Our samples from Chamber Tomb X belong to Skeletons 1, 2 and 5. Skeleton 1 (**CGG_2_022122/Rise 1423**) was in a fragmentary state, possibly female (the genetic sex actually resulted to be female), c. 30 years old with no severe pathological conditions. Some teeth show carious lesions and are covered with calculus resulting in periodontal disease. The other analysed skeletons from this tomb are Skeletons 2 (**CGG_2_022123/Rise 1424**) and 5 (**CGG_2_022126/Rise 1427**); their genetic sex resulted to be female and male respectively.

Judging from local and imported pottery and supported by radiocarbon, Tomb X was in use from roughly 1550 to 1200 BCE, i.e. from Late Cypriot IB to IIC. Skeleton 1 was radiocarbon dated^84^ to 3006±36BP (cal. BP 3337-3072, see Suppl. Tab. S9) thus in line with the chronology based on the study of the ceramic material and other finds..

**Chamber Tomb RR**

Chamber Tomb RR is another rich tomb south of Chamber Tomb X^85,86^. It has a figure-of-eight shape with two interconnected chambers (3.5 by 2.5 m in total). It contained the skeletal remains of 60 individuals (minimum number of individuals) and 186 objects representing personal belongings and tomb gifts. Some skeletons are articulated but most of them were non-articulated comprising all age categories from infants to mature adults. Tomb RR provided indications of ritual activities involving fire. In absolute terms, most of the ceramics and other finds can be dated from 14^th^ to the 13^th^ century BCE. There are Mycenaean, Minoan, Anatolian, Levantine and Egyptian imports. Zoological, botanical, and other remains from the tomb include skull fragments of cattle with horns, a fragment of a notched sheep scapula, a complete fish skeleton with the scales intact, fish bones including those of Nile perch, murex shells, plant seeds, and pieces of ochre.

Our samples belong to Skeletons 1, 2, 3, 6, 15 and 20. Skeleton 1 (**CGG_2_022924/Rise 1889**) is a female individual of 18-22 years. The individual shows traces of calculus, caries, enamel hypoplasia, and periostitis (ulna). Skeleton 15 (**CGG_2_100074**) was estimated a possible male adult (now disproved by genetics), of undetermined age with the following pathological finds: caries, periodontitis, periapical abscess, *cribra orbitalia*. Skeleton 20 (**CGG_2_100076**) is badly preserved. Sex could not be determined osteologically of this 25-30 years old individual. Skeleton 2 (CGG_2_022925/Rise 1890) is a 16-20 old female with the following pathologies: calculus, caries, LEH, and *cribra orbitalia*. Skeleton 6 (**CGG_2_022928/Rise 1893)** is an adult (35-40 years). Osteologically, the sex could not be determined. The individual suffered from calculus, caries, and periodontitis. Skeleton 3 (**CGG_2_022926/Rise 1891**) is a child 8± 2 years old. Skeleton 1 (**CGG_2_022924**) has been radiocarbon dated to uncal. BP 3085±15 (cal. BP 3363-3241, see Suppl. Tab. S9).

#### **2.3. Egypt**

##### ***2.3.1. Maasara, Helwan, Cairo Governorate***

Latitude: 29.883, Longitude: 31.3015

Sample provider: Medelhavsmuseet, Stockholm

Serena Sabatini

The cemetery at Maasara, Helwan, south of Cairo, is the largest necropolis of the Early Dynastic Period in Egypt with more than 10.000 tombs^87^. It was first investigated by the Swedish archaeologist Hjalmar Larsen in 1937^87^(see also the Medelhavsmuseet’s archive [Carlotta database]). Larsen organised just one excavation season and uncovered six graves from the Early Dynastic period^87^. The sample included in this study comes from grave 6 and it is supposed to be individual a. It is an adult, possibly a female (**CGG_2_022554**). In the archive of the Medelhavsmuseet the following is reported:

“*According to the literature tomb 6 should contain[sic] of one individual with fragmentary upper limbs caused by an assumed looting. However, within the material two individuals was[sic] located with the same designation, that is: grav 6, lik 6 [= Swedish meaning tomb 6, body 6]. To separate them they have the suffix a and b respectively, the latter with the inventory number: MME 2006:001H. The bone elements were wrapped in newspapers from 1937 as well as 1955*.”

#### **2.4. France**

##### ***2.4.1. Lano Cave, Lano commune, Haute-Corse Department of France, Corsica***

Latitude: 42.37955375060895; Longitude: 9.240410810529346

Sample provider: Patrice Courtaud

Patrice Courtaud

The burial cavity of Lano is located in the centre of Corsica at 835 m of altitude above sea level. The cave is on an impressive limestone cliff. The opening is at c. 20 m above the bottom of the cliff and it must have been very difficult to access. Exceptional sedimentary conditions made possible the preservation of two wooden coffins associated with human remains. One of the coffins is dated to 1200 cal. BCE (unpublished data, courtesy of P. Courtaud).

Analyses of the wood essence show that one of the coffins is made of yew (*Taxus baccata*). The only comparison for the use of wooden coffins so far, both from a typological and chronological point of view, can be made with the Early Bronze Age oak coffins discovered in Denmark.

The human remains of a minimum of six individuals, without anatomical connections, have been uncovered. They are composed of two children, one of them a perinatal, and four adults or sub-adults. The radiocarbon dates from the human bones spread over the 11^th^ and 10^th^ century BCE (unpublished data, courtesy of P. Courtaud). Preservation of bones from the stratigraphic horizons which yield wood specimens was excellent as some soft tissue remains (cartilage, cerebral material) were also found. The field observations suggest primaries and successive burials secondary dislocated by the taphonomic factors. A total of 374 bones and fragments were collected and numbered individually.

Five skulls (bone #108, # 131, #132, #177 and #317) were sampled and are presented in this study (**CGG_2_021108/Rise 1185**, **CGG_2_021109/Rise 1186**, **CGG_2_021110/Rise 1187**, **CGG_2_022152/Rise 1433**, **CGG_2_022153/Rise 1434**).

##### ***2.4.2. Migennes, Yonne department in Bourgogne-Franche-Comté***

Latitude: 47.968158135953104; Longitude: 3.5159857185051036

Sample provider: Lorenz Rahmstorf

Rebeca Peake, Lorenz Rahmstorf

The Late Bronze Age cemetery of Migennes «Le Petit Moulin» (France), excavated by Inrap in 2004, is located on the banks of the Armançon River about 3 km upstream from its confluence with the Yonne in northern Burgundy. With its 61 burials (Fig. 6), it is one of the largest cemeteries dating to the early phase of the Late Bronze Age (14^th^-13^th^ century BCE) in central-eastern France. Two burial groups make up the cemetery, a first group comprising 25 inhumations and 10 cremation burials dispersed around a circular ditch monument and a second group located about 50 m to the south with 21 cremations and 4 inhumations detected to north of two circular ditch monuments. All of the burials contain a large array of objects, adornments (bronze bracelets, pins and beads), some weapons (daggers, arrowheads and sword) and in some cases a small box or leather bag holding a standardised tool kit which includes a bronze knife or dagger, a sharpening stone, flint lighters and iron pyrite.

Two of the burials stand out by their unusual grave goods, each containing weighing equipment, which consists of small bone or antler weighing beams (about 10 to 12 cm in length) and weights, either purposely made or of a more opportunistic nature^88,89^. One of the burials (tomb nr. 251) contained a weighing beam and a bronze Rixheim type sword. Two weighing kits as well as specialised metalworking tools were found in the second burial (tomb nr. 298). This inhumation, housed in a wood lined burial pit, is that of an adult male in a lateral position with two groups of objects, placed on the left hand side of the body. A cylindrical bone beam with flared ends and several weights, as well as fragments of gold, bronze tweezers, a dagger and three arrowheads minus their shafts were contained in a small box with an elaborate bronze fastening placed at the waist. A collection of larger objects found further down the body includes a rectangular shaped balance beam made from antler and decorated with eyespot motifs, a socketed bronze hammer, a small bronze dagger, stone tools for abrasion and a sphendonoid shaped stone weight similar to Late Bronze Age finds in the Eastern Mediterranean (e.g. Uluburun shipwreck). This unique find underlines the specialised knowledge of the individual buried in this particularly rich grave and suggests links to the Mediterranean world whilst raising questions about the origin, status and mobility of bearers of weighing equipment during the Late Bronze Age. These people were specialist artisans with extensive knowledge of metalworking, as well as being merchants capable of acting as intermediaries in the trade and exchange of precious materials.

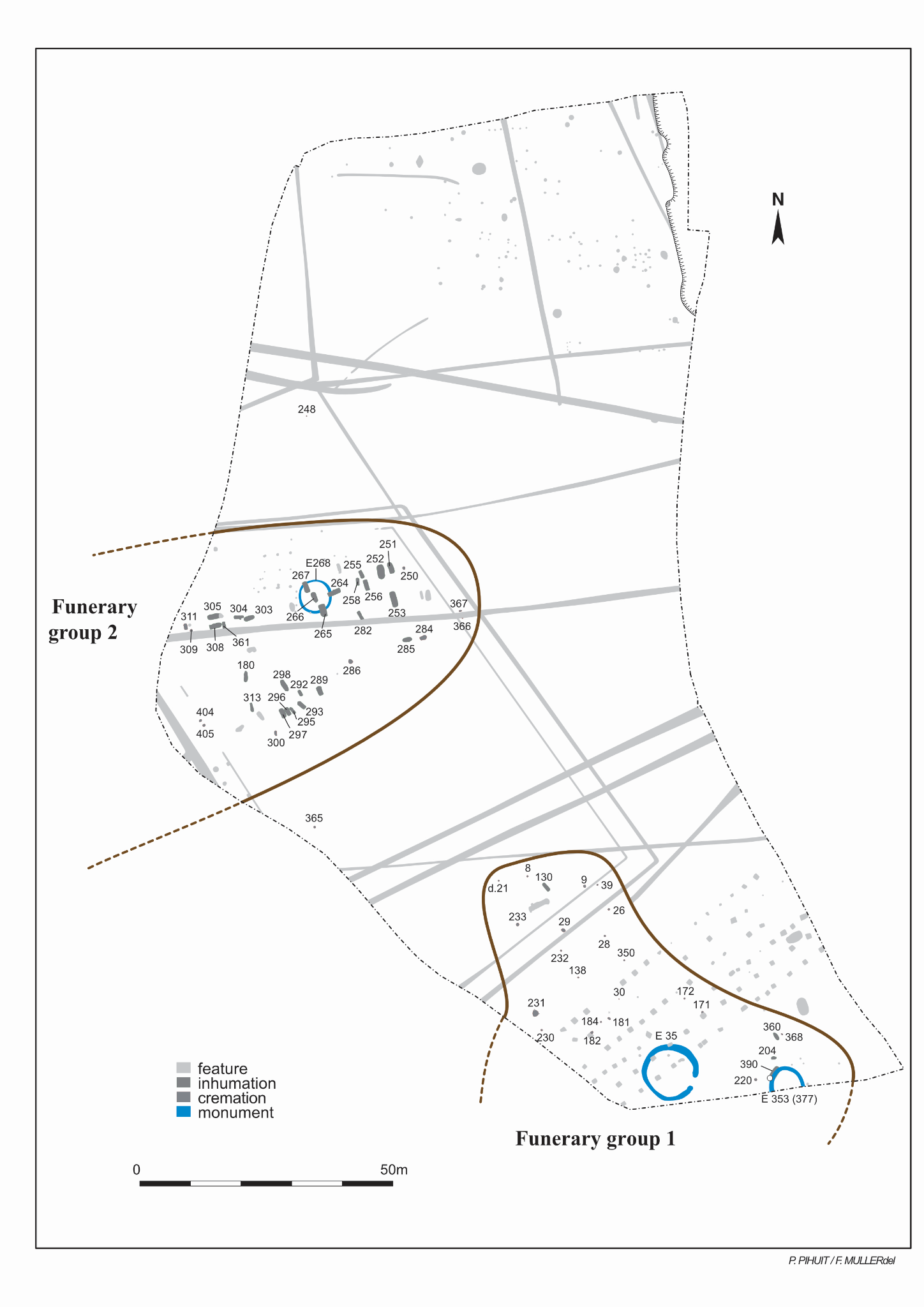

Fig. 6. Plan of the cemetery at Migennes (courtesy of Rebeca Peake and Lonrenz Rahmstorf)

At Migennes there are both inhumations and cremations. All the individuals analysed in this project come from inhumation burials (Table 5).

| **Grave nr., CGG nr.** | **sex/age** | **Grave goods** | **Radiocarbon dates**  **(courtesy of the sample providers)** |
| --- | --- | --- | --- |
| 360A  **CGG_2_021372** | -/Juvenile | Dagger |  |
| 180A **CGG_2_021374** | F/Young adult | Goblet + amber + small conical headed pin+ bronze earrings + bronze bead | 3045±30 (Lyon14455/SacA-50983)and 2σ cal. To 1401-1220 BCE (95,4%) |
| 303A  **CGG_2_021375** | -/Infans II | Goblet + jug + lighter + silex | 3050±30 (Gr-A29960) and 2σ cal. To 1405-1223 BCE (94,5%) |
| 251A  **CGG_2_021376** | -/Juvenis | Goblet + sword + chape + Saint-Gervais type pin + bone balance beam + small bronze cylinders + bronze button | 3050±30 (GrA-30457) |
| 267A  **CGG_2_021377** | M/Mature adult |  | 3010±30 (Lyon14458/SacA-50986) |
| 305A  **CGG_2_021379** | M/Mature adult (c.30 years) | Goblet + disc headed pin + silex | 3010±30 (Lyon14463/SacA-50991) |
| 293A **CGG_2_021381** | -/Infans II, c. 7 y. ± 24 months | Goblet + fossil + bronze earrings + amber bead + limestone bead |  |
| 258A  **CGG_2_021385** | -/Infans I | Goblet + small conical headed pin |  |
| 256B **CGG_2_021387** | -/Juvenis c. 15 y. ± 24 months | wheel pendant + urchin fossil + six swine canines + antler bone + sharpening stone + silex + dagger + arrowhead + spiral headed pin + bronze object | 2910±35 and 2σ cal. To 1220-1003 BCE (95,4%) |
| 252A  **CGG_2_021388** | M/  Mature (elderly) adult | Jug + goblet + quiver + disc headed pin + bronze arrow heads + bronze handle + lighter + gold + silex + deer point + bone button | 3020±40 (GrA-30477) and 2σ cal. To 1400-1156 BCE (91,4%) 1147-1127 (4%) |
| 255A  **CGG_2_021389** | -/Infans II c. 8 y. ± 24 months | Gobelet + pin with trumped shaped head |  |
| 297A **CGG_2_021390** | -/Infans II c. 12 y. ± 30 months | Goblet + lighter + bronze bracelet + bronze neck ring + spiral headed pin + bronze beads + amber bead + silex |  |
| 297B  **CGG_2_021391** | -/Infans I  c. 3 y. ± 12 months | torc + amber + spiral headed pin |  |
| 265A **CGG_2_021392** | M/Mature adult | Lighter + 2 bronze spirals + silex | 2960±35 and 2σ cal. To 1283-1048 BCE (95,4%) |
| 265B **CGG_2_021393** | M/Mature (elderly) adult | - | 2960±35 and 2σ cal. To 1283-1048 BCE (95,4%) |
| 298A  **CGG_2_021394** | M?/  Mature adult | jar + stone anvil + bone balance beams + small bronze weights + sharpeners + set of stones + gold scraps + silex + bronze hammer + bronze arrow beads + bronze tweezer + bronze handle + bronze fittings for box + bronze awl + small bronze items + amber beads + small lead pieces | 3020±30(Lyon14461/SacA-50989) |

Table 5. Synoptic table with data about the sampled individuals from Migennes and their context of origin.

#### **2.5. Greece**

##### ***2.5.1. Apollo Maleatas, Argolis, Peloponnese***

Latitude: 37.5988; Longitude: 23.0855

Sample provider: Ioanna Moutafi, Vassilis Lambrinoudakis

Ioanna Moutafi

The site of Apollo Maleatas at Kynortion Mount, Epidavros, Argolid, is located at a close distance from the famous sanctuary of Asclepios and it is mostly known for the Archaic and Classical sanctuary of Apollo Maleatas on the north slope of the hill. At the top hill of Kynortion Mount, a small Early Bronze Age settlement was also found (lastly excavated by Prof. V. Lambrinoudakis and A. Theodorou)^90^. The settlement developed during the Early Helladic (EH) period, with the earliest human activity at the site dated to the EH I (including three human burials, initially attributed to that phase). However, most of the building phases date to the EH II. After abandonment of the settlement, some, possibly ritual, activity is attested locally and dated to the Middle Helladic (MH) times over the EH ruins^91^. During the following centuries, ritual activities continued; they now take place lower down the hill (during Mycenaean and Geometric times), and intensify in historical times with the establishment of the Apollo Maleatas Sanctuary.

The three burials were located on the Northeastern edge of the hilltop, adjacent to the EH buildings. They consisted of single burials placed in pits, covered by rough limestone slabs (Graves 1-3). The bodies were placed on their sides, in contracted positions. Grave 1 did not contain any grave goods, but the other two did (mostly tools and ornaments), while broken pots and tools were also placed outside of Grave 2. The burials were initially dated in EHI, based on stratigraphic information, as they were associated with the earliest phase of occupation on the hill. However, radiocarbon dating of both samples analysed in this study, as well as an independent dating of a sample from Grave 2 (see below), place all three burials in the 19^th^ century BCE, and thus the MH period.

Individuals from all three graves have been sampled for this study.

**Grave 1.** The sample (**CGG_2_023928-23929**) comes from the temporal bone (petrous portion) of the skeleton. The skeleton belongs to a middle-adult male individual (c. 40 years). The grave contained no grave goods. The radiocarbon date of the skeleton is 3535± 15BP (cal. BP to 3882-3723, see Suppl. Tab. S9).

**Grave 2.** The sample (**CGG_2__2_023931**) comes from the temporal bone (petrous portion) of the skeleton. The skeleton belongs to a young adult (c. 25 years) of indeterminate sex. The grave contained a few grave offerings in association with the body: an obsidian blade, a clay whorl, and a stone grinder/crusher. In close proximity to this grave, and probably associated with it, a pottery assemblage was placed under two stones, with three stone tools on top. The radiocarbon date of the skeleton is 3479±28BP (cal. BP 3835-3646, see Suppl. Tab. S9)

**Grave 3.** The sample (**CGG_2_023933**) comes from the temporal bone (petrous portion) of the skeleton. The skeleton belongs to a young female adult (c. 18-20 years). The body was buried with a copper pin holding the cloth on her shoulder and a pendant around her neck. The radiocarbon date of the skeleton is 3490± 15BP (cal. BP 3831-3698, see Suppl. Tab. S9).

##### ***2.5.2. Ayios Vasileios, Laconia, Peloponnese***

Latitude: 36.9798; Longitude: 22.4782

Sample provider: Ioanna Moutafi, Sofia Voutsaki

Ioanna Moutafi

Ayios Vasileios is the site of the recently (2008) discovered palace of Mycenaean (Late Bronze Age, c. 1700-1100 BCE) Laconia, southern Greece. The site lies on the low hill of Ayios Vasileios, within the modern district of Laconia, at a distance of about 12 km south of the modern town of Sparta. It has been excavated since 2010 by the Archaeological Society at Athens under the direction of former Ephor of Antiquities A. Vasilogamvrou. The site was occupied already in the Early Bronze Age (2^nd^ millennium BCE), when a small village was standing on top of the hill. It may have been abandoned during most of the Middle Bronze Age and resettled at the end of the period (around 1700 BCE). It grew in significance during the early phases of the Late Bronze Age (LH I - LH II), when a cemetery consisting of stone-built graves (cists) and pits, the so-called North Cemetery, was in use next to the settlement. By the middle of the period (LH IIIA), a monumental complex consisting of a central court and impressive, elaborately decorated, buildings was standing on the site. While only a small part of the palatial complex has been excavated so far, all the functions usually associated with a Mycenaean palace (including evidence of central administration and the use of Linear B tablets) are attested. Interestingly, however, Ayios Vasileios differs from other Mycenaean palatial sites in two important respects: a) its development; the site was destroyed around 1250 BCE, some 50 years earlier than other mainland palaces, like Mycenae, Tiryns and Pylos, and b) its spatial organisation; the town surrounding it is relatively small and unfortified.

The excavation of the North Cemetery was conducted between 2010-2016 under the direction of Prof. S. Voutsaki, University of Groningen. 22 graves (12 cists, 1 built-tomb, and 9 pits) and two ”burials” (free-standing bone assemblages on top of another grave) have been found^92^. The period of use of the cemetery spans the Middle Helladic III-Late Helladic II periods (in ceramic terms). The relative (ceramic) dating of each grave cannot be established with great precision, as the graves are either unfurnished or contain very few and modest offerings. However, radiocarbon dating was conducted for most of the skeletons by the Centre for Isotope Research, University of Groningen, as well as 14C Chrono Laboratory at Queen’s University, Belfast and Keck Carbon Cycle AMS Laboratory, UCI confirming the dating of the period of use at c. 1700-1450 BCE. The graves of Ayios Vasileios mostly follow the older (MH) tradition but also display novel features pointing to the Mycenaean era (introduction of elaborate cists and one built-tomb, destined for multiple burials). Similarly, the burial practices expressed in these graves are variable, including both the most traditional forms of single primary burials and innovations such as collective depositions of multiple successive interments, with both primary and secondary burials within the same grave. The total MNI (= Minimum number of individuals) of the cemetery is c. 80 individuals, with most graves containing 1-4 burials, some a few more, and one notable exception (Tomb 21) with MNI >25. Both sexes and all age categories are included in the cemetery, although infants are significantly underrepresented and receive a slightly different treatment^93^.

**Ayios Vasileios, Grave 3**

From this grave we have sampled individual 3.2 (**CGG_2_022385**), which has been osteologically determined as a young adult (c. 25 years) male.

The burial was found in a single secondary bone deposit in cist Grave 3 (with total MNI: 2). It is the earliest burial of this grave.

**Ayios Vasileios, Grave 15**

Burial 15 is a bone assemblage of commingled disarticulated remains, placed as a secondary deposit on top of Grave 20, the grave of a mature adult male. The assemblage of Burial 15 has MNI:2, including a middle adult male (individual 15.1), c. 40 years of age at death, and a child (individual 15.2), 7-8 years old. From this grave we have sampled individual 15.1 (**CGG_2_022360**), which is dated to 3390±15BP (cal. BP 3689-3572, see Suppl. Tab. S9).

**Ayios Vasileios, Grave 21**

Tomb 21 is a built-tomb of unique design compared to all the others in Ayios Vasileios. It contains a large quantity of multiple successive burials (MNI: 25), placed in various contexts in two successive floors . Most of them are dated (based on Radiocarbon dates results) between 1600 and 1450 BCE. From this grave we have sampled three individuals (21.1, 21.6, and 21.8).

Individual 21.1 (**CGG_2_022367**), which has been osteologically determined as an adult female (of c. 35-45 years, comes from a secondary deposit of the upper floor (thus dated among the later burial groups of the grave, but is earlier than Burial 21.6 and 21.8 below). A reversed ceramic vessel (alabastron) was found among the bones of this secondary deposit.

Individual 21.6 (**CGG_2_022370**), which has been osteologically determined as a young adult male (of c. 25 years), comes from the main upper floor deposit of four primary burials, interred at the same time. Individual 21.6 was the very last interment of the four *in situ* skeletons. This individual has been radiocarbon dated to 3201± 33BP (cal. BP 3467-3364, see Suppl. Tab. S9).

Individual 21.8 (**CGG_2_022372**), which has been osteologically determined as an adult female of c. 30 years of age-at-death, is one of the latest interments in the tomb. It was found semi-articulated and slightly displaced on the upper floor, only slightly pre-dating the final collective deposit of four primary burials, which was dated around 1500-1450 BCE. There is no radiocarbon date for this skeleton.

**Ayios Vasileios, Grave 24**

Individual 24.1 (**CGG_2_022383**) represents the primary burial of the pit-grave 24 (MNI:2), which contained two infant burials, one in primary and the other in secondary deposit. 24.1 was the last interment in the tomb and has been radiocarbon dated to 3120± 15BP (cal. BP 3385-3259). Individual 24.1 has been osteologically determined as a neonate; it was buried with a necklace of five stone beads, it also had a cup upside down on the grave’s covering slab.

**Ayios Vasileios, Grave 25**

Individual 25.5.(**CGG_2_023947**) or Skull 5, bone 158 comes from a pit of commingled disarticulated remains, Burial 25, found south of Tomb 21, and probably associated with it as a container of secondary burials coming from Tomb 21. The pit, with MNI: 7, contained both adult and sub-adult skeletal remains. It also contained four small ceramic vessels that could not be associated with specific individuals. Individual 25.5 was an adult, whose sex was osteologically indetermined. There is no radiocarbon date for this skeleton.

##### ***2.5.3. Kalyvia, Elis, Western Greece***

Latitude: 37.8865; Longitude: 21.3839

Sample provider: Ioanna Moutafi, Jörg Rambach

Ioanna Moutafi

The site of Kalyvia lies in the region of Elis, Western Greece, and came to light in 2004 during a rescue excavation directed by Dr. Jörg Rambach^94^. The site was located on a low hill in ancient Elis. It is a Final Neolithic (Chalcolithic) – Early Bronze Age cemetery with 24 graves, mostly rock-cut chamber-tombs, but there are also a few associated pit-graves, arranged in two parallel rows. The graves contained modest offerings, mostly ceramic vessels. They are dated around the transition from the fourth to the early third millennium BC (3400-2800 BCE, covering the Final Neolithic/Early Helladic I period). This cemetery together with Kalamaki in Western Achaea are the only cemeteries of that period discovered in the Peloponnese, and thus of significant importance. The chamber-tomb graves consist of a *dromos* (entrance way), a *stomion* (entrance), and a chamber, which represent a novelty for this period, and were clearly intended to accommodate multiple burials. The chamber tombs (not the pits) were used for successive burials, and contained both primary and secondary skeletal deposits, of 2-16 individuals each, both adults and sub-adults of both sexes. A preliminary estimation of the total MNI of the cemetery is around 150, but the osteological material has not been studied yet.

The sample presented in this study comes from grave 14, and belongs to Skeleton 1 of group 8 (Skull 1, **CGG_2_023925**). The MNI of the tomb was 5. This is a young female (adolescent, c. 15 years) and was the last, and only primary, burial in this grave. The tomb contained a bone palette and four small clay vessels, but no grave goods could be attributed with certainty to specific individuals. The radiocarbon date of the skeleton (obtained by UCI Keck AMS Facility) is 4550±15BP (cal. BP 5316-5059, see Suppl. Tab. S9), thus in line with the archaeological chronology.

##### ***2.5.4. Kirrha, Phocis, Central Greece***

Latitude: 38.431; Longitude: 22.444

Sample provider: Ioanna Moutafi, Anna Lagia, Raphaël Orgeolet

Ioanna Moutafi, Raphaël Orgeolet

The prehistoric archaeological site of Kirrha, in the south-east edge of the plain of Itea, Phokis, Greece, occupies a low mound c. 230 m from the northern shore of the Corinthian Gulf. Two intense excavation campaigns were undertaken by the French School of Athens in the late 1930s. They uncovered extensive areas of a Middle Helladic (c. 2100–1700 BCE) to Late Helladic (c. 1700–1075 BCE) settlement and numerous graves of the same period, whereas deep soundings, halted by the water table, revealed a long-lived tell, already established by the end of the Early Helladic period. Recent geoarchaeological investigations have shown that the foundation of the settlement took place earlier in the neolithic times^95^. With the publication of the first excavations^96^, Kirrha became a key site for the Aegean Prehistory, and one of the most important for the understanding of the Middle Helladic (MH) period in Central Greece. After a long break, archaeological activity resumed in the 1960s, with a series of numerous rescue excavations^97^. Eventually, a new research programme was set up in 2009, under the co-direction of the French School of Athens and the Ephorate of Antiquities of Delphi^97^. Among other goals, a better understanding of the funerary practices through time was considered as a significant issue and tackled through a close cooperation between archaeologists, anthropologists, and bioarchaeologists.

The funerary remains of Kirrha come from at least 40 contexts, associated with various grave forms (stone cists, few shaft graves, burial pits, and few pot burials). In the MH period, the tombs are generally single intramural simple graves, while in the transitional period (Middle Helladic III – Late Helladic I/II) the burial grounds extend over abandoned MH houses and display a variety of funerary forms. These include single and collective burials of both sexes and all ages, in various types of graves and disposal^98^. Large collective later Mycenaean shaft graves were also uncovered, whose relation to the settlement still needs clarification. The total number of individuals is not yet estimated as the osteological material is currently under study, but it certainly surpasses 80. 11 of the sampled individuals (Table 6) provided results.

| **Tomb** | **Sex/Age** | **Observations** |
| --- | --- | --- |
| 150.1 | M/Young adult (c. 20 years) | Burial 150.1 (**CGG_2_022400/Rise 1605**) comes from cist grave 150, with a collective burial [MNI]: 3), arranged in two levels: two secondary burials (150.2 and 150.3) placed in two different compartments on the lower floor, and one intact primary burial (150.1) on the upper floor, laying in flexed position on the right side. No grave goods. The genetics revealed a 1st degree relation with 152.1  **14C**: 3305±20 BP, cal. BP 3567-3466 (see Suppl. Tab. S9) |
| 150.2 | F/Older adolescent (c. 17 years) | Burial 150.2 (**CGG_2_022401/Rise 1606**) comes from the lower level of grave 150. These skeletal remains belonged to a fully disarticulated skeleton, placed as a secondary deposit in a stone compartment below the floor on which Burial 150.1 was lying. No grave goods.  **14C**: 3310±15BP, cal. BP 3567-3481 (see Suppl. Tab. S9) |
| 151.1 | F/Neonate | The bones of burial 151.1 come from a disturbed grave (probably of pit type), containing the remains of two neonates. It is unclear if the skeleton of B151.1 (**CGG_2_022403/Rise 1608**) belonged to a disturbed primary burial or to a secondary deposit. No grave goods. |
| 152.1 | M/Middle adult | Burial 152.1 (**CGG_2_022404/Rise 1609**) comprised the secondary burial of a single, fully disarticulated, skeleton in a pit grave next to grave 150. It is plausible that this skeleton was first interred in grave 150. No grave goods. 1st degree relation with 150.1  **14C**: 3315±15 BP, cal. BP 3567-3483 (see Suppl. Tab. S9) |
| 238.1 | F/Neonate | Burial 238.1 (**CGG_2_022409/Rise 1614**) is an intact primary burial from grave 238, a simple pit.The body was lying flexed on its right side. No grave goods. |
| 401.6 | Young child (c. 3.5 years) | The burial comes from grave 401, a pit deposit of commingled disarticulated remains (MNI: 7). This secondary burial included four adults (401.1-4), a neonate (401.5), a child (401.7) and the young child 401.6 (**CGG_2_022392/Rise 1597**). Stratigraphically, this is one of the latest burials of the Kirrha cemetery. Grave goods: Late Helladic I-II jug. |
| 459.2 | M/Young adult | Burial 459.2 (**CGG_2_022405/Rise 1610**) comes from cist grave 459, containing exclusively secondary remains (MNI: 5), in two successive layers. The upper layer included the disarticulated remains of two individuals (459.1 and 459.2); whether these remains originated from bodies originally placed in this grave is under investigation. The lower layer contained the secondary remains of another three individuals: a carefully placed secondary burial of a young adult female in the middle (459.3) and bones from another two adult individuals towards the sides of the grave (459.4-5). No grave goods. |
| 504.1 | M/Prime adult | Burial 504.1 (**CGG_2_022411/Rise1616**) comes from grave 504/505, a mortar-lined pit containing a secondary deposit of commingled disarticulated remains (MNI: 3). No grave goods  **14C**: 3336±24 BP, cal. BP 3677-3481 (see Suppl. Tab. S9) |
| 513.1 | F/Adult | This sample (**CGG_2_022413/Rise 1618**) comes from a secondary assemblage of disarticulated bones, probably associated with the high-status female burial of L515. The MNI is not yet finalised as the assemblage is still under analysis.  **14C**: 3355±15 BP, cal. BP 3684-3493 (see Suppl. Tab. S9) |
| 753.1 | M/Neonate | Burial 753.1 (**CGG_2_022396/Rise 1601**) is an intact primary burial from a small cist, grave 753. The body was lying flexed on its right side. No grave goods.  **14C**: 3130±15 BP, cal. BP 3393-3266 (see Suppl. Tab. S9) |
| 762.1 | M/Young child (c. 3 years) | Burial 762.1 (**CGG_2_022398/Rise 1603**) is the single primary burial of a young child, placed in pit grave 762. The body was lying flexed on its left side. No grave goods.  **14C**: 3345±15BP, cal. BP 3677-3489 (see Suppl. Tab. S9) |
| 790.1 | M/Older child (c. 6-9 years) | Burial 790.1 (**CGG_2_022415**) This is the single primary burial of an older child, placed in pit grave 790. The body was lying flexed on its right side. No grave goods. |

Table 6. Synoptic table with data about the sampled individuals from Kirrha and their context of origin.

##### ***2.5.5. Voudeni, Achaea, Western Greece***

Latitude: 38.2540, Longitude: 21.7819

Sample provider: Ioanna Moutafi (excavation director: Lazaros Kolonas)

Ioanna Moutafi

The Mycenaean site of Voudeni is located in western Achaea, 7km north–east of the modern city of Patras. The site consists of a cemetery and the associated settlement, the latter only minimally excavated. The cemetery was discovered in 1987, and it has been systematically excavated since by the local Ephorate of Antiquities, under the direction of Honorary General Director of Antiquities Lazaros Kolonas. Voudeni is one of the largest Mycenaean cemeteries in the region of Achaea and was in use for more than three centuries, covering the entire time span between the end of Late Helladic (LH) IIB and the LH IIIC/Sub-Mycenaean period (c. 1450–1050 BCE). It consists of more than 80 chamber tombs, of which c. 75% have been fully excavated. 35 chamber tombs (Tombs 1–44) have been fully published by the excavator^99^ and the osteological material of 20 of those has been analysed and published by Moutafi^100^. The study of the remaining tombs is currently in progress.

The chamber tombs were designed to receive multiple successive burials, and many of them were in use for the entire life-span of the cemetery. A range of 2-27 Minimal Number of Individuals (MNI) per tomb was attested, with an average of 10.3. Both sexes and all age categories were included in the tombs, with adult burials predominating. In accordance with the typical Mycenaean funerary treatment, the tombs included exclusively inhumations, arranged in both primary and secondary burial assemblages, accompanied by grave goods (mostly ceramic vessels, but also jewellery, other ornaments, weapons, and other objects). Mortuary practices were variable, with the emphasis shifting between individual and collective notions, both between certain tombs and specific time periods.

A large number of samples from Voudeni is included in this study (Tab. 7), coming from several different graves and contexts (see table below).

| **Grave nr.**  **(sample nr)**  **Relative chronology** | **Sex/Age** | **Observations/Grave goods/14C (BP)/ 2σ cal.** **(BCE)** |
| --- | --- | --- |
| Τ4Γ  (**CGG_2_023811**)  LHIIIB |  | Tomb 4 was one of the two largest (and oldest) tombs of the Voudeni cemetery (the other being Tomb 75), with evidence of continuous use from the LHIIIA1 to the LHIIIC Late. Despite this long period of use, the osteologically attested MNI was only 7 due to the very poor bone preservation, but the true number of interments must have been much higher. The skeletal material was found in various, both primary and secondary, burial contexts. Three primary burials of the LHIIIC period were placed at the eastern part of the chamber (T4/A, T4/B and T4/Δ), together with a single secondary deposit of the LHIIIB (T4/Γ). At the south, three primary burials dated to the last phase use, LHIIIC Middle/Late, were located (T4/E, T4/ΣΤ, Τ4/Ζ). Finally, the western part was occupied by a large secondary pile of bones dated in the LHIIIA period, all extensively damaged, and the eastern part by some more scattered bones (T4/H and east secondary deposit) dated in LHIIIB to LHIIIB times.  This sample comes from Burial T4/Γ, the single secondary deposit dated to LHIIIB2.  Grave goods: Two ceramic vessels, a few beads, and a clay button were attributed to this burial.  **14C**: 3030±15BP, cal. BP 3331-3170 (see Suppl. Tab. S9) |
| T4 - East chamber secondary deposit, Individual Ε2_sample 2  (**CGG_2_023814**)  LH IIIA-B | F/Young adult | This sample originates from the secondary deposit of scattered bones at the east chamber (see above), with no securely associated grave goods. The spatial relationship of these bones to later burials corroborates the radiocarbon dating, with this sample predating the last phase of use in the tomb.  **14C**: 3055±15BP, cal. BP 3347-3210 (see Suppl. Tab. S9) |
| Τ5/Θ-ΙΓ (Λ1)_bone 380  (**CGG_2_023816**)  LH IIIA-LH IIIC EARLY | M/Adult | Tomb 5 was in use in LHIIIA and then again in LHIIIC. The tomb included various funerary deposits in two successive layers. The top layer was used for at least two burials (Τ5/Α-Β) of the LHIIIC Middle and Late period, while the lower floor included six primary burials (Τ5/Γ-Τ5/Η) of the LHIIIIC Middle period (possibly buried in one event). Also, a pit (T5/Θ-ΙΓ) with commingled secondary remains of at least 12 individuals was discovered, containing grave goods mostly of the LHIIIA period, but also some of the LHIIIIC Early. Therefore, the pit skeletal remains cannot be securely associated with one phase or the other.  All samples included in this study from Tomb 5 come from the pit assemblage (Τ5/Θ-ΙΓ).    **Grave goods:** The pit assemblage included several grave goods (ceramic vessels, jewels etc.) that cannot be securely attributed to specific individuals. |
| Τ5/Θ-ΙΓ (Λ1)_bone 216  (**CGG_2_023818**)  LH IIIA-LH IIIC EARLY | M/Adult | See above |
| Τ5/Θ-ΙΓ (Λ1)_bone 297  (**CGG_2_023819**)  LH IIIA -LH IIIC EARLY | F/Adult | See above |
| T9/Γ + sec.dep_bone 100  (**CGG_2_023820**)  LH IIIA -LH IIIC | M/Adult | Tomb 9 was continuously used during the entire LHIII period. It contained two disturbed primary burials (T9/A-T9/B) and a large secondary deposition in a pile on the floor (T9/Γ & secondary deposit) of MNI: 6, but no artefacts can be securely associated with specific skeletons, therefore the exact dating of the bones is uncertain, as the grave goods accompanying the assemblage are of mixed date.  The sample of this study comes from the secondary deposition, and its 14C date places it in the LHIIIB period.    **Grave goods:** A very large quantity of grave offerings were found in this context, including ceramic vessels, bronze weapons, bronze and silver jewels and ornaments, glass and gold plates, sheets, and various beads. It is impossible to directly associate any of them with specific individuals.  **14C**: 3020±15BP, cal. BP 3329-3163 (see Suppl. Tab. S9) |
| T14/A-H (IND. C)_bone 3 (Skull 1) (**CGG_2_2023829**)  LH IIIA2/IIIB | M/Middle Adult | This is a small tomb that, based on ceramic evidence, was used only during the LH IIIC (Early and Middle) period. The tomb’s MNI was 13, and the skeletal remains were found in one large secondary deposit (T14/Α-Η, MNI: 10), three extra (scattered) crania (T14/Θ-Κ), and a disturbed primary burial (T14/Λ) of the LH IIIC Middle date, preserving only the lower skeleton. The samples used here come from the secondary deposit, associated with artefacts of the LH IIIC Early date. However, sample Ind. C_bone 3 shows a 14C date of the LH IIIA2/LH IIIB date.    **Grave goods:** Five ceramic vessels of the LH IIIC Early period, a bronze ring and a pin were found in this assemblage. It is not possible to directly associate any of them with specific individuals.  **14C**: 3095±15BP, 3370-3245 (see Suppl. Tab. S9) |
| T14/A-H (IND. E)_bone 40 (Skull 3), (**CGG_2_023830**)  LH IIIC EARLY | F/Prime to Middle Adult | This sample also comes from the secondary deposit (T14/Α-Η). |
| Τ14/Α-Η_bone 76 (**CGG_2_023831**)  LH IIIC EARLY | M/Adult | This sample also comes from the secondary deposit (T14/Α-Η). |
| T16/Γ-IND. A_sample 1_bone 610 (**CGG_2_023835**) & Γ-IND. A_sample 2_bone 579  (**CGG_2_023836**)  LH IIIC MIDDLE-LATE | F/Middle Adult | Tomb 16 was in continuous use during the entire LH III period. The tomb contained the largest number of burials in the Voudeni cemetery (MNI: 27). The deceased included two sub-adults and 25 adults of all ages and both sexes. The burials were placed in various funerary deposits, arranged in two successive layers. The lower floor included a large secondary bone pile (T16/ΣΤ-Μ), with secondary remains of at least 12 people spanning the period LH IIIA2-LH IIIC Early; plus two smaller deposits of secondary remains placed in pits: pit I (T16/O) with MNI: 5 dated in the same time span, and the earliest pit II (T16/Π-Υ) with MNI: 3, dated in LH IIIA2. Three isolated crania were also found on the lower floor (T16/N and T16/Ξ). The upper floor contained two primary burials (T16/A and T16/B) of the LH IIIC Late period, and one secondary deposit (T16/Γ), slightly earlier in date (LH IIIC Middle/Late) with MNI: 2 (including mostly remains of one mature adult female, and only few other extra bones (not counting for the total tomb MNI).    This sample comes from the main individual of the secondary deposit T16/Γ, dated in LH IIIC Middle/Late.    **Grave Goods:** Five stirrup jars and three clay buttons, possibly associated with this individual.  **14C**: 2950±15BP, cal. BP 3173-3005 (see Suppl. Tab. S9) |
| Τ16/ΣΤ-Μ – IND. E  (**CGG_2_023840**)  LH IIIA-LH IIIC EARLY | M/Middle Adult | This sample comes from the large secondary deposit of the lower floor (see above). It is not possible to associate it with specific grave goods.    **Grave goods:** Several grave goods accompanied this secondary assemblage (ceramic vessels, clay buttons, beads, ornaments) but it is impossible to be directly associated with specific individuals. |
| Τ16/ΣΤ-Μ – IND. G_bone 450 (**CGG_2_023841**)  LH IIIA-LH IIIC EARLY | F/Prime Adult | See above |
| Τ16/ΣΤ-Μ – IND. Η  (**CGG_2_023842**)  LH IIIA-LH IIIC EARLY | F/Middle Adult | See above |
| Τ16/N-Ξ-IND. A_bone 542_Skull AB  (**CGG_2_023843**)  LH IIIA-LH IIIC EARLY | F/Middle Adult | This sample comes from the most complete cranium of the secondary deposit T16/Ν-Ξ (see above).    **Grave goods:** Few ceramic vessels accompanied this deposit, but it is impossible to be directly associated with specific individuals.  **14C**: 3020±15BP, cal. BP 3329-3163 (see Suppl. Tab. S9) |
| T17/Λ_sample 1_bone 557  (**CGG_2_023846**)  LH IIIC EARLY | F/Young Adult | Tomb 17 had a long and continuous use during the entire LHIII period, with total MNI: 19, in various burial deposits. A pit with a large quantity of skeletal remains was located at the *dromos*, in front of the tomb’s entrance (T17/A-K) containing commingled disarticulated bones from at least 15 individuals, with the skeletal material dated in the LHIIIA-B period. Inside the chamber, four primary burials of the LHIIIC period were found: T17/Ξ was first in the sequence, placed in a pit sometime during the LHIIIC Early period or earlier; T17/Λ was the next interment placed on the floor on top of the pit, also dated in the same period based on ceramic evidence; T17/M and finally T17/N followed in LHIIIC Late period.    **Grave Goods:** Two ceramic vessels  This sample comes from the skeleton of the primary burial T17/Λ.  **14C**: 2951±25BP, cal. BP 3207-3003 (see Suppl. Tab. S9) |
| T17/Ν_sample 1_bone 651  (**CGG_2_023849**)  LH IIIC MIDDLE-LATE | M/Prime Adult | This sample comes from the final primary burial of the tomb, T17/N (see above).    **Grave goods:** Four ceramic vessels and some clay buttons  **14C**: 2921±29BP, ca. BP 3164-2965 (see Suppl. Tab. S9) |
| T17/Ξ_sample 1 (**CGG_2_023851**)  LH IIIA2/IIIB | F/Middle Adult | This sample comes from the oldest primary burial of the chamber, placed in a pit. 14C dating places it in the LH IIIA2/LH IIIB period, even though it was relatively dated in the LH IIIC Early as no offerings of earlier periods were found inside the chamber. No grave goods were found  **14C**: 3060±15BP, cal. BP 3347-3214 (see Suppl. Tab. S9) |
| T17/A-K (Λ1) – IND. B_bone 278 (**CGG_2_023853**)  LH IIIA-LH IIIB | M/Old Adult | This sample comes from the pit secondary deposit of the dromos T17/A-K, comprising the earliest burials of the tomb (see above).    **Grave goods:** A pair of bronze tweezers, several ceramic vessels, and some buttons were found in this deposit, but it is not possible to associate them with specific individuals  **14C**: 3135±20BP, cal. BP 3441-3261 (see Suppl. Tab. S9) |
| T17/A-K (Λ1) – IND. C  (**CGG_2_023854**)  LH IIIA-LH IIIB | F/Prime Adult | As above |
| T17/A-K (Λ1) – IND. D  (**CGG_2_023855**)  LH IIIA-LH IIIB | F/Prime adult | As above |
| T17/A-K (Λ1) – IND. E  (**CGG_2_023856**)  LH IIIA-LH IIIB | M/Prime Adult | As above |
| T17/A-K (Λ1) – IND. F  (**CGG_2_023857**)  LH IIIA-LH IIIB | M/Middle Adult | As above |
| T17/A-K (Λ1) – IND. H  (**CGG_2_023859**)  LH IIIA-LH IIIB | M/Middle Adult | As above |
| T17/A-K (Λ1) – IND. I (**CGG_2_023860**)  LH IIIA-LH IIIB | M/Middle Adult | As above |
| Τ20/Β-Δ (Λ1)_bone 173  (**CGG_2_023868**)  LH IIIB | M/Adult | Tomb 20 contained only two burial assemblages: The secondary deposit of disarticulated commingled bones in a pit (T20/Β-Δ, MNI: 9), with the skeletal remains dated in the LH IIIB period and comprising both sexes, adults and sub-adults; and the male primary burial Τ20/Α, dated in the LH III Early period.  This sample comes from the pit secondary deposit of LH IIIB remains.    **Grave goods:** One ceramic vessel, bronze tweezers, buttons, and beads were found in this deposit, but it is not possible to associate them with specific individuals  **14C:** 3070±15BP, cal. BP 3355-3225 (see Suppl. Tab. S9) |
| Τ20/Β-Δ (Λ1)_bone 203/250  (**CGG_2_023869**) | F/young adult | As above |
| Τ22/Α-Β (Λ1)_bone 352  (**CGG_2_023875**)  LH IIIA-LH IIIC EARLY | ?/Adult | Tomb 22 was used continuously during the entire LH III period. The tomb included a MNI: 15, placed in various burial deposits. A pit at the end of the *dromos* (T22/A-B) contained the disarticulated remains of several burials (MNI: 11) dated from LH IIIA1-LH IIIC Early. The majority of these bones should be dated in the palatial period (LH IIIA-B) based on ceramic evidence. Inside the chamber, four primary burials of the LH IIIC Late/Sub-Mycenaean period were placed on the floor (T22/Γ was the first of this series of burials, followed by T22/Δ, T22/E and T22/ΣΤ).  The sample in this study comes from the secondary pit deposit, and thus the most probable dating is that of the LH IIIA-B period.    **Grave goods:** Several ceramic vessels of the LH IIIA and LH IIIB, one of the LH IIIC, clay and steatite buttons, as well as carnelian beads were found in this deposit, but it is not possible to associate them with specific individuals. |
| T24/A-E4_bone 257 (**CGG_2__2_023877**)  LH IIIA or LH IIIC | F/Middle Adult | Tomb 24 presented evidence of use in LH IIIA and LH IIIC, but not in LH IIIB. Skeletal remains were found in two contexts inside the chamber: a large pile of commingled disarticulated remains (T24/A) at the east with MNI: 6, including four adults, one child and one adolescent, which was accompanied by artefacts of both LH IIIA and LH IIIC period; and a smaller pile of a single secondary deposit of a male middle adult (T24/B) at the west, also accompanied by vessels of both periods.  The sample in this study comes from the large secondary deposit T24/A-B.    **Grave goods:** Several ceramic vessels of both LH IIIA and LH IIIC date were found in this deposit, together with steatite buttons, beads and a bronze pin. It is not possible to associate the grave goods with specific individuals |
| Τ28/Β-Ζ (Λ1) _bone 179 (**CGG_2__2_023883**)  LH IIIA | F/Adult | Tomb 28 presented evidence of use in LH IIIA and the LH IIIC Middle and Late periods. The chamber contained no primary burials, only two secondary deposits: one pit (T/28-B-Z) with various commingled remains (MNI: 13, including 9 adults of both sexes, 1 child and 3 infants), dated in the LH IIIA1 period; and a small bone assemblage in the form of a pile on the floor (T28/A) with grave goods of the LH IIIC Middle/Late. Bone remains from the latter were not recovered, so the osteological study concerns only the contents of the pit, including all samples in this study.    **Grave goods:** Three ceramic jars, clay buttons, and various beads were found in this deposit, but it is not possible to associate the grave goods with specific individuals.  **14C**: 3060±28BP, cal. BP 3360-3177 (see Suppl. Tab. S9) |
| Τ28/Β-Ζ (Λ1) – IND. B_sample 1_bone 139  (**CGG_2_023884**)  LH IIIA | F/Child (7-8y) | See above  **14C**: 3213±27BP, cal. BP 3466-3375 (see Suppl. Tab. S9) |
| Τ28/Β-Ζ (Λ1) – IND. D_bone 41 (**CGG_2_023887**)  LH IIIA | F/Infant  (8-12 months) | See above |
| Τ40/Δ-Ι_bone 1128 (**CGG_2_023899**)  LH IIIA | F/Adult | Tomb 40 was used exclusively during the LH IIIA period. It contained a very large quantity of commingled disarticulated human remains (T40/Δ-Ι) with MNI: 14, including two infants, one child, and 11 adults of both sexes. Three primary burials (T40/A, T40/B, T40/Γ) of adult females, dated in the same period, were also placed next to the large bone mass and partially mixed with it. The sample in this study comes from the large secondary deposit (T40/Δ-Ι).    **Grave goods:** Several grave goods accompanied this secondary assemblage (various ceramic vessels, a bronze dagger, clay buttons, beads, metal ornaments and jewels); however, it is not possible to associate any of them directly with specific individuals. |
| Τ42/Α_sample 2 (**CGG_2_023902**)  LH IIIC MIDDLE-LATE | M/Young Adult | Tomb 42 presented evidence of use in the LH IIIA2 and the LH IIIC Middle and Late periods. The tomb included various burial deposits, both in the chamber and the *dromos*. At the *dromos*, a pit with commingled secondary remains (T42/Pit II) from at least two adults (male and female) was found but could not be relatively dated due to the lack of datable grave goods. However, radiocarbon dating of the bones dates the contents in the LH IIIA period. Inside the chamber, one intact (T42/A) and one disturbed (T42/B) primary burials, and a single secondary deposit (T42/Γ) of the LH IIIC Middle/Late period were found, as well as a pit with a large number of commingled, disarticulated bones (T42/Β-Δ), dated to the LH III period (both by relative and absolute dating). The pit’s MNI was 10, including one infant and 9 adults of both sexes.  This sample comes from the LH IIIC primary burial T42/A.    **Grave goods:** Various ceramic vessels, a bronze knife, and a clay button. |
| Τ42/Γ_sample 1_bone 172  (**CGG_2_023903**)  LH IIIA2/IIIB | F/Young Adult | This sample comes from the single secondary deposit T42/Γ (see above) that was dated to the LH IIIC period based on the grave goods associated with it. However, the radiocarbon dating suggests an earlier date (LH IIIA2/IIIB), which is compatible with the secondary character of this assemblage.    **Grave goods:** A stirrup jar of the LH IIIC Late period, and a steatite button were associated with this context, and thus the human bones were presumably dated in that period. However, this is not confirmed by the radiocarbon date  **14C**: 3100±20 BP, cal. BP 3375-3244 (see Suppl. Tab. S9) |
| Τ42/Δ-Θ (Λ1) –IND. E_Skull 5  (**CGG_2_023909**)  LH IIIA | F/Middle Adult | This sample comes from the secondary pit deposit of the chamber (T42/Δ-Θ), whose contents were dated in the LH IIIA period (see above). |
| Τ42/Δ-Θ (Λ1) –IND. Η_bone 575  (**CGG_2_023912**)  LH IIIA | F/Infant (6  months) | As above |
| Τ42/Λ2 - ΙΝD. Α_bone B151  (**CGG_2_023913**)  LH IIIA | M/Prime Adult | This sample comes from the *dromos* pit secondary deposit (T42/Pit II, see above).    **Grave goods:** Only few sherds of indeterminate date    **14C**: 3120±20BP, cal. BP 3390-3253 (see Suppl. Tab. S9) |
| Τ42/Λ2[IM18] -ΙΝD. Β_bone B128  (**CGG_2_023914**)  LH IIIA | F/Adult | **14C**: 3032±31BP, cal. BP 3351-3150 (see Suppl. Tab. S9) |
| Τ75/Bur. layer B/ Skull Γ_sample 1  (**CGG_2_023920**)  LH III? | M/Adult | This tomb is, together with Tomb 4, the largest in size in the Voudeni cemetery, showing evidence of use in all phases of LH III. The tomb is not fully studied yet, so detailed information on its contexts is lacking.    **Grave goods: ?** |
| Τ75/Bur. layer B/ Skull E  (**CGG_2_023922**)  LH III? | F/Adult | As above  **Grave goods:?**  **14C**: 3080±15BP, cal. BP3360-3236 (see Suppl. Tab. S9) |

Table 7. Synoptic table with data about the sampled individuals from Voudeni and their context of origin.

##### ***2.5.6. Eleon, Boeotia, Central Greece***

Latitude: 38.355898, Longitude: 23.48093

Sample provider: Brendan Burke, Bryan Burns, Nick Herrmann

Brendan Burke

The chief goal of the most recent excavations at ancient Eleon in central Greece, from 2015–2018^101^, was the exploration of the Late Helladic I (c. 1700 BCE) funerary construction we call the Blue Stone Structure (BSS). The BSS (Fig. 7) contains both Individual tombs and constructions. They are grouped into two major phases: the period of tomb use and perimeter wall enclosure (left), and the subsequent period when stone pavements and a clay tumulus closed off access to tombs below.

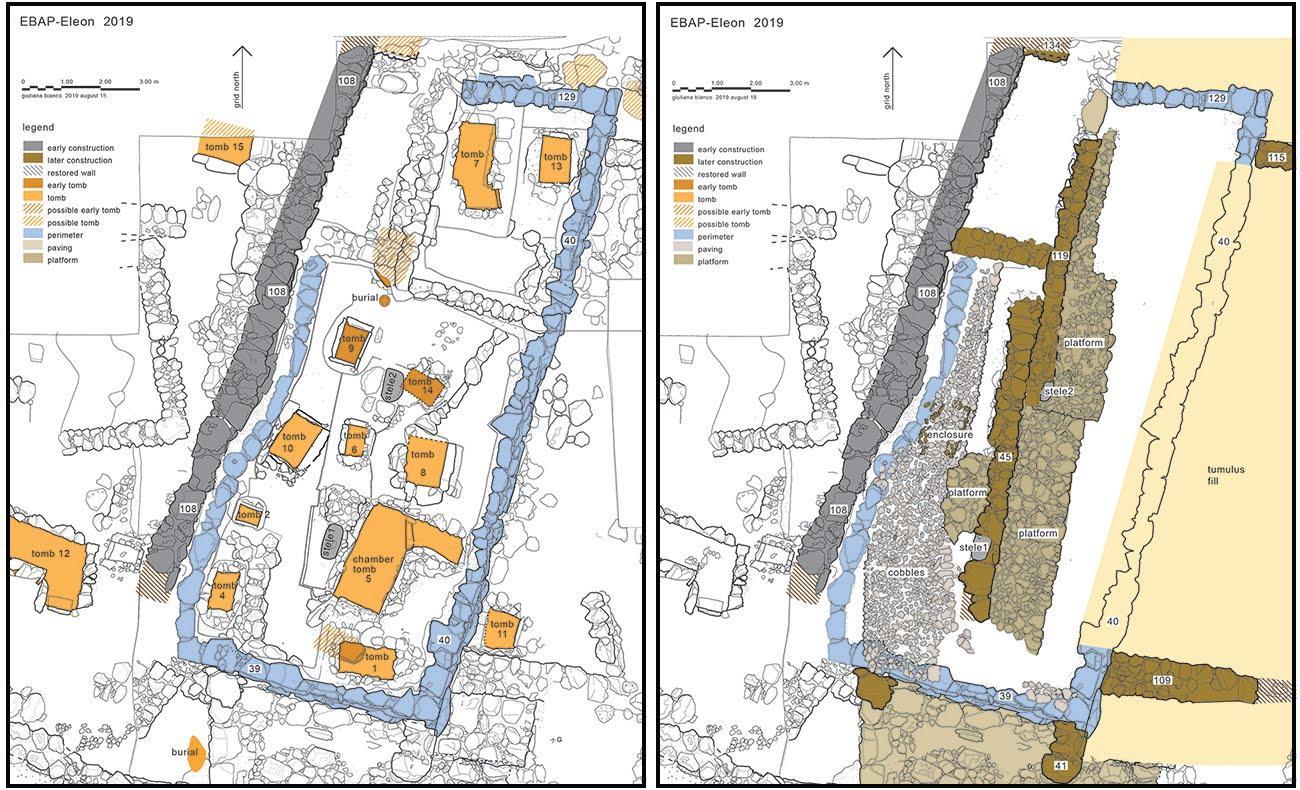

Fig. 7. The Blue Stone Structure – Early Mycenaean c. 1700-1600 BCE (courtesy of Brendan Burke)

All of the dental and osseous samples bones come from burials associated with the BSS, which is an enclosure, demarcating a space for the dead apart from the living, rather than a freestanding structure. Three joining walls make the west, south, and east sides of a rectangular enclosure measuring 10 m across its southern end and extending for at least 17 m north–south. The north side is less well defined. The enclosure consists of stone walls around a select group of Early Mycenaean burials under a low mound of earth and clay. Within its boundaries, numerous tombs were indicated by rings of stone around the grave cuttings; cobbled surfaces were laid at several elevations in fill above the stone marking the tombs; and two stelae, roughly carved markers about 1.3–1.5 m high, stood in situ. The recovery of human remains, directed by Herrmann, followed a protocol to excavate, record, map, and identify individual bones as they were removed regardless of the conditions of the bones. After burials ceased, the subsequent creation of a tumulus over the BSS further unified the monumental complex.

Within the confines of the BSS side walls, we found well-laid cobblestone surfaces at varying levels but not directly above individual tombs. These well-built surfaces inhibited access to the tombs and perhaps served as platforms for activity focused on the dead sealed below. In the west half of the BSS, a long, narrow stretch of cobblestone paving was preserved above Tombs 2, 4, 9, and 10; in the east half, similar platforms were built of larger stones mainly above Tombs 5, 8, and 14.

Within the BSS, we have excavated 11 built tombs of diverse construction types ranging from a small clay cist (Tomb 2) to a large built chamber tomb (Tomb 5). Most of the other tombs are a single chamber (cist) formed by drystone walls and covered by one or more large cover stones. The BSS is a significant burial enclosure within what may be a larger Middle Helladic to LH I cemetery occupying the northeast edge of the acropolis. Two burials (Tomb 12 in SWA1c and Tomb 15 in NWA1d) were located west of the BSS. Both graves appear to be L-shaped tombs and are among the largest of all tombs discovered at the site. Along the exterior eastern wall of the BSS, we found Tomb 11, a stone-lined ossuary containing the remains of at least 30 individuals—the largest number in any tomb—and the greatest number of grave goods, despite it being one of the smallest tombs in size. Taken together, Tombs 11 and 12 suggest the existence of a larger cemetery that likely extended all the way to the clay cist burial recovered in NWB1b, if not farther. The location of the BSS within a larger, earlier cemetery would follow the example found at Mycenae of Grave Circles A and B near the earlier Middle Helladic cemetery.

The samples presented in this study come from Tomb 2, 9 and 13.

**Tomb 2** was a small stone cist with the single inhumation of a child, 2-4 years old (**CGG_2_105016**) buried without grave goods^101^.

**Tomb 9** was a stone cist with the single inhumation of a young adult buried with a murex shell. Under the individual a small pit was discovered containing the disarticulated remains of an adult individual (**CGG_2_105024**) without grave goods^101^, which was sampled for this study.

**Tomb 13** predates the eastern perimeter of BSS and it was included intentionally in it. Three individuals were found lying on the floor of the context. Under them two pits contained the commingled remains of three more individuals^101^. This tomb, which is dated to Late Helladic I (c. 1700-1600 BCE), contained six individuals; two of them (**CGG_2_105030,** [IND3] and **CGG_2_105031,** [IND 5]**)** were sampled.

#### **2.6. Hungary**

##### ***2.6.1. Pitvaros, Csongrád County***

Latitude: 46.331880; Longitude: 20.735700

Sample provider: György Pálfi, John M. O’Shea, Györgyi Parditka

John M. O’Shea, Györgyi Parditka

The cemetery of Pitvaros is located on the northern edge of the village of Pitvaros, along the Pitvaros-Ambrózfalva road, roughly 48 km east of Szeged. The site occupies a sandy stream levee and was discovered during the course of sand quarrying. Móra Ferenc visited the site on 2 August, 1926. Six graves had already been unearthed. The grave goods from each had been kept separate and were turned over to Móra. Since the site was in imminent danger, excavation began immediately and continued through to the 31st of August. In total, the excavations produced 49 graves, 43 that could be attributable to the Bronze Age, and six to the migration period. The Pitvaros cemetery represents the first instance where the full Maros funerary program of body placement and artefact accompaniment is observed^102^.

The primary published source on the Pitvaros excavations is Bóna^103^. Bóna employed both the field diary from the excavation and the museum catalogue in his descriptions. The finds are also listed in Banner's summary of the Maros group cemeteries^104^. In his synthesis Bóna concluded that the site falls chronologically into the Early Bronze Age (chronology confirmed by 14C), based on the ceramic assemblage which shares affinities with EBA Nagyrév pottery, and that the individuals interred at Pitvaros had migrated into the region from the lower Danube^103^.

Sources of information on the skeletal remains include field assessments of skeletal age, which distinguished adults from sub-adults in 37 instances^103^, and formal osteological analyses conducted by Farkas^105,106^ and Rega^107^. However, only 14 Bronze Age crania survived for modern assessment, while postcranial elements were preserved from only two cases.

**Pitvaros, grave 3**

Burial 3 contained the skeleton (**CGG_2_104049**) of an adult female in a flexed posture, however no information is available regarding the orientation of the deceased as the grave was excavated by workmen. A small ceramic bowl was deposited in the grave. The sample has been 14C dated and produced a date to 3615±20BP, cal. BP 3980-3849 (see Suppl. Tab. S9).

**Pitvaros, grave 15**

Burial 15 contained the flexed skeleton (**CGG_2_104061**) of an adult male that was placed on its right side, which is contrary to the gender based system of body placement and orientation. The individual was interred with a necklace of faience beads, two copper bracelets, and with a medium sized bowl containing animal remains. The sample has been 14C dated and produced a date to 3645±15BP, cal. BP 4075-3897 (see Suppl. Tab. S9).

**Pitvaros, grave 29**

Burial 29 contained the flexed skeleton (**CGG_2_104071**) of a mature male, placed on his left side, and oriented NE-SW. No grave goods were recorded from this burial. The sample has been 14C dated and produced a date to 3695±15BP, cal. BP 4090-3976 (see Suppl. Tab. S9).

##### ***2.6.2. Szőreg C cemetery, Szeged, Csongrád County***

Latitude: 46.21873, Longitude: 20.19886

Sample provider: György Pálfi, John O’Shea, Györgyi Parditka

John O’Shea, Györgyi Parditka

The Szőreg C cemetery is located in the village of Szőreg, immediately to the south and east of Szeged in southeastern Hungary, and is the closest of the Maros cemeteries to the Tisza-Maros confluence. The site was excavated by F. Móra between March 1928 and October 1931^102,104,108,109^. Approximately 230 graves were unearthed during this time and Foltiny estimated there may have been an additional 330-400 graves remaining to be recovered^108^. The first and most complete bio-anthropological analysis of the material was conducted by Gy. Farkas ^106^, with additional analyses conducted by Rega^107^ and Nagy^110^. There is overall good agreement among all these sources. When significant differences among the determinations are observed, the agreed determination of two of the three investigators is given. The ceramics from Szőreg appear to span the entire Maros sequence, except for the earliest Early Bronze Age varieties observed at Pitvaros^103,111^, which extends from the Early Bronze Age to the end of the Middle Bronze Age in Hungary. In calibrated radiocarbon years the cemetery appears to have been used continuously between about 2100 BC to about 1600 BCE^112^.

**Szőreg, grave 33**

Burial 33 contained the flexed skeleton (**CGG_2_104001**) of a mature man. The body was placed with the head to the north and was interred without grave goods.

**Szőreg, grave 70**

Burial 70 is the grave of an adult female (**CGG_2_104154**). The woman was interred in a flexed posture on her right side with her head to the South, such that she faced east. She was interred with a bone needle and three ceramic vessels; a bowl and two one handled cups. The burial was 14C dated in an earlier study (sample Oxa-31099) to 3716 ± 32BP^112^, cal. BP 4152-3934 (see Suppl. Tab. S9).

**Szőreg, grave 74**

Burial 74 is the grave of an adult male (**CGG_2_104164**). The man was interred in a flexed posture on his left side with his head to the East, such that he faced south. Although the side of placement for the man fits the normative Maros pattern of grave geometry, the East-West orientation represents an alternative to the normative body positioning. The man was interred with three ceramic vessels, a large bowl, a small biconical beaker, and a larger jug of Nagyrév type. All these ceramics are characteristic of the Early Bronze Age phase of site use.

**Szőreg, grave 80**

Burial 80 is the grave of an adolescent of undetermined sex (**CGG_2_104167**). The body was placed in a flexed posture on its right side, but no other information is present on its orientation or facing. Given the side of placement the adolescent may have been a female. The adolescent was interred with two ceramic vessels, a bowl and a pitcher, both of Early Maros style.

**Szőreg, grave 89a**

Grave 89 contained the remains of two individuals. Individual 89a is an adult female that was placed in a flexed posture on her right side, with her head to the south and facing east. Individual 89b was the remains of a child. The grave contained a baroque style late phase pitcher and the fragments of a bowl. The sample taken for this study belongs to the adult woman (**CGG_2_105421**).

**Szőreg, grave 97**

Grave 97 contained the skeleton of an adult woman (**CGG_2_105413**). The skeleton was placed on the right side but with the legs spread apart. No ceramics were found with the burial, but the grave did contain a bone fastener that functioned like a belt buckle.

**Szőreg, grave 125**

Grave 125 belonged to a child (**CGG_2_104132**). The body was placed on its right side, but no other information is present on its orientation or facing. The child was buried with three ceramic vessels; a jug, a bowl and a late Szőreg (Phase 5) mug^111^.

**Szőreg, grave 147**

Grave 147 contained the remains of an adult male (**CGG_2_105451**). The body was placed in a flexed posture on the right side, but was oriented with the head to the north, resulting in the body facing west. The grave contained a cup, a small bowl, and a baroque style late phase pitcher. The grave was radiocarbon dated in an earlier study (sample OxA-30989) to 3402 ± 34BP^112^, cal. BP 3821-3514 (see Suppl. Tab. S9).

**Szőreg, grave 156**

Grave 156 was represented by the skeleton of an adult male (**CGG_2_105409**). The skeleton was placed in a flexed posture on its right side, but without further information on its orientation or facing. The grave contained a late phase cup and a bone fastener that was located near the hips and functioned like a belt buckle.

##### ***2.6.3. Tápé – Széntéglaégető cemetery, Szeged, Csongrád County, Hungary***

Latitude: 46.27511; Longitude: 20.19980

Sample provider: György Pálfi, John O’Shea, Györgyi Parditka

John O’Shea, Györgyi Parditka

The Tápé-Széntéglaégető cemetery is located in southeast Hungary, on the west side of the River Tisza, in close vicinity of the Tisza-Maros confluence. Close to 700 graves were excavated in the 1960s by Ottó Trogmayer. The site documentation with initial evaluation of the data and bio-anthropological analysis was published in 1975^113,114^. In 2015 and 2016 a new macro-morphological analysis of the skeletal material was conducted by Olga Spekker (Spekker n.d.). There is a good general agreement between the two determinations, but when there are significant differences, the more recent determination is reported. In several instances there are inconsistencies in the published orientation of the individual graves. These have been corrected in the descriptions that follow.

The cemetery was assigned to the Reinecke BB2 – BC periods, based primarily on metal artefacts, and was associated with the Tumulus culture by the excavator^113^. While different authors argued for various use-life periods for Tápé^115^, recent absolute dating of the cemetery suggests a range of 1501 BCE to 1233 BCE^112^.

The metal finds in the cemetery represent classical items of the so-called Tumulus type, such as the *Petschaftkopf* pins. In case of the ceramic finds more variation was noted, including some connections to the local Middle Bronze Age traditions, as well as similarities with the Slovakian Carpathian Tumulus Culture. Trogmayer argued however that Tápé represented a distinct cultural entity.

Tápé has been the subject of numerous studies focusing on both archaeological and bio-anthropological questions^115–123^. Among the archaeological studies^116,121,123^ considered Tápé in depth.

**Tápé-Széntéglaégető, grave 157**

Inhumation burial. The skeleton of a mature male was placed in a flexed posture on its right side. The orientation of the grave was NE-SW. The individual (**CGG_2_103950**) was buried with a cup, a bronze awl with a bone handle and a bronze blade (?) fragment.

**Tápé-Széntéglaégető, grave 168**

Inhumation burial. The body of an adult female was placed in a flexed posture on its right side. The orientation of the grave was SE-NW. The individual (**CGG_2_103916**) was buried with a pot, a bowl and a cup.

**Tápé-Széntéglaégető, grave 209**

Inhumation burial. The body of an adult male was placed in a flexed posture on its right side. The orientation of the grave was SW-NE (212-32°). The grave orientation was rectified. The body (**CGG_2_103834**) was interred without grave goods.

**Tápé-Széntéglaégető, grave 250**

Inhumation burial. The upper body of a mature woman was lying on the stomach. The legs were in a flexed position, on the left side. The orientation of the grave was E-W. The individual (**CGG_2_103857**) was buried with three mugs, a unio shell, a piece of stone, two rings of bronze found close to the skull, 7 pierced sea shells and an amber bead. The burial was 14C dated in an earlier study (sample UGAMS-30831) to 3170±25BP^112^, cal. BP 3450-3355 (see Suppl. Tab. S9).

**Tápé-Széntéglaégető, grave 333**

Inhumation burial. The upper body of an adult male was lying on its back. The legs were flexed, lying on the right side. The orientation of the body was SE-NW. The individual (**CGG_2_103901**) was buried with a cup and one seal-headed pin (*Petschaftkopf*).

**Tápé-Széntéglaégető, grave 346**

Inhumation burial. The body of a juvenile female was placed in a flexed posture on its right side. The orientation of the body was N-S. The individual (**CGG_2_103882**) was buried with a wire ring.

**Tápé-Széntéglaégető, grave 354**

Inhumation burial. The body of an adult woman (**CGG_2_103822**) was placed in a flexed posture on its left side. The orientation of the body was S-N. A wire ring, a bronze spiral, the tip of a pin-like object and the fragments of a mug were found in the grave.

**Tápé-Széntéglaégető, grave 491**

Inhumation burial. The body of a young adult male (**CGG_2_103928**) was placed in a flexed posture on its left side. The orientation of the body was E-W. A small pedestaled vessel with a lid was included in the grave. The burial was 14C dated in an earlier study (sample UGAMS-30836) to 3060±20BP^112^, cal. BP 3355-3210 (see Suppl. Tab. S9).

**Tápé-Széntéglaégető, grave 534**

Inhumation burial. The body of a mature man was placed on its back, in an extended posture. The orientation of the body was E-W. The individual (**CGG_2_103885**) was buried with a dagger. In addition, multiple bronze tutuli were found in the burial that were interpreted as relating to the presence of a belt . The burial was 14C dated in an earlier study (sample UGAMS-236660) to 3187±24BP^112^, cal. BP 3451-3370 (see Suppl. Tab. S9).

**Tápé-Széntéglaégető, grave 685**

Inhumation burial. The body of an adult male was placed in a flexed posture on his right side. The orientation of the grave was SE-NW. The individual (**CGG_2_103942**) was interred without grave goods.

#### **2.7. Italy**

##### ***2.7.1. Coppa Nevigata, Foggia province, Puglia***

Latitude: 41.558106; Longitude: 15.833939

Sample provider: Giulia Recchia, Alberto Cazzella, Mary Anne Tafuri

Serena Sabatini, Giulia Recchia

Coppa Nevigata is a fortified long-lasting settlement^124^, located on the Adriatic coast just below the Gargano promontory, which was continuously occupied from the 18th to the 8th century BCE. Unlike the coeval settlements in southern Italy it has yielded a number of human bones (more than 300). These mostly come from different 15th century BCE layers and belong to three main contexts:

- Two formal burials located each on a different postern of the predating 17^th^ century BCE fortification wall. The sample CN15 (**CGG_2_100648**) analysed for this project belongs to one of these formal burials.
- A cluster of human bones (chiefly finger bones and other small bones) located in a space in between the two above mentioned posterns with burials.
- Scattered human bones (chiefly mandibles, teeth and long bones) found in the filling (made of soil and crushed limestone) of the 15^th^ century BCE defensive line. The sample CN 11 (**CGG_2_100647**), belongs to this group of finds.

Furthermore, a few scattered human bones come from both earlier and subsequent layers of the settlement. In particular, some bones of infants come from the 16^th^ century BCE layers. Sample CN7 (**CGG_2_100646**) belongs to the Early Subapennine (13^th^ century BCE) layers. CN7 has been radiocarbon dated to 3095±20BP, cal. BP 3371-3241 (see Suppl. Tab. S9).

##### ***2.7.2. Lavello, Potenza province, Basilicata***

Latitude: 41.06364028318502; Longitude: 15.819151268308177

Sample provider: Alessandro Canci, Mirella Cipolloni Sampò

Serena Sabatini

Grave 743 is a large hypogea with several rooms, whose complex design cannot be fully reconstructed as it has been partly destroyed by looters. At Lavello there were several deposition layers and traces of ritual activities^125^. The latter took place in the dromos leading to the grave. The latest find from the grave is a typical Final Bronze Age fibula. The earliest material seems to be dated to the Early Middle Bronze Age (c. 1650-1550 BCE). The grave had been disturbed by looter and not all the depositions may have been preserved when the excavations took place, c. 50 individuals were recovered, the sample analysed in this study is deposition 53 (**CGG_2_022564/Rise 1702**), from layer 6, anthropologically determined as a male individual of c. 25 years.

##### ***2.7.3. Lucone, Brescia province, Lombardy***

Latitude: 45.548241; Longitude: 10.494240

Sample provider: Alessandro Canci

Serena Sabatini

The individual (**CGG_2_022565/Rise 1703**) analysed within the present project is an infant c. 2.5-3 years old. The bones were dated to 1967±10 BCE by means of dendrochronology. The individual was found in the settlement settings of the so-called Lucone D. Specific parts of this skull were accurately deposited right above a destruction layer (destruction caused by fire, which probably destroyed most of the village). The destruction phase is dated to the Italian Early Bronze Age (c. 2150-1650 BCE). The bones were covered with a layer of bark stripes. Both the bones and the bark were not affected by fire and clearly deposited some time after the destruction. C. 9 m from this deposition, a few long bones of an individual of the same age (the same as the skull?) were also found together with animal bones of various kinds. There is little doubt that the skull was the object of a ritual of some sort. The way the bones of the skull were found suggest that soft tissue of the head was not completely dissolved when it was separated from the rest of the body. Different possible scenarios for the ritual or sacrifice that took place there have been proposed on the basis of the osteological evidence. Skull-related cults are known in the broader Mediterranean and Near Eastern world and attested in other Neolithic and Early Bronze age lake dwellings of the Alpine region^126^.

##### ***2.7.4. Mereto, Udine province, Friuli- Venezia Giulia***

Latitude: 46.041417; Longitude: 13.05617

Sample provider: Alessandro Canci, Elisabetta Borgna, Susi Corazza

Serena Sabatini, Susi Corazza, Elisabetta Borgna

The grave at Mereto is a complex monument^127^. The final stage of the monument - as it appeared to the excavators in the first place - is a large tumulus. It is important to say that it was well-preserved with a height of 6.5 m and a diameter of 25-26 m. It was carefully excavated and revealed a complex biography^127^.

A single inhumation grave was at the bottom of the monument and dated to a later part of the Italian Early Bronze Age (c. 2150-1650 BCE). The monument consisted of a large stone platform that was progressively constructed over the tomb and likely marked its presence in the landscape^127,128^. Various activities of apparently ritual character took place on the monument and the monument grew in connection to them well into the advanced Middle Bronze Age. At the end, a large mound in which layers of gravel and clay alternate covered the whole platform. Traces of wooden structures of some sort could be also found in the upper stratigraphy of the mound. The grave was, however, not the first structure present in the area, micro-morphological analyses, fragmentary finds and sparse human remains demonstrate that ritual activity of some sort took place at the same location before the inhumation was laid down^54^.

The individual (**CGG_2_022253/Rise 1503**) was in a supine position. The careful excavation of the bones has given the possibility to reconstruct the way in which the body was deposited^128^. The unnatural position of the shoulder and of the legs, and the position of the head and the jaw suggest that the body, including the head, was maybe put in a coffin and also wrapped in something of organic origin, which kept the bones in place even after the decomposition of the soft tissues. The anatomic position of the feet also suggests that the body was wearing shoes or rather boots too. The individual is a young adult (c. 16-19 years) probably male and had no grave goods with the exception of two stone tools.

Several 14C dates have been gained for the monument^129^. The bones of the individual from the grave have been dated to 3450±40BP, cal. BP 3832-3583 (see Suppl. Tab. S9).

##### ***2.7.5. Narde, Rovigo province, Veneto***

Latitude: 45.027941657128856; Longitude: 11.656466525667486

Sample provider: Claudio Cavazzuti

Serena Sabatini

The necropolis at Narde is a large cemetery likely belonging to the Final Bronze Age settlement of Frattesina di Fratta Polesine. Around this large and complex settlement two different burial grounds have been discovered so far. One is located c. 500 m south-east of the site and it is called the cemetery of Fondo Zanotto, while c. 500 m north of the site there is the Narde cemetery, which is divided in two sectors Narde I and II. Fondo Zanotto has c. 150 depositions^130^, while at Narde I and II over 900 depositions have been recovered. The chronology at the Narde cemetery spans from the Final Bronze Age to the beginning of the Early Iron Age.

An extensive study of the Narde graves suggests a sensible difference as to health and life expectancy between the deceased buried in Narde I and those from Narde II. The individuals from Narde II seem to have had worse living conditions than those buried in the Narde I^130^. The analyses of the grave goods seems to confirm a wealthier status (in the sense of larger numbers of grave goods and of artefacts made with more precious materials) for the individuals from Narde I.

The vast majority of the graves are cremation burials^131^. Of the c. 600 burials from Narde I only 3 are inhumations, while out of c. 300 burials from Narde II, the inhumation were only 18. This is a very important factor to keep in mind when assessing the social and cultural status of those individuals. Those graves have no grave goods and with the exception of some of them, that appear to be in particular relation to the cremation graves it is not possible to date them with certainty. It is supposed that those individuals might have been of a very low social class or slaves. Two individuals were sampled for this project: the individual from grave 23 (**CGG_2_022895/Rise 1882**), which was an adult male of c. 40-50 years, and the one from grave 59 (**CGG_2_022897/Rise 1884**), which was also an adult male, but of c. 30 years. The genetic sex of both individuals resulted also to be male. Due to the position of the skeleton it is argued that the individual in grave 59 was likely wrapped in a sort of textile at the time of the burial: in the grave a bronze fragment and a ceramic shard were also recovered^131^.

##### ***2.7.6. Olmo di Nogara, Verona province, Veneto***

Latitude: 45.176747; Longitude: 11.055126

Sample provider: Alessandro Canci, Luciano Salzani

Serena Sabatini

Olmo di Nogara is the name of a large bi-ritual cemetery close to the modern town of Nogara in northern Italy. It lies on the right bank of the Tartaro River^59^. It was excavated through an intense series of campaigns in the 1980s and 1990s, when the majority of the known depositions were unearthed^59,132^. The last excavation took place in 2009 returning a limited number of new graves^133^. All in all, 471 inhumation graves and 62 cremations have been documented at Olmo di Nogara. The chronology of the site goes approximately from the Middle Bronze Age IIA to the Recent Bronze Age 1 (c. 1550-1225/1200 BCE). The graves were at times well preserved and furnished in many cases with grave goods such as bronze swords, bronze pins, jewellery, amber beads and other materials, which the community at Olmo likely received from long-distance trade and exchange. The characteristics of the skeletal remains show that several individuals had survived violent conflicts or died violently suggesting that warfare was a steady presence in the life of the community^134^.

The site lies in a north-west/south-west axis following the river morphology. Overall, one can say that the cemetery had a complex topography. It was delimited by a sort of funerary road, which probably also represented the access to it; to the east of the documented graves and along the entire length of the Area C, two small V-shaped ditches were found six metres apart from each other. They have been interpreted as the remains of two palisades, which might have flanked/defined the funerary road to the site^59^.

Four different large parts of the cemetery were excavated. The lots are close to each other, but separated by parts of land, which have not been investigated mostly due to the presence of existing constructions, thus the original number of tombs was likely much larger. The northernmost lot is called area A, followed (going south) by area B, Area C and finally area D.

Area A is the northernmost investigated part of the cemetery. It has a surface of c. 2100 m^2^. Differently from the other ‘areas’, it includes a sort of circular structure surrounded by a narrow ditch. The structure was unfortunately badly damaged by agricultural works and no information seems available as to its use. One grave was found in the south-west part of the ditch. From this area, 7 individuals have been analysed; they come from graves: 3-5, 7, 9-10, and 15-16

Area B is in-between modern buildings and roads. The investigated surface is of c. 4000 m^2^. Adjustments of the area during different historical periods have damaged part of the contexts. From this area, 17 individuals have been analysed (Tab. 8); they come from graves: 388, 392, 401, 404, 408, 412, 417, 423, 430, 433, 439, 459, 462, 471, 478, 493, 499.

Area C is located in between area B and D. It has a surface of c. 6240 m^2^. There are abundant traces (dated to many different historical periods) of recurrent use and occupation of the area including building activities and agricultural work; some parts of the area were heavily damaged. A general(?) levelling of the area at the end of the 19th century must have destroyed and/or damaged a large number of incinerations and of children burials, which are normally at a lower depth than those of the other inhumations. From this area, 49 individuals have been analysed (Tab. 8); they come from graves: 28, 30, 32, 43-45, 53, 57-58, 104-105, 110-111, 118, 127, 133, 136-137, 159, 180, 183-184, 197, 199-200, 207, 211, 213, 224, 226, 228, 230, 234-235, 274, 282, 287, 289, 293, 296-297, 301, 313-314, 318, 321, 324, 362, 372. In the main dataset we also have CGG_2_22658 (OdN 45A), although the sample is from Olmo, but the context is not certain. One grave (OdN209) from Area C was archaeologically dated to the Early Middle Age. Two of our samples (see below) from Area C have also been radiocarbon dated to the same period, suggesting that area C was sparsely used as a cemetery during Late Antiquity/Early Middle Age; one should therefore not exclude the possibility that some of the contexts without grave goods might potentially belong to a later period than Bronze Age.

Area C west. In 2009, a new rescue excavation was carried out unveiling another part of the so-called funerary road and more graves. Being this new area close to Area C it has been called Area C north-west^133^. For this area, four graves have been analysed (Tab. 8): 541, 543-544, 547

Area D is the southmost area, it has a size of c. 7500 m^2^. None of the graves from this area seems to pertain to the Bronze Age. The contexts from Area D have been dated to the Chalcolitic and to the Roman period^59^.

| **OLMO DI NOGARA - AREA A** | | | |
| --- | --- | --- | --- |
| **Grave nr** | **Project ID** | **SEX/AGE** | **Context** |
| 3 | CGG_2_022657  Rise1795 | F?/  14±4 years | Supine body positioned E-W; head to the right/north. No grave goods^59^**.**  **Chronology**: MBA3? |
| 4 | CGG_2_022567  Rise 1705 | F?  7±2 years | Supine body positioned SE-NW; head to the left/south. Two fragments of a bronze pin and 3 of bronze ring, a bronze pendant, bottom of a vase (Salzani 2005, 22). Archaeologically suggested date: MBA 3-RBA^59^  **14C**: 3135±15BP, cal. BP 3396-3268 (see Suppl. Tab. S9). **Chronology**: MBA3 |
| 5 | CGG_2_022568  Rise 1706 | F?  12±3 years | Supine body positioned E-W; head straight. There were apparently no grave goods^59^.  **14C**: 3155±15BP, cal. BP 3445-3350 (see Suppl. Tab. S9). **Chronology**: MBA2B-3A |
| 7 | CGG_2_022569  Rise 1707 | M  15±3 years | Supine body positioned E-W; head straight. There were apparently no grave goods^59^.  **14C**: 3040±20BP, cal. BP 3340-3170 (see Suppl. Tab. S9). **Chronology**: MBA3A-RBA 1 |
| 9 | CGG_2_022570  Rise 1708 | -  4±1 years | Supine body positioned E-W. Grave damaged by ploughing, no grave goods^59^.  **14C**: 3115±20BP, cal. BP 3387-3251 (see Suppl. Tab. S9). **Chronology**: MBA3 |
| 10 | CGG_2_022571  Rise 1709 | M?  7-8 years | Body positioned SE-NW. Grave damaged by ploughing, no grave goods^59^**.**  **Chronology**: MBA3? |
| 15 | CGG_2_022572  Rise 1710 | -  4±1 years | Supine body positioned SE-NW. Grave damaged by ploughing, no grave goods^59^**.**  **14C**: 3105±15BP, cal. BP 3375-3251 (see Suppl. Tab. S9). **Chronology**: MBA3 |
| 16 | CGG_2_022573  Rise 1710 | F  55-65 years | Supine body positioned E-W; straight head, left hand on the hip and legs convergent at the feet. Two bronze plait clips found around the skull^59^.  **Chronology**: MBA3? |
| **OLMO DI NOGARA - AREA B** | | | |
| **Grave nr** | **Project ID** | **SEX/AGE** | **Context** |
| 412 | CGG_2_022635  Rise 1773 | M  30-35 years | Supine body positioned E-W; convergent legs at the feet. Damaged grave. Only the lower half of a bronze sword is preserved^59^. The sword piece gives no dating.  **14C**: 3140±20BP, cal. BP 3444-3265 (see Suppl. Tab. S9). **Chronology**: MBA2B-3A |
| 388 | CGG_2_022630  Rise 1678 | -  7±1 years | Supine body positioned E-W; head to the right/north. Decorated horn comb, two fragmentary pins, pin of the *Monte Lonato* type B, discoid amber bead, bronze plait clips (*fermatrecce*), bronze spring (*saltaleone*), bronze pendant. The grave goods suggest a date to the MBA 2^59^.  **Chronology**: MBA2 |
| 392 | CGG_2_022659 | M  40-50 years | Supine body positioned NE-SW; head straight. 16 bronze buttons in a circle of dark soil above the head. A bronze sword on the left part of the chest seems typical of the tumulus culture dating to the MBA 2^59^  **Chronology**: MBA2 |
| 471 | CGG_2_022643  Rise 1781 | F  20-25 years | Supine body positioned NE-SW; head to the left/south. Two bronze pins with embossed neck that date the context to the MBA 2^59^**.**  **Chronology**: MBA2 |
| 401 | CGG_2_022632  Rise 1770 | F  40-50 years | Supine body positioned E-W; head to the right/north.  The tomb is partly damaged. There were apparently no grave goods^59^**.**  **14C**: 3190±15BP, cal. BP 3449-3375 (see Suppl. Tab. S9). **Chronology**: MBA2B-3A |
| 408 | CGG_2_022634  Rise 1772 | -  4±1 years | Supine body positioned NE-SW; head to the left/south and legs bent to the left/south. There were apparently no grave goods, but three medium-large stones were piled up by the right elbow^59^.  **14C**: 3120±15BP, cal. BP 3385-3259 (see Suppl. Tab. S9). **Chronology**: MBA2B-3B |
| 430 | CGG_2_022638  Rise 1776 | -  5±1 years | Supine body positioned SE-NW; head straight. No grave goods^59^.  **14C**: 3180±15BP, cal. BP 3447-3370 (see Suppl. Tab. S9). **Chronology**: MBA2B-3A |
| 493 | CGG_2_022646 Rise 1784 (=Rise 461) | F  25-35 years | Supine body positioned E-W; head straight. The grave goods included two bronze pins of the *Montata* type, one bronze pin with a rolled head, two amber beads of truncated conical shape, one bronze plait clip (*fermatrecce*); the pins suggest a date MBA 3^59^  **14C**: 3172±28BP, cal. BP 3452-3354 (see Suppl. Tab. S9). **Chronology**: MBA2B-3A |
| 499 | CGG_2_022647 Rise 1785 | -  8±1 years | Supine body positioned NE-SW; head to the right/west. Damaged grave. No grave goods^59^ .  **Chronology**: ? |
| 404 | CGG_2_022633  Rise 1771 | M  > 50 years | Supine body positioned E-W; head to the left/south. No grave goods^59^.  **Chronology**: ? |
| 423 | CGG_2_022637  Rise 1775 | F  25-35 years | Supine body positioned NE-SW; head to the left/south. Damaged grave. No grave goods^59^.  **Chronology**: ? |
| 433 | CGG_2_022639  Rise 1777 | -  11±3 years | Supine body positioned NE-SW; head to the left/south. No grave goods^59^ .  **Chronology**: ? |
| 439 | CGG_2_022640  Rise 1778 | F  > 50 years | Supine body positioned SE-NW; head to the left/south. No grave goods^59^**.**  **Chronology**: ? |
| 459 | CGG_2_022641  Rise 1779 | F  25-35 years | Supine body positioned NW-SE; head straight. No grave goods^59^ (Salzani 2005, 85-86). Probably pregnant, the bone of the newborn lies on the pelvis.  **Chronology**: ? |
| 462 | CGG_2_022642Rise 1780 | -  1 year | Supine body positioned E-W. Grave heavily damaged. No grave goods^59^**.**  **Chronology**: ? |
| 478 | CGG_2_022645  Rise 1783 | M  30-40 years | Supine NE-SW. Grave heavily damaged only the feet were in situ. No grave goods^59^**.**  **Chronology**: ? |
| **OLMO DI NOGARA - AREA C** | | | |
| **Grave nr** | **Project ID** | **SEX/AGE** | **Context** |
| 28 | CGG_2_022574  Rise 1712 | M  24-30 years | Supine body positioned E-W; head to the right/north and towards the hilt of the sword. Sword blade with rivets of the *Bigarello* type parallel to the right arm. The *Bigarello* swords are very few and not of secure dating, possibly MBA or MBA 2^59^.  **Chronology**: MBA (MBA?) |
| 32 | CGG_2_022576  Rise 1714 | F  40-50 years | Supine body positioned SE-NW; head to the left/south. Grave damaged by ploughing. Two bronze pins with a double hole, one small bronze pin with two rings, two cylindrical amber beads, one fragmentary bronze pin/needle, one bronze stud. The pins may be dated to the MBA 2B-3^59^**.**  **Chronology**: MBA2B-3 |
| 45 | CGG_2_022579  Rise 1717 | -  1,5 year | Probably supine body positioned NE-SW. Grave damaged by grave 46 and by ploughing, no grave goods^59^**.**  **14C**: 3160±15BP, cal. BP 3445-3350 (see Suppl. Tab. S9). **Chronology**: MBA2B-3A |
| 57 | CGG_2_022581  Rise 1719 | F  30-40 years | Supine body positioned E-W; head to the left/south and towards grave 58. The legs lie over the stratigraphically lower grave 58. Two bronze pins, one with 3 rings and one with a spiral of the *Santa Caterina* type A; the latter suggest a date to MBA 3^59^  **Chronology**: MBA3 |
| 58 | CGG_2_022582  Rise 1720 | -  5±2 years | Supine body positioned SE-NW; head to the right/north and towards grave 57. There were apparently no grave goods^59^. The legs of the individual in grave 57 were above the legs of this body, which is therefore contemporary or earlier, possibly MBA3.  **Chronology**: MBA3? |
| 137 | CGG_2_022594  Rise 1732 | F  35-45 years | Supine body positioned NE-SW; head to the right/north; hand on the hip and convergent legs at the knees. Two bronze pins, one fragmentary and one with a spiral head, and a bronze rod. Differently from the 14C, the pins would suggest a date to the RBA^59^**.**  **14C**: 3135±20BP, cal. BP 3441-3261 (see Suppl. Tab. S9). **Chronology**: MBA2B-3B |
| 197 | CGG_2_022600 Rise 1738 | -  5± ? months | Supine body positioned E-W, resting above the legs of the body in grave 199. Grave damaged by ploughing, no grave goods^59^**.**  **14C/2σ cal.**: 3145±20BP, cal. BP 3444-3269 (see Suppl. Tab. S9). **Chronology**: MBA2B-3B |
| 207 | CGG_2_022603  Rise 1741 | M  > 50 y. | Supine body positioned E-W; head to the right/north; convergent legs at the knees. A fragmentary bronze stud (*borchia*), a bronze arrowhead in between the legs^59^  **14C**: 3190±15BP, cal. BP 3449-3375 (see Suppl. Tab. S9). **Chronology**: MBA2A-2B |
| 224 | CGG_2_022606  Rise 1744 | F  45-55 years | Right lateral recumbent body with bent legs, S-N. Right arm along body, left arm bent with whistle under the chin. Unusual position as in grave 156. There were apparently no grave goods^59^.  **14C**: 3220±15BP, cal. BP 3459-3391 (see Suppl. Tab. S9). **Chronology**: MBA2A-3A |
| 235 | CGG_2_022611  Rise 1749 | -  c. 3 years | Supine body positioned E-W; head straight; Graves heavily damaged. Partly superimposed to grave 236. There were apparently no grave goods^59^.  **14C**: 3160±15BP, cal. BP 3445-3350 (see Suppl. Tab. S9). **Chronology**: MBA2B-3A |
| 234 | CGG_2_022610  Rise 1748 | M  15±1 years | Supine body positioned NE-SW; head to the left/north and convergent legs at the knees. Left hand on the hip. There were apparently no grave goods^59^  **14C**: 3110±15BP, cal. BP 3378-3254 (see Suppl. Tab. S9). **Chronology**: MBA3 |
| 282 | CGG_2_022614  Rise 1752 | -  Fetus/ Newborn | Supine body positioned NE-SW; head to the right/north; There were apparently no grave goods^59^.  **14C**: 3105±15BP, cal. BP 3375-3251 (see Suppl. Tab. S9). **Chronology**: MBA3A-3B |
| 318 | CGG_2_022623  Rise 1761 | -  11±1 years | Supine body positioned NE-SW; head to the right/north and convergent legs at the knees. There were apparently no grave goods^59^**.**  **14C**: 3100±15BP, cal. BP 3372-3250 (see Suppl. Tab. S9). **Chronology**: MBA3A-3B |
| 230 | CGG_2_022609  Rise 1747 | -  6±3 months | Supine body positioned E-W. Grave damaged by ploughing, no grave goods^59^.  **14C**: 3100±20BP, cal. BP 3375-3244 (see Suppl. Tab. S9). **Chronology**: MBA3A-3B |
| 321 | CGG_2_022624  Rise 1762 | F  35-45 years | Supine body positioned NE-SW; head to the left/south and right hand on the hip; There were apparently no grave goods^59^  **14C**: 3090±15BP, cal. BP 3367-3243 (see Suppl. Tab. S9). **Chronology**: MBA3A-3B |
| 30 | CGG_2_022575  Rise 1713 | F  20-30 years | Supine body positioned SE-NW. Grave damaged by ploughing. Two bronze pins of the *Franzine* type, one bronze ring, one amber bead. The bronze pin suggests a chronology to the RBA^59^**.**  **14C**: 3060±20BP, cal. BP 3355-3210 (see Suppl. Tab. S9). **Chronology**: MBA3B-RBA1 |
| 301 | CGG_2_022620  Rise 1758 | F  40-50 years | Supine body positioned NE-SW; head to the left/south and convergent legs at the feet. There were apparently no grave goods^59^**.**  **14C**: 3090±20BP, cal. BP 3368-3239 (see Suppl. Tab. S9). **Chronology**: MBA3A-RBA1 |
| 287 | CGG_2_022615  Rise 1753 | F  35-45 years | Supine body positioned NE-SW; The head rests on the body of the child in grave 288, which is stratigraphically lower; the feet overlap. The grave goods icnluded one bronze pin of the *Cataragna* type. One bronze pin with a rolled head. The pins suggest a date to RBA^59^**.**  **14C**: 3080±15BP, cal. BP 3360-3236 (see Suppl. Tab. S9). **Chronology**: MBA3A-RBA1 |
| 226 | CGG_2_022607  Rise 1745 | F  30-40 years | Supine body positioned NW-SE; head to the right/east and convergent legs at the feet. Fragments of a bronze pin/needle^59^.  **14C**: 3070±15BP, cal. BP 3355-3225 (see Suppl. Tab. S9). **Chronology**: MBA3B-RBA1. |
| 118 | CGG_2_022589  Rise 1727 | -  C. 1 year | Supine body positioned E-W; head to the right/north. In the grave there was a bronze pendant, whose shape suggests a chronology to the MBA 3^59^**.**  **14C**: 3065±20BP, cal. BP 3355-3215 (see Suppl. Tab. S9). **Chronology**: MBA3B-RBA1 |
| 136 | CGG_2_022593  Rise 1731 | -  Newborn | Grave completely destroyed, no grave goods^59^**.**  **14C**: 3070±25BP, cal. BP 3362-3213 (see Suppl. Tab. S9)  **Chronology**: MBA3B-RBA1 |
| 297 | CGG_2_022619  Rise 1757 | F  50-60 years | Supine body positioned E-W; head to the left/south and both hands on the hips; the lower part of the legs is over grave 304. One bronze needle. Two bronze pins with a spiral head of the *Santa Catarina* type suggest a MBA 3-RBA date^59^**.**  **Chronology**: MBA3-RBA |
| 228 | CGG_2_022608  Rise 1747 | -  5±1 years | Supine body positioned NE-SO; head to the right/north; probably convergent legs. Grave damaged by ploughing. Two perforated bone discs. Bronze flange-hilted dagger of *Bertarina* type, which according to is to be dated to the RBA^59,135^.  **Chronology**: RBA? |
| 110 | CGG_2_022585  Rise 1723 | F  20-25 years | Supine body positioned NE-SW; head straight and left hand on the hip, while on the right arm lies a newborn; the legs are convergent at the knees. One bronze *Cataragna* pin and one *Colombare* type A pin, and one bronze stud. Both pins suggest a date to RBA^59^  **Chronology**: RBA? |
| 105 | CGG_2_022584  Rise 1722 | F  15±2 years | Prone body positioned E-W; head to the right/north; the hands are under the hips and the legs convergent at the feet. Bronze plait clip by the head, one pin of the *Cataragn*a type, and one small pin with a similar perforation as the previous one. The *Cataragna* pin suggests a date to RBA^59^.  **Chronology**: RBA |
| 296 | CGG_2_022618  Rise 1756 | -  7±2 years | Supine body positioned NE-SW; head to the right/east and legs slightly bent to the left. A decorated bronze comb. Two bronze (*saltaleoni*) spring wires. The grave is in the south part of area C, where there seem to be only RBA graves^59^  **Chronology**: RBA? |
| 180 | CGG_2_022597  Rise 1735 | F  35-45 years | Supine body positioned E-W. Grave damaged by ploughing, no grave goods^59^.  **14C**: 2945±25BP, cal. BP 3205-3000 (see Suppl. Tab. S9). **Chronology**: RBA1-FBA |
| 43 | CGG_2_022577  Rise 1775 | M  35-45 years | Supine body positioned E-W; head to the right/north-west and convergent legs at the feet. No grave goods^59^  **Chronology**: ? |
| 44 | CGG_2_022578  Rise 1716 | -  12±3 years | Grave heavily damaged by recent digging^59^**.**  **Chronology**: ? |
| 53 | CGG_2_022580  Rise 1718 | M  25-30 years | Supine body (almost sitting) positioned E-W; head to the left/north; No grave goods^59^**.**  **Chronology**: ? |
| 104 | CGG_2_022583  Rise 1721 | F  35-45 years | Supine body positioned SE-NW; head to the left/south, the right hand is on the hip. There were apparently no grave goods^59^**.**  **14C**: 3055±20BP, cal. BP 3350-3184 (see Suppl. Tab. S9). **Chronology**: MBA3B-RBA1 |
| 111 | CGG_2_022586  Rise 1724 | -  5±1 years | Supine body positioned NE-SW; head to the left/north. Grave damaged by ploughing, no grave goods^59^  **Chronology**: ? |
| 127 | CGG_2_022591  Rise 1729 | M  45-50 years | Supine body positioned NE-SW; head to the right/north; the left hand is under the hip and the legs are convergent at the knees. No grave goods^59^.  **Chronology**: ? |
| 133 | CGG_2_022592  Rise 1730 | -  c. 5 years | Grave completely destroyed, no grave goods^59^  **Chronology**: ? |
| 159 | CGG_2_022595  Rise 1733 | F  40-50 years | Supine body positioned E-W; head to the left/south; right hand on the hip. No grave goods^59^  **Chronology**: ? |
| 183 | CGG_2_022598  Rise 1736 | -  6 months- 1 years | Supine body positioned E-W. Grave damaged by ploughing^59^  **Chronology**: ? |
| 184 | CGG_2_022599  Rise 1737 | -  Newborn | Grave completely damaged by ploughing^59^  **Chronology**: ? |
| 199 | CGG_2_022601  Rise 1739 | F  30-40 years | Supine body positioned NE-SW; head to the right/north; the legs are below the body in grave 197. The grave goods included two bronze pins with a rolled head, a fragment of a needle close to the head. The pins with rolled head suggest a date to RBA, however Grave 197 above this is 14C dated to MBA 2B-3B^59^.  **Chronology**: MBA? |
| 200 | CGG_2_022602  Rise 1740 | F  50-60 years | Supine body positioned E-W; head straight and right hand on the hip. No grave goods^59^**.**  **Chronology**: ? |
| 213 | CGG_2_022605  Rise 1743 | M  50-60 years | Supine body positioned E-W; head to the left/south; right hand over and left hand under the pelvis. No grave goods^59^**.**  **Chronology**: ? |
| 274 | CGG_2_022613  Rise 1751 | -  Fetus/ Neborn | Grave damaged by ploughing. No grave goods^59^ **Chronology**: ? |
| 289 | CGG_2_022616  Rise 1754 | M  40-50 years | Supine body positioned W-E; head to the left/north; convergent legs at the knees. There were apparently no grave goods^59^  **14C**: 1275±20BP, cal. BP 1280-1175 (see Suppl. Tab. S9). **Chronology**: Late Antiquity/Early Middle Age |
| 293 | CGG_2_022617  Rise 1755 | -  6-18 months | Supine body positioned NE-SW; head to the left/south. No grave goods; however, there were sparse body parts of two female individuals (incomplete) one of them had 2 fibula. The fibula is not published^59^  **Chronology**: ? |
| 313 | CGG_2_022621  Rise 1759 | F  20-30 years | Supine body positioned NE-SW; head to the right/north. A fragmentary bronze pin/needle^59^  **Chronology**: ? |
| 314 | CGG_2_022622  Rise 1760 | -  Newborn | Supine body positioned NE-SW. The grave is heavily damaged^59^  **Chronology**: ? |
| 324 | CGG_2_022625  Rise 1763 | F  > 50 years | Supine body positioned NE-SW; head to the left/south; convergent feet. Two bronze pins of the *Cataragna* type suggest a date to the end of MBA 3 and RBA^59^.  **Chronology**: MBA3-RBA |
| 362 | CGG_2_022626  Rise 1764 | -  2-3 years | Supine body positioned E-W; head to the right/north. No grave goods^59^ **Chronology**: ? |
| 372 | CGG_2_022628  Rise 1766 | -  2±2 years | Supine body positioned NE-SW; convergent feet. No grave goods^59^.  **Chronology**: ? |
| 378 | CGG_2_022629  Rise 1767 | -  Newborn | Supine body positioned NE-SW. No grave goods^59^.  **Chronology**: ? |
| **OLMO DI NOGARA - AREA C NORTH-WEST** | | | |
| 543 | CGG_2_022649  Rise 1787 | -  2±1 years | Supine body positioned E-W. Heavily damaged grave, no grave goods^133^.  **14C**: 3145±15BP, cal. BP 3441-3274 (see Suppl. Tab. S9). **Chronology**: MBA2B-3B |
| 541 | CGG_2_022648  Rise 1786 | -  5±2 years | Supine body positioned E-W; head to the left/south; the phalanges of both hands and feet are missing. There were apparently no grave goods^133^.  **14C**: 3115±15BP, cal. BP 3381-3255 (see Suppl. Tab. S9). **Chronology**: MBA3 |
| 544 | CGG_2_022650  Rise 1788 | -  1,5 years | Prone body positioned NE-SW. The prone position is known only in Olmo grave 56, 105, 225 and 554. There were no grave goods^133^**.**  **14C**: 3110±15BP, cal. BP 3378-3254 (see Suppl. Tab. S9). **Chronology**: MBA3 |
| 547 | CGG_2_022651  Rise 1789 | M  25-35 years | Supine body positioned NE-SW; head straight; convergent kegs at the knees. Bronze sword with *codolo a spina* of the *Chiese* type, which suggests a date to the MBA 3^133^**.**  **Chronology**: MBA3 |

Table 8 Synoptic table with data about the sampled individuals from Olmo di Nogara and their context of origin, presented separately in Area A, B and C.

##### ***2.7.7. Pian Sultano, Rome province, Latium***

Latitude: 42.0443385569645; Longitude: 11.976588998481512

Sample provider: Flavia Trucco

Serena Sabatini, Flavia Trucco

Pian Sultano is a unique setting. It is not a grave rather an area where ritual activities took place including the deposition of selected human remains. The remains were not cremated. The site is very difficult to access being at the bottom of a narrow (c. 45 cm wide) natural crevice in the bedrock. The crevice is c. 10 m deep and it was not only very difficult to excavate the site in modern times, but it must have been difficult to access it in ancient times as well. The excavations brought to light human remains, a conspicuous number of animal remains (c. 400 fragments) and ceramics (probably originally c. 40 vases). All the material was in fragmentary conditions and spread all over. The ceramic material suggests that the site was in use at the end of the Early Bronze Age. Some of the vases are difficult to date and could be from the beginning of the Middle Bronze Age.

The osteological investigations of the sampled material^136^ suggest they belong to three different individuals out of a total of eight that were found in the crevice. Individual A (**CGG_2_101262**) was an adult (c. 30-40 years) probably female with evident signs of anaemia. Individual B (**CGG_2_101264** and **CGG_2_101265**) was a young adult (c. 18-25 years) probably female. Individual C (**CGG_2_101266**) was juvenile (c. 12-16 years).

##### ***2.7.8. Scalvinetto, Padova province, Veneto***

Latitude: 45.12271146984293; Longitude: 11.281629277018682

Sample provider: Claudio Cavazzuti

Serena Sabatini

Scalvinetto is a large bi-ritual cemetery. It is similar in many ways to Olmo, which is only about 22 km away. The site is just 400 m north-west of the large settlement of Fondo Paviani (cf. Cupitò et al. 2015) and it is most likely one of its cemeteries. Fondo Paviani is a site in use from the Middle Bronze Age 3 (c. 1450 BCE) whose size grows exponentially from c. 7 to 15-20 ha during the Recent Bronze Age. Differently from the Terramare south of the Po River it does not collapse at the end of the period, but rather continues to be inhabited during the subsequent Final Bronze Age (c. 1115-950/900 BCE). Fondo Paviani is known among other things also for its late Mycenaean imports. Most of the known fragments of Mycenaean pottery from northern Italy come from this site. It was discovered in 1989 and excavated in 1991-1994. 243 tombs (192 cremations and 51 inhumations) were excavated. In 2000-2003, new excavations took place^60^; 462 graves (245 cremations and 217 inhumations) were then investigated. The inhumations represent 38% of the total. Although these data are preliminary, it might be that inhumations or at least some of them precede cremations. The crematory ritual might therefore have been introduced at a slightly later stage^137^. The chronology of the cemetery goes from the Middle Bronze Age to the Recent Bronze Age 2. The topography of the site is interesting. The inhumations are mostly concentrated in the western part of the cemetery, while cremations are in the eastern part of it.

Strontium isotope analyses of the individual (adult male c. 30-40 years) in grave 405 (**CGG_2_022900/Rise 1887**) revealed that the ^87^Sr/^86^Sr values of his M2 tooth were considerably higher (0.719292) then those in his M3 tooth (0.711976), thus it has been concluded^61^ that he must have moved in young age from an Alpine area to an area closer to the Fondo Paviani/Scalvinetto. Our second sample (grave 285, **CGG_2_022898/Rise 1885**) belongs to an adult female of c. 30-40 years. Only the M2 sup tooth was analysed for this individual and gave a strontium value compatible with that of the Po plain (^87^Sr/^86^Sr 0.709620), at least as far as we understand it now. The third sample comes from grave 401 (**CGG_2_022899/Rise 1886**). All three individuals were deposited supine, in graves with an east-west orientation, apparently without grave goods.

##### ***2.7.9. Sedegliano, Udine province, Friuli- Venezia Giulia***

Latitude: 46.007928733115655; Longitude: 12.969459519598159

Sample provider: Alessandro Canci, Elisabetta Borgna, Susi Corazza

Serena Sabatini, Susi Corazza, Elisabetta Borgna

Four simple graves have been recovered from the embankment that surrounded the local *castelliere* settlement (Castellieri is the name given to the Bronze Age fortified settlements from northeastern Italy); the tombs were dug out in the early nucleus of the rampart. Tomb 1 contained a male individual of c. 45-55 years (**CGG_2_022653/Rise 1791**). The position of the bones suggested that the body was interred in a wooden coffin or wrapped in a thick organic material which decomposed after the body influencing the movement of the bones once the soft tissues had disappeared^138^.

The sample has been radiocarbon dated to 3420±40^139^, cal. BP 3827-3565 (see Suppl. Tab. S9), thus confirming the chronology of the foundation of the rampart in the Early Bronze Age, as indicated by the other tombs, ranging from earlier in the EBA (ca 1880) well into the early MBA.

##### ***2.7.10. Selvis, Udine province, Friuli-Venezia Giulia***

Latitude: 46.07; Longitude: 13.32 (approximately)

Sample provider: Alessandro Canci, Elisabetta Borgna, Susi Corazza

Serena Sabatini, Susi Corazza, Elisabetta Borgna

The individual (**CGG_2_022923/Rise 1888**) analysed in this project comes from a burial mound destroyed in the 1980s by agricultural works. Skeletal remains (from a grave of the Roman period) noticed during the removal of the tumulus suggested that it could have been an ancient grave. Rescue excavations were therefore carried out and could investigate the central grave of the tumulus belonging to a male individual of c. 20-25 years^138^. The body was deposited on a bed of selected stones. Judging from the position of the bones it was probably originally wrapped in organic material. It had a stone pendant by the ankle and a bronze dagger on the chest which after decomposition slid towards the left elbow. The deposition was dated to an advanced phase of the Italian Early Bronze Age (c. 2150-1650 BCE). The sample has been radiocarbon dated in 2014 at Centro di Datazione e Diagnostica (CEDAD) dell’Università del Salento to 3515±50, cal. BP 3960-3642(see Suppl. Tab. S9). A second 14C analysis was carried out in this study (UCIAMS-226880) and gave a similar result 3530±2BP, cal. BP 3885-3719 (see Suppl. Tab. S9). Both are in agreement with the chronology provided by the archaeological record confirming it is an Early Bronze Age burial.

##### ***2.7.11. S. Osvaldo, Udine province, Friuli- Venezia Giulia***

Latitude: 46.030851519247655; Longitude: 13.22755382838825

Sample provider: Alessandro Canci, Elisabetta Borgna, Susi Corazza

Serena Sabatini, Susi Corazza, Elisabetta Borgna

The individual (**CGG_2_022652/Rise 1790**) analysed in this project comes from a burial mound with a circular base and a stone core or cairn, typical of the area. The mound covered a single burial with an adult individual of c. 25-35 years, likely male (the genetic sex of this individual also resulted to be male). The individual was in a slightly crouched position on his left side. The study of the remains suggested that the body has been buried in a wooden coffin embedded within the stones of the cairn and forming a small built chamber. With the degradation of the coffins the stones had fallen on the remains causing the modifications visible at the time of the excavation. The bones have been 14C dated to 3580±50BP^140^, cal. BP 4076-3717 (see Suppl. Tab. S9).^140^

##### ***2.7.12. Toppo Daguzzo, Potenza province, Basilicata***

Latitude: 41.000730801709935; Longitude: 15.729134513424887

Sample provider: Alessandro Canci, Mirella Cipolloni Sampò

Serena Sabatini

The site at Toppo Daguzzo includes a large inland settlement (probably dominating trade routes between the Tyrrhenian, the Adriatic and the Ionian Sea in southern Italy) and its cemeteries. The site was apparently in use for a long time from the Copper Age to the Iron Age. Several graves were excavated at the site. They are monumental hypogea dug out of the rocky side of the local acropolis. The remains of looted or partially destroyed contexts suggest that there were originally larger numbers of such graves. The majority of them must have been destroyed by the modern stone quarry established along one side of the hill. Of those who survived it is interesting to notice that not all hypogea had a primary funerary purpose; some were clearly used for cultic activities.

Tomb 3 is one of the most impressive among the excavated hypogea. It occupies a prominent position on the south-eastern slope of the hill, but very close to the top. It was accessed through a long imposing dromos, also dug out of the bedrock; it had a rectangular chamber of noteworthy dimensions, probably divided in two similarly sized rooms by a sort of internal wooden wall. Three phases could be investigated in the tomb. The archaeological record from the most recent layer must have been looted already in antiquity. Looters took with them all possible grave goods making it almost impossible to date this last phase. Below this looted layer, and after a sterile stratum, a second burial phase could be detected. This middle phase is archaeologically and 14C dated to the Early Middle Bronze Age (1650-1550 BCE) and contained 11 individuals (five adult males, four females and two young individuals of c. 5-6 and 12-13 years respectively, cf.^141^. Individuals 4 and 11 from this stratum have been sampled for this project. Individual 4 (**CGG_2_022655/Rise 1793**) should be an adult male^141^ of c. 30-40 years accompanied by a spearhead^142^ (fig. 8: 4) Individual 11 (**CGG_2_022656/Rise 1794**) is supposed to be an adult male of c. 25-35 years buried with a sword of Pertosa type^142^ (fig. 9:3). The archaeological evidence from the lowest and earliest stratum of the hypogeum suggests that during this early phase, the context was used for ritual purposes. The anthropological characteristics of most of the individuals from both the looted most recent phase and the second middle burial phase suggest genetic affinity; in facts, all individuals appear characterised by robust traits and significant height^125,141^; another feature that suggest genetic affinity is the relatively low age of death and the spread presence of tumour(?) among the deceased^141^. The Early Middle Bronze Age at Toppo Daguzzo tomb 3 shows a strict funerary ritual with rigid gender connotations embodied by a sword or daggers for the male individuals and bone spindle whorl and *exotica,* amber and glass, for the female individuals; no other grave goods were recovered in the grave with the exception of three small vases deposited in close vicinity to the only child in the middle of the inner room of the grave.

#### **2.8. Lebanon**

##### ***2.8.1. Sidon, Sidon district, South Governorate***

Latitude: 33.5615957598156; Longitude: 35.3712556821047

Sample provider: Claude Serhal

Claude Serhal

On the coast of Lebanon lies the city of Sidon mentioned 38 times in the Old Testament and considered by some to be the more ancient and the most prominent of the Canaanite/Phoenician coastal cities. Highlighted in biblical accounts as being the “first born of Canaan”, Sidon’s inhabitants were the most well-known of Phoenician communities, to such an extent in fact that the designations Phoenician and Sidonian were often considered synonymous. Thus far the main sources of information on Sidon’s history have been based on finds not from the city itself but from the surrounding area. This situation changed in 1998 when the excavation project which is still ongoing started on Sidon’s ancient mound or tell, right in the heart of the city. The site which is called ‘College site’ (Fig. 8; Fig. 9) after a couple of 19^th^ century schools which were built there and then later demolished around 1967, is part of three downtown pieces of land expropriated by the Lebanese Department of Antiquities for the specific purpose of archaeological research. It is therefore not subject to any limit or any pressure from commercial developers.

The Sidon excavation revealed an exceptional continuity in occupation. Once settled in their bedrock-based round houses at the end of the Chalcolithic Period, the Sidonians never left this location and the site was to remain occupied almost continuously throughout the third and 2^nd^ millennium BCE, well into the 1^st^ millennium BCE, and encompassed the Persian and Hellenistic Periods which were succeeded by the Roman and Crusader events, and whereby all these Sidonians at one time or another, saw fit to take up residence.

The Middle Bronze Age in Canaan marks the beginning of a new urbanised society and an era of international trade and cultural exchange in the Eastern Mediterranean. This time period is represented in Sidon by a sand level ranging from 90 cm to 1.40 m. This sand was extremely fine and has been brought to the site from the nearby sea shore. Clearly at the end of the third and the beginning of the second Millennium BC the site undertakes a change of function. Sidon’s 3^rd^ millennium occupation level is covered with sand and the site becomes a burial site. 172 burials were found to date in the sand (phase 1-3) and in the occupation levels on top of it which was resumed sometime around 1750 BCE (level 4) allowing to follow the development of mortuary practices until the end of the Middle Bronze Age (levels 5 to 8).

A variety of grave types is evidenced over time starting with the constructed graves of warriors. Burial type changes and the most common burial types after 1750/1700 BCE is the jar burial of neonates and infants, very rarely the burial in the ground of an adult in a flexed position and constructed multiple burials. On the basis of the mortuary evidence, it is possible to draw certain inferences regarding the social structure of the Sidonian community.

Burial 91 is the only burial which corresponds to a re-deposition of bones in the Iron Age.

As the excavation is still ongoing most of the burials have been so far published in preliminary reports^143–150^.

**Phase 1**

*Burial 78.* Consisted of a rectangular mud brick-lined grave. The excavation of the burial revealed a thick layer of grey sticky mud covering the skeleton. The individual (**CGG_2_104308**) was an adolescent possibly between 13 and 18 years old. The body was in a flexed position, aligned on an East-West axis, with the head to the east, the face to the north and the hips/feet to the west, with the individual resting on its right side. The legs were tightly flexed. A miniature spearhead was exposed in front of the individual’s face in the North-East corner of the grave, just above the right hand and a bronze miniature duckbill axe was recovered just behind the skull in the South-East corner of the grave. The individual was wearing a bracelet with carnelian beads and scarabs around his left hand. Animal remains were found under the legs.

**Phase 3**

*Burial 99.* Burial 99 was a constructed grave of both stone and mud brick. The alignment of the stones was roughly NE/SW-E/W. The body appeared to be aligned along the same axis in a flexed position with the head to the east facing S/SE. The young adult female individual, aged around 30 to 35 years (**CGG_2_104307**) was placed lying on her left side with the knees towards the S/SW. A large scarab was recovered from next to the mid-shaft of the lower left arm and a second scarab was recovered from amid the finger bones of the left hand. The deliberate lining of the grave floor with a thin sandy layer was noted.

**Level 4**

*Burial 70.* This well-built grave, constructed using a rectilinear stone lining, contained at least a single young adult male individual *in situ* aged approximately 25 to 35 years with disarticulated adult and subadult remains and animal bones also present. The grave was very deep with its stone lining still intact to 1m high. The grave’s long axis was oriented roughly along an EW alignment. The articulated remains were placed along a similar EW line, with the feet to the west and the upper body to the east. The upper torso and head were missing, most likely truncated, with no definite articulation observed in the poorly preserved skeletal remains in this area at the time. Both legs appear to have been articulated in situ. The right leg was articulated with the right hip *in situ* with the knee flexed and in an upright position. The right femur proceeded roughly W/NW, with the right lower leg returning towards SW. The left leg was also articulated with the left hip in situ, again with the knee flexed and somewhat upright. The left femur proceeded roughly W/SW, with the right lower leg returning towards NW. The distal ends of both extant lower legs were deliberately placed on top of a large angular stone at the western end of the stone-lined grave.

The disarticulated human remains represented at least an additional two individuals. A knife was found on the pelvis under a skull in a jumbled area of bones indicating the presence of both subadult and adult remains. A bronze spearhead was found near the surface of the fill and is probably a disturbed grave good from the original burial. The twine that attached the shaft to the blade was extant and was used to secure the end of the socket to the joint between the edges of the socket and the shaft. The upper fill (1083/1109) consisted of a loose dark orange-brown silty sand which was not distinguishable in colour or texture from the lower fill (1136). After backfilling, the eastern area was spread with a rectangular layer of white plaster which covered the remains of animal bones embedded underneath. These consisted of seven individuals namely, three caprines and one adult *Bos taurus.* The sample in this study is the individual 70B (**CGG_2_104314**).

*Burial 93.* Burial 93 was a steep-sided oval cut gradually descending to a break of slope at the base containing the burial jar. The top of the jar was truncated by burial 84a/b’s cut. The jar was oriented along a North-South axis and contained the remains of what might have been a subadult (early to late childhood) of less than 8 years and which were also oriented along the same NS axis as the jar itself. The skull (**CGG_2_104312**) was to the south and facing north (i.e. towards the neck of the burial jar). The individual appears to have been placed lying on the right side with the upper body almost supine. The legs were very tightly flexed. Inside the jar were several medium-sized stones, deliberately and carefully placed over and around the body, possibly to protect it. There were also 4 adult hand phalanges found together in the upper fill of the jar burial against the eastern internal side.

*Burial 103.* This multiple burial (103-104-105-107-109) was constructed along an East-West axis. Burial 103 was an old adult male (**CGG_2_104258**) inhumation 60 to 80 years old aligned along the same East-West axis as the stone-lined grave with the head to the east facing north, the legs in a flexed position which in turn appeared to be propped directly against the articulated individual of burial 104 suggesting all articulated individuals were likely to have been placed inside this grave at the same time.

**Stratum 5**

*Burial 6.* The grave was a 1.35 m x 70 cm deep pit dug in yellow sand with a large amount of mixed charcoal and limestone. The skeleton of a young, probable male, adult (**CGG_2_104281**) was lying on its left side in a flexed position to the south with the head to the east. The arms were crossed on the chest. Beach pebbles, most probably brought to the site, were associated with the burial. Disarticulated adult human bones appeared after lifting the skeleton.

*Burial 44.* This jar burial consisted of a subadult (perinate) in a North-South position with the body in the same axis. The bones of the child (**CGG_2_104257**) had been crushed by the jar fragments. A carinated bowl was resting on top of the base of the jar, another bowl covered the bones. A jug was also found in the jar.

*Burial 47.* This jar burial was deposited in a relatively small but deep rectangular cut which truncated the western side of another cut containing burials 45 and 46. Found intact and undisturbed, this jar was oriented along an EW axis with the neck towards the east and the base towards the west. The skeletal remains (**CGG_2_104256**) of the infant (birth to 1 year) covered by a clayey backfill, were lying on the bottom of the jar with the cranium slightly crushed and the frontal/parietals rather flattened out. The body was aligned in an East-West position with the head to the east. The face was facing W/NW. The legs were partially flexed with the right knee towards the north. The left leg was less flexed with the femur along an East-West axis and the left tibia flexed towards the N/NE. The spine was more or less along an East-West axis and suggests the individual was placed supine inside the jar.

*Burial 69.* Continued excavation under burials 63 and 67 in a stone-lined grave cut revealed two further articulated inhumations, 69 and 74, as well as a collection of disarticulated remains (burial 75) in the SW corner of the grave. Burial 69 was that of a subadult female individual (late childhood) between 6 and 12 years in a supine position buried head to the north facing west, with the knees in a flexed position facing west. The skeleton (**CGG_2_104300**) was interred with the head to the north and the cranium facing west. The thorax and ossa coxae were in a supine position, although the orientation of both sides of the rib cage towards the west suggests that the body may have been interred with its left side slightly elevated. The right upper limb was extended with the right hand underlying the right thigh. The left hand was observed to be clenched and closed. Both left and right lower limbs were semi-flexed at the hips and flexed at the knees. The knees were oriented towards the west. The feet were in anatomical alignment being oriented inferiorly towards the south. A spearhead with a slender blade and a socket retaining evidence of a hole for a transverse peg was found near the legs of the deceased.

*Burial 77.* Burial 77 was a grave cut roughly ovoid in shape containing a jar burial and appeared to slightly cut burial 70 to the north. The jar was lying along an East-West axis with the base to the east. The human remains (**CGG_2_104299**) were those of a young subadult individual (early childhood), between 1 to 5 years with the body also aligned along an East-West axis with the head to the west, the face towards the east. The remains were in a flexed position with the head to the west, feet and hips to the east and legs tightly flexed with the knees to the North/North-West. The right forearm was roughly aligned along a South-West/North-East axis. The left arm was extended to the north of the skull. Several small rock crystal beads were recovered from the neck area.

*Burial 82.* This jar burial belonged to a female adolescent 16 to 17 years old (**CGG_2_104313**) . The jar was aligned along a north south axis with the opening sealed by mud brick and with two limestones (18 x 17 x 6 cm - 16 x 16 x 10 cm) placed on either side of it. The skeleton was oriented with the head to the north. The thorax was in supine position with the left and right arms lying along the rib cage. The left and right forearms were gently flexed. The skull had rolled forward and lay on its right side, facing west. The neck had also rolled forward.

*Burial 84.* Burials 84A and 84B consisted of an unlined grave pit containing two adult individuals, one on top of the other. Burial 84A (**CGG_2_104303**) was that of an adult elderly male 50 to 70 years old, oriented along a North-South axis with the head to the north facing upwards and the feet to the south with the knees towards the east. The body was in a flexed position. The upper body appeared more or less supine with the legs on their side, the knees toward the west and the feet proceeding back toward the South/South-East. Burial 84B (**CGG_2_104306**) contained a flexed female adult individual 60 to 80 years old, in a position almost identical to that of burial 84A, except that the individual had been rotated 180° with the head to the south, facing east and the feet and hips facing north. There was a small concentration of charcoal in front of the individual’s forehead and burnt clay below the mouth and on top of the neck area.

*Burial 86.* Burial 86 (**CGG_2_104315**) was the jar burial of an infant (birth to 1 year) interred with its head to the south. The skull at the base of the jar was facing north. The thorax lay on the right side with the ribs oriented towards the east of the vessel.

*Burial 87.* This jar burial contained a subadult (early childhood) child of 5 years (**CGG_2_104301**) . The burial lay beneath three large rocks and some burnt mud bricks which were placed at the mouth of the vessel in the north. The body lay on the right side and was oriented with the head to the south, facing east. The right forearm was slightly flexed at the elbow. The left arm lay superior to the left ribs and the left forearm was flexed at the elbow. The left and right hands were absent apart from one right distal phalanx I (thumb). The left and right thighs were flexed at the hips. The knees were oriented to the east of the jar. The left and right legs were flexed at the knees. The left and right feet were extended at the ankles, oriented to the west of the jar. The foot phalanges were absent. A juglet sat atop the base of the jar, to the east of the ossa coxae, south of the hands. A carinated bowl sat on top of the legs. The bowl was crushed *in situ* as it was directly beneath large rocks and a tannour was overlying the northern end of the burial. The legs and feet were also crushed due to this overburden. Osseous preservation was fair although almost all skeletal elements were fragmented.

*Burial 89*. This burial contained the remains of a perinate (**CGG_2_104316**) lying on the right side in a flexed position, with the knees to the north. The jar was aligned along an EW axis, with the base to the east. The west end was truncated. The majority of the remains *in situ* were articulated and the body was also aligned EW with the head to the east, facing north. The lower legs were truncated and only the left femur was present. The left thigh was flexed at the hip, with the knee oriented north. The burial was truncated at this point and thus the left leg, the left foot and the right lower limb were absent. Osseous preservation was good although many skeletal elements were fragmented *in situ*. The matrix was of fine dark grey soil which was dry and compacted. Occasional small flecks of charcoal were observed in the matrix.

*Burial 95.* This burial comprised the plain unlined burial of a young adult male (**CGG_2_104304**). This individual was oriented along a NS axis with the head towards the north facing west. The burial appeared to have been disturbed when the pit for the stone-lined grave of burial 67 was cut. This, however, only seems to have clipped the lower legs of the individual of burial 95 and care seems to have been taken in placing the tibiae and fibulae back in with the body in approximately the correct position (albeit the wrong way around), behind the eastern wall of the stone-lined grave of burial 67. The individual was lying on the right side with the no-longer flexed legs now laying with the knees towards the west and the feet proceeding towards the S/E. The left arm was tightly flexed. The individual was buried in a charcoal-flecked grey fill.

**Stratum 6**

*Burial 63.* This burial consisted of an unusually large sub circular grave into which was placed a very large jar burial containing the remains of at least two individuals. This large jar burial with a decorative incised rope decoration around its circumference was aligned along an EW axis with the jar neck towards the east. It contained the partially articulated body remains (A) of an adolescent of indeterminate sex placed in the upper fill above the articulated remains of a middle adult female 30 to 45 years old (B). Whilst there seems to have been a preference for jar burials to contain infants, a practice also observed for the late Chalcolithic of the Jordan valley and in the Middle Bronze Age in Palestine, evidence from Sidon suggests that the custom of jar burials was not necessarily confined to certain age ranges but may have also involved not only infants and children but adults as well. The body of 63B (**CGG_2_104302**) was aligned similarly east-west within the jar with the head to the east, facing west and the body flexed/curled on its left side.

*Burial 73.* This interment consisted of a highly fragmented and disarticulated burial of a 6-month-old infant in a jar (**CGG_2_104298**) . The fragmented cranium was located in the south of the jar. The cranium was inferiorly oriented and faced east. The left femur and os coxae were observed in the north of the burial. The right femur, fragmented ribs and other unidentified long bones were observed in the west. Post depositional movement was substantial. The mandible and several vertebral hemiarches were retrieved from inside the cranial space.

**The Iron Age**

*Burial 91.* A group of re-deposited remains dating back to the Iron Age. Some long bones were piled together (at least 2 tibiae orientated at opposite ends to each other; 2 fibulae). A large fragment of an ossa coxae (pelvis) was found next to the proximal head of a right femur (unfused head). The remains were mixed with animal bone. The sample from this burial is a petrous bone (**CGG_2_104309**)

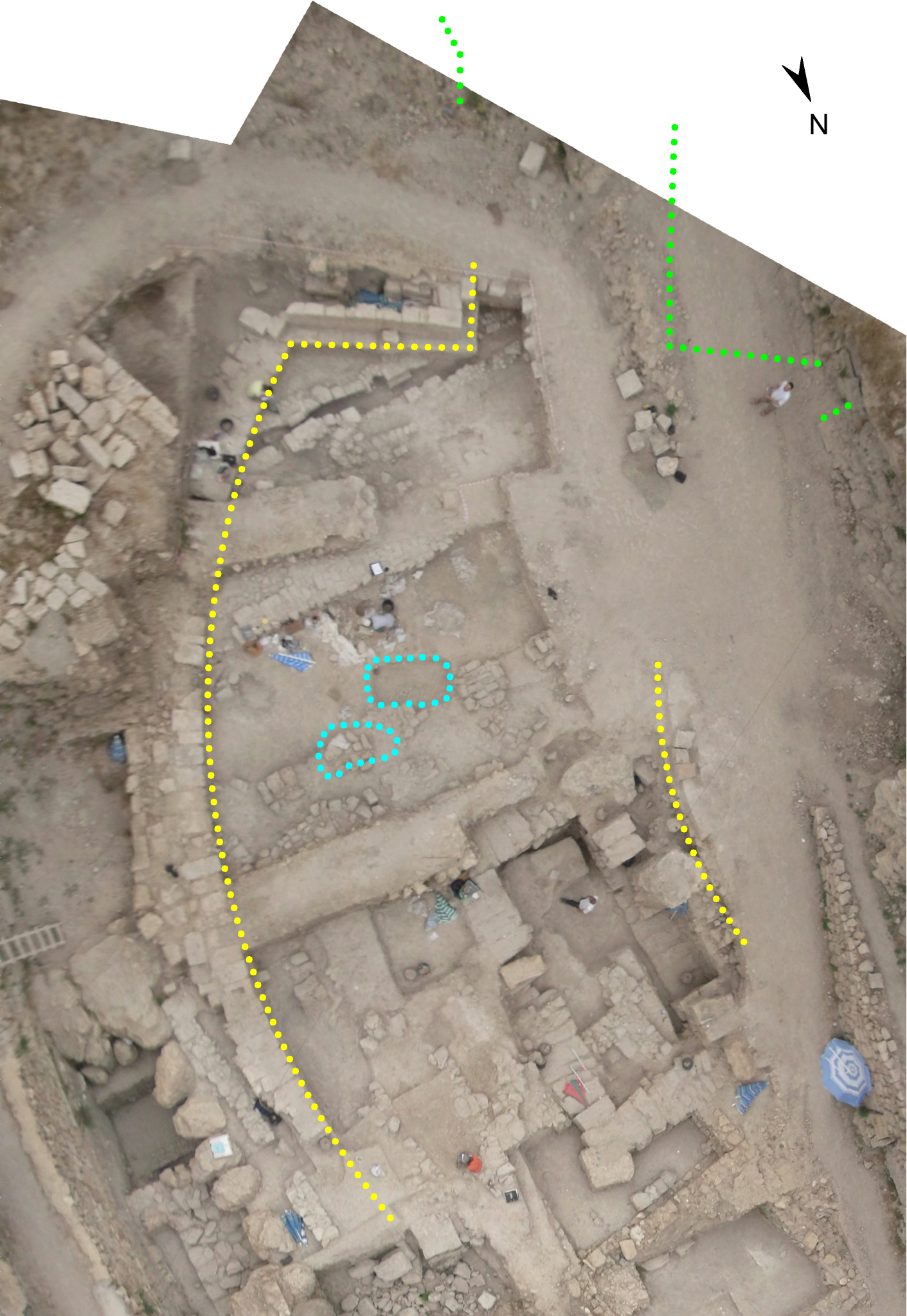

Fig. 8. The Medieval city wall on either side of the Medieval moat where burials dating from the Crusader period were found (courtesy of Claude Serhal).

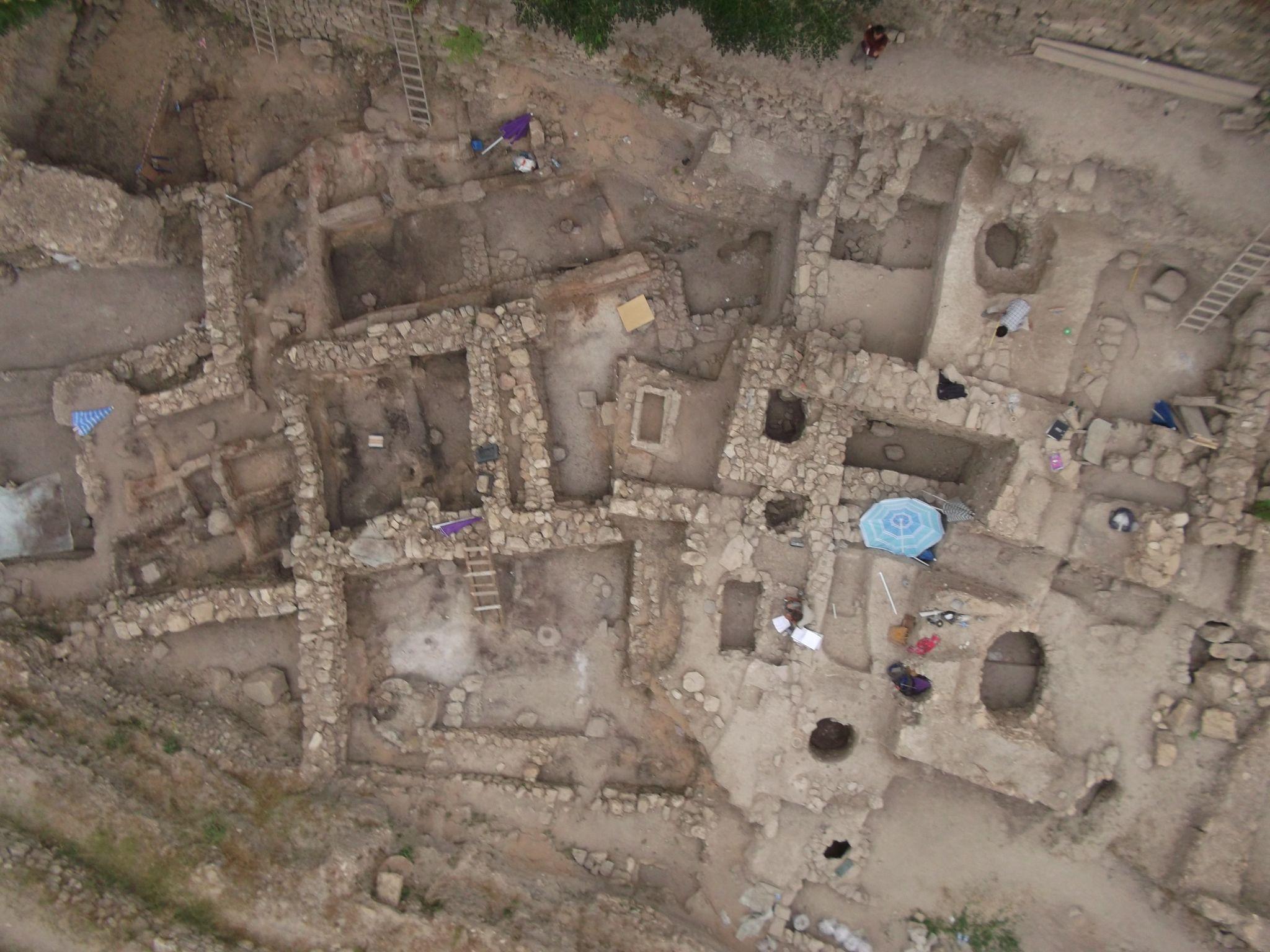

Fig. 9. Aerial view of the “College site” (courtesy of Claude Serhal).

#### **2.9. Moldova**

##### ***2.9.1. Balabanu II, Taraclia district, southwestern Moldova***

Latitude: 45.92666667; Longitude: 28.59194444

Sample provider: Alexander Varzari, Eugen Sava

Alexander Varzari, Igor V. Deyneko

In total, 9 mounds to the north-east of the village Balabanu were excavated in 1984^151^. They are located in the valley of the Yalpug river, on the right-hand side, in the Budjak steppe (western part of the Pontic–Caspian steppe zone). The excavations were carried out under the direction of E. Sava.

**Balabanu II 1984 T6M2 (=Kurgan 6, grave 2)**

The kurgan 6 was partially destroyed by the installation of a geodetic survey tower, which was removed before the excavation started. By the beginning of the excavation, the height and the diameter of the kurgan were c. 1 and 36 metres respectively. Five burials/graves were found in the kurgan.

The Grave/Burial 2 (Multi-cordoned Ware culture [KMK], archaeologically dated to 2100-1600 BCE) was in the northeastern sector of the mound. The burial pit was roughly circular in shape (1.92 m in diameter; 1.14 m deep) with a ledge. The burial chamber had an oval layout (1.46 x 0.88 m; 0.6 m deep), oriented SW-NE. Under the skeleton, the trace of a decayed rug was recorded.

The individual presented in this study (**CGG_2_103663**) was found lying in a crouched position on its left-hand side (left-sided crouched), SW-NE oriented, skull to the NE. Anthropological determination: male, aged 50‐55.

**Balabanu II 1984 T7M2** **(= Kurgan 7, grave 2)**

The kurgan 7 is located near the village Balabanu and belongs to the same group of burial mounds as the kurgan 6. At the time of excavation, kurgan 7 had a preserved diameter of 30 m and a height of 0.2 m. The mound covered two burials.

The Grave/Burial 2 (Multi-cordoned Ware culture [KMK]) was a primary grave archaeologically dated to 2100-1600 BCE. It was found in the eastern sector of the mound/kurgan. The burial pit was oval-shaped (1.4 x 0.9 m; 0.7 m deep), oriented SW-NE, with vertically cut sides.

The individual presented in this study (**CGG_2_103667**) was found lying in a crouched position on its left-hand side (left-sided crouched), SW-NE oriented, skull to the NE. Anthropological determination: male, aged 50‐60.

##### ***2.9.2. Cahul/Crihana Veche, Cahul district***

Latitude: 45.85833333; Longitude: 28.255

Sample provider: Alexander Varzari

Alexander Varzari, Igor V. Deyneko

Cahul/Crihana Veche was excavated under the direction of S.I. Kurceatov in 1992^152^. The name of the site is Cahul/Crihana Veche 1992 T1M6.

Kurgan 1 near the village of Crihana Veche was part of a group of three kurgans. The group of kurgans was located at a distance of 3 kilometres North-East from the village Crihana Veche and 1.5 km North-West from the Cahul city airport on the southwestern edge of the Southern Moldovan Plain in the West Pontic grass steppe zone. The kurgan 1 with a height of 1.5 m and a diameter of 60 m contained 23 burials.

The chamber/burial 6 (Yamnaya culture, probably late phase, archaeologically dated to 2600-2200 BCE) is a secondary burial found in the southwestern sector of the mound. The burial chamber was of irregular rectangular shape with slightly rounded corners (chamber size: 1.8 x 0.95 m, 0.6 m deep), SW-NE oriented. The burial 6 shared a common burial pit/grave (pit size: 3.6 x 2.2 m) and a common ledge platform (0.45 m deep) with the burial 5 (Yamnaya culture), SW-NE oriented. Two burials were at 0.4 m distance from each other with the burial 6 located on the NW part and the burial 5 – on the SE part of the grave. Dark brown rot/decay is marked at the bottom of the graves.

The individual sampled in this study (**CGG_2_103620**) was buried on its back with bent legs (crouched position on the back), SW-NE oriented, skull to the NE. The skeleton was slightly coloured with red ochre. Anthropological determination: male, aged 40‐45.

##### ***2.9.3. Cazaclia, Ceadîr-Lunga district, Southwestern Moldova***

Latitude: 45.97313889; Longitude: 28.62277778

Sample provider: Alexander Varzari, Sergey Agulnikov

Alexander Varzari, Igor V. Deyneko

The Сazaсlia mound group was located on a watershed hill between the Yalpug and Lunguta rivers, 3.5 km southwest of the village of Cazaclia and was extended in a chain up to 6 km long in a SE - NW direction. The studied group included 29 burial mounds. The site is situated on the Budjak steppe, which is a western part of the North Pontic Lowland. It was excavated under the direction of S. Agulnikov in 1984^153^.

**Cazaclia 1984 T6M5 (= Kurgan 6, grave 5)**

The kurgan 6 had a maximum height of 0.7 m and a diameter of roughly 40 m, and contained 6 burial pits (graves). The Grave/Burial 5 (Multi-cordoned Ware culture [KMK], archaeologically dated to 2100-1600 BCE) was in the northeastern sector of the mound.

The burial consisted of a rectangular pit 2.25 m long, 1.9 m wide and 0.2 m deep, SE-NW oriented. The burial chamber was within the pit, 1.57 by 1.0 m in size, and was surrounded by a ledge of 0.6 m deep. At the level of the ledge, the chamber was roofed by 7 wooden logs, laid across. The walls of the pit were covered with a white coating. The skull and the bottom of the pit were coloured brown by decay/rot.

A biconical vessel with a curved corolla was near the skull at the SE corner of the chamber. The surface of the vessel is grey with black and yellow spots. The ceramic dough/compound is black with an admixture of fine chamotte. The vessel’s dimensions are: height - 18.5 cm, bottom diameter - 10 cm, body diameter - 17 cm and corolla diameter - 14.5 cm. Next to the vessel, a fragment of a treated limestone of a rectangular shape, 6 cm long and 3.5 cm wide, was found.

The individual presented in this study (**CGG_2_103707**) was found lying in a crouched position on its left-hand side (left-sided crouched), skull to the S-SE. Anthropological determination: male, aged 50‐60.

**Cazaclia 1985 T5aM2 (= Kurgan 5a, Grave 2)**

The kurgan 5a belongs to the same Cazaclia group of burial mounds as the kurgan 6 (Ceadir-Lunga district, southwestern Moldova). The kurgan with a height of 0.6 m and a diameter of 25 m consisted of two burial pits (graves). The Grave/Burial 2 (Yamnaya culture, probably Late, archaeologically dated to 2600-2200 BCE primary/central) was found in the northern sector of the mound. The form of the pit was roughly rectangular, with dimensions of 1.36 × 1.0 m, vertical walls and a depth of 0.75 m from the identification/fixation level, E-W oriented. The bottom of the burial pit was covered by dark-brown decay/rot.

The individual presented in this study (**CGG_2_103703**) was found lying in a crouched position on its right-hand side (right-sided crouched), skull to the E. The bones, and especially the skull, were coloured with brown-red ochre pigment. Anthropological determination: male, aged 50‐55.

##### ***2.9.4. Constantinovca, Slobozia district, Transnistria, southeastern Moldova***

Latitude: 46.85958333; Longitude: 29.84888889

Sample provider: Alexander Varzari, Sergey Agulnikov

Alexander Varzari, Igor V. Deyneko

The Kurgan was situated at a distance of 250 m from the northern outskirts of the village Constantinovca on the watershed hill between the Dniester and Kuchurhan rivers in the West Pontic grass steppe zone. Base diameter and height of the kurgan were 45 m and 1.2 m, respectively; it contained 13 burials and was excavated under the direction of S. Agulnikov in 1987^154^.

Grave/Burial 6 (**Constantinovca 1987 T1M6 [ =Kurgan 1, grave 6]**) - Yamnaya culture, probably late (archaeologically dated to 2600-2200 BCE). A molar (**CGG_2_103753**) was sampled from this individual. Anthropological determination: male.

Grave/Burial 8 (**Constantinovca 1987 T1M8 [= Kurgan 1, grave 8]**) - Yamnaya culture, probably late (archaeologically dated to 2600-2200 BCE). Grave 8 is a secondary burial and was identified in the southwestern sector of the mound. It was trapezoidal in shape (pit size: 1.7 x 1.3 x 1.2 x 1.05 m; 0.25 m deep), NW‐SE oriented. The bottom of the burial pit was covered by dark-brown decay/rot. The body was laid on its back and inclined on its left side (crouched position on the back inclined on the left), skull to the NW, legs strongly crouched on its left side, arms slightly bent towards the pelvis. The bones, and especially the skull and the left arm, were deep-red stained with ochre. A molar (**CGG_2_103750**) was sampled from this individual. Anthropological determination: male, aged 55‐70.

##### ***2.9.5. Cotiujeni, Șoldănești district, Northern Moldova***

Latitude: 47.86777778; Longitude: 28.57583333

Sample provider: Alexander Varzari, Sergey Agulnikov

Alexander Varzari, Igor V. Deyneko

A group of 5 kurgans was located at a distance of 2.5 km northeast from the village Cotiujeni (Șoldănești district) on a watershed hill in the forest-steppe zone of northern Moldova. With a height of 1.5 m and a diameter of 44 m, the kurgan 1 was the biggest one in the group. Eight burials were found in the mound. The excavation was conducted under the direction of S. Agulnikov in 1986^155^.

Burial 5 (**Cotiujeni 1986 T1M5 [= Kurgan 1, Grave 5]** - Yamnaya culture, probably late archaeologically dated to 2600-2200 BCE) was a secondary burial in the northeastern sector of the mound. The pit was rectangular in shape with slightly rounded corners (pit size: 1.58 x 1 m, 0.35 m deep), SWW-NEE oriented. Dark brown rot/decay is marked at the bottom of the grave. In the north-western part, the burial was overlapped by another burial (Grave 4, Yamnaya culture).

A scraper of light grey flint was found 0.15 m to the right of the skull.

The individual presented in this study (**CGG_2_103674**) was found lying in a crouched position on its left-hand side (left-sided crouched), skull to the NE-E. The right arm is bent with the hand on the knee; the left arm is straight; legs bent at the knee. Anthropological determination: presumably a woman, aged 40-50.

##### ***2.9.6. Taraclia, Taraclia district, southern Moldova***

Latitude: 45.91877778; Longitude: 28.62833333

Sample provider: Alexander Varzari, Sergey Agulnikov, Eugen Sava

Alexander Varzari, Igor V. Deyneko

The kurgan group near Taraclia city was located to the south of the Cazaclia mounds, forming a single complex of mounds along the watershed between the Yalpug and Lunguta rivers. The excavation was carried out under the direction of S. Agulnikov and E. Sava between 1980-1984^153^.

**Taraclia 1982 T3M14 (= Kurgan 3, grave 14)**

The kurgan 3 measured 40 m in diameter, had a height of 1.1 m and contained 19 burials. The Grave/Burial 14 (Multi-cordoned ware [KMK]/Sabatinovka culture, archaeologically dated to 2100-1200 BCE, secondary) was in the southern part of the mound 3. The burial pit was of an irregular rectangular shape, 1.1 m long, 0.8 m wide and 0.2 m deep from the identification/fixation level, E-W oriented.

The individual presented in this study (**CGG_2_103777**) was found lying in a strongly crouching position on its left-hand side (left-sided crouched) with the skull to the E and the femoral bones touching the elbow joints. Anthropological determination: male, aged 45‐55.

**Taraclia 1982 T3M16 (= Kurgan 3, grave 16)**

The Grave/Burial 16 (Multi-cordoned ware [KMK]/Sabatinovka culture, archaeologically dated to 2100-1200 BCE, secondary) was in the southern part of the kurgan 3, at a distance of 2 metres or less from the burial 1s4. The pit contour was not fixed.

The individual presented in this study (**CGG_2_103781**) was found lying in a strongly crouching position on its left-hand side (left-sided crouched), with the skull to the E, the left femur touching the bones of the forearms and the hands placed in front of the face. Anthropological determination: male, aged 30‐35.

#### **2.10. Spain**

##### ***2.10.1 Argar, Antas, Almería, Andalusia***

Latitude: 37.24741; Longitude: -1.9137

Sample provider: Alfredo Mederos Martín, Victoria Peña Romo, Nicolas Cauwe

Alfredo Mederos Martín

The El Argar settlement is located next to the town of Antas (Almería), occupying a 2.19-hectare plateau next to the Antas River. It was excavated by Louis and Henri Siret (1887) between 1884 and 1889. Under the houses, 1032 tombs were found in urns, cists and vaults. Most of it is preserved in the MRAH in Brussels, tombs 1-972, and the last 155, tombs 973-1032^156^, excavated between 1887 and 1889, in the National Archaeological Museum, Madrid. Being the site with the largest number of tombs studied from the Early Bronze Age of the Southeast Iberian Peninsula, the name El Argar is also used to define the main archaeological group of the period. After the Siret brothers, archaeological excavations took place at the site only once in 1990^157^. The dates of the graves, carried out within the project, range between 2135-1625 BCE.

Tomb 153 is a burial in an urn or pithos, without grave goods^158^, which contains the remains of a mature adult (**CGG_2_022310/Rise 1515**), probably a man, between 30 and 55 years old. Unidentified bones of a 3-year-old infant^159^ have also been reported from this context. The DNA sample was taken from a lower premolar.

##### ***2.10.2. Can Martorell, Dosrius, Barcelona, Catalonia***

Latitude: 41.594; Longitude: 2.414

Sample provider: Juan Francisco Gibaja

Oriol Mercadal (┼), Sara Aliaga, Juan F. Gibaja, Maria Eulàlia Subirà, Bibiana Agustí

The site of Costa de Can Martorell (Dosrius, Barcelona) is located in the northeast of the Iberian Peninsula. It is a hypogeum or artificial sepulchral cave with a megalithic door and a corridor that leads to a 7 m² circular burial chamber^160^. It was excavated 1995-1996. 227 individuals lay buried in the grave; many of them would have died in youth and early adulthood^161^. Grave goods are almost exclusively arrowheads, up to 68, many of which present fractures produced by their use as projectiles^162^.

14C dating of four human samples from the vestibule (done by the sample providers) gave the possibility to date the use of the structure during the period 2640-2060 cal. BCE. It is believed that part of the buried died due to an episode of violence^160^ because of the age of the individuals, the position of the arrowheads relative to the bodies (many of them under the bodies).

Three out 26 samples provided results (SJ345 UE9085, CM28, and CM34) from this site:

SJ345 UE9085 (**CGG_2_021255/Rise 1345**), dental pieces

CM28 (**CGG_2_021260/Rise 1350**), dental pieces 23-46

The sample preserves maxillary and mandibular remains. We determined that they belong to a young adult with masculine features.

CM31 (**CGG_2_021274**), dental pieces 17-18

The sample preserves maxillary and mandibular remains. We determined that it corresponds to a young adult without sexual markers.

CM34 (**CGG_2_021277**), dental pieces 47-48

The sample preserves maxillary and mandibular remains. It has a type A *Criba Orbitalia*. We determined that it corresponds to an adolescent.

##### ***2.10.3. Cantorella, Maldà, LLeida province, Catalonia***

Latitude: 41.552; Longitude: 1.022

Sample provider: Andreu Moya i Garra, Juan Francisco Gibaja

Òscar Escala Abad, Andreu Moya i Garra, Enric Tartera Bieto, Ares Vidal Aixalà, Núria Armenatano Oller.

Cantorella is located on an old river terrace of the Corb, in the lowlands of the Segre valley, in the northeast of the Iberian Peninsula. The site was excavated in 2010 and 2011 as a result of an archaeological rescue intervention. It is an open air settlement with two phases of occupation during the Late Neolithic and the Early Bronze Age. The most characteristic structures of both periods are the storage pits, which are often reused as waste pits.

During the recent phase, eight of the silos were reused as funerary tombs, most of which become collective graves. The anthropological study of the group has established a minimum number of 43 individuals. Two of them have been radiocarbon dated by the excavators to the middle of the first half of the 2nd millennium BCE or 1886-1689 cal. BCE (courtesy of the sample providers).

The depositions generally correspond to primary burials, although secondary deposits are also found as isolated remains. The registry describes a group with the presence of both sexes and practically all age categories. Adult individuals predominate, although most did not reach maturity. The documented skeletal pathologies correspond to degenerative processes associated with continued physical activity. From an anthropological point of view, common and proper processes and alterations of collective graves and empty space depositions are documented. In general, the anatomical positions of the skeletons are followed, although accumulation and interlocking of the remains is important. From an taphonomic point of view, the disarticulation of entire anatomical parts and the alteration and manipulation of some isolated bone remains were confirmed. The disorder and neglect that the bodies present suggest posthumous actions and intentional disturbances derived from sepulchral overtures that involved cadaverous manipulations.

The osteological analyses conducted on the samples used the following terminology/Age groups: Infans 1 (Inf 1) >7 years; Infans 2 (Inf 2) 7-14 years; Juvenilis (Juv) 14-20 years; Adultus (Ad) 20-40 years; Maturus (Mat) 40-60 years; Senilis (S) >60 years; UND: indeterminate.

Four samples from this site are included in this study. They come respectively from Silo J-29, SJ-31, Sj-42 and Silo SJ-47.

**Silos SJ-29**

Individual 9 or 29-1230 (**CGG_2_021206/Rise 1296**), who is an adult (c. 35-42 years old) female

**Silos SJ-31**

Individual 31-1158 (**CGG_2_021209/Rise1299**), who is an adult female, c. 30-40 years old.

**Silos SJ-42**

Individual 42-1189 (**CGG_2_021211/Rise 1301**), who is an adult, probably female ≤30 years old

Individual 42-1248 (**CGG_2_021213/Rise 1303**), who an adult individual, probably female

Individual 42-1250 (**CGG_2_021215/Rise 1305**), who is Infans I (>7 years) of undetermined sex

Individual 42-1264 (**CGG_2_021214/Rise 1304**), who is Infans I (4±1 years old) of undetermined sex

**Silos SJ-47**

Individual 47-1151 (**CGG_2_021218/Rise 1308**), who is an adult, probably male, c. 35-45 years old

##### ***2.10.4. Castellon Alto, Galera, Granada, Andalusia***

Latitude: 37.7400778; Longitude: -2.565888

Sample provider: Alfredo Mederos Martín, Victoria Peña Romo, Fernando Molina González, María Oliva Rodríguez Ariza,

Alfredo Mederos Martín

The settlement of Castellón Alto (Galera, Granada), is located on the left bank of the Castillejar river. It has an estimated size of 0.50 ha. It was excavated by the University of Granada in 1983^163^, and then again for several months between 2001 and 2002. The last excavation included the restoration of the site^164^. The houses are distributed in several terraces of the steep hill. Under the houses, 130 tombs have been documented, of which about 40 had been looted. The dates of the site oscillate between 2030-1660 BCE ^165^. The samples come from three double or reused graves (tombs 11, 36 and 93).

**Tomb 11**

Tomb 11 was excavated in 1983. It contained the two adult burials, possibly one male and one female. The sample was obtained from the left third molar of 11A, the adult male (**CGG_2_023797/Rise 1972**).

**Tomb 36**

Tomb 36 contained the remains of three individuals, a possibly male adult (36A), from which the lower left second molar was sampled (**CGG_2_023799/Rise 1974**), and two mature adults, between 30 and 55 years old, possibly male and female.

**Tomb 93**

Tomb 93, which lacked grave goods, also contained two individuals, an adult, perhaps male, (93A), from which a sample of the upper right third molar was taken (**CGG_2_023795/Rise 1970)**, and an infant I, less than 6 years old.

##### ***2.10.5. Cerro de la Virgen, Orce, Granada, Andalusia***

Latitude: 37.726667; Longitude: -2.513889

Sample provider: Alfredo Mederos Martín, Victoria Peña Romo, Fernando Molina González, Juan Antonio Cámara

Alfredo Mederos Martín

The fortified settlement of Cerro de la Virgen is an important site during the Bell Beaker period and the subsequent Early Bronze Age. It has an estimated size of 1.20 ha and lies close to the Orce River. It was initially excavated in four campaigns (1963, 1965, 1967 and 1970) led by W. Schüle and M. Pellicer^166^, followed by a more recent one in 1986 (Sáez, Schüle, 1987). It presents a significant Bell Beaker occupation with large circular huts^166,167^, which was replaced during the Early Bronze Age by rectangular houses typical of the Argaric groups^168^. The dates of the Bell Beaker phase range between 2450-2200 BC^169^, while those of the Bronze Age tombs are between 2225-1625 BC^168,170^. The samples come from tombs 6, 8, 15, 21, 26, and 32).

**Tomb 6** is a double burial with two adult individuals. The sample was obtained from the individual 2, an adult female (**CGG_2_024002**). The sample has been radiocarbon dated by the excavators to 3500±35BP (Ua39402^168^), cal. BP 3873-3648 (see Suppl. Tab. S9).

**Tomb 8** is the deposition of an adult individual (**CGG_2_024003**). The grave has been 14C dated by the excavators to 3426±34 BP (Ua39403^168^), cal. BP 3825-3571 (see Suppl. Tab. S9).

**Tomb 15** is a double burial. The sample was obtained from the first lower left molar of individual 15A (**CGG_2_023764/Rise 1939**), possibly a young adult female, c. 19-29 years old. The sample has been 14C dated by this project to 3490±15BP, cal. BP 3831-3698 (see Suppl. Tab. S9). It was accompanied by a mature adult (between 30 and 55 years old), possibly male.

**Tomb 21** is a monumental tomb with one older deposition at the bottom of individual 21A (adult/elderly masculine of c. 60-65 years) on which a second tomb is allocated at a later stage. The second deposition belongs to an elderly female individual of c. 55-60 years. Our sample is supposed to belong to the earliest deposition (**CGG_2_024001**) which has been 14C dated (Ua39410) by the excavators to 3586±36 BP^168^, cal. BP 3984-3727 (see Suppl. Tab. S9).

**Tomb 26** is the deposition of an adult/mature male individual (**CGG_2_023768**). The grave has been dated by means of 14C (Ua39415) to 3429 ± 31 BP^168^, cal. BP 3824-3574 (see Suppl. Tab. S9)

**Tomb 32** is the deposition of an infant II c. 7-12 years old. The lower left second molar of the individual has been sampled for this project (**CGG_2_023770/Rise 1945**). The grave has been dated by means of 14C (Ua39422) to 3406 ± 30BP^168,170^, cal. BP 3817-3567 (see Suppl. Tab. S9).

##### ***2.10.6. Cerro de la Encantada, Granátula de Calatrava, Ciudad Real***

Latitude: 38.81688154137975; Longitude: -3.7288116650308187

Sample provider: Alfredo Mederos Martín, Victoria Peña Romo, Catalina Galán, José Sánchez Meseguer and Armando González

Alfredo Mederos Martín

Encantada hill is a fortified settlement from the Bronze Age identified in 1976, which rises 75 m above the Jabalón river valley. Between 1977 and 1978 two major excavation campaigns were carried out by Nieto Gallo and Sánchez Meseguer (1980) published in a monograph with the first 7 tombs. New annual excavation campaigns were carried out between 1979 and 1991, identifying up to tomb 52. Excavations resumed with regular annual excavation campaigns on the fortification wall between 1998 and 2009, which make it, together with the Motilla del Azuer, the most important Bronze of La Mancha site. Within the settlement, 83 graves have been located in pits, cists and pithoi, of which the first 37 were the subject of a preliminary assessment^171^ and more recently a detailed study of the children's graves has been published^172^. It presents two large levels of occupation, stratum II dated ca. 2050-1875 BCE and stratum IIIa-b ca. 1875-1600 BCE^173,174^. The sample comes from tomb 37B (**CGG_2_023754**), excavated in 1982, corresponds to a young man between 19-29 years old, also lacking funerary goods. The sample has been 14C dated by this project to 3410±20BP, cal. BP 3811-3573 (see Suppl. Tab. S9).

##### ***2.10.7. Cuesta del Negro, Purullena, Granada, Andalusia***

Latitude: 37.33543878810364; Longitude: -3.2326500014826887

Sample provider: Alfredo Mederos Martín, Victoria Peña Romo, Fernando Molina González, Syvia Jiménez Brobeil

Alfredo Mederos Martín

The settlement of Cuesta del Negro is close to the Fardes river. It has an estimated size of c. , with 1.50 ha. It was excavated by the University of Granada in three campaigns between 1971 and 1972. Only grids 4-7 and 10 have been published^175^ and part of the data from the necropolis^176^. The chronology of the necropolis, where 36 tombs have been documented^177^, extends between 1850-1600 BCE, characterising the Middle-Late Bronze Age phase of the site, until, but just a few burials, 1550-1400 BCE^165^.

The analysed sample comes from grave 4, which was in a vault and contained a double burial. Both individuals were 14C dated^165,178^. The first individual 4A (**CGG_2_023745/Rise1920**) was radiocarbon dated to 3375±32BP, cal. BP 3569-3412 (see Suppl. Tab. S9). Individual 4A is a mature man between 30 and 55 years old, with a set of two silver earrings, a copper dagger and a ceramic bowl. The sample analysed in this project is the lower right third molar of this individual 4A. The second individual (4B) from the grave is an infant II (c. 7-12 years old), who had a silver earring, a bottle and a pedestal cup, and was perhaps buried somewhat later than 4A^165,178^.

##### ***2.10.8. Fuente Álamo, Cuevas del Almanzora, Almería, Andalusia***

Latitude: 37.32945; Longitude: -1.858202

Sample provider: Alfredo Mederos Martín, Victoria Peña Romo

Alfredo Mederos Martín

The settlement of Fuente Álamo is located on a high hill that rises 65 m above the surrounding territory, and organised in stepped terraces. It has an estimated size of 1.83 ha. It was excavated and published by Louis and Henri Siret (1887) in the same year 1887. They documented 48 graves. New investigations were resumed in 1977 by the German Archaeological Institute, carrying out 7 excavation campaigns in 1977, 1979, 1985, 1988, 1991, 1996 and 1999 led by H. Schubart, O. Arteaga and V. Pingel^179–181^. The new excavations identified 64 new graves in cist or urn up to grave 112^182^, which make it the best-studied site in the Argaric group of the Iberian southeast. The chronology of the site ranges from 2450-1625 BCE^169,179^.

While other Argaric collections are well preserved in the Museum at Brussels, there are hardly any identifiable bones from the first 48 tombs excavated at Fuente Álamo in 1887. It is therefore difficult to assign the osteological remains to specific contexts. The grave from which our sample was taken lacked a number and was given the number 0. The sample (**CGG_2_022307/Rise 1512**) is a third lower molar of a mature adult, perhaps female.

##### ***2.10.9. Los Torcales 6, Beas, Huelva, Andalusia***

Latitude: 37.426801153027; Longitude: -6.8076867749241

Sample provider: Ana Pajuelo Pando, Alfredo Mederos Martín, Victoria Peña Romo and José Miguel Morillo León

Serena Sabatini, Alfredo Mederos Martín

The site called Los Torcales is located in the village of Beas, Huelva province. The site was excavated in 2015^183^ and has been divided into various subsites, Los Torcales 1 and 6, with the classical cist tombs of the Middle Bronze Age of SW Iberia. The burials from which our individuals were samples come from the sub-site called Los Torcales 6. Four burial contexts were excavated in Los Torcales 6: Cista 2, 3, 4 and 5. The only available 14C date by this project for the site corresponds to the individual 2 in Cista 2, which provided a date to 3420±30 BP or 1870-1631 (2σ cal.) BCE^183^.

Our samples belong to individual 1 in Cista 2 (**CGG_2_021224/Rise 1314**) and individual 1 in Cista 3 (**CGG_2_021225/Rise 1315**).

##### ***2.10.10. Minferri, Juneda, LLeida, Catalonia***

Latitude: 41.557062679604584; Longitude: 0.8380653764285175

Sample provider: Joan B. López Melcion (1957-2020), Natàlia Alonso Martínez, Andreu Moya i Garra, Bibiana Agustí Farjas

Serena Sabatini

Minferri is located on an old river terrace of the Femosa, in the lowlands of the Segre valley, in the northeast of the Iberian Peninsula. The site has been intermittently excavated from 1993 to 2006 by the Grup d’Investigació Prehistòrica de la Universitat de Lleida (GIP-UdL). Various publications about the site are available^184–186^.

It is an open air settlement that corresponds to the scattered village pattern with two phases of occupation during the Late Neolithic and the Early Bronze Age. The most characteristic structures of both periods are the storage pits, which are often reused as waste pits.

During the recent phase, dated by 14C throughout the first half of the 2^nd^ millennium (2132-1456 2σ cal. BCE) 24 of the silos were reused as funerary tombs, most of which become collective graves. The funerary register of Minferri is the most extensive in the western Catalan plain for the Early Bronze Age. The anthropological study of the group has established a minimum number of 56 individuals.

The population group is made up of both female (or probably female) and male (or probably female) individuals (n. 18 and 14, respectively), with a high rate of undetermined individuals (n. 24). All age categories with a predominance of adults (n. 15) and matures (n. 14) are represented, a significant presence of perinatal (n. 5) and children (Inf. 1: n. 8; Inf. 2: n. 9), but with a low representation of young people (n. 4) and only a single case of a senile individual.

The burial record reveals a wide variety of funerary practices. There is no single pattern of skeletal positions –lateral decubitus, supine, prone, sitting– although in the niches all the bodies are lateralized and with forced positions. In addition, possible bundle deposits are identified and decompositions are documented in both empty and filled spaces. On the other hand, the most common is that the buried individuals are not associated with other types of depositions, so the presence of funerary trousseaux and offerings is not a common or widespread phenomenon within the Minferri funeral rite. The documented examples are the deposition of ceramic vessels and meat offerings, which distinguishes and gives meaning to the burials with which they are associated. The ceramic vessels are deposited near the bodies. Instead, the deposits of fauna in a sepulchral context are formalised in the same silos with burials or in structures adjacent to the tombs. They mainly correspond to the deposition of complete animals or partial remains in anatomical connection of cattle, ovicaprid, canids and, singularly, some birds. Ten individuals were sampled for this study (Table 9).

| **Context/Sample** | **CGG_2_ nr.** | **Rise nr.** | **Osteological age/sex** |
| --- | --- | --- | --- |
| SJ418, UE5312 | CGG_2_021233 |  | Maturus (30-40 y)/F |
| SJ418, UE5311 | CGG_2_021234 |  | Infans 1 (2-3 y)/Und. |
| SJ418, UE5310 | CGG_2_021235 |  | Infans 1 (6-7 y.)/Und. |
| SJ418, UE5264, EN 421 | CGG_2_021237 |  | Maturus (40-50 y.)/Probably F |
| SJ296, UE8587 | CGG_2_021240 | 1330 | Maturus (40-60 y.) /Probably F |
| SJ385, UE5152 | CGG_2_021247 |  | Adultus (40-60 y.)/F |
| SJ373, UE 10041 | CGG_2_021249 | 1339 | Adultus (30-40 y.)/M |
| SJ95, UE7086 | CGG_2_021253 | 1343 | Maturus (40-50 y.)/M |
| SJ354, UE9080 | CGG_2_021254 |  | Juvenilis (15-17 y.)/Probably F |
| SJ354, UE9085 | CGG_2_021255 |  | Maturus (50-60 y.)/M |

Table 9. Synoptic table with data about the sampled individuals from the burials at Miniferri and their context of origin.

Sample **CGG_2_021249** has been 14C dated in this study to 3400±15 BP, cal. BP 3692-3575 (see Suppl. Tab. S9), supporting the archaeological chronology of the site in the first half of the 2^nd^ millennium BCE.

##### ***2.10.11. Motilla del Azuer, Daimiel, Ciudad Real province, Castilla y La Mancha***

Latitude: 39.043267; Longitude: -3.497400

Sample provider: Alfredo Mederos Martín, Victoria Peña Romo, Trinidad Nájera Molina, Fernando Molina González

Alfredo Mederos Martín

The site at Motilla del Azuer is one of the two most important Bronze Age settlements in La Mancha region. It is a fortified site, with two walled enclosures that surround a central tower. The first stage of the excavations was carried out between 1974 and 1986^187,188^. Those investigations (between 1982 and 1984) also identified a deep well in the central patio. Field investigations were resumed between 2000-2010, with long annual campaigns that included the restoration of the tower and of the surrounding walls. Altogether there have been 24 annual excavation campaigns^189^.

63 graves have been excavated at the site. They contained a total of 65 individuals, which indicates that, with few exceptions, they are individual graves^190,191^. The graves were generally located under the houses on the outer perimeter, and some within the fortified enclosure. The dates of the fortified enclosure range between 2200-1600 BCE, with a final phase well into the Middle-Late Bronze Age between c. 1600 and 1350 BCE^192^.

Tomb 60 (G01) was excavated in 2008, and corresponds to a young adult male, between 19-20 years old. The only grave good in the burial is a globular vessel. The context has been 14C dated (Ua-38416) to 3591±37BP, cal. BP 4065-3728 (see Suppl. Tab. S9). The body shows various wounds produced by a dagger^193^. The sample (**CGG_2_023808/Rise 1983**) was taken from the lower third molar.

##### ***2.10.12. Necrópolis de los Algarbes, Tarifa, Cádiz***

Latitude: 36.075567317771046; Longitude: -5.699202386612876

Sample provider: V. Castañeda, J.M. Morillo León

Vincent Castañeda, Josè M. Morillo León

The so-called Necropolis of Los Algarbes is a funerary site from Late Chalcolithic- Early Bronze Age (second half of the III millennium, according to the 14C dates obtained) at the southernmost side of the Iberian Peninsula (Tarifa Cadiz).

While the site was long known since the *hispanista* P. Paris identified in the decade of 1920, no particular attention was paid to it until forest-related works unearthed a probable hypogeum burial in 1963^194^.

During the 1970s, and then in the 1990s, conservation and rescue operations were carried out. It was, however, not until 2012-2014 that a proper scientific research project was implemented. The material studied in this project resulted from the outcomes of this modern project^195^.

The necropolis by itself comprise three types of burials: several individual pits excavated in the rock, with roughly anthropomorphic shape, hypogea with lateral entrance, also excavated in the rock, and one pseudo megalithic grave partially excavated in the bedrock (Structure 1-2). The latter seems to be the main funerary structure of the site, acting as an articulation element of the necropolis.

While the most common ritual is the collective burial, there is at least one case of individual burial in the Structure 1-2 (the individual pits have not been excavated). This has led to the interpretation that the context of this site is marked by the transition from the collective ritual to the rise of forms of individuality in society, with all the strong implications that this process has in sociocultural terms. Structure 14 belongs to the second type, containing up to eight individuals, three of them still preserving several anatomical connections, accompanied by findings such as a bronze axe and several shell beads^194^.

Concerning to its geocultural context, this area has been considered peripheral to the stronger centres of the Iberian Megalithic Southwest (that comprises sites in mid-south Portugal and the Guadalquivir Valley, with sites like Valencina de la Concepción or Perdigoes) and the Eastern Spanish Chalcolithic and Early Bronze Age (cultures of Los Millares and el Argar). However, the geographical position of this site makes it especially significant, as it is not only placed between the two above-mentioned main southern centres/cultural complexes of the Iberian Prehistory, but less than 20 km from the North African shores. Also, the collective ritual and the hypogeic character of the burials links this site not only with its Iberian counterparts, but also with the whole Mediterranean Basin^196^, as opposed to the Northern and Central European contexts, dominated by the individual ritual^197^.

The monumentality of the site of Los Algarbes suggests that it functioned as a territory marker for the inhabiting communities, in a similar way as the abundant natural picks containing rock art spread all over in the Gibraltar Strait region.

Finally, we can conclude that, whether or not the local communities occupied a central or peripheral position in respect to the mentioned main cultural complexes of the Iberian Peninsula, this area represents a key bottleneck where influences from Western Iberia, the Mediterranean Basin and North African influences converge, creating a potential melting pot of great relevance for the understanding of transregional relations during Late Prehistory.

The samples from Los Algarbes necropolis included in this study come from:

Estructura 1-2. Corredor. 0014 (**CGG_2_100579**), indiv. A was 14C dated (CNA-5365) to 3340±30 BP, cal. BP 3684-3481 (see Suppl. Tab. S9); and also (Beta-564943) to 3300±30 BP, cal. BP 3636-3466 (see Suppl. Tab. S9).

Estructura 14 (ALG-2013) 14006. Cuadrìcula A. Indiv. 1 (**CGG_2_100581**), probably a female. If our individual 1 and individual A in Castañeda et al. 2022 are the same person, it is dated (CNA-2736) to 3961±35 BP^198^, cal. BP 4353-4293 (see Suppl. Tab. S9). The individual from Estructura 14. Hornacina 010. CuadrÌcula B (CGG_2_100584) was excluded because contaminated.

##### ***2.10.13. Terrera del Reloj, Dehesas de Guadix, Granada province, Andalusia***

Latitude: 38.49; Longitude: -3.43

Sample provider: Alfredo Mederos Martín, Victoria Peña Romo, Fernando Molina, Syvia Jiménez Brobeil

Alfredo Mederos Martín

The Argaric settlement of Terrera del Reloj is located on a prominent hill at 590 metres above sea level, controlling the confluence of the Fardes River into the Guadiana Menor River. After its identification during an inspection visit in 1980^199^, it was the object of an emergency excavation in 1983 as it was affected by a stone quarry and clandestine excavations. A stepped habitat was documented in 6 terraces where 17 tombs were located inside the habitat. The adult tombs all correspond to pits, with wooden planks attached to their walls, while the children's tombs are deposited in small ceramic urns or pithoi^200^. Human remains 6215 M8 (**CGG_2_023793**) correspond to bones from a possibly destroyed tomb on terrace 3, section 6.

##### ***2.10.14. Valencina de la Concepción, Calle Trabajadores 14-18, Seville, Andalusia***

Latitude: 37.417; Longitude: -6.076

Sample provider: Alfredo Mederos Martín, Victoria Peña Romo, Pedro López Aldana, Ana Pajuelo Pando

Alfredo Mederos Martín

The Copper Age settlement of Valencina de la Concepción is the largest in the Iberian Peninsula and also in Western and Central Europe (see also the description of Valencina de la Concepción, Calle Dinamarca 3 y 5, Sevilla, Andalusia, below). The settlement has an estimated size of c. 185-235 ha, while the adjacent necropolis extends between 162 and 233 ha^201^. In the necropolis there are large tombs in tholoi such as La Pastora, Matarrubilla^202^ or Montelirio^203^, discovered in 1860 and onwards, while the settlement was not identified until 1971^204^.

More than 130 rescue excavations have been carried out at site over the years^205^ and since 2014 there has been a systematic research project in the settlement^206,207^. Not all researchers would agree that this was a proper settlement and the hypothesis that it was only a large aggregation space for certain celebrations has been put forward as well^208^. The chronology of the settlement ranges between 3400 and 2250 BCE^206,209^, while the necropolis dates between 3200 and 2400 BCE^210^.

The excavation has also documented 336 Bell Beakers fragments, mainly of the so-called maritime style, and mostly from structure 136, which is 4 x 2.40 m large and 3.30 m deep.

Structure 1

In 2008, the rescue excavation of Calle Trabajadores was carried out, a sector with Bell Beaker occupation, where 30 structures excavated in the ground were documented^211^. The samples analysed in this project were obtained from structure 1 (a 4 m in diameter and only 0.30 m deep structure, which contained 9 human skulls together in UE 2). Five radiocarbon dates from the site indicate a chronology ranging between 2573 and 2297 BCE^208^, and thus clearly dates the structure to the Late Chalcolithic.

The samples from the 2008 structure 1 UE 2 are:

Skull, adult, pars petrosa (**CGG_2_023780/Rise 1955**)

Skull B, adult pars petrosa (**CGG_2_023781/Rise 1956**)

Skull associated with B (**CGG_2_023782/Rise 1956**)

Skull C, adult pars petrosa (**CGG_2_023783/Rise 1963**)

Skull K, adult, pars petrosa of the right temporal bone (**CGG_2_023786**)

Skull L, young adult, pars petrosa (**CGG_2_023787**) has been 14C dated (OxA-28244) to 3967±29 BP ^210^, cal. BP 4523-4298 (see Suppl. Tab. S9).

Skull M, adult, pars petrosa of the right temporal bone (**CGG_2_023788**)

**Tomb 6**

From the necropolis also individual 6 (Skull II) from tomb 6 was sampled; individual 6 (I) is an infant II between c. 7 and 8 years old. The *pars petrosa* of the left temporal bone was taken (**CGG_2_022341/Rise 1546**)

##### ***2.10.15. Valencina de la Concepción, Calle Dinamarca 3-5, Seville, Andalusia***

Latitude: 37.409835; Longitude: -6.076716

Sample provider: Jose Miguel Murillo León, Alfredo Mederos Martín, Victoria Peña Romo, Ana Pajuelo Pando, Pedro López Aldana

Jose Miguel Murillo León, Alfredo Mederos Martín

Since the discovering of the dolmen of La Pastora in 1860, the site of Valencina de la Concepción is one of the most emblematic of the Iberian Chalcolithic (see also the description of Valencina de la Concepción, Calle Trabajadores 14-18, Sevilla, Andalusia, above). However, it wasn´t until the decade of 1980 when extensive excavations started being carried on, mostly related with rescue archaeology^212^.

Comprising more than 400 ha, the site was divided in two or three sectors (habitational, productive, necropolis; however, this division is under debate), showing an enclosures division that is currently under study^207^. The site occupied a strategic position on the South-eastern border of a natural plateau in the palaeo-estuary of the Guadalquivir River, connecting the mouth of the river with the Northern ores of the Ossa-Morena mining district in South Western Iberia, from approximately 3000 BCE to the first half of the 2^nd^ millennium^210^.

While the character of this site is still under discussion (habitational, ceremonial, etc.), it is clear that its funerary sector had a long life. It shows a strong variability in terms of funerary structures, from megalithic tholoi to hypogea excavated in the subsoil, with different plant shapes. The burial ritual is dominated by the collective form, often subjected to remotions and reconditioning of the osteological remains. To this variability we have to add the abundant finding of isolated or partial human remains that are not restricted to the necropolis area, but spread all across the site^213^. All this suggests a strong social complexity that is still only partially understood.

In the considered habitational sector, the archaeological register is characterised by negative, sub circular structures excavated in the local clayish subsoil, showing different sized and, probably different purposes related both with occupation and storage.

This site is notorious not only for its extension or the relevance of its megalithic burials, but also due to the important amount of exotic materials found during the many excavations carried on since the 1980´s, like exotic rocks, amber or African and Asian elephant ivory.

The material under study belongs to a burial complex placed at the South of the site, consisting in three negative structure complexes (UE-5, UE-28, UE-51), not connected. The typology of those structures can be included in the hypogea that are spread across the Mediterranean Basin^196^. It represents a potential kinship group of a subsidiary class during approximately 100 years^214^.

The samples from this site come from the structure UE-5:

S-14, main chamber, UE 60, skull 30, pars petrosa (**CGG_2_100096**);

S-15, main chamber, UE 60, skull 33, par petrosa (**CGG_2_100097**);

S-16, main chamber, UE 60, skull 34, pars petrosa (**CGG_2_100098**);

S-17, main chamber, UE 60, skull 5, pars petrosa (**CGG_2_100099**), which has been 14C dated (OxA-28240) to 4221±30 BP^210^, cal. BP 4855-4628 (see Suppl. Tab. S9).

Also from structure UE-28:

S-35, UE 32, skull 11, *pars petrosa* (**CGG_2_100109**);

S-36, UE 32, skull 1, *pars petrosa* (**CGG_2_100110**), has been 14C dated three times (see Suppl. Tab. S9):

(SUERC-60398) to 4470±31BP^210^, cal. BP5289-4975

(OxA-32306) to 4423±31BP^210^, cal. BP 5275-4869.

(OxA-30336) to 4429±29BP^210^, cal. BP 5276-4874.

##### ***2.10.16. Cueva de la Carada, Huéscar, Granada, Andalusia***

Latitude: 37.79055556; Longitude: -2.53638889

Sample provider: Alfredo Mederos Martín, Victoria Peña Romo, Fernando Molina González, Sylvia Jiménez Brobeil

Alfredo Mederos Martín

Collective burial in an artificial cave 3 km from the town of Huéscar, which was the subject of an emergency excavation between April and July 1980 by Angela Mendoza and Leovigildo Sáez, although it remains unpublished^215,216^. No articulated skeletons were preserved. It shows an occupation during the Early and Middle Chalcolithic, c. 3000-2500 BCE. The sample (**CGG_2_023951**) was a third upper molar.

#### **2.11. Syria**

##### ***2.11.1. Tall Sūkās, Latakia Governorate***

Latitude: 35.306306; Longitude: 35.922194

Sample provider: Marie Louise Schjellerup Jørkov

Serena Sabatini

The Tall Sūkās (Zoukas) with great detail Henrik Thrane describes in the excavation reports^217^ how the work at the grave was not only demanding, but difficult and hampered by time limits. The author is therefore aware that some information might be missing, but also that the context was extremely complex to excavate and showed a high degree of post depositional activities, which included movement of both human remains and grave goods.

Three main layers were recognized in the tomb. There is evidence for 41 skeletons of which only 12 were more or less anatomical order. As the material was very difficult to make sense of, all bones and skulls were given individual numbers. Skeletons also got numbers. Skeletons 1-10 belonged to the first uppermost layer. After a layer consisting of sherds level two was identified. Skeletons 11 and 12 were deposited in this layer. Underneath those individuals there was a layer of sterile clay with a loose stone packing^217^. From this level 25 skulls were recovered.

The context is dated to the Syrian Middle Bronze Age, and no 14C dates are available for this material. Traditionally the West Syrian Middle Bronze Age is considered c. 2000-1600 BCE^218^.

Five individuals from this grave are analysed in this project. All of them come from the top layer of the grave^217^ Cranium 8 A5 (**CGG_2_101879**) belonging to a child of c. 11-12 years, Cranium 9 (with mandible 45) A7-15 (**CGG_2_101880**) belonging to a child of c. 8 years, Cranium 4 - Skeleton 2 (**CGG_2_101883**) belonging to an adult male (?) c. 35-40 years, Cranium 20 A10 (**CGG_2_101884**) belonging to a female of c. 25-35 years; an adult female is also Cranium 24, Skeleton VIII (I) A13 (**CGG_2_101885**), cranium and skeletons number for this individual are reported according to the data from the Anthropological Collection register; it is possible that change of numbers occurred at the time of registration as in the publication^217^ it is reported skull 24 belongs to skeleton I, which is an adult individual possibly female.

#### **2.12. Turkey**

##### ***2.12.1. Antandros, North-western Anatolia***

Latitude: 39,575833 Longitude: 26,790556

Sample provider: Gürcan Polat

Fulya Eylem Yediay

The Antandros necropolis is located west of Kaletasi Hill, Altinoluk, Balikesir province, also known as Troas region. Excavations at the site have been directed by Prof. Dr. Gürcan Polat and the Museum Kuva-yi Milliye since 2001; in total 558 tombs have been excavated, dated from the Late Geometric to Early Roman period.

**Grave M455**

The grave M455 was unearthed in 2014 in the southeast of Necropolis. The skeleton (**CGG_2_023618/Rise1901**) was found lying on the ground with an unguentarium vessel which belongs to Early Hellenistic period on the right shoulder and a bronze mirror on the left arm, a pair of gold Eros earrings at the level of ears, some bone objects on the legs and ribs, silver pendants, bronze ring, blue necklace bead and a bronze coin on pelvis. The skeleton was dated to 2304±29, cal. BP 2358-2179 (see Suppl. Tab. S9).

**Grave M520**

Grave M520 was excavated in 2017-2018. It is a so-called tile grave. The skeleton (**CGG_2_023623/Rise1906**), which was radiocarbon dated to 2290±32, cal. BP 2354-2157 (see Suppl. Tab. S9), belongs to an adult lying in an east-west direction, with the head looking east. One bronze coin and a piece of bronze were found under the jaw bone.

**Grave M524**

Grave M524 was excavated like M520 in 2017-2018. It is an inhumation grave without grave goods. The individual (**CGG_2_023624/Rise 1907**) was lying on the face with the head and the bent legs looking south; it has been radiocarbon dated 2184±31BP, cal. BP 2318-2073 (see Suppl. Tab. S9).

##### ***2.12.2. Resuloğlu, Çorum province***

Latitude: 40.426, Longitude: 34.193

Sample provider: İzzet Duyar, Derya Atamtürk, and Tayfun Yıldırım.

İzzet Duyar, Derya Atamtürk

The Resuloğlu cemetery and the nearby settlement are situated in the province of Çorum (Central Anatolia, Turkey) and dated to the Early Bronze Age (EBA III). The archaeological excavations started by Tayfun Yıldırım in 2003 and were completed in 2019. Over 200 graves have been unearthed during excavations in the necropolis area. Although the existence of different types of graves in the necropolis has been identified^32^, approximately ¼ of the skeletons were buried in pithos type tombs. The second most common burial type is the cist grave with a rate of 21.7%^219^.

Anthropological and osteological investigations of human skeletons recovered from the cemetery are ongoing. The preliminary analysis of 115 skeletons has been completed so far^219–221^. The series includes individuals of all ages. A significant part of the skeletons died in middle and/or young adulthood, and the number of subadult skeletons is 34 of the 80 skeletons whose sex was determined, 45 are female and 35 are male. 13 individuals were sampled for the present study (Table 10).

Paleopathological observations revealed that joint degenerations are quite common (52.5%) among Resuloğlu residents^219^. In addition, the presence of infectious and metabolic diseases has also been detected. Genetic originated disorders or anomalies such as sacralization and spina bifida are also seen in the population^219^. The most important point that draws attention in the examination of the Resuloğlu skeletons in terms of oral health is the high rate of dental calculus. Dental calculus developed in 970 (79.77%) of the 1216 teeth examined. The rate of tooth decay in the population is 3.74%, abscess is 2.34% and ante mortem tooth loss is 3.22%^220^.

| **Year of excavation** | **Grave nr./type/sample nr.** | **Burial gifts** |
| --- | --- | --- |
| 2003 | M2 (Pithos grave) (**CGG_2_022280**) | A white-black stone, carnelian necklace.  **14C:** 4036±36BP, cal. BP 4784-4416 (see Suppl. Tab. S9) |
| 2003 | M11 (Cist grave) (**CGG_2_022289**) | Two bronze pins, necklace’ beads made of faience, bronze, carnelian, frit |
| 2003 | M16 (Pithos grave) (**CGG_2_022276**) | A spouted red juglet.  **14C:** 4068±34BP, cal. BP 4798-4425 (see Suppl. Tab. S9) |
| 2003 | M20/1 (Cist grave) (**CGG_2_022291**) | A black slipped and polished vase, stone beads, a head of pin, a golden plate, a piece of broken bronze cup.  **14C:** 3860±36BP, cal. BP 4410-4154 (see Suppl. Tab. S9) |
| 2004 | M37 (Pithos grave) (**CGG_2_022284**) | No grave goods.  **14C:** 4172±36BP, cal. BP 4834-4580 (see Suppl. Tab. S9) |
| 2006 | M120 (Pithos grave) (**CGG_2_022281**) | Necklace' beads made of stone, frit, and bronze, two earrings, a bronze pin |
| 2006 | M148 (Pithos grave) (**CGG_2_022285**) | A bronze earring, a necklace made of carnelian, bronze, shells, and frit.  **14C:** 3844±33BP, cal. BP 4405-4151 (see Suppl. Tab. S9) |
| 2006 | M149 (Pithos grave) (**CGG_2_022288**) | No grave goods |
| 2006 | M151 (Pithos grave) (**CGG_2_022278**) | No grave goods |
| 2007 | M164 (Jar grave) (**CGG_2_022283**) | Frit necklace, two bronze pins |
| 2007 | M170 (Jar grave) (**CGG_2_022277**) | Two earrings, stone beads.  **14C:** 3852±33BP, cal. BP 4406-4153 (see Suppl. Tab. S9) |
| 2007 | M179/1 (Pithos grave) (**CGG_2_022279**) | Two bronze pins.  **14C:** 3864±37BP, cal. BP 4411-4155 (see Suppl. Tab. S9) |
| 2007 | M190/1 (Pithos grave) (**CGG_2_022282**) | Two bronze pins, a torque, a hair ring made of silver |

Table 10. Synoptic table with data about the sampled individuals from the cemetery at Resuloğlu and their context of origin.

##### ***2.12.3. Kaman-Kalehoyük Kazisi, Kırşehir province***

Latitude: 39.363392303714804; Longitude: 33.78664611534262

Sample provider: Sachihiro Omura

Fulya Eylem Yediay, Kimiyoshi Matsumura

Kaman-Kalehöyük is situated 3 km east of the city of Kaman in Kırşehir province, approximately 100 km southeast of Ankara. It is a circular tell mound with a diameter of 280 m^222^. The excavations in Kaman-Kalehöyük have been carried out by Dr. Sachihiro Omura since 1986 in collaboration with the Japanese Anatolian Archeology Institute.

The mound was in use for a long time; four main strata were excavated. According to the archaeological and architectural evidence from each stratum, the main levels are further subdivided into Stratum Ia and Ib - Ottoman period; Stratum IIa, IIc, and IId - from the Hellenistic period to the Early Iron Age; Stratum IIIa - Hittite Empire period; IIIb - Old Hittite period; IIIc - Karum (Assyrian Colony) period; Stratum IVa - Intermediate Period; Stratum IVb - Early Bronze Age.

The samples analysed in this project were found predominantly in stratum IIa, IIc, IIIb and IIIc.

**Kaman-Kalehoyük Stratum IIa (Late Iron Age) IIa period is dated from 700 BC to 200 BC (Early Hellenistic period)**

Sample HS 11-2 KL-110809 (**CGG_2_022174/Rise 1453**) comes from sector; N, VII, Grid; XXXII-54.55 (Q, R), XXXIII-54.55 (S, T), provisional layer 99 P3142. The sample was found in Sector VII in pit 3142. It belongs to a child skeleton found in the pit as decomposed.

**Kaman-Kalehoyük Stratum IIc (Middle Iron Age)**

Sample HS 12-2 KL-120731 (**CGG_2_022175/Rise 1454**), comes from Sector; N, VIII, provisional layer 192. It belongs to an infant skeleton found in pit 3171.

**Kaman-Kalehoyük Stratum IIIb (Middle Bronze Age: the Old Hittite period)**

Sample HS-03-01 KL 030809 (**CGG_2_022183/Rise1462**), comes from Sector; N, X, Grid; XXVII 54 (35), provisional layer 10. It belongs to an adult male in a prone burial alongside the Wall 2 (Hittite enclosure wall). This sample has been radiocarbon dated to 2250±30BP, cal. BP 2341-2153 (see Suppl. Tab. S9). Judging from the radiocarbon date, this individual would rather belong to strata IIa.

Sample KL 910717 (**CGG_2_022193/Rise1472**) comes from Strata III?, Sector; N, VI, west section. there is no provisional layer info. This sample has been radiocarbon dated to 2138±26 BP, cal. BP 2298-2001 (see Suppl. Tab. S9). Judging from the radiocarbon date, this individual would rather belong to strata IIa.

**Kaman-Kalehöyük Stratum IIIc (Middle Bronze Age: Karum period)**

Sample KL 940826 (**CGG_2_022180/Rise1459**) comes from Sector; N, I W4-W7, Grid; XLV-54 (GG). Four skeletons were found between wall4? and room153. Our sample belongs to Skeleton 3, who might be an outlier/foreigner..

Sample HS 11-3 KL 110816 (**CGG_2_022181/Rise 1460**) comes from Sector; N, VIII, Grid; XXX 54.55, XXXI 54.55, provisional layer P144. Three skulls were found in the burned layer (HS 11-03, HS 11-04, HS 11-05) in sector VII near room 390. Our sample comes from HS 11-03.

Sample HS-05-03 KL 050920 (**CGG_2_022182/Rise1461**) comes from Sector; N VIII, Grid; XXXI 55 (YY), provisional layer 91. It belongs to an adult male and was found in room 390. This sample has been radiocarbon dated to 3448±31BP, cal. BP 3830-3589 (see Suppl. Tab. S9).

Sample KL 940815 (**CGG_2_022188/Rise1467**), dated Strata IIIc, Sector; N, XII-0. Grid; XLVII 55 (LL), XLVIII 55 (PP), KL 940811, Sector N-XII, XLVIII-55 (PP) PL14, Sample S1. This sample has been radiocarbon dated to 3548±33 BP, cal. BP 3963-3718 (see Suppl. Tab. S9).

Sample KL 900719 (**CGG_2_022184/Rise1463**) comes from Strata IIIc?, Sector; N, III, Grid; XLI 54 (C), PL51. It belongs to a child skeleton. This sample has been radiocarbon dated to 3368±38BP, cal. BP 3694-3485 (see Suppl. Tab. S9).

##### ***2.12.4. Küllüoba Kazısı, Eskişehir province***

Latitude: 39.55665826645916, Longitude: 30.744497808841015

Sample provider: Murat Türkteki

Murat Türkteki, Fulya Eylem Yediay

The settlement of Küllüoba is located at the western end of the Sakarya Basin, northeast of the Eskişehir Province-Seyitgazi District. It has been excavated since 1996 under the direction of Prof. Turan Efe and now of Prof. Murat Türkteki. The geographical area where the settlement was established has highly fertile agricultural soils throughout its history. This geographical area is also a natural transportation route that can provide passage between Central Anatolia and the Marmara region. Founded 930 m above sea level on a slight elevation on the northern side of the Kireçkuyusu Creek, which is completely dried up today, the settlement has an area of 350 x 250 m and the thickness of the culture fill is 10 m. The mound, inhabited uninterruptedly for 1250 years between 3200-1950 BCE, includes all phases of the Early Bronze Age. It is an important settlement that provides information on the local urbanisation phenomenon^223,224^, the Early Bronze Age chronology of western Anatolia, and the trade between Syro-Cilicia and western Anatolia at the end of the Early Bronze Age^225^.

The cemetery area belonging to the beginning of the Early Bronze Age, which is the earliest phase of the settlement, was discovered in 2019 and it is the earliest known extramural cemetery area in western Anatolia so far^226^. The burials are of different types such as pithos, mudbrick cists, stone cists, and simple inhumations in hocker position.

**CGG_2_022156/Rise1435**, AI-AJ13/35, Küllüoba excavation. It was unearthed in 2014, Early Bronze Age I,pseudo-pithos type, hocker position burial. Dated to 4307±32BP, cal. BP 4960-4832 (see Suppl. Tab. S9). This sample is located within the boundaries of the EBA I cemetery area discovered in 2019.

**CGG_2_022159/Rise1438**, AA19/145. It was excavated in 2003 Early Bronze Age burial. Three skeletons were found (143, 144, 145).

Skeletons AA 19, 143-144 and 145 are the skeletons of an adult individual, one in the form of a simple inhumation and the other of a child in a pithos. Of these, skeleton AA19 143 belongs to an adult. Of the other examples, AA19 144 belongs to a child in a pithos. The skull of this skeleton was taken with the number AA19 145 and therefore actually belongs to specimen 144. All of the skeletons found at this site can be dated to the Middle Bronze Age Transition phase, in other words Late EB III. Both skeletons were buried in the hocker position. Although there is no architectural finding in the trench where the burials are found, it is understood that both burials belong to an intramural context, depending on the fact that the structures were found in both the previous and later phases in the mound.

**CGG_2_022160/Rise1439**, AC14/16. One skeleton was found on the north of sector AC14 with a deep bowl grave gift which is very characteristic of Early Bronze Age III period^227^. The sample was radiocarbon dated to 3936±32BP, cal. BP 4420-4238 (see Suppl. Tab. S9).

**CGG_2_022161/Rise1440**, AB13/20, radiocarbon dated to 3733±35BP, cal. BP 4231-3976 (see Suppl. Tab. S9). It is a skeleton found very close to the surface.

##### ***2.12.5. Keçiçayırı Kazısı, Eskişehir province***

Latitude: , Longitude: 39.320567, 30.768130

Sample provider: Murat Türkteki

Murat Türkteki, Fulya Eylem Yediay

Keçiçayırı is located in the mountainous southern part of Eskişehir province known as the Phrygian Plateau, 22 km south of Seyitgazi and 4.5 km southwest of Bardakçı village. The Bardakci Stream flowing through the village merges with the Seyit Stream, a tributary of the Sakarya River, in the north and then merges with the Eşen Stream coming from Yazılıkaya in the village of Bardakci. The valley splits into three bifurcations about 1 km before reaching the plain known as Keçiçayırı Mevkii in the southwest. The Eşen Stream crossing the plain and its tributaries from the south and north flow through these valleys. Between the Eşen Stream and its tributaries are two rocky hills called Cıbırada Hill and Aralıkada Hill, which border the plain from the east^228^.

The plain called Keçiçayırı Mevkii is surrounded by not very high mountains. The settlement area is mainly concentrated on the Cıbırada hill, the western slopes of this hill and the fields in the northeast of the plain. Early Neolithic, Early Bronze Age, Phrygian, Roman and Early Christian periods are represented here. Although the Phrygian period could not be clearly identified in terms of pottery during the excavations, a fibula, mentioned in the excavation reports but not yet published, is a typical Phrygian find from one of the destructions on the Cıbırada hill.

Another importance of the Cıbırada fortress is that it is understood that there were strong fortified castles on the rocks in the mountainous parts of the Eskişehir region. These fortresses may have been built for the exploitation and protection of the deposits of flint, marble and copper, as well as for the accommodation of merchants travelling between the regions and for the security of the road route^229^. It is also possible that the Cıbırada castle was used for burial structures in the Phrygian period.

**CGG_2_022162/Rise1441**, This sample is from Keçiçayırı Salvage excavation. During the excavations of the Early Bronze Age fortification on the hill identified as Cıbırada Hill in Keçiçayırı, the remains of a 2.5 x 1.5 m grave surrounded by stones were uncovered. AT4/8, was 14C dated to 2600±33BP, cal. BP 2775-2543 (see Suppl. Tab. S9).

201.	Mejías García, J. C. Formaciones sociales del III milenio A.N.E. en Valencina. (University of Sevilla, 2017).

202.	Collantes de Terán, F. El dolmen de Matarrubilla. in *Actas del V Symposium Internacional de Prehistoria Peninsular. Tartessos y sus Problemas (Jerez, 1968)* 47–61 (Francisco Collantes de Terán Barcelona, 1969).

203.	Fernández, A., García-Sanjuán, L. & Díaz-Zorita, M. Montelirio: un gran monumento megalítico de la Edad del Cobre. *Arqueología Monografías, Junta de Andalucía, Consejería de Cultura, Sevilla* (2016).

204.	Mata, D. R. & Martín, A. M. La primera campaña de excavación en el poblado calcolítico de Valencina de la Concepción (Sevilla). El corte estratigráfico 1, 1971. Fases del Calcolítico Inicial y Campaniforme. 55–70 (2020).

205.	Vargas Jiménez, J. M. *Elementos Para La Definición Territorial Del Yacimiento Prehistórico de Valencina de La Concepción*. (2003).

213.	Cruz-Auñón, R. & Mejías, J. C. *Diversidad de Prácticas Funerarias E Identidades En El Yacimiento de Valencina de La Concepción*. (Sevilla, 2013).

214.	Pando, A. P. & Aldana, P. M. L. La necrópolis de cuevas artificiales y fosas de c/ Dinamarca 3 y 5 (Valencina de la Concepción, Sevilla). *de Valencina de la Concepción (Sevilla …* 281–292 (2013).

215.	Mendoza Enguaras & Saéz Pérez, L. *Informe de Las Excavaciones En El Yacimiento Cueva Carada*. (Huéscar (Granada, 1980).

216.	Brobeil, J. *Estudio Antropológico de La Necrópolis de La Carada*. (Huéscar (Granada, 1983).

217.	Thrane, H. *Sūkās. 4. A Middle Bronze Age Collective Grave on Tall Sūkās*. (Munksgaard, 1978).

218.	Morandi Bonacossi, D. The Northern Levant (Syria) During the Middle Bronze Age. in *The Oxford Handbook of the Archaeology of the Levant: c. 8000-332 BC* (eds. Killebrew, A. E. & Steiner, M.) 414–433 (Oxford, 2014).

219.	Atamtürk, D. & Duyar, İ. Resuloğlu (Uğurludağ, Çorum) iskeletlerinin antropolojik analizi. *Arkeometri Sonuçları Toplantısı* **25**, 311–328 (2009).

220.	Atamtürk, D. & Duyar, İ. Resuloğlu erken tunç çağı topluluğunda ağız ve diş sağlığı. *Hacettepe Üniversitesi Edebiyat Fakültesi Dergisi* **27**, (2010).

221.	Duyar, İ. & Atamtürk, D. Erken Tunç Çağında Orta Anadolu’da ölü gömme âdeti, sağlık yapısı ve yaşam biçimi: Resuloğlu örneği. *Çorum Kazı ve Araştırmalar Sempozyumu* (2011).

222.	Sachihiro, O. Kaman-kalehöyük Excavations in Central Anatolia. in *The Oxford Handbook of Ancient Anatolia: (10,000-323 BCE)* (eds. McMahon, G. & Steadman, S.) 1095–1111 (Oxford University Press, 2011).

223.	Efe, T. Küllüoba and the initial stages of urbanism in Western Anatolia. in *From Primary Villages to Cities. Essays in Honour of U. Esin* (eds. Özdoğan, M., Hauptmann, H. & Başgelen, N.) 265–282 (2003).

224.	Efe, T., Türkteki, M., Şahoğlu, V. & Sotirakopoulou, P. Early Bronze Age Architecture in the Inland Western Anatolian Region. in *Across: the Cyclades and Western Anatolia during the 3rd Millennium BC* (eds. Sotirakopoulou, P. & Şahoğlu, V.) (2011).

225.	Türkteki, M., Sarı, D., Şahin, F., Türkteki, S. & Tuna, Y. Anadolu’da Bir İlk Tunç Çağı Kenti: Küllüoba Genel Değerlendirme ve 2020 Yılı Çalışmaları. *Lycus Dergisi* 105–128 (2021).

226.	Türkteki, M., Sarı D.,, Erdal, Y.S., Türkteki., S., Gündem,C.Y., Tuna,Y., Koruyucu,M.M., Delibaş,D., Dengiz, O. Küllüoba Kazısı 2022 Yılı Çalışmaları. in *43. Kazı Sonuçları Toplantısı* 295–305.

227.	Efe, T. Küllüoba’dan Çift Kulplu Fincan Formuna ait iki Örnek ve Düşündürdükleri. *Mustafa Büyükkolancı’ya Armağan/Essays in Honour of* (2015).

228.	Efe, T. & Türkteki, M. Keçiçayırı (Seyitgazi-Eskişehir) 2007 Yılı Kurtarma Kazıları. 75–84 (2007).

229.	Efe, T., Sarı, D. & Fidan, E. The significance of the Keçiçayırı excavations in the Prehistory of inland northwestern Anatolia. in *Archaeological Research in Western Central Anatolia (Third International Symposium of Archaeology)* 9–28 (2011).

1. Please note that while authors provided the full description of the various sites, all texts have been edited by Serena Sabatini with the contribution from Fulya Eylem Yediay and Fabrice Demeter. Editing included dating, calibration data, cross-reference to the whole work, revision of the references, partial rearrangement and development of the provided material and texts to create a uniform and consequent document.
